# Improved robustness to gene tree incompleteness, estimation errors, and systematic homology errors with weighted TREE-QMC

**DOI:** 10.1101/2024.09.27.615467

**Authors:** Yunheng Han, Erin K. Molloy

## Abstract

Summary methods are widely used to reconstruct species trees from gene trees while accounting for incomplete lineage sorting; however, it is increasingly recognized that their accuracy can be negatively impacted by incomplete and/or error-ridden gene trees. To address the latter, Zhang and Mirarab (2022) leverage gene tree branch lengths and support values to weight quartets within the popular summary method ASTRAL. Although these quartet weighting schemes improved the robustness of ASTRAL to gene tree estimation error, implementing the weighting schemes presented computational challenges, resulting in the authors abandoning ASTRAL’s original search algorithm (i.e., computing an exact solution within a constrained search space) in favor of search heuristics (i.e., hill climbing with nearest neighbor interchange moves from a starting tree constructed via randomized taxon addition). Here, we show that these quartet weighting schemes can be leveraged within the Quartet Max Cut framework of Snir and Rao (2010), with only a small increase in time complexity compared to the unweighted algorithm, which behaves more like a constant factor in our simulation study. Moreover, our new algorithm, implemented within the TREE-QMC software, was highly competitive with weighted ASTRAL, even outperforming it in terms of species tree accuracy on some challenging model conditions, such as large numbers of taxa. In comparing unweighted and weighted summary methods on two avian data sets, we found that weighting quartets by gene tree branch lengths improves their robustness to *systematic* homology errors and is as effective as removing the impacted taxa from individual gene trees or removing the impacted gene trees entirely. Lastly, our study revealed that TREE-QMC is highly robust to high rates of missing data and is promising as a supertree method. TREE-QMC is written in C++ and is publicly available on Github: https://github.com/molloy-lab/TREE-QMC

---

Over the last decade summary methods have been widely adopted in species tree estimation pipelines, starting from the identification of orthologus genomic regions from target capture sequencing data or assembled genomes (Steenwyk et al., 2023, 2024). Next, an evolutionary history for each region (referred to as a gene tree) is reconstructed typically in two steps: first a multiple sequence alignment (MSA) is built from the unaligned sequences and second a phylogeny is estimated from the MSA via maximum likelihood under standard models of molecular sequence evolution (e.g., the GTR model; Tavaré, 1986). Finally, the estimated gene trees are given as input to summary methods, many of which are designed to reconstruct the species tree while appropriately accounting for gene tree discordance due to incomplete lineage sorting (ILS), unlike concatenation (Roch and Steel, 2015). Examples of such summary methods include MP-EST (Liu et al., 2010) and STELAR (Islam et al., 2020), which are based on triplets implied by rooted gene trees, NJst (Liu and Yu, 2011) and ASTRID (Vachaspati and Warnow, 2015), which are based on pairwise internode distances implied by unrooted gene trees, and ASTRAL (Mirarab et al., 2014) and TREE-QMC (Han and Molloy, 2023), which are based on quartets implied by unrooted gene trees.

Despite the wide spread use of summary methods, it is increasingly recognized that their accuracy depends on gene tree quality. In simulations, summary methods typically decrease in accuracy as gene tree estimation error (GTEE) or incompleteness increases (Xi et al., 2015; Mirarab and Warnow, 2015; Molloy and Warnow, 2018; Nute et al., 2018). Moreover, a growing number of systematic studies show that homology errors can impact species tree estimates produced by different summary methods (Simmons et al., 2022; Springer and Gatesy, 2024).

The dominant approach for combating poor quality gene trees is *gene tree filtering*, in which entire gene trees are removed from the data set typically based on some threshold for the proportion of missing taxa (Hosner et al., 2016; Cunha et al., 2021) and/or proxies for GTEE (Simmons et al., 2016). However, simulation studies have suggested that filtering can reduce the accuracy of summary methods and that only gene trees with very high GTEE should be removed entirely (Molloy and Warnow, 2018). This finding motivates the use of less aggressive approaches, for example contracting very low support branches in gene trees (Zhang et al., 2018; Simmons and Gatesy, 2021) and/or removing specific taxa from individual gene trees. Taxa are typically removed based on the fraction of gapped sites in the gene MSA (referred to as fragmentary missing data by Sayyari et al., 2017 and the One Thousand Plant Transcriptomes Initiative, 2019) or based on (long) outlier branch lengths in the gene tree (Mai and Mirarab, 2018; One Thousand Plant Transcriptomes Initiative, 2019). Critically, these approaches require substantial effort and decision-making on the part of researchers, who must choose how to quantify error as well as select thresholds for filtering gene trees, contracting branches, and/or removing individual taxa. Often a range of parameter settings are explored, leading to many different estimates of the species tree (e.g., Hosner et al., 2016).

To address GTEE, Zhang and Mirarab (2022) proposed to leverage gene tree branch lengths and support values within the popular summary method ASTRAL. ASTRAL seeks a species tree that maximizes the number of quartets (unrooted four-leaf trees) implied by the input gene trees. This optimization problem is NP-hard (Lafond and Scornavacca, 2019), so ASTRAL solves the problem within a constrained search space built from the input gene trees (Mirarab et al., 2014; Mirarab and Warnow, 2015; Zhang et al., 2018). The goal of weighted ASTRAL is to *down-weight quartets with low support on the internal branch* (an indicator that the quartet cannot be confidently resolved due to insufficient phylogenetic signal) and/or *long lengths on the terminal branches* (an indicator that the quartet may be impacted by long branch attraction). However, these quartet weighting schemes increase the complexity of ASTRAL’s original search algorithm. To achieve greater computational efficiency, weighted ASTRAL employs heuristic search, specifically hill climbing with nearest neighbor interchange moves from a starting tree constructed via randomized taxon addition. This new heuristic, often referred to as ASTER, has been found to be more robust to incomplete gene trees than the original ASTRAL algorithm (Zhang and Mirarab, 2022; Morel et al., 2023).

Following the success of weighted ASTRAL/ASTER, Liu and Warnow (2023) incorporated weighting schemes based on gene tree branch lengths or support values but not into the distance method ASTRID. Overall, leveraging gene tree branch lengths and support values is a promising path forward but weighted summary methods are still in their infancy. Comparatively little is known about how they perform on real data, and there are significant challenges to developing efficient algorithms.

In this paper, we introduce a new weighted summary method, called weighted TREE-QMC, which integrates the quartet weighting schemes of Zhang and Mirarab (2022) into the Quartet Max Cut framework (QMC) of Snir and Rao (2010) while also efficiently addressing the issue of quartet weight normalization for incomplete gene trees. A previous challenge in using QMC as a summary method was that it required researchers to explicitly extract quartets from gene trees prior to species tree estimation; we recently addressed this issue, introducing TREE-QMC Han and Molloy (2023). Weighting quartets based on gene tree branch lengths and support values presents computational challenges, and perhaps surprisingly, we achieve an algorithm with only a small increase in time complexity and no increase in storage complexity compared to the unweighted TREE-QMC method. Moreover, our simulation study showed that weighted TREE-QMC was fast and highly competitive with weighted ASTRAL/ASTER in terms of species tree accuracy, even outperforming weighted ASTRAL/ASTER on some challenging conditions, such as large numbers of taxa.

We then evaluated weighted summary methods on three recently published data sets: one for plants with high rates of missing taxa (Morel et al., 2023) and two for birds, *Palaeognathae* (Cloutier et al., 2019) and *Neognathae* (Wu et al., 2024a), both of which have been found to contain homology errors (Simmons et al., 2022; Springer and Gatesy, 2024). The homology errors for Palaeognathae were biased towards two species (Chicken and White-Throated Tinamou), which clustered together off a long branch in the 105 affected ultraconserved element (UCE) gene trees (Simmons et al., 2022). We found that TREE-QMC and ASTRAL/ASETER with hybrid or length weighted recovered the same species tree topology as the unweighted methods *after removing* Chicken and White-Throated Tinamou from the 105 affected UCE trees or *after filtering* the 105 affected UCE trees entirely. Otherwise, an alternative resolution of the focal branch was favored by the unweighted summary methods. To evaluate whether biased homology errors had a systematic impact on the reconstructed species tree, we implemented Partitioned Coalsecence Support (PCS) (Gatesy et al., 2017, 2019) within TREE-QMC and applied it to the focal branch in Palaeognathae. The PCS analysis showed that the 105 UCE trees with homology errors consistently supported an alternative resolution of the focal branch; however, the length and hybrid weighting schemes mitigated this issue. Quartet weighting schemes did not have a big impact on TREE-QMC for the other avian data set, where homology errors were not clearly biased towards particular species. For this data set and the plant data set, TREE-QMC produced trees that were closer to the concatenation tree and reference taxonomies than those produced by ASTRAL/ASTER, which produced in a highly imbalanced (i.e., caterpillar-like) branching pattern not seen in the other estimated species trees. Overall, our results suggest that TREE-QMC is efficient, easy-to-use, and improves robustness to gene tree incompleteness, estimation error, and homology errors.

## Materials and Methods I. Weighted TREE-QMC

### Overview of unweighted TREE-QMC

To introduce weighted TREE-QMC, we briefly review the original (unweighted) method (Han and Molloy, 2023), which leverages the Quartet Max Cut (QMC) framework (Snir and Rao, 2010, 2012; Avni et al., 2014). The idea behind QMC is to reconstruct the species tree in a divide-and-conquer fashion from a set of input quartets. At each step of the divide phase, a graph is built with vertices representing taxa. The edges between vertices (taxa) are weighted by the relevant input quartets. A cut of the **quartet graph** partitions the taxa into two disjoint subsets, corresponding to a bipartition (associated with an internal branch) in the unrooted species tree. The algorithm continues by recursion on subproblems defined by the taxa on each side of the bipartition.

Termination occurs when there are three or fewer taxa in the subproblem as the solution (i.e., a phylogenetic tree on the taxon set) is trivial (Fig. 6a in Appendix A). Importantly, **artificial taxa** are added to subproblems at each step in the divide phase to represent the taxa on the other side of the bipartition (Fig. 6a in Appendix A). During the conquer phase, subproblem trees are connected at artificial taxa, ultimately producing an unrooted, binary species tree on the complete set of taxa (Fig. S1 in Han and Molloy, 2023).

The original (unweighted) TREE-QMC method improves upon QMC in two ways. First, QMC operates on quartets, which must be extracted from gene trees prior to species tree estimation. Given *k* gene trees on *n* taxa, the preprocessing phase alone has time complexity Ω(*n*^4^*k*) and storage complexity Ω(*n*^4^) (note that we do not include the storage required for input gene trees). In contrast, TREE-QMC builds the quartet graph directly from the input gene trees, having time complexity *O*(*n*^3^*k*), with some assumptions on subproblem decomposition, and storage complexity 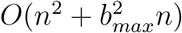, where *b*_*max*_ is the maximum number of artificial taxa for any subproblem. To summarize, the time complexity of TREE-QMC is lower than the quartet extraction step required for QMC, and the reduction in storage complexity should further improve efficiency by reducing cache misses. More importantly, our experimental evaluation showed TREE-QMC scaled to large data sets (with 1000 taxa and 1000 genes on which we could not run QMC) and achieved faster runtimes than ASTRAL-III, the leading method at the time (Fig. 2B in Molloy et al., 2022).

Second, TREE-QMC normalizes quartet weights with respect to artificial taxa so that each gene tree gets one vote for every subset of four taxa present in the tree, although the vote could be split across the three possible quartets when gene trees are multi-labeled by artificial taxa introduced from the QMC framework, multiple gene copies (Legried et al., 2021), or multiple individuals/alleles per species (Rabiee et al., 2019). TREE-QMC can be run without normalization (**n0**), with uniform normalization (**n1**), or with non-uniform normalization (**n2**), which leverages the structure implied by the introduction of artificial taxa to *downweight quartets that are less relevant for resolving a particular subproblem (i.e*., *internal branch in the species tree)* (Fig. 6a–d in Appendix A). In our simulation study, normalization was critical for accurate species tree estimation under challenging model conditions, such as high mean GTEE, with the n2 normalization scheme yielding the best accuracy (Fig. 4A in Molloy et al., 2022). Notably, TREE-QMC (with n2 normalization) outperformed ASTRAL-III (Zhang et al., 2018) on challenging model conditions, ranging from large numbers of taxa (200–1000) to very high levels of ILS (60%) and mean GTEE (60%) (Figs. 2 and 4A in Han and Molloy, 2023, respectively).

Based on the promising performance, we were motivated to further evaluate and improve upon the TREE-QMC method, especially with respect to gene tree estimation error and incompleteness, which are well-established obstacles to phylogenomics.

### Normalizing Quartet Graph for Incomplete Gene Trees

Our first improvement addresses the issue of incomplete gene trees, specifically their impact on quartet weight normalization. As previously mentioned, we say the weights are normalized if each gene tree votes once for each subset of four taxa present in the tree. We show that the normalization factors derived for the complete taxon set do not constitute a valid normalization scheme for incomplete gene trees (Fig. 6d–f in Appendix A), and **sharing** these normalization factors across all gene trees reduces species tree accuracy, as we show in our simulation study. In Appendix A, we introduce an efficient algorithm for computing the correct normalization values for each gene tree on the fly. This new approach does not increase the storage or time complexity of quartet graph construction and reduces the number of memory allocations and thus cache misses compared to earlier versions of the TREE-QMC code. Lastly, TREE-QMC now outputs a polytomy when there are no quartets for the subproblem; this can occur when the majority of taxa are missing from gene trees, for example the real and synthetic data sets assembled by Morel et al. (2023), which we use for benchmarking.

### Weighting Quartet Graph based on Gene Tree Branch Lengths and Support Values

Our second improvement addresses the issue of GTEE by leveraging the quartet weighting schemes of Zhang and Mirarab (2022).

Recall that the vertices in the quartet graph represent taxa from the subproblem and edges between taxa are weighted based on quartets implied by the input gene trees, with leaves mapped to taxa by the function *L*. Then, each quartet *q* = *x, y*|*z, w* implied by some gene tree *T* contributes two **bad edges** between each of the sibling pairs (i.e., (*L*(*x*), *L*(*y*)) and *L*(*z*), *L*(*w*)) as well as four **good edges** between each of the non-sibling pairs (i.e., (*L*(*x*), *L*(*z*)), (*L*(*x*), *L*(*w*)), (*L*(*y*), *L*(*z*)) and (*L*(*y*), *L*(*w*))), provided that the leaves are uniquely labeled (i.e., *L*(*x*) ≠ *L*(*y*) ≠ *L*(*z*) ≠ *L*(*w*)). In the unweighted algorithm, all good and bad edges have weight one; otherwise, good and bad edges inherit the weight of the quartet that contributed them. In the **length, support**, and **hybrid weighting schemes** of Zhang and Mirarab (2022), the weights of the quartet *q* implied by gene tree *T* are defined as

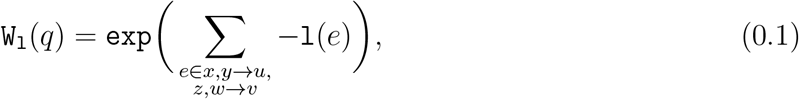

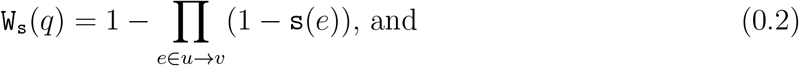

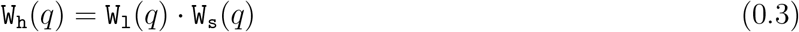

respectively, where `l`(*e*) denotes the length of branch *e* in gene tree *T*, `s`(*e*) denotes the support value of branch *e* in *T*, and *u* and *v* denote the **anchor** vertices of quartet *q* = *x, y*|*z, w* in *T*, with *u* closer to sibling pair *x, y* and *v* closer to sibling pair *z, w*. Note that *u* → *v* is the set of edges on the path between *u* and *v*, which corresponds to the **internal branch** of quartet *q* implied by *T* (Fig. 1a–b; also see Fig. 1 in Zhang and Mirarab, 2022). Similarly, *x, y* → *u* and *z, w* → *v* is the set of edges on the paths from each of the four leaf vertices to their respective anchor, which corresponds to the four terminal branches of quartet *q* implied by *T*.

**Fig. 1.**
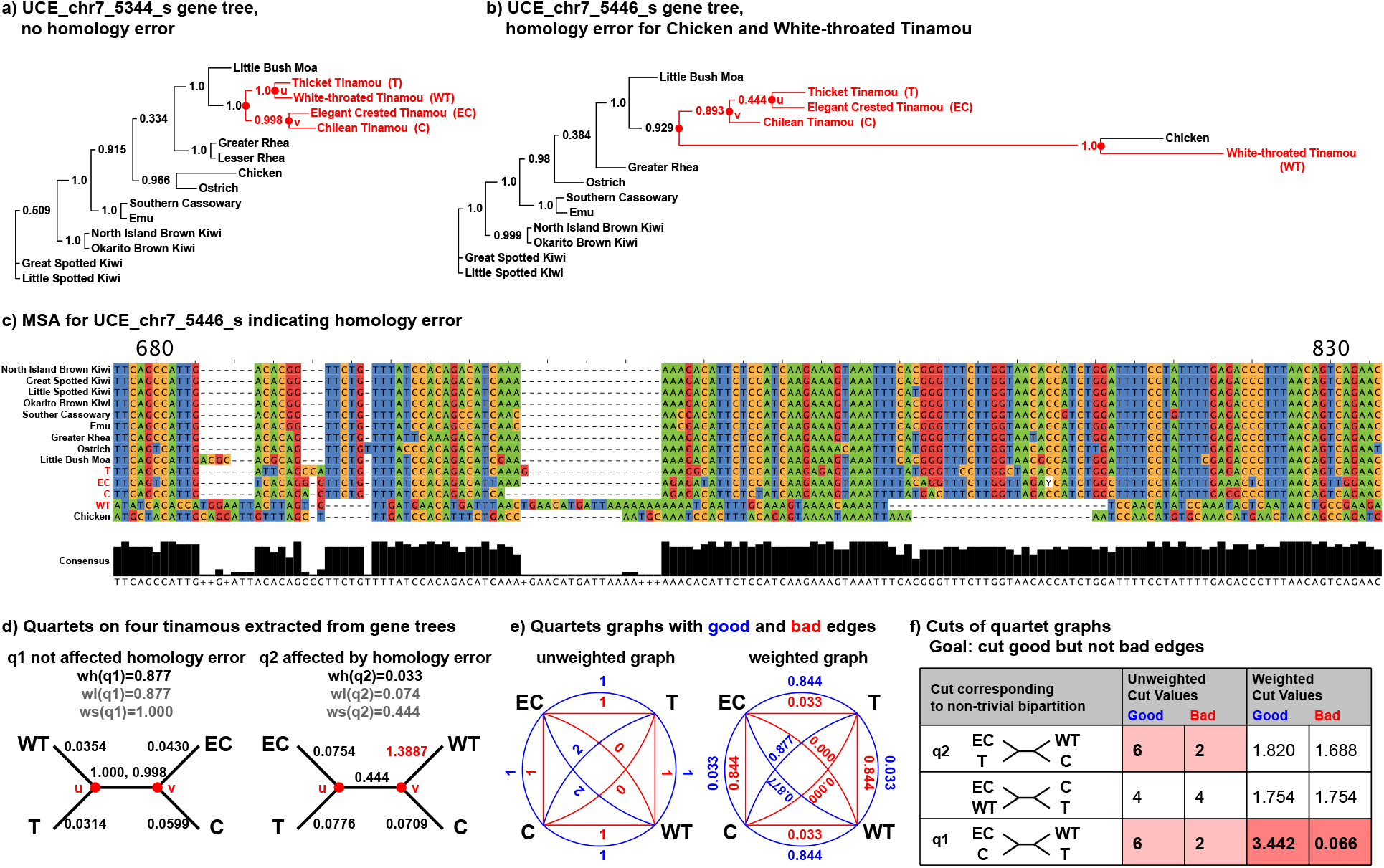
Impact of hybrid weights on quartet graph and its best cut. **a)** Gene tree estimated on UCE chr7 5344 s by Cloutier et al. (2019), where *u* and *v* denote the anchor vertices for the quartet on the four Tinamou species: Thicket Tinamou (T), White-throated Tinamou (WT), Elegant Crested Tinamou (EC), and Chilean Tinamou (C). **b)** Gene tree estimated on UCE chr7 5446 s with long branch to Chicken and White-Throated Tinamou. **c)** MSA for UCE chr7 5446 s suggests homology errors for Chicken and White-Throated Tinamou, as noted by Simmons et al. (2022) (visualized with JalView; Waterhouse et al., 2009). d) Quartets *q*1 and *q*2 on the four Tinamous species extracted from gene trees and chr7 5344 s and chr7 5446 s, respectively, along with their hybrid, length, and support weights. **e)** Quartet graphs built from both *q*1 and *q*2 without and with hybrid weighting. **f)** A cut of the quartet graph partitions the species into two disjoint sets. There are only three cuts that yield non-trivial bipartitions on four species, corresponding to quartets. Cut quality is related to the ratio of good and bad edges removed (shown in table); the goal is to remove good edges but not bad edges thus preserving sibling relationships (Snir and Rao, 2010). For the unweighted graph, there are two best cuts (highlighted). For the weighted graph, there is one best cut corresponding to *q*1 (highlighted), which seems reasonable as *q*2 is affected by homology errors.

Figure 1 shows the impact of quartet weighting schemes on the quartet graph constructed by TREE-QMC, taking two gene trees from the Palaeognathae data set (Cloutier et al., 2019) as input, one with and one without identified homology errors (Simmons et al., 2022). For simplicity, we examine the quartet graph on the four Tinamou species, finding that the three weighting schemes successfully down-weight the quartet implied by the gene tree with homology errors compared to the gene tree without homology errors, which in turn impacts the best cut(s).

The naive way to construct the quartet graph is to extract all weighted quartets from the gene trees (Fig. 1 in the Supplementary Materials); however, this preprocessing step is computationally intensive and requires a large amount of storage even for unweighted quartets, as previously mentioned. The challenge is building the quartet graph efficiently (i.e., directly from gene trees without extracting all possible quartets) while weighting quartets based on gene tree branch lengths and support values. In Appendix B, we introduce efficient algorithms to construct the weighted quartet graph for a subproblem directly from gene trees in *O*((*a*^2^*b* + *ab*^2^*n* + *b*^3^*n*)*k*) time and *O*(*a*^2^ + *ab* + *b*^2^*n*) space, where *a* is the number of non-artificial taxa from the subproblem (referred to as **singletons**), *b* is the number of artificial taxa from the subproblem, and *n* is the total number of species, which we assume corresponds to the maximum number of leaves in any gene tree for simplicity (Theorem 4 in the Supplementary Materials).

### New features in weighted TREE-QMC

Lastly, we implement several new features within weighted TREE-QMC. *Bioconda*. TREE-QMC is now installable via bioconda.

#### Multi-labeled and non-binary gene trees

TREE-QMC now extends its framework for handling artificial taxa to accommodate multi-labeled gene trees as well as non-binary gene trees being given as input.

#### Characters and “BP” mode

TREE-QMC now allows character data to be given as input; see the `--chars` and `--char2tree` options. TREE-QMC treats each character (i.e., site) in the data set as an unrooted tree with internal branches separating taxa assigned the same state from all other taxa. This functionality is useful because quartet-based summary methods are statistically consistent estimators of species trees for binary (0/1) characters evolving under the neutral Wright-Fisher model (Mendes and Hahn, 2017; Molloy et al., 2022) even when there is error and missingness provided it is unbiased (Han and Molloy, 2024). Springer et al., 2020 refer to this approach as summary methods in “bipartition mode” or “BP mode”. If the `--bp` option is used with TREE-QMC instead of the `--chars` option, branch lengths in coalescent units will be computed under the neutral Wright-Fisher model with a fast maximum likelihood estimator, assuming a constant mutation rate (Molloy et al., 2022); otherwise, branch lengths are not computed for character data.

#### Quadrapartition Quartet Support (QQS)

TREE-QMC now enables users to output **Quadrapartition Quartet Support (QQS)** for the three quartet topologies “around” each internal branch (quadripartition) for unweighted or weighted quartets (Sayyari and Mirarab, 2016; Mirarab et al., 2024); see the `--support` or `--supportonly` options.

Additionally, maximum likelihood branch lengths in coalescent units are reported, along with the effective number (**EN**) of gene trees that provide information (i.e., quartets) for resolving the branch. Typically a branch is considered well supported when EN is sufficiently large and discordance follows signatures of ILS (i.e., the QQS value correspond to the branch in the estimated species tree is higher than the QQS values for the two alternative branch resolutions, which should be roughly equal to each other). Note that QQS and EN values are used by ASTRAL to compute branch support as the local posterior probability (Local PP) (Sayyari and Mirarab, 2016).

#### Partitioned Coalescence Support (PCS)

TREE-QMC now enables users to compute **Partitioned Coalescence Support (PCS)** for a specified focal branch in an input species tree (Gatesy et al., 2017, 2019). Specifically, the `--pcsonly` option returns the QQS values for each gene tree for each of the three possible resolutions of the focal branch in the species tree, enabling users to identify outlier gene trees that strongly or weakly favor a particular resolution of the focal branch compared to other gene trees. If applied to characters or gene trees sorted by their position in the genome, the output QQS plots can highlight interesting regions of the genome, for example a region of suppressed recombination in the avian tree of life (Fig. 2 in Mirarab et al., 2024) as well as enrichment for deep coalescence at the MHC locus in primates (Fig. PanGenomeS1 in Yoo et al., 2024).

**Fig. 2.**
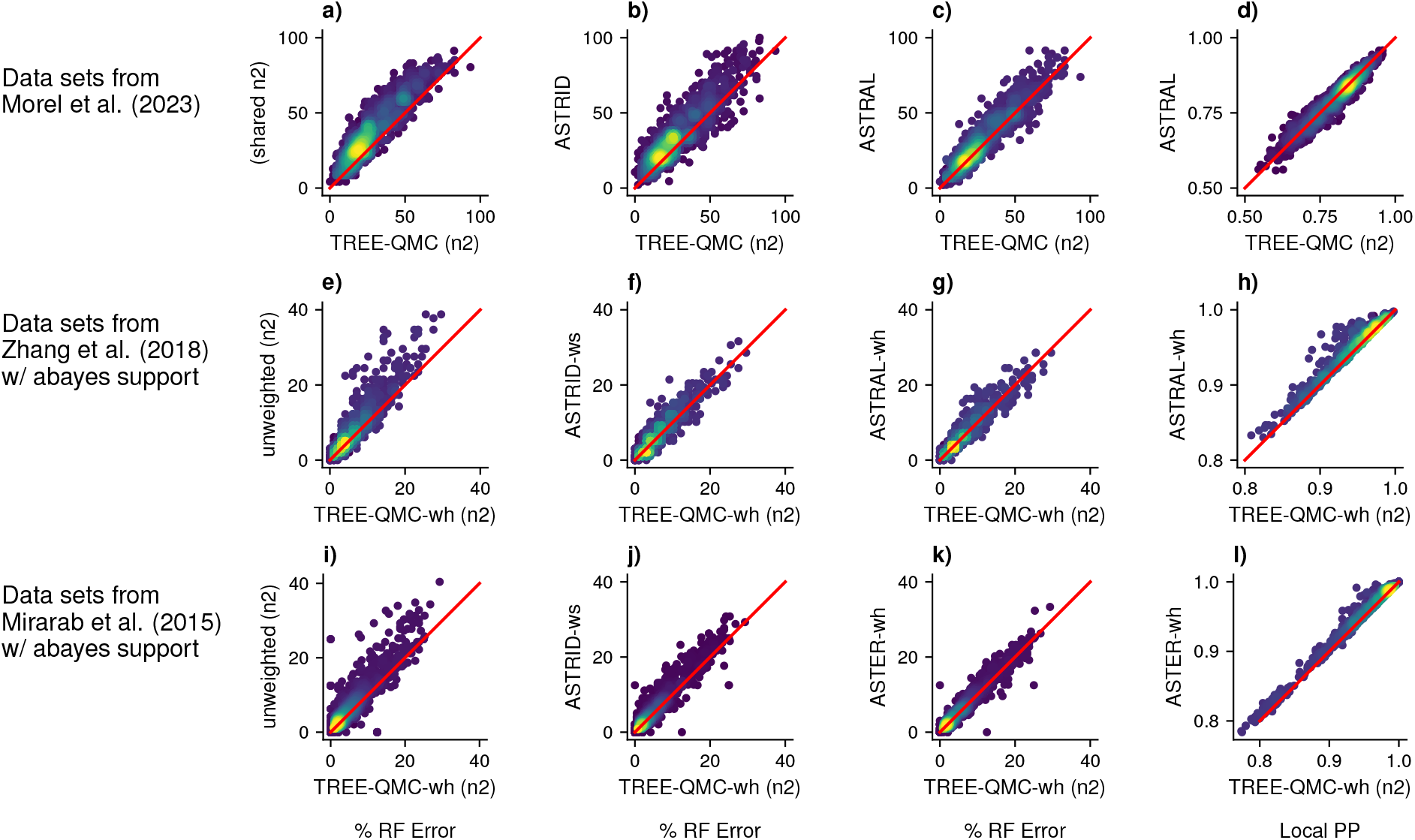
Rows of subplots show results on data sets simulated from three prior studies. Each dot in the scatter plot is a data set, with its position based on the evaluation metric (indicated below columns of subplots) for the method on the *x*-axis and the method on the *y*-axis. Red lines indicate there is no difference between methods. Data points are colored based on density, with lighter colors indicating higher density and darker colors indicating lower density. For % RF error, higher density of data points above the red line indicates that the version of TREE-QMC on the *x*-axis achieves lower error (and thus is more accurate) than the method on the *y*-axis.

## Materials and Methods II. Simulation Study

We now describe our performance study for evaluating the utility of weighted TREE-QMC on simulated data sets from prior studies focusing on gene tree estimation error and incompleteness, specifically the ASTRAL-II (Mirarab and Warnow, 2015), ASTRAL-III (Zhang et al., 2018), and Asteroid (Morel et al., 2023) data sets. Scripts are available at https://github.com/molloy-lab/tree-qmc-study/tree/main/han2024wtreeqmc.

### Species Tree Estimation Methods

A large number of summary methods have been developed over the last decade. To make the simulation study manageable, we focused on **weighted summary methods**: weighted ASTRAL (i.e., ASTER) and weighted ASTRID. We also included Asteroid (Morel et al., 2023), as it is similar to (unweighted) ASTRID but with improvements for incomplete gene trees. All summary methods were run in default mode with the parameter settings summarized below. Detailed repository information, version/commit numbers, and software commands are available in the Supplementary Materials.

For the Asteroid data sets, summary methods were run in unweighted mode, as the estimated gene trees did not include branch support and no alignments were available.

Additionally, the Asteroid data sets had high rates of missing taxa, so we ran TREE-QMC on simulated data sets with and without updating the (n2) normalization values for incomplete gene trees (Fig. 6). The latter is denoted as **n2 shared** to indicate the same normalization values are shared across all gene trees regardless of their taxon sets.

For the ASTRAL-II/III data sets, summary methods were run with their best weighting option. We ran ASTRAL with hybrid weights (denoted **ASTER-wh**), as this weighting scheme yielded the best accuracy in Zhang and Mirarab (2022). Motivated by these prior results, we ran TREE-QMC with hybrid weights (denoted **TREE-QMC-wh** or **TQMC-wh**) as well as in unweighted mode as an comparison. ASTRID does not implement the hybrid weighting scheme, and thus we ran ASTRID with support weights (denoted **ASTRID-ws**), as this weighting scheme yielded the best accuracy in Liu and Warnow (2023). **Asteroid** does not implement any weighting schemes; thus we ran Asteroid in default (unweighted) mode. For all weighted summary methods, we set the minimum/maximum values for gene tree branch support values to 0.333/1 for data sets with abayes support and 0/100 for data sets with bootstrap support, as indicated in the user manuals. On the smaller (i.e., Asteroid and ASTRAL-III) simulated data sets, we ran TREE-QMC with each of its three quartet weight normalization schemes: **n0** (none), **n1** (uniform), and **n2** (non-uniform). The non-uniform (n2) yielded the best accuracy, so we used it for the larger (i.e., ASTRAL-II) simulated data sets and for biological analyses.

### Simulated Data Sets

We evaluated summary methods using 3719 data sets simulated in three prior studies. These simulations were conducted by (1) simulating species trees under the Yule model (Yule, 1925), (2) simulating gene trees within the species tree under the Multi-Species Coalescent (MSC) (Rannala and Yang, 2003), which models ILS, and (3) simulating sequences down gene trees under standard models of molecular sequence evolution (e.g., the GTR model; Tavaré, 1986), which produces an MSA because there are no insertions or deletions. Different model conditions were created by varying a one or two model parameters at at time while keeping the others fixed. Multiple replicate data sets were simulated for each model condition. To enable comparisons of model conditions across studies, we empirically evaluated ILS, GTEE, and missingness (MISS) for each data set, reporting the average values across data sets with the same model condition. **ILS** was evaluated as the normalized RF distance between the true species tree and true gene trees, averaged across all gene trees. **GTEE** was evaluated as the normalized RF distance between each true and estimated gene tree, averaged across all gene trees. **MISS** was evaluated as the fraction of missing taxa from each gene tree, averaged across all gene trees. We summarize the model conditions and empirical properties below.

#### Asteroid simulated data sets

The Asteroid data sets were simulated by Morel et al. (2023) to evaluate Asteroid in the context of missing data. After deleting sequences from gene MSAs, they estimated gene trees with maximum likelihood (ML) under the GTR+GAMMA4 model (Yang, 1993) using ParGenes (Morel et al., 2018). The empirical properties of the Asteroid data sets are given in Table S1 in the Supplementary Materials. Overall, the data sets were characterized by very high missingness (0.76–0.84), very low ILS (0.05–0.08), and medium GTEE (0.30–0.45), with some exceptions depending on the model parameter being varied. Model conditions (50 replicates each) were simulated by varying the

1. **effective population size** and thus ILS level: 10 (ILS: 0), **50 million (default; ILS: 0.06)**, 100 million (ILS: 0.11), 500 million (ILS: 0.39), and 1 billion (ILS: 0.56)
2. **number of taxa**: 25, **50 (default)**, 75, 100, 125, 150
3. **number of genes**: 250, 500, **1000 (default)**, and 2000
4. **sequence length** and thus GTEE level: 50 bp (GTEE: 46), **100 bp (GTEE: 0.35; default)**, 200 bp (GTEE: 0.25), and 500 bp (GTEE: 0.16)
5. **gene tree branch length scalar** and thus GTEE level: 0.05 (GTEE: 0.55), 0.1 (GTEE: 0.47), **1 (GTEE: 0.35; default)**, 10 (GTEE: 0.42), 100 (GTEE: 0.70), and 200 (GTEE: 0.77)
6. **missingness parameter**: 0.5 (MISS: 0.74), 0.55 (MISS: 0.78), **0.6 (MISS: 0.81; default)**, 0.65 (MISS: 0.84), 0.7 (MISS: 0.86), and 0.75 (MISS: 0.88)

where default indicates the value used in all other model conditions. There are 26 unique model conditions, after removing duplicates from the default parameters, and 1300 replicate data sets in total.

#### ASTRAL-III (S100) simulated data sets

The ASTRAL-III datasets with 100 taxa were simulated by Zhang et al. (2018) to evaluate ASTRAL-III in the context of GTEE. They estimated gene trees with maximum likelihood under the GTR+GAMMA model using FastTree-2 (Price et al., 2010). After which, branch support was estimated via bootstrapping with 100 replicates; later Zhang and Mirarab (2022) estimated abayes branch support (Anisimova et al., 2011) with IQ-TREE-2 (Minh et al., 2020). The empirical properties of the ASTRAL-III (S100) data sets are given in Table S2 in the Supplementary Materials. Overall, the data sets were characterized by medium-high ILS (0.46). Four model conditions (50 replicates each) were generated by varying the sequence length and thus the GTEE level: 200 bp (GTEE: 0.56), 400 bp (GTEE: 0.42), 800 bp (GTEE: 0.31), and 1600 bp (GTEE: 0.23). The number of genes given to methods as input varied from 50, 200, 500, and 1000, yielding 16 model conditions and 800 replicate data sets in total.

#### ASTRAL-II simulated data sets

The ASTER-II data sets (no missingness) were simulated by Mirarab and Warnow (2015). We estimated abayes branch support for these data sets (see Supplementary Materials for software command), following the recommendation by Zhang and Mirarab (2022). The empirical properties of the ASTRAL-III data sets are in Table S3 of the Supplementary Materials. Model conditions (50 replicates each) were simulated by varying the species tree height and speciation rate (deep/shallow) with a fixed number of taxa (200), which in turn varied ILS and GTEE. At the shortest species tree height (called 0.5*×*), the ILS and GTEE were 0.68/0.69 and 0.44/0.44, respectively. At the longest species tree height (called 5*×*), the ILS and GTEE were 0.09/0.21 and 0.28/0.21, respectively. Conversely, the speciation rate and species tree height was fixed and the number of taxa was varied: 50, 100, 200, 500, 1000 (ILS: 0.31–0.35; GTEE: 0.26–0.30). The number of genes given to methods as input varied from 50, 200, and 1000 genes, yielding 33 model conditions and 1640 replicate data sets (note that 33 data sets were excluded based on criteria described in the Supplementary Materials).

### Evaluation Metrics

We compared summary methods in terms of species tree error, runtime, quartet score, and mean branch support.

#### Species tree error

We reported the normalized Robinson-Foulds (RF) error rate (Robinson and Foulds, 1981), which is equivalent to the false negative (FN) or false positve (FP) error rates when both the true and estimated species trees were binary (note that the FP/FN error rate are defined as the number of branches in the estimated/true species tree that are missing from the true/estimated species tree divided by the number of branches in the estimated/true species tree). All methods returned binary trees, except TREE-QMC, which returned non-binary trees for the Asteroid data sets when there were no quartets for resolving one or more subproblems. To ensure the RF distance was comparable to the other methods, we refined polytomies arbitrary. Additionally, we reported false positive (FP) and false negative (FN) error rates separately in the Supplementary Materials.

For each model condition (with replicate data sets simulated under the same conditions), we tested for significant differences between some pairs of methods using two-sided, paired Wilcoxon signed-rank tests, as implemented in the R coin library. To be conservative, we corrected for ties between methods in two different ways: the Wilcoxon method (Wilcoxon, 1949) and the Pratt method (Pratt, 1959), taking the higher of the two *p*-values. Lastly, we corrected for multiple comparisons based on Bonferroni correction, dividing the *p*-value by the number of tests performed for the experiment (note that we treated data sets simulated for the ASTRAL-II, ASTRAL-III (S100), and Asteroid studies as three different experiments). If the *p*-value was less than 0.05 after Bonferroni correction, we said there was a **significant difference** between two methods for the model condition.

#### Runtime

We recorded the wall clock time (i.e., the total time for the method to run on an input data set from start to finish). When collecting runtime data, we gave all methods exclusive access to the compute node (architecture: AMD EPYC-7313 with 32 CPUs; maximum RAM: 64 GB). ASTRAL/ASTER did not complete within our maximum wall clock time of 20 hours for three data sets when given a single-thread, so we also ran it with 16 threads. Otherwise, methods were run with a single thread. Branch support calculations were turned off to enable fair runtime comparisons.

#### Quartet Score and Branch Support Values

For the quartet-based methods (i.e., TREE-QMC and ASTRAL/ASTER), we reported the total number of quartets in the input gene trees that were satisfied by each estimated species tree as well as mean branch support. Branch support was computed using ASTRAL’s Local PP (Sayyari and Mirarab, 2016), which is based on QQS values from either unweighted or weighted quartets. Both total quartet score and mean branch support were computed with the same quartet weighting scheme as was used for species tree estimation.

## Results on Simulated Data Sets

We now report the results of benchmarking (weighted) summary methods on three collections of data sets simulated for prior studies.

### Asteroid Data Sets

The Asteroid data sets were used to evaluate unweighted summary methods under very high levels of missing taxa (see Figure 2a–c as well as Figures S2–S7 and Tables S4–S11 in the Supplementary Materials). We found that TREE-QMC (n2), without shared normalization factors, achieved lowered species tree (RF) error in 86% of data sets compared to TREE-QMC (n2 shared) (Fig. 2a). Moreover, TREE-QMC (n2) achieved significantly lower FN and FP error than no normalization (n0) for 17 and 16 of the 26 model conditions, respectively (Table S11). There were no significant differences between non-uniform (n2) and uniform (n1) normalization (Table S10); thus, we compared TREE-QMC (n2) against the other summary methods.

Asteroid was the only method that achieved significantly lower FN error than TREE-QMC (n2), which occurred for 3 of the 26 model conditions (Table S6). Asteroid was not significantly better than TREE-QMC when considering FP error. The two methods tied on 20% of data sets in terms of RF error. Asteroid was better on 46% of data sets and TREE-QMC (n2) was better on the remaining 34%.

In comparison to ASTRID and ASTRAL/ASTER, TREE-QMC (n2) outperformed or tied with them in terms of RF error on more than 75% of data sets (it was better on 74% and 62% of data sets and tied on 9% and 14% of data sets, respectively) (Fig. 2b–c). TREE-QMC (n2) achieved significantly lower FN and FP error compared to ASTRID for 18 of the 26 model conditions (Table S6–S8). Although TREE-QMC (n2) achieved significantly lower FP error than ASTRAL/ASTER on 17 of the 26 model conditions, respectively (Table S6–S8), this dropped to 3 model conditions when considering at FN error. Interestingly, TREE-QMC and ASTRAL/ASTER performed similarly in terms of mean branch support (Fig. 2d), but ASTRAL/ASTER achieved a higher total quartet score on 97% of data sets.

#### ASTRAL-III (S100) data sets

The ASTRAL-III (S100) data sets were used to evaluate weighted summary methods under varying levels of GTEE (see Figure 2d–f as well as Figure S8 and Tables S12–S15 in the Supplementary Materials). We compared weighted methods given gene trees with abayes support values, as bootstrap support values resulted in worse species tree accuracy for all methods, as previously observed by Zhang and Mirarab (2022). We found that TREE-QMC-wh (n2) lowered or tied for species tree (RF) error on 57% and 27% of data sets, compared to TREE-QMC (n2), without hybrid weighting (Fig. 2e). Moreover, TREE-QMC-wh (n2) typically achieved the lowest mean species tree (RF) error per model condition compared to TREE-QMC’s other normalization schemes as well as the other summary methods (Fig. S8 and Table S12). TREE-QMC-wh (n2) achieved significantly lower RF error than ASTRID-ws and ASTRAL/ASTER-wh in 4 and 3 of the 16 model conditions, respectively (Tables S13–S14), whereas ASTRID-ws and ASTRAL/ASTER-wh were not significantly better than TREE-QMC-wh (n2) for any model conditions. Overall, TREE-QMC-wh (n2) outperformed or tied with ASTRID-ws and ASTRAL/ASTER-wh on more than 75% of data sets (it was better on 53% and 44% of data sets and tied on 26% and 35% of data sets, respectively) (Fig. 2f–g). However, TREE-QMC-wh (n2) never achieved a higher quartet score than ASTRAL/ASTER-wh and had better mean branch support on just 5% of data sets (Fig. 2h). Note Asteroid should perform the same as (unweighted) ASTRID, as there are no missing taxa and that ASTRID performed worse than ASTRID-ws on these data sets in the study by Liu and Warnow (2023).

#### ASTRAL-II data sets

The ASTRAL-II data sets were used to evaluate weighted summary methods under varying levels of ILS and numbers of taxa (see Figure 2g–h as well as Figures S9–S10 and Tables S16–S23 in the Supplementary Materials). Similar to S100 data sets, we found that TREE-QMC-wh (n2) lowered or tied for species tree (RF) error of TREE-QMC (n2) in 58% and 24% of data sets, compared to TREE-QMC (n2), without hybrid weighting (Fig. 2i). In comparison to ASTRID-ws and ASTRAL/ASTER-wh, TREE-QMC-wh (n2) outperformed or tied with them on more than 75% of data sets (it was better on 54% and 46% of data sets and it tied on 30% and 37%) (Fig. 2j–k). Moreover, TREE-QMC (n2) achieved significantly lower RF error than ASTRID-ws and ASTRAL/ASTER-wh on 16 and 10 of the 33 model conditions, respectively (Tables S19–S20 and S22–S23). The difference between these pairs of methods was the most pronounced for model conditions with 500 and 1000 taxa, where TREE-QMC had a clear advantage. However, TREE-QMC-wh (n2) only achieved a higher quartet score than ASTRAL/ASTER-wh on just 1 of the 1619 data sets and had better mean branch support on just 5% of data sets (Fig. 2l).

Mirarab and Warnow (2015) also published the concatenation (CA-ML) trees (except the 1000-taxon, 1000-gene model condition). In comparison to CA-ML TREE-QMC (n2) was significantly better in terms of RF error on 19 of the 33 model conditions (Tables S18 and S21). The only condition CA-ML had a notable advantage over TREE-QMC (n2) was when the species tree height was longer (in generations) and speciation occurred closer towards the root (Table S18), although it is worth noting that the ILS level for these data sets was only 9%.

#### Runtime

Lastly, we evaluated the runtime of summary methods on the ASTRAL-II data sets with varying numbers of taxa and 1000 gene trees. At 1000 taxa, the fastest method was ASTRID-ws, followed by ASTRAL/ASTER-wh (16 threads), TREE-QMC (n2), TREE-QMC-wh (n2), and then ASTRAL/ASTER-wh (1 thread) at 0.0, 0.35, 0.36, 1.24, and 8.00 hours, respectively (Fig. 3a). As the number of taxa increased from 100 to 1000 (so by a factor of 10), the runtime ratio between the weighted and unweighted TREE-QMC algorithms only increased from 2.9 to 3.4 (so by a factor of 1.2) (Fig. 3b). Thus, although TREE-QMC-wh (n2) runtime was slower than the original (i.e., unweighted) algorithm, it still scaled to large numbers of taxa, where it had the greatest advantage in terms of species tree accuracy compared to ASTRID-ws and ASTRAL/ASTER-wh.

**Fig. 3.**
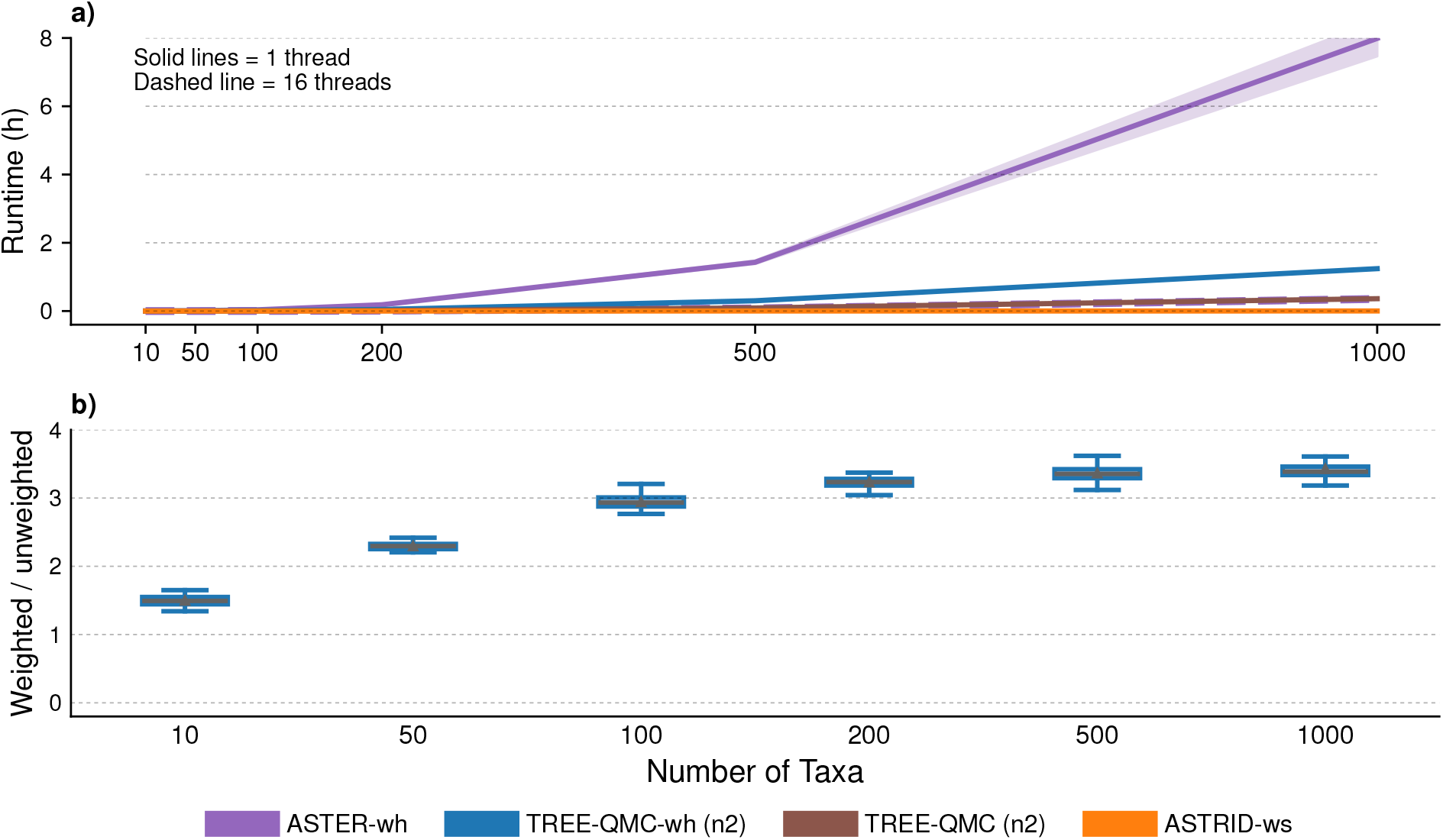
Runtime for ASTRAL-II data with varying numbers of taxa and 1000 gene trees with abayes support. Subfigure (a) shows the average runtime in hours for each of summary method (shaded region is standard error). Note that unweighted TREE-QMC (n2) is directly beneath ASTER-wh (16 threads). Subfigure (b) shows the ratio between the runtime for hybrid weighted TREE-QMC (n2) and the original TREE-QMC (n2) method.

**Fig. 4.**
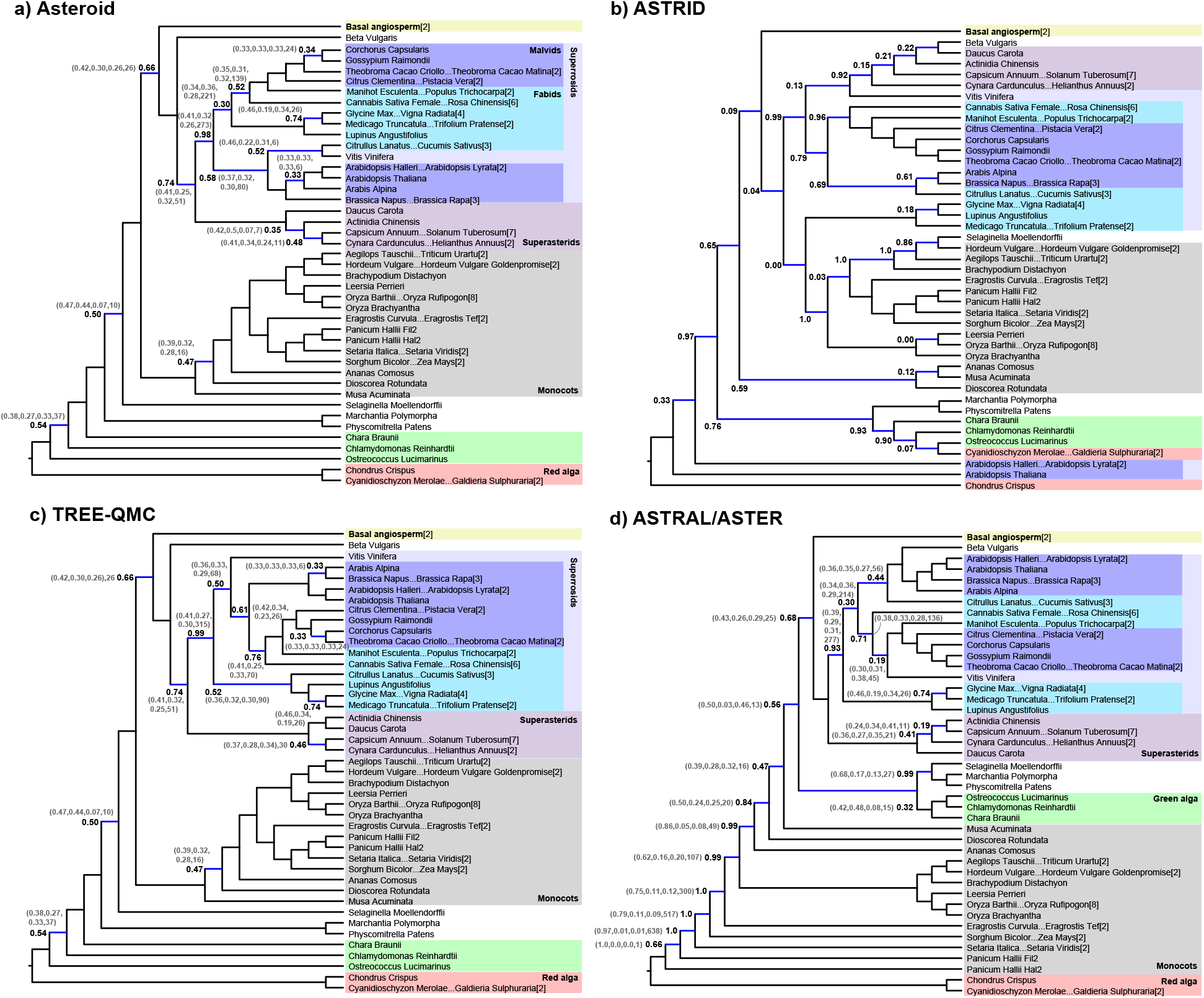
Species trees estimated from data curated by Morel et al. (2023). Subfigures (a), (b), (c), and (d) show species trees we estimated using Asteroid, ASTRID, TREE-QMC, ASTRAL/ASTER, respectively. Blue branches are not in the RAxML tree from Morel et al. (2023). Branch support (i.e., Local PP) is shown for branches that disagree with the RAxML tree. QQS values and EN are given in parentheses. All species trees are shown on the same reduced the leaf set created based on subtrees in the RAxML that were shared by all estimated species trees (leaf labels have the form *X*…*Y* [*N*] indicating a subtree in the RAxML tree with *N* taxa, starting taxon *X* and ending with taxon *Y*). The reduction in leaf labels from 81 to 43 indicates that the estimated trees are on many of the lower order relationships.

## Biological Analyses

We now describe our re-analysis of three published biological data sets using summary methods Asteroid, ASTRID, TREE-QMC, and ASTRAL/ASTER. Software commands are similar to those used in the simulation study. Scripts used to analyze biological data sets are available at https://github.com/molloy-lab/tree-qmc-study/tree/main/han2024wtreeqmc.

### Plant data set from Morel et al., (2023)

We first re-analyzed the 81-taxa, 6176-gene plant data set curated by Morel et al. (2023) to evaluate their method Asteroid. This data set was characterized by extreme missing data, with the majority of gene trees have 9 or fewer taxa. Specifically, the number of gene trees with 30–35 taxa (MISS: 57–63%), 20-29 taxa (MISS: 64–75%), 10-19 taxa (MISS: 77–88%), and 4–9 taxa (MISS: 89–95%) was 4, 95, 490, and 5587, respectively. Overall, the mean percentage of missing taxa across the 6176 gene trees was 95%. The high rates of incompleteness make the plant data set more similar to a supertree analysis (rather than a coalescent analysis) so we ran summary methods in unweighted mode only. After species tree estimation, we estimated branch support using ASTRAL’s Local PP (Sayyari and Mirarab, 2016).

We compared the estimated species trees to the (non-binary) NCBI reference taxonomy as well as the (binary) concatenation tree from Morel et al. (2023), which was missing 7 out of 56 (12.5%) internal branches in the NCBI reference tree. Overall, Asteroid, TREE-QMC, ASTRAL/ASTER, and ASTRID returned binary trees missing 14 (18%), 15 (19%), 20 (26%), and 29 (37%) of branches in the concatenation tree, respectively. In comparison to the reference tree, Asteroid, TREE-QMC, ASTRAL/ASTER, and ASTRID were missing 6 (12.5%), 6 (12.15%), 17 (30%), and 24 (43%) branches. All methods recovered Basal Angiosperm, although only two taxa were sampled from this group. All methods but ASTRID recovered Superastrids and Red Alga. Asteroid and TREE-QMC recovered Monocots as well as Superrosids, unlike ASTRAL/ASTER. Only ASTRAL/ASTER recovered Green Alga but this clade was separated from Red Alga by all of Monocots, which formed a highly imbalanced (caterpillar-like) topology. The branching ordered for Monocots differed from all other methods but was highly supported (Local PP), although the effective number (EN) of gene trees with information around some of these branch was quite low (e.g., 1–49), in which case ASTRAL recommends ignoring them. The ASTRAL/ASTER tree satisfied 3790, 4815, and 26141 more quartets than TREE-QMC, Asteroid, and ASTRID trees (note that the fraction of quartets satisfied was above 0.89 for all methods).

### Avian (Neognathae+Palaeognathae) data set from Wu et al., (2024)

Next, we re-analyzed the avian data set curated by Wu et al. (2024a), which included 5756 coding sequences (CDs), 4871 introns, and 2384 intergenic segments for 124 avian species. Later, homology errors in CDs were identified by Springer and Gatesy (2024), although these errors did not appear biased towards any particular set of taxa (Fig. 1B in Springer and Gatesy, 2024 and Fig. 1A in Wu et al., 2024b). We used IQ-TREE-2 (version 2.3.5) to compute abayes support on the best maximum likelihood (ML) gene trees estimated by Wu et al. (2024a) and then estimated species trees using Asteroid, ASTRID, TREE-QMC, and ASTRAL/ASTER (with branch support calculations were turned off to enable fair runtime comparisons). Species tree estimation was restricted to the 10,627 CDs and introns, as these markers may be more robust to orthology errors than intergenic regions (personal communication with Dr. Mark Springer and Dr. John Gatesy). All three weighted summary methods were run with and without their different weighting schemes. For unweighted analyses only, we also filtered the input gene trees by removing outlier taxa with TreeShrink version 1.3.9 (options: -m “per-species” -q 0.05) (Mai and Mirarab, 2018). We confirmed that TreeShrink successfully removed the homology errors from the CDs highlighted in Figure 1B in Springer and Gatesy (2016). After species tree estimation, we estimated branch support as Local PP using the same quartet weighting scheme that was used for species tree estimation (Sayyari and Mirarab, 2016). For weighted methods, we used the original gene trees when computing Local PP and QQS values; for unweighted methods only, we used gene trees with outlier taxa removed.

#### Comparison of unweighted versus weighted summary methods

ASTRAL/ASTER produced four species trees: one tree for hybrid weighting, one tree for length weighting, one tree for support weighting OR no weighting after taxon filtering, and one tree for no weighting before taxon filtering. The ASTRAL/ASTER with length weighting differed from unweighted ASTRAL/ASTER after taxon filtering (Fig. S11d in Supplementary Materials) by just one internal branch. Similarly, unweighted ASTRAL/ASTER (after taxon filtering) differened from ASTRAL/ASTER with hybrid weighting (Fig. 5d) by just three branches, but this difference jumped to six branches when taxon filtering was not used for the unweighted method. In contrast, TREE-QMC produced two species trees: one tree for hybrid and length weighting (Fig. 5c) and one tree for all other ways of running TREE-QMC (Fig. S11b in Supplementary Materials). The two TREE-QMC trees differed from each other on just three branches. ASTRID also produced two species trees: one tree for length weighting and one tree for all other ways of running ASTRID (Fig. 5b). The two ASTRID trees differed on 18 internal branches, with the length weighting tree being much farther from the NJst tree (thus we did not consider the length weighting ASTRID tree in further comparisons). Lastly, Asteroid recovered the same tree as ASTRID if taxon filtering was used; otherwise, the Asteroid tree differed from the ASTRID tree by four branches.

**Fig. 5.**
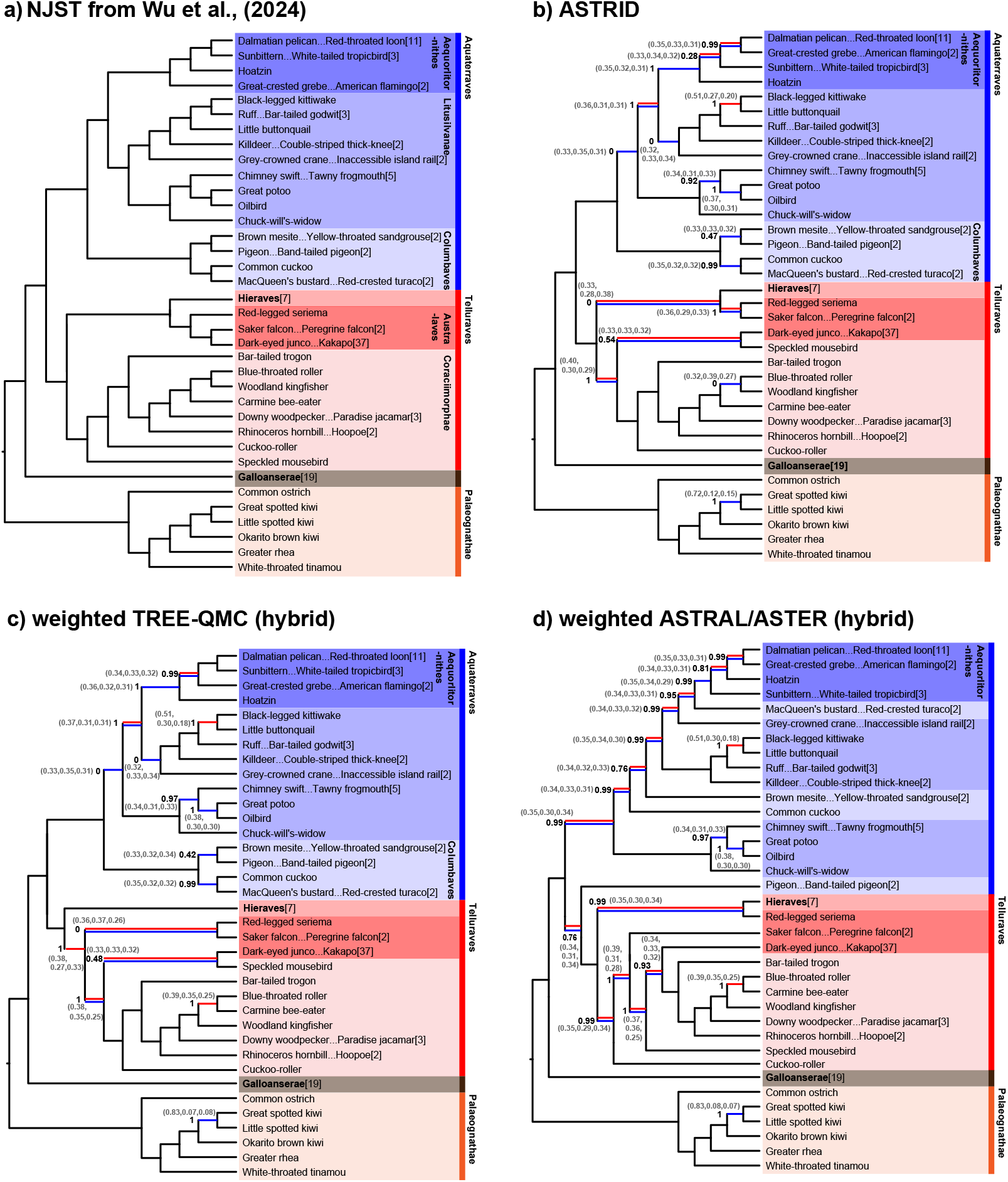
Species trees estimated from data curated by Wu et al. (2024a). Subfigure (a) is the NJst tree from Figure 2 in Wu et al. (2024a). Subfigure (b), (c), and (d) show species trees we estimated on CDs and introns using ASTRID (no weighting and support weighting scheme), weighted TREE-QMC (hybrid or length weighting scheme), weighted ASTRAL (hybrid weighting scheme), respectively. Red and blue branches are not in the NJst and RAxML trees from Wu et al. (2024a), respectively. Branch support (i.e., Local PP) is shown for branches that disagree with the NJst and/or RAxML trees. QQS values (in parentheses) are based on unweighted quartets (with taxon filtering) for subfigure (b) and hybrid weighted quartets (without taxon filtering) for subfigures (c) and (d). All species trees are shown on the same reduced leaf set created based on subtrees in the NJst that were shared by all estimated species trees. Leaf labels have the form *X*…*Y* [*N*] indicating the subtree in the NJst tree has *N* taxa, starting taxon *X* and ending with taxon *Y*. The reduction in leaf labels from 124 to 36 indicates that the estimated trees agree on most of the lower order relationships.

#### Comparison to published species trees

Wu et al. (2024a) published two main species trees: one from applying NJst (Liu and Yu, 2011) (Fig. 5a and Fig. 2 in Cloutier et al., 2019) and one from applying RAxML (Stamatakis, 2014). The NJst and RAxML trees, which differed by 13 internal branches, were given different collection inputs than our methods, as they were given all three marker types (CDs, introns, and intergenetic regions) after removing 30% outlier genes (i.e., genes with the highest quartet distance between the best ML gene tree and the estimated species tree were removed). Our estimated species trees differed from the NJst (and RAxML) trees by 8 (16), 8 (13), and 16 (18) branches for ASTRID/Asteroid, TREE-QMC-wh (n2), and weighted ASTRAL/ASTER-wh, respectively. Thus, the TREE-QMC tree was closer to NJst and RAxML trees than the ASTRAL/ASTER tree by 8 and 5 branches, respectively.

When considering major clades, all methods tested in our study recovered *Palaeognathae/Neognathae, Galloanserae/Neoaves, Aequorlitornithes*, and *Hieraves*. ASTRAL/ASTER did not recover *Aquaterraves* (because it clustered the two pigeons into Telluraves) or *Columbaves*, unlike ASTRID and TREE-QMC. No methods recovered *Listusilvae, Coraciiorphae*, or *Australaves*, but the ASTRID and TREE-QMC trees required fewer edit moves (either contractions/refinement moves or subtree prune and regraft moves) to achieve monophyly for these groups than the ASTRAL/ASTER tree. On the other hand, ASTRAL/ASTER always achieved the highest quartet score and its branches that disagreed with NJst or RAxML had high branch support (Local PP), although the QQS values around these branches were all close to one third, which is consistent with a rapid radiation for Neoaves, as demonstrated by many prior studies (Jarvis et al., 2014; Wu et al., 2024a; Stiller et al., 2024). Lastly, ASTRAL/ASTER recovered a highly imbalanced (i.e., caterpillar-like) branching pattern for *Aquaterraves* minus the two pigeons, which was striking compared to the other estimated species trees.

#### Runtime

ASTRID completed in just a few seconds, weighted TREE-QMC completed in 11 minutes, weighted ASTRAL/ASTER-wh completed in 2 hours, and ASTRAL/ASTER completed in 2.6 hours (both with and without taxon filtering the input using TreeShrink). The wall clock time of ASTRAL/ASTER dropped to 5–8 minutes when using 16 threads instead of one thread.

### Avian (Palaeognathae) data set from Cloutier et al., (2019)

Lastly, we re-analyzed the avian (Palaeognathae) data set curated by Cloutier et al. (2019), with 3,158 ultraconserved elements (UCEs) for 15 species. Systematic homology errors were identified in 105 of the UCEs by Simmons et al. (2022), with the two impacted taxa, White-Throated Tinamou and Chicken, clustering together. We used IQ-TREE-2 to compute abayes support on the best maximum likelihood (ML) gene trees estimated by Cloutier et al. (2019) and then estimated species trees with the four summary methods, comparing weighting schemes as well as filtering practices for unweighted methods. All analyses produced one of two trees, called “A” and “B,” which differed by a single focal branch (quadrapartition). In tree A, the focal branch split Tinamous plus Kiwis+Cassowary+Emu from Rheas plus Ostrich+Chicken, whereas in tree B, Tinamous swapped with Rheas so they were on the same side of the focal branch as Chicken (Fig. S12 in Supplementary Materials). Notably, the 105 UCEs with homology errors strongly favored tree B, but the hybrid quartet weighting scheme reduced the magnitude of support for tree B, as shown by partitioned coalescent support (PCS) analysis around the focal branch (Fig. S13 in Supplementary Materials).

Weighted TREE-QMC and ASTRAL (hybrid and length) returned tree A, as did unweighted TREE-QMC and ASTRAL after filtering the UCE data set (either removing the two impacted taxa from the 105 gene trees with homology errors or removing the gene trees entirely). Otherwise, tree B was returned. Branch support (i.e., Local PP) for the two resolutions of the focal branch was low (*<*0.5; see Fig. S12). Filtering taxa or entire gene trees resulted in tree A satisfying 2507 and 2269 more quartets than tree B respectively.

Likewise, tree A satisfied 972.5 more quartets than tree B when using hybrid weights; otherwise it satisfied 1783 fewer quartets. The concatenation trees from UCEs estimated by Cloutier et al. (2019) and Springer and Gatesy (2016) (who removed homology errors) resolved the focal branch in the same way as tree A but differed from tree A on one branch, placing Emu and Cassowary as sister to Tinamous instead of Kiwis.

## Discussion

Recently, Zhang and Mirarab (2022) introduced quartet weighting schemes to improve the robustness of the popular method ASTRAL to gene tree estimation error. Although their quartet weighting schemes are straightforward to describe, they present challenges for developing efficient summary methods. Here, we show that these weighting schemes can be effectively integrated into the Quartet Max Cut framework of Snir and Rao (2010), introducing weighted TREE-QMC. Our theoretical and empirical study shows that weighted TREE-QMC is a highly promising approach and may even be the leading weighted summary method developed to date.

### Scalability

Weighted TREE-QMC only increases the time complexity of quartet graph construction by a factor of *b* compared to the original (unweighted) algorithm, where *b* = *O*(*n*) is the number of artificial taxa for the subproblem (see proof of Theorem 1 in Han and Molloy, 2023). Interestingly, the empirical runtime of weighted TREE-QMC did not grow much compared in our computational experiments (Fig. 3b). We conjecture this difference between our theoretical and empirical study is due to the number *b* of artificial taxa per subproblem growing sub-linearly in the number *n* species or behaving more like a constant factor in practice. Under the latter assumption, both weighted and unweighted TREE-QMC have storage complexity *O*(*n*^2^) and time complexity *O*(*n*^2^ log(*n*)*k*) with some assumptions on subproblem decomposition (Theorem 5 in Supplementary Materials). Taken together, these theoretical and empirical results on the scalability of weighted TREE-QMC are promising.

It is worth noting that weighted ASTRAL/ASTER (with 16 threads) was faster than weighted TREE-QMC (with one thread) in our computational experiments (Fig. 3a). Weighted ASTRAL uses multi-threading to speed up the quartet score calculation, parallelizing the computation across gene trees, which contribute independently to the quartet score. Parallelism across gene trees could also be exploited by weighted TREE-QMC’s during graph construction algorithm, and future versions of the software should take advantage of multi-threading and vector instructions available on modern processors.

### Missingness and supertrees

Quartet weight normalization was an important algorithmic development from the original TREE-QMC method. Here, we showed that incomplete gene trees require their own normalization factors to be computed, which means that normalization cannot be correctly performed on quartets that have been extracted from gene trees, as required by QMC or related algorithms. We introduced a more efficient algorithm for computing the correct normalization factors for each gene tree and found in simulations that correct normalization is critical for species tree accuracy when gene trees are incomplete (Fig. 2a). Because the simulated data sets were characterized by extreme missingness and relatively low ILS, our results suggest that TREE-QMC is a promising supertree method. We found similar results for the plant data set, for which TREE-QMC successfully recovered many established clades in the NCBI taxonomy similar to Asteroid and concatenation. The ASTRAL/ASTER tree, in contrast, was farther from the NCBI taxonomy and concatenation tree, displaying a highly imbalanced (caterpillar-like) branching pattern towards the root. Although this topology was better supported (Local PP) and achieved a higher quartet score, our results on simulated data sets do not suggest that higher mean branch support and/or total quartet score are always indicators of greater species tree accuracy.

### Species tree accuracy and robustness to gene tree errors

As in Zhang and Mirarab (2022), we found that the hybrid quartet weighting scheme typically improved the accuracy of TREE-QMC compared to the unweighted version (Figs. 2e and 2f). A limitation of the simulated data sets is that error is due to insufficient phylogenetic signal or other issues arising during gene tree estimation. Such errors may follow the “MSC+Error+Support” model, under which Zhang and Mirarab (2022) prove weighted ASTRAL/ASTER is statistical consistent, unlike the unweighted method. However, these errors do not fully encompass those observed on real data sets due to incorrect orthology inference, poor quality multiple sequence alignments, and violations in the model of molecular sequence evolution. Although the effectiveness of methods on biological data is difficult to discern, it is promising that the hybrid and length weighting schemes made TREE-QMC and ASTRAL/ASTER more robust to *systematic* homology errors in the Palaeognathae data set (Cloutier et al., 2019). However, we caution that there was still insufficient signal in this data set to resolve this part of the avian evolutionary history (also see Simmons et al., 2016).

### Species tree topological stability

We also reanalyzed a recent avian data set curated by Wu et al. (2024a). A major finding of this study is the separation of *Neoaves* into two major clades: *Telluraves* (land birds) and *Aquaterraves* (waterbirds and relatives). Springer and Gatesy (2024) later identified homology errors in the data set and showed that ASTRAL-III on the cleaned data set did not recover the *Telluraves* and *Aquaterraves* split. Subsequently, Wu et al. (2024a) responded that NJst on cleaned data set still recovered the split and suggested that ASTRAL-III was less stable than NJst.

Our reanalysis also suggests that ASTRAL/ASTER is less stable than the other methods, as ASTRAL/ASTER returned different trees when outlier taxa were removed with TreeShrink, unlike ASTRID and TREE-QMC, which returned the same tree regardless of taxon filtering. When considering the five ways of running weighted summary methods (with different inputs and different weighting schemes), ASTRAL/ASTER produced four different species trees (differing by up to six branches), whereas TREE-QMC and ASTRID each produced two trees (differing by three branches). This may have interesting implications for branch support estimated by bootstrapping procedures.

The trees produced by ASTRAL/ASTER were distinct from those produced by TREE-QMC and the other methods, including concatenation. Not only did ASTRAL/ASTER not recover the Telluraves/Aquaterraves split, it also displayed a highly imbalanced (caterpillar-like) branch patterning at the deeper nodes of Aquarterraves minus pigeons. Although the ASTRAL/ASTER trees were better supported and achieved the highest quartet scores, as previously mentioned, our results on simulated data sets do not suggest that higher mean branch support and total quartet score are always indicators of greater species tree accuracy. Interestingly, the TREE-QMC tree was closer to the trees produced by concatenation as well as the distance methods NJst and ASTRID. It is possible that distance methods and TREE-QMC’s divide-and-conquer framework might behave more similarly to each other than ASTRAL/ASTER’s heuristic search, although we note that TREE-QMC was more robust to missing data than ASTRID in our simulation study (Fig. 2b). It is also worth noting that TREE-QMC does optimize the same criterion score as ASTRAL/ASTER because it downweights quartets based on subproblem decomposition, which could result in a scoring function that is more local or diffuse across the species tree, akin to likelihood scores. Future work should further study ASTRAL/ASTER and TREE-QMC to better understand the impact of objective function and search heuristics on species tree accuracy.

## Appendix A: Efficient Quartet Weight Normalization given Incomplete Gene Trees

In this appendix, we describe our approach for normalizing quartet weights for a subproblem given incomplete gene trees.

### Preliminaries

A quartet is an unrooted, binary tree on four leaves {*a, b, c, d*}; thus it takes on one of three possible topologies, denoted *a, b*|*c, d* or *a, c*|*b, d* or *a, d*|*b, c*. Let *L* denote the mapping function from leaf vertices to taxon labels for the subproblem. To avoid confusion, we use *lower case letters to indicate leaf vertices* and *upper case letters to indicate leaf labels, also referred to as taxa or species*. This distinction is necessary because the introduction of artificial taxa results in multiple leaves being labeled by the same label (Fig. 6a–c).

**Fig. 6.**
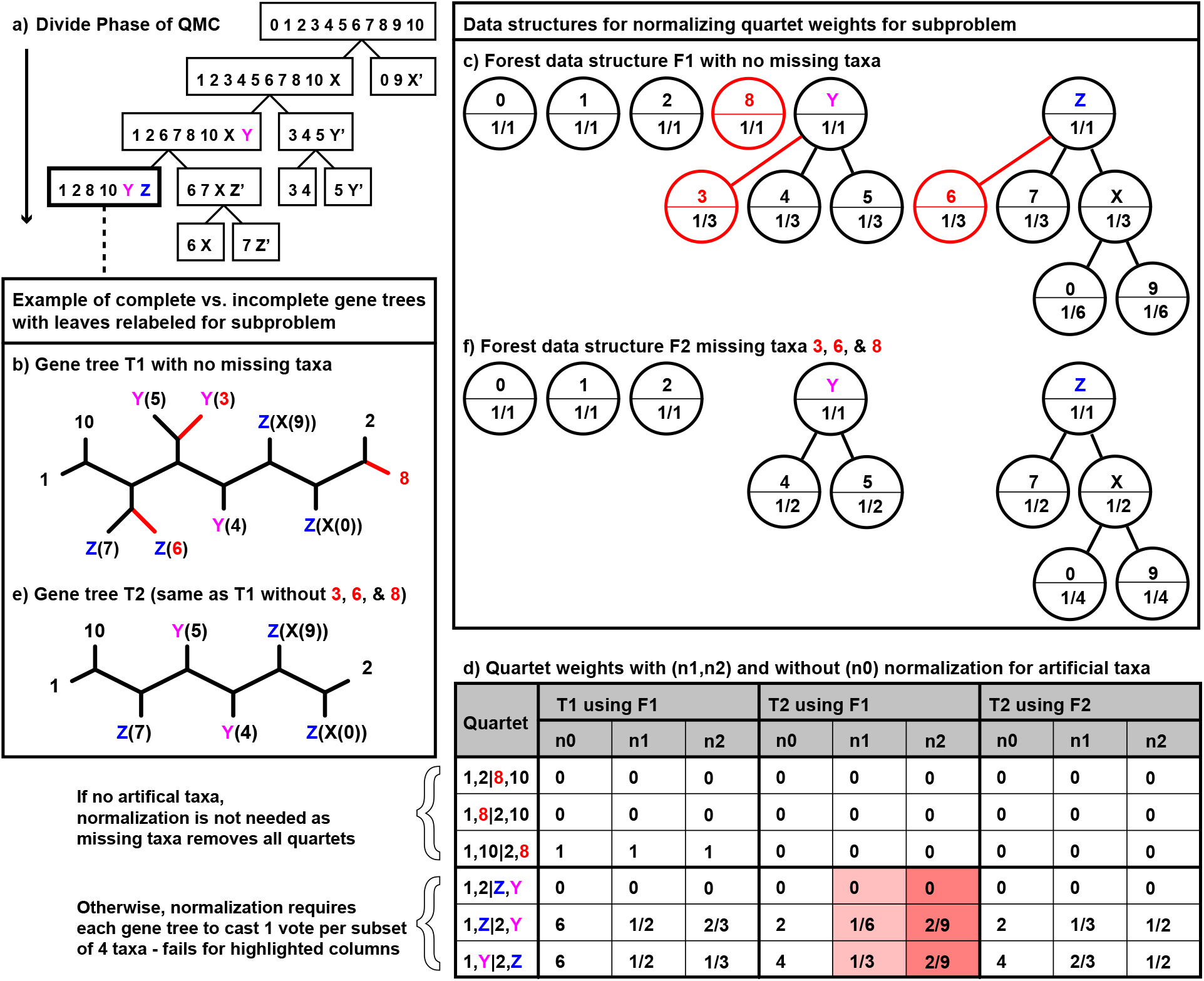
Normalization for artificial taxa. **a)** The divide-and-conquer algorithm starts with a set of input gene trees on 11 taxa. Taxa are split into two disjoint subsets at each subproblem generated during the divide phase. Take the subproblem on taxon set *{*1, 2, 8, 10, *Y, Z}* as an example, where *X, Y*, and *Z* are artificial taxa representing *{*0, 9*}, {*3, 4, 5*}*, and *{*6, 7, *X}*, respectively. **b)** A complete gene tree *T* 1 with leaves relabeled for the subproblem. **c)** Forest data structure *F* 1 for the subproblem. **d)** Quartets on *{*1, 2, *Y, Z}* implied by gene tree *T* 1 have weights summing to 1 after normalization if using forest data structure *F* 1 to compute importance values. **e)** An incomplete gene tree *T* 2 with leaves relabeled for the subproblem (note that *T* 2 is the same as *T* 1 but missing taxa 3, 6, and 8). Two quartets (1, 7|2, 5 and 1, 7|2, 4) in gene tree *T* 2 correspond to 1, *Z*|2, *Y* after relabeling for the subproblem. If using *F* 1 importance values for n2 normalization, I(1, 7, 2, 5) = I(1, 7, 2, 4) = 1 *·* 1*/*3 *·* 1 *·* 1*/*3 = 1*/*9. For n1 normalization, the importance values are I(1, 7, 2, 5) = I(1, 7, 2, 4) = 1 *·* 1*/*4 *·* 1 *·* 1*/*3 = 1*/*12. Thus, the total weight of quartet 1, *Z*|2, *Y* is 1*/*6 and 2*/*9 after n1 and n2 normalization, respectively. Repeating this process for the other two topologies 1, 2|*Z, Y* and 1, *Y* |2, *Z* does not produce weights that sum to 1 (highlighted columns). **f)** Forest data structure *F* 2 for subproblem but updated to exclude taxa 3, 6, and 8. Quartets on *{*1, 2, *Y, Z}* implied by gene tree *T* 2 have weights summing to 1 after normalization if using *F* 2.

### Normalization, Importance Values, and Forest Data Structure

A key observation in the original TREE-QMC method is that gene trees can contribute more than one quartet per subset of four species when they are multi-labeled (Han and Molloy, 2023). Thus, we wish to construct the normalized quartet graph, with quartet weights normalized so that each gene tree votes once for each subset of four labels present in the tree. To implement normalization, TREE-QMC assigns an **importance value** `I`(*x*) to each leaf vertex *x* and then **normalization factor** for each quartet *q* = *x, y*|*z, w* by multiplying the importance values of its leaves:

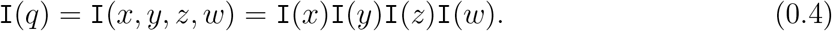

Later, we will use similar notation to indicate importance values for two or three leaves being multiplied together (e.g., I(*x, y*) = `I`(*x*)`I`(*y*) and `I`(*x, y, z*) = `I`(*x*)`I`(*y*)`I`(*z*)). It is easy to show that if

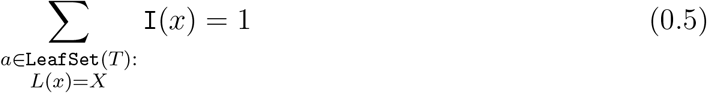

for every taxon label *X*, then the importance values form a valid normalization scheme, specifically

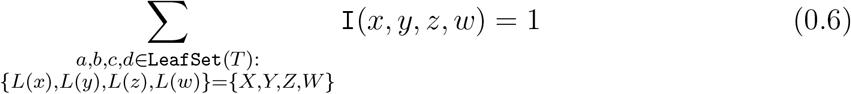

for selections of four unique taxon labels {*X, Y, Z, W*} from the subproblem. Note that in the **unnormalized** case (**n0**), the importance values are set to one for all leaves, which does not satisfy Equation 0.5 beyond the first subproblem in which all leaves are uniquely labeled.

TREE-QMC implements two normalization schemes: **uniform** (**n1**) and **non-uniform** (**n2**). The latter requires storing the hierarchical relationships between artificial taxa and real taxa for the subproblem as a collection of disjoint trees *ℱ* = {*F*_1_, *F*_2_, …, *F*_*x*_}, which we refer to as a **forest data structure**. The roots of trees in *ℱ* are taxa in the current subproblem, whose importance values sum to one following Equation 0.5. Leaf vertices of the trees in *ℱ* are singleton taxa for the subproblem, which appear at most once in any gene tree; all other vertices represent artificial taxa for the subproblem.

To construct the quartet graph for the subproblem, the leaves of each gene tree are labeled by the roots of trees in *ℱ* based on ancestor-descendant relationships (Fig. 6a–c). Importance values can be computed using *ℱ* for both normalization schemes. In uniform normalization (n1), all leaves in the same tree *F*_*i*_ ∈ *ℱ* are assigned the same importance value equal to the inverse of the number of leaves in *F*_*i*_. In non-uniform normalization (n2), the importance values are computed via the recurrence:

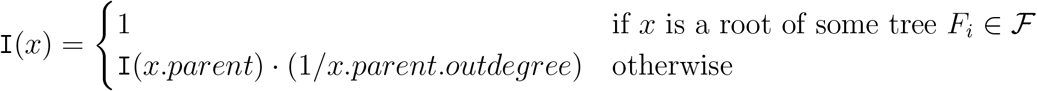

where *x.parent.outdegree* returns the number of outgoing edges for the parent of *x* in *F*_*i*_ (Fig. 6c). The recurrence can be implemented via a preorder traversal of each tree in *ℱ*.

### Normalization for Incomplete Gene Trees

If there are no missing taxa, the same importance values can be used for all gene trees (Fig. 6a–d). However, if a gene tree is incomplete, the importance values for subsets of four taxa may not sum to one, violating the requirement of normalization (Fig. 6d–e). To normalize correctly, importance values need to be computed for each gene tree based on its missing taxa (Fig. 6f). Early versions of the TREE-QMC code addressed the issue of incomplete gene trees by copying the forest data structure for each gene tree and then deleting the missing taxa while updating the importance values accordingly. This naive approach suffers due to frequent memory allocation and thus cache misses.

To improve performance, the current version of the TREE-QMC computes importance values from the forest data structure for the subproblem for each gene tree on the fly in two steps (see Algorithm 1 in the Supplementary Materials). First, the outdegree of internal nodes in the forest is updated to account for missing data via postorder traversal. Second, the importance values are updated according to the recurrence via preorder traversal. The time complexity to get the normalization values per gene tree is thus *O*(*n*), which is lower than constructing the quartet graph from the gene tree (see Appendix B). To implement normalization, we maintain the forest data structure for each subproblem on the stack during the divide phase of TREE-QMC. As the storage complexity of each forest is *O*(*n*) and the maximum depth of recursion is *O*(*n*), the total storage complexity for the forest data structure is *O*(*n*^2^), which is the same as the quartet graph on the complete species set. Thus, our normalization approach does not increase the time or storage complexity of TREE-QMC.

## Appendix B: Efficient Normalized Quartet Graph Construction leveraging Gene Tree Branch Lengths and Support Values

In this appendix, we describe how to construct the (normalized) quartet graph using the weighting schemes of Zhang and Mirarab (2022) without explicitly extracting all weighted quartets from the gene trees (Fig. S1 in Supplementary Materials). All referenced Definitions and Lemmas, Theorems, and Corollaries in this section are provided in Section 2 of the Supplementary Materials.

### Strategy for Computing Weighted Bad Edges and Good for Subproblem

Recall that vertices in the quartet graph represent taxa from the subproblem and edges between taxa are weighted by quartets implied by the gene trees, whose leaves map to taxa. If the leaves of some quartet *q* = *a, b*|*c, d* implied by a gene tree *T* have distinct labels, then *q* contributes two **bad** edges (*L*(*x*), *L*(*y*)) and (*L*(*z*), *L*(*w*)) as well as four **good** edges (*L*(*x*), *L*(*z*)), (*L*(*x*), *L*(*w*)), (*L*(*y*), *L*(*z*)), and (*L*(*y*), *L*(*w*)) to the quartet graph, provided that *L*(*x*) ≠ *L*(*y*) ≠ *L*(*z*) ≠ *L*(*w*); otherwise *q* does not contribute any good or bad edges. Good and bad edges inherit the weight of the quartet that contributed them, with the weight set to one in the unweighted algorithm and set using Equations 0.1, 0.2, or 0.3 for the length, support, or hybrid weighting schemes, respectively. Observe that under the hybrid weighting scheme, branch support weighting can be turned off by setting all support values to one and similarly length weighting can be turned off by setting all branch lengths to zero. Moreover, **non-binary gene trees** can be handled by simply refining polytomies and setting the resulting edges to have support zero and length zero (all other edges have support one and length zero in the unweighted algorithm or inherit their respective values from the unrefined tree in the weighted algorithm). As non-binary trees as well as length and support weights can be handled via the hybrid weighting scheme, we focus on hybrid weights henceforth.

For ease of computation, we split the calculation of the hybrid weight into two parts:

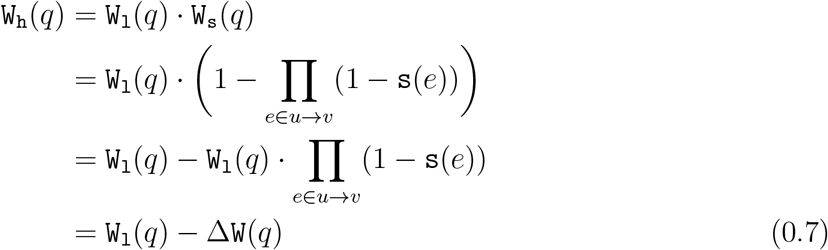

where *u* and *v* denote the anchor vertices for quartet *q* in gene tree *T* (Fig. 1). Then, the total weight of bad edges between *X* and *Y* in the quartet graph computed from gene tree *T* becomes

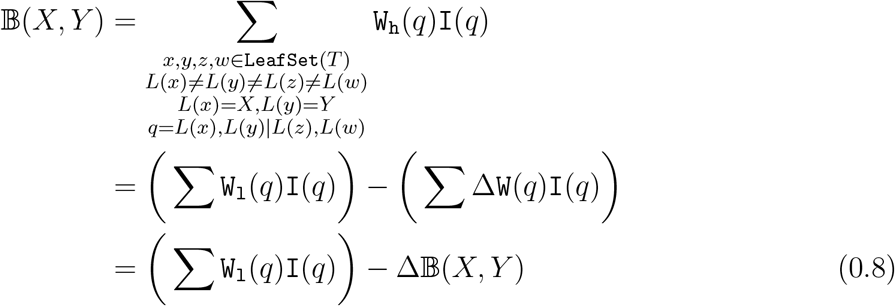

where `I`(*q*) corresponds to the importance value for normalizing the weight of each quartet *q* (Equation 0.6) and the summations loop over all ways of selecting four unique leaves *x, y, z, w* from `LeafSet`(*T*) such that the implied quartet has distinct labels and separates two leaves labeled by *X* and *Y* from the other two leaves. Observe that an algorithm for computing the left term ∆ 𝔹 (*X, Y*) given gene tree *T* can be used to compute the right term after setting all branch support values in *T* to zero. Summing the left and right terms together for all pairs of taxa (*X, Y*) and repeating across all *k* gene trees yields the bad edges 𝔹 in the weighted quartet graph for the subproblem. The good edges, denoted 𝔾, can be computed in a similar fashion.

We therefore focus on algorithms for computing ∆𝔹 (*X, Y*) and ∆𝔾 (*X, Y*) from a single unrooted gene tree *T* henceforth. We assume that *T* is binary (because any non-binary tree can be transformed into a binary tree with support values of zero on its refined edges, as previously mentioned) with *n* leaves mapping to *s* = *a* + *b* taxon labels for the subproblem, where *a* denotes the number of singleton taxa and *b* denotes the number of artificial taxa (Fig. 6). For ease of computation, we root *T* arbitrarily, letting *h* denote its height *h* (note that in our complexity analysis *h* and *n* are taken to be the maximum values across the *k* gene trees). We let *t.p, t.s, t.l*, and *t.r* denote the parent, sibling, left child, and right child of vertex *t*, respectively. We let `below`(*t*) denote the set of leaves “below” the subtree rooted at vertex *t* (i.e., they are descendants of vertex *t*). Conversely, we let `above`(*t*) denote the set of leaves “above” or “outside” the subtree rooted at *t* (i.e., they are in the set `LeafSet`(`T`) \ `below`(*t*)).

To provide some intuition for computing ∆𝔹 (*X, Y*) and ∆𝔾 (*X, Y*), consider a pair of leaves *x* and *y* in `LeafSet`(*T*) such that *L*(*x*) = *X* and *L*(*y*) = *Y*. Let *x* → *y* denote the (unique) path between *x* and *y* in *T*, and let *ℱ* denote the set of subtrees obtained from deleting edges on the path from *x* → *y* as well as their endpoints from *T*. To form a bad or good quartet for leaf pair *x, y*, we must select two leaves *z* and *w* from *ℱ* such that *L*(*z*) *≠ L*(*w*) *≠ L*(*x*) *≠ L*(*y*) (Fig. 6A in Han and Molloy, 2023). All pairs of *z* and *w* selected from the same subtree in *ℱ* correspond to a “bad quartet for *X, Y* ” because *x, y* are siblings (one anchor vertex is on the path from *x* → *y* and the other anchor vertex is within the subtree from which *z, w* were selected). Likewise, all pairs of *z* and *w* selected from different subtrees in *ℱ* correspond to a “good quartet for *X, Y*” because *x, y* are NOT siblings (both anchor vertices of the quartet are on the path *x* → *y*).

Building upon this idea, we propose to compute ∆𝔹 (*X, Y*) and ∆𝔾 (*X, Y*) by summing up the weights from bad and good quartets for *X, Y* at their anchor vertices in gene tree *T*. Specifically, we let ∆𝔹_*t*_(*X, Y*) and ∆𝔾_*t*_(*X, Y*) correspond to the weights from bad and good quartets for *X, Y* such that one leaf is below the left child of *t* (call *x*), one leaf is below the right child of *t* (call *y*), and one of *x, y* is labeled by *X* and the other is labeled by *Y* (w.l.o.g. *X* is the taxa labeling *x* and *Y* is the taxon labeling *y*; see Definitions 1 and 2). Because *T* can be multi-labeled (e.g., due to the introduction artificial taxa), we must consider all pairs *x, y* such that *x* ∈ {*L*(*l*) = *X* : *l* ∈ `below`(*t.l*)} and *y* ∈ {*L*(*l*) = *Y* : *l* ∈ `below`(*t.r*)}. To complete the bad or good quartet, we must consider all ways of selecting two additional leaves *z* and *w* from *T* such that *L*(*z*) *≠ L*(*w*) *≠ L*(*x*) *≠ L*(*y*). This allows us to break the computation of ∆𝔹_*t*_(*X, Y*) and ∆𝔾_*t*_(*X, Y*) into six cases:

a. *z, w* ∈ `below`(*t.l*),
b. *z* ∈ `below`(*t.l*) and *w* ∈ `above`(*t.p*),
c. *z, w* ∈ `above`(*t.p*),
d. *z* ∈ `above`(*t.p*) and *w* ∈ `below`(*t.r*) (symmetric with case (b)),
e. *z, w* ∈ `below`(*t.r*) (symmetric with case (a)), (f) *z* ∈ `below`(*t.l*) and *w* ∈ `below`(*t.r*).

It is easy to see that case (c) is not relevant for the computation of ∆𝔾 (*X, Y*) because *z* and *w* are drawn from the same subtree off the path from any *x, y* under consideration, thus forming a bad quartet. We define equations to compute the weights from each of these cases and then sum the relevant cases to get ∆𝔹_*t*_(*X, Y*) and ∆𝔾_*t*_(*X, Y*) (Theorems 1 and 2). Our equations rely on the precomputation of six auxiliary values, summarized below.

- 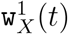 and 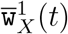: **singlet auxiliary values** give the sum of length weights from leaves labeled by taxon *X* in the subtree below and above vertex *t*, respectively (Definition 3 and 4). Singlets are useful for computing the branch lengths of terminal edges to leaves labeled *X* in the quartets. They can be computed for all taxa *X* and for all vertices *t* ∈ *V* (*T*) in *O*((*a* + *b*)*n*) time and require *O*((*a* + *b*)*n*) storage to access later (Lemmas 1 and 2).
- 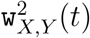 and 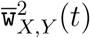: **doublet auxiliary values** give the sum of weights corresponding to pairs of leaves labeled by taxa *X, Y* in the subtree below and above vertex *t* (Definition 5 and 6). Doublets are useful for accessing the complete branch lengths of terminal edges for leaf pairs labeled *X, Y* attached to the same in anchor in a quartet and computing the branch support for the internal edge. They can be computed for all pairs of taxa *X, Y* and for all vertices *t* ∈ *V* (*T*) in *O*((*a* + *b*)^2^*n*) time and require *O*((*a* + *b*)^2^*n*) storage to access later (Lemmas 3 and 4).
- 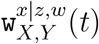 : **bad triplet auxiliary values** give the sum of weights from three leaves *x, z, w* in the subtree below *t* such that the triplet corresponds to a bad quartet for *X, Y* (Definition 7). Bad triplets are useful for accessing the branch lengths for the three terminal edges and the branch support for the internal edge in a bad quartet. They can be computed for all pairs of taxa *X, Y* and all vertices *t* ∈ *V* (*T*) in *O*((*a* + *b*)^4^*n*) time and require *O*((*a* + *b*)^2^*n*) storage to access later (Lemma 5).
- 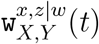 : **good triplet auxiliary values** give the sum of weights from three leaves *x, z, w* in the subtree below *t* such that the triplet corresponds to a good quartet for *X, Y* (Definition 8). Good triplets can be computed for all pairs of taxa *X, Y* and for all vertices *t* ∈ *V* (*T*) in *O*((*a* + *b*)^4^*n*) time and require *O*((*a* + *b*)^2^*n*) storage to access later (Lemma 6).

After precomputation, ∆𝔹_*t*_(*X, Y*) and ∆𝔾_*t*_(*X, Y*) can be computed in constant time. Summing across all vertices in *T* enables us to build ∆𝔹 (*X, Y*) and ∆𝔾 (*X, Y*) for all *X, Y* in *O*((*a* + *b*)^2^*n*) time and *O*((*a* + *b*)^2^) space (Theorem 3). This does not exceed the requirements for precomputation, putting the final time and space complexity for computing ∆𝔹 and ∆𝔾 given gene tree *T* at *O*((*a* + *b*)^4^*n*) and *O*(*a*^2^*n* + *abn* + *b*^2^*n*), respectively.

### Efficient Algorithm for Computing Weighted Bad Edges and Good for Subproblem

Our core contribution is reducing the final time and space complexity to *O*(*a*^2^*b* + *ab*^2^*h* + *b*^3^*n*) and *O*(*a*^2^ + *ab* + *b*^2^*n*), respectively (Theorem 4). We summarize our strategy below.

### Reducing the time complexity of triplet auxiliary values by aggregating singletons

Our first goal is to reduce the time complexity of computing triplets. To achieve this goal, we introduce a new label 1 to represent all singleton taxa and then extend the computation of singlets and doublets to this new label in the natural way (Definitions 9-11 and Corollaries 1–3), effectively aggregating the computation across all singleton taxa. We then reorganize the computation of triplet recurrences to leverage the aggregated singlets and doublets, along with some other tricks. This drops the time complexity of the triplet recurrences from *O*((*a* + *b*)^2^) to *O*(*b*) (Lemmas 7–10 and Corollaries 4–5 and 7–8), enabling us to compute triplet auxiliary values for all vertices in *T* and for all taxa *X, Y* in *O*((*a* + *b*)^2^*bn*) time (Corollaries 6 and 9) instead of *O*((*a* + *b*)^4^*n*) time.

### Reducing the storage and time complexity of auxiliary values for singleton taxa

Our second goal is to reduce the storage and time complexity of auxiliary values for singleton taxa. To achieve this goal, we break the computation of ∆𝔹 (*X, Y*) and ∆𝔾 (*X, Y*) into three cases (i)–(iii), depending on whether zero, one, or both of *X* and *Y* are singleton taxa. In case (i), there are multiple leaves labeled *X*, each with its own path to the root of *T*, and similarly for the leaves labeled *Y*. This means there are multiple vertices (lowest common ancestors of *X, Y*) in gene tree *T* where we need to compute ∆𝔹_*t*_(*X, Y*) and ∆𝔾_*t*_(*X, Y*), with the number of vertices is *O*(*n*). In case (ii), there are multiple leaves labeled *Y*, each with its own path to the root, but there is only one path from the unique leaf labeled *X* to the root. Again, there are multiple vertices in gene tree *T* where we need to compute ∆𝔹_*t*_(*X, Y*) and ∆𝔾_*t*_(*X, Y*), but the number of vertices is *O*(*h*). Lastly, in case (iii), there is one path from the unique leaf *x* labeled *X* to the root and one path from the unique leaf *y* labeled *Y* to the root. So there is just one vertex in gene tree *T* where we need to compute ∆𝔹_*t*_(*X, Y*) and ∆𝔾_*t*_(*X, Y*). The same analysis holds for the auxiliary values.

We can exploit path uniqueness for singleton taxa to reduce the storage and time complexity of auxiliary values. Take the singlet auxiliary value 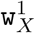 as an example. Fortunately, the singlet 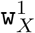 is only evaluated at *t.l* and *t.r* when computing ∆𝔹_*t*_(*X, Y*) and ∆G_*t*_(*X, Y*) (Theorems 1 and 2). If *X* is a singleton taxon, there are three possible cases at vertex *t*: (1), *X* labels a leaf below *t.l* so 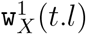 exists and 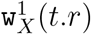 is undefined, (2) *X* labels a leaf below *t.r* so 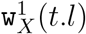 is undefined and 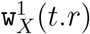 exists, and (3) *X* labels a leaf above *t* so both 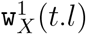and 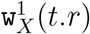 are undefined. ∆𝔹_*t*_(*X, Y*) and ∆𝔾_*t*_(*X, Y*) are also undefined in case (3), so we only need to consider cases (1) and (2). In both cases, we can compute ∆𝔹_*t*_(*X, Y*) and ∆𝔾_*t*_(*X, Y*) in a postorder traversal, so that we do not need to store 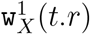 or 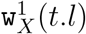 after processing vertex *t*. Before continuing the traversal from *t*, we need to compute and store 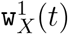 to access it at *t.p*. In case (1), we can compute 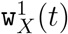 from 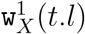, and in case (2) we can compute 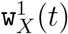 from 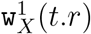. Thus, if *X* is a singleton, we can simplify the recurrence for 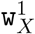 (denoted 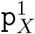), and we can overwrite the old value at *t.l* or *t.r* with the new value at *t*. This reduces the storage and time complexity of 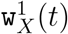 from *O*(*an* + *bn*) to *O*(*a* + *bn*) (Corollary 10). Similar results can be given for the other auxiliary quantities (Corollaries 11–14).

Ultimately, the auxiliary values with at least one singleton taxon can be computed on the fly during weighted graph construction and do not affect time or storage complexity (proof of Theorem 4). Thus, we only need to compute and store auxiliary values for artificial taxa during precomputation. This reduces the time complexity of precomputation from *O*((*a* + *b*)^4^*n*) to *O*(*b*^3^*n*) and reduces the storage complexity from *O*((*a* + *b*)^2^*n*) to *O*(*b*^2^*n*) (proof of Theorem 4).

#### Wrapping up

Lastly, the computation of ∆𝔹_*t*_(*X, Y*) and ∆𝔾 _*t*_(*X, Y*) can be improved by applying the previously introduced tricks based on aggregating singleton taxa and splitting up the computation into three cases based on whether *X* and/or *Y* are singletons (Corollaries 15 and 16). After putting everything together, it takes *O*((*a*^2^*b* + *ab*^2^*h* + *b*^3^*n*)*k*) time and *O*((*a* + *b*)^2^ + *b*^2^*n*) = *O*(*a*^2^ + *ab* + *b*^2^*n*) space to construct the weighted quartet graph for the subproblem given *k* gene trees, including precomputation (Theorem 4). The only work remaining for the subproblem is seeking a max cut. The heuristic (Dunning et al., 2018) we use to seek a max cut of the weighted quartet graph does not exceed the time complexity of graph construction (Theorem 2 in Han and Molloy, 2023).

## Acknowledgements

We thank Dr. Mark Springer and Dr. John Gatesy for sharing the estimated gene trees used in their reply to Wu et al. (2024a) and for helpful discussions about the data. We thank the editor Dr. Matthew Hahn and the anonymous reviewers for detailed feedback that improved our study.

## References

Anisimova, M., M. Gil, J.-F. Dufayard, C. Dessimoz, and O. Gascuel. 2011. Survey of Branch Support Methods Demonstrates Accuracy, Power, and Robustness of Fast Likelihood-based Approximation Schemes. Systematic Biology 60:685–699.

Avni, E., R. Cohen, and S. Snir. 2014. Weighted quartets phylogenetics. 64:233–242.

Cloutier, A., T. B. Sackton, P. Grayson, M. Clamp, A. J. Baker, and S. V. Edwards. 2019. Whole-genome analyses resolve the phylogeny of flightless birds (Palaeognathae) in the presence of an empirical anomaly zone. Systematic Biology 68:937–955.

Cunha, T. J., J. D. Reimer, and G. Giribet. 2021. Investigating Sources of Conflict in Deep Phylogenomics of Vetigastropod Snails. Systematic Biology 71:1009–1022.

Dunning, I., S. Gupta, and J. Silberholz. 2018. What works best when? a systematic evaluation of heuristics for Max-Cut and QUBO. 30:421–624.

Gatesy, J., R. W. Meredith, J. E. Janecka, M. P. Simmons, W. J. Murphy, and M. S. Springer. 2017. Resolution of a concatenation/coalescence kerfuffle: partitioned coalescence support and a robust family-level tree for mammalia. Cladistics 33:295–332.

Gatesy, J., D. B. Sloan, J. M. Warren, R. H. Baker, M. P. Simmons, and M. S. Springer. 2019. Partitioned coalescence support reveals biases in species-tree methods and detects gene trees that determine phylogenomic conflicts. Molecular Phylogenetics and Evolution 139:106539.

Han, Y. and E. K. Molloy. 2023. Improving quartet graph construction for scalable and accurate species tree estimation from gene trees. Genome Research Pages gr–277629.

Han, Y. and E. K. Molloy. 2024. Quartets enable statistically consistent estimation of cell lineage trees under an unbiased error and missingness model. Algorithms for Molecular Biology 18:19.

Hosner, P. A., B. C. Faircloth, T. C. Glenn, E. L. Braun, and R. T. Kimball. 2016. Avoiding missing data biases in phylogenomic inference: An empirical study in the landfowl (Aves: Galliformes). Mol. Biol. Evol. 33:1110–1125.

Islam, M., K. Sarker, T. Das, R. Reaz, and M. S. Bayzid. 2020. STELAR: a statistically consistent coalescent-based species tree estimation method by maximizing triplet consistency. BMC Genomics 21:136.

Jarvis, E. D., S. Mirarab, et al. 2014. Whole-genome analyses resolve early branches in the tree of life of modern birds. Science 346:1320–1331.

Lafond, M. and C. Scornavacca. 2019. On the weighted quartet consensus problem. 769:1–17.

Legried, B., E. K. Molloy, T. Warnow, and S. Roch. 2021. Polynomial-time statistical estimation of species trees under gene duplication and loss. 28:452–468.

Liu, B. and T. Warnow. 2023. Weighted astrid: fast and accurate species trees from weighted internode distances. Algorithms for Molecular Biology 18:6.

Liu, L. and L. Yu. 2011. Estimating species trees from unrooted gene trees. 60:661–667.

Liu, L., L. Yu, and S. V. Edwards. 2010. A maximum pseudo-likelihood approach for estimating species trees under the coalescent model. BMC Evolutionary Biology 10:302.

Mai, U. and S. Mirarab. 2018. Treeshrink: fast and accurate detection of outlier long branches in collections of phylogenetic trees. BMC Genomics 19:272.

Mendes, F. K. and M. W. Hahn. 2017. Why concatenation fails near the anomaly zone. Systematic Biology 67:158–169.

Minh, B. Q., H. A. Schmidt, O. Chernomor, D. Schrempf, M. D. Woodhams, A. von Haeseler, and R. Lanfear. 2020. IQ-TREE 2: New Models and Efficient Methods for Phylogenetic Inference in the Genomic Era. Molecular Biology and Evolution 37:1530–1534.

Mirarab, S., R. Reaz, M. S. Bayzid, T. Zimmermann, M. S. Swenson, and T. Warnow. 2014. ASTRAL: genome-scale coalescent-based species tree estimation. Bioinformatics 30:i541–i548.

Mirarab, S., I. Rivas-González, S. Feng, J. Stiller, Q. Fang, U. Mai, G. Hickey, G. Chen, N. Brajuka, O. Fedrigo, G. Formenti, J. B. W. Wolf, K. Howe, A. Antunes, M. H. Schierup, B. Paten, E. D. Jarvis, G. Zhang, and E. L. Braun. 2024. A region of suppressed recombination misleads neoavian phylogenomics. Proceedings of the National Academy of Sciences 121:e2319506121.

Mirarab, S. and T. Warnow. 2015. ASTRAL-II: coalescent-based species tree estimation with many hundreds of taxa and thousands of genes. Bioinformatics 31:i44–i52.

Molloy, E. K., J. Gatesy, and M. S. Springer. 2022. Theoretical and practical considerations when using retroelement insertions to estimate species trees in the anomaly zone. Systematic Biology 71:721–740.

Molloy, E. K. and T. Warnow. 2018. To include or not to include: The impact of gene filtering on species tree estimation methods. 67:285–303.

Morel, B., A. M. Kozlov, and A. Stamatakis. 2018. ParGenes: a tool for massively parallel model selection and phylogenetic tree inference on thousands of genes. Bioinformatics 35:1771–1773.

Morel, B., T. A. Williams, and A. Stamatakis. 2023. Asteroid: a new algorithm to infer species trees from gene trees under high proportions of missing data. Bioinformatics 29.

Nute, M., J. Chou, E. K. Molloy, and T. Warnow. 2018. The performance of coalescent-based species tree estimation methods under models of missing data. 19(Suppl 5):286.

One Thousand Plant Transcriptomes Initiative. 2019. One thousand plant transcriptomes and the phylogenomics of green plants. Nature 574:679–685.

Pratt, J. W. 1959. Remarks on zeros and ties in the wilcoxon signed rank procedures. Journal of the American Statistical Association 54:655––667.

Price, M. N., P. S. Dehal, and A. P. Arkin. 2010. FastTree 2 – approximately maximum-likelihood trees for large alignments. PLOS ONE 5:1–10.

Rabiee, M., E. Sayyari, and S. Mirarab. 2019. Multi-allele species reconstruction using ASTRAL. 130:286–296.

Rannala, B. and Z. Yang. 2003. Bayes estimation of species divergence times and ancestral population sizes using DNA sequences from multiple loci. Genetics 164:1645–1656.

Robinson, D. and L. Foulds. 1981. Comparison of phylogenetic trees. Mathematical Biosciences 53:131–147.

Roch, S. and M. Steel. 2015. Likelihood-based tree reconstruction on a concatenation of aligned sequence data sets can be statistically inconsistent. 100:56–62.

Sayyari, E. and S. Mirarab. 2016. Fast coalescent-based computation of local branch support from quartet frequencies. 33:1654–1668.

Sayyari, E., J. B. Whitfield, and S. Mirarab. 2017. Fragmentary Gene Sequences Negatively Impact Gene Tree and Species Tree Reconstruction. Molecular Biology and Evolution 34:3279–3291.

Simmons, M. P. and J. Gatesy. 2021. Collapsing dubiously resolved gene-tree branches in phylogenomic coalescent analyses. Molecular Phylogenetics and Evolution 158:107092.

Simmons, M. P., D. B. Sloan, and J. Gatesy. 2016. The effects of subsampling gene trees on coalescent methods applied to ancient divergences. Molecular Phylogenetics and Evolution 97:76–89.

Simmons, M. P., M. S. Springer, and J. Gatesy. 2022. Gene-tree misrooting drives conflicts in phylogenomic coalescent analyses of palaeognath birds. Molecular Phylogenetics and Evolution 167:107344.

Snir, S. and S. Rao. 2010. Quartets MaxCut: A divide and conquer quartets algorithm. 7:704–718.

Snir, S. and S. Rao. 2012. Quartet MaxCut: A fast algorithm for amalgamating quartet trees. 62:1–8.

Springer, M. S. and J. Gatesy. 2016. The gene tree delusion. 94:1–33.

Springer, M. S. and J. Gatesy. 2024. A new phylogeny for aves is compromised by pervasive misalignment and homology problems. Proceedings of the National Academy of Sciences USA 121:e2406494121.

Springer, M. S., E. K. Molloy, D. B. Sloan, M. P. Simmons, and J. Gatesy. 2020. ILS-Aware analysis of low-homoplasy retroelement insertions: Inference of species trees and introgression Using quartets. Journal of Heredity 111:147–168.

Stamatakis, A. 2014. RAxML version 8: a tool for phylogenetic analysis and post-analysis of large phylogenies. Bioinformatics 30:1312–1313.

Steenwyk, J., G. Martínez-Redondo, T. Buida, E. Gluck-Thaler, X.-X. Shen, T. Gabaldón, A. Rokas, and R. Fernández. 2024. Phykit: A multitool for phylogenomics. Preprints.

Steenwyk, J. L., Y. Li, X. Zhou, X.-X. Shen, and A. Rokas. 2023. Incongruence in the phylogenomics era. Nature Reviews Genetics 24:834–850.

Stiller, J., S. Feng, A.-A. Chowdhury, I. Rivas-González, D. A. Duchêne, Q. Fang, Y. Deng, A. Kozlov, A. Stamatakis, S. Claramunt, J. M. T. Nguyen, S. Y. W. Ho, B. C. Faircloth, J. Haag, P. Houde, J. Cracraft, M. Balaban, U. Mai, G. Chen, R. Gao, C. Zhou, Y. Xie, Z. Huang, Z. Cao, Z. Yan, H. A. Ogilvie, L. Nakhleh, B. Lindow, B. Morel, J. Fjeldså, P. A. Hosner, R. R. da Fonseca, B. Petersen, J. A. Tobias, T. Székely, J. D. Kennedy, A. H. Reeve, A. Liker, M. Stervander, A. Antunes, D. T. Tietze, M. F. Bertelsen, F. Lei, C. Rahbek, G. R. Graves, M. H. Schierup, T. Warnow, E. L. Braun, M. T. P. Gilbert, E. D. Jarvis, S. Mirarab, and G. Zhang. 2024. Complexity of avian evolution revealed by family-level genomes. Nature 629:851–860.

Tavaré, S. 1986. Some probabilistic and statistical problems in the analysis of DNA sequences. 17:57–86.

Vachaspati, P. and T. Warnow. 2015. ASTRID: Accurate species trees from internode distances. BMC Genomics 16:S3.

Waterhouse, A. M., J. B. Procter, D. M. A. Martin, M. Clamp, and G. J. Barton. 2009. Jalview version 2–a multiple sequence alignment editor and analysis workbench. Bioinformatics 25:1189–1191.

Wilcoxon, F. 1949. Some Rapid Approximate Statistical Procedures Page 6. American Cyanamid Company, New York.

Wu, S., F. E. Rheindt, J. Zhang, J. Wang, L. Zhang, C. Quan, Z. Li, M. Wang, F. Wu, Y. Qu, S. V. Edwards, Z. Zhou, and L. Liu. 2024a. Genomes, fossils, and the concurrent rise of modern birds and flowering plants in the late cretaceous. Proceedings of the National Academy of Sciences 121:e2319696121.

Wu, S., F. E. Rheindt, J. Zhang, J. Wang, L. Zhang, C. Quan, Z. Li, M. Wang, F. Wu, Y. Qu, S. V. Edwards, Z. Zhou, and L. Liu. 2024b. Reply to springer and gatesy: The impact of long branches and misalignments on phylogenetic analysis is minimal. Proceedings of the National Academy of Sciences 121:e2409344121.

Xi, Z., L. Liu, and C. C. Davis. 2015. Genes with minimal phylogenetic information are problematic for coalescent analyses when gene tree estimation is biased. 92:63–71.

Yang, Z. 1993. Maximum-likelihood estimation of phylogeny from DNA sequences when substitution rates differ over sites. Molecular Biology and Evolution 10:1396–1401.

Yoo, D., A. Rhie, P. Hebbar, F. Antonacci, G. A. Logsdon, S. J. Solar, D. Antipov, B. D. Pickett, Y. Safonova, F. Montinaro, Y. Luo, J. Malukiewicz, J. M. Storer, J. Lin, A. N. Sequeira, R. J. Mangan, G. Hickey, G. M. Anez, P. Balachandran, A. Bankevich, C. R. Beck, A. Biddanda, M. Borchers, G. G. Bouffard, E. Brannan, S. Y. Brooks, L. Carbone, L. Carrel, A. P. Chan, J. Crawford, M. Diekhans, E. Engelbrecht, C. Feschotte, G. Formenti, G. H. Garcia, L. d. Gennaro, D. Gilbert, R. E. Green, A. Guarracino, I. Gupta, D. Haddad, J. Han, R. S. Harris, G. A. Hartley, W. T. Harvey, M. Hiller, K. Hoekzema, M. L. Houck, H. Jeong, K. Kamali, M. Kellis, B. Kille, C. Lee, Y. Lee, W. Lees, A. P. Lewis, Q. Li, M. Loftus, Y. H. E. Loh, H. Loucks, J. Ma, Y. Mao, J. F. I. Martinez, P. Masterson, R. C. McCoy, B. McGrath, S. McKinney, B. S. Meyer, K. H. Miga, S. K. Mohanty, K. M. Munson, K. Pal, M. Pennell, P. A. Pevzner, D. Porubsky, T. Potapova, F. R. Ringeling, J. L. Rocha, O. A. Ryder, S. Sacco, S. Saha, T. Sasaki, M. C. Schatz, N. J. Schork, C. Shanks, L. Smeds, D. R. Son, C. Steiner, A. P. Sweeten, M. G. Tassia, F. Thibaud-Nissen, E. Torres-González, M. Trivedi, W. Wei, J. Wertz, M. Yang, P. Zhang, S. Zhang, Y. Zhang, Z. Zhang, S. A. Zhao, Y. Zhu, E. D. Jarvis, J. L. Gerton, I. Rivas-González, B. Paten, Z. A. Szpiech, C. D. Huber, T. L. Lenz, M. K. Konkel, S. V. Yi, S. Canzar, C. T. Watson, P. H. Sudmant, E. Molloy, E. Garrison, C. B. Lowe, M. Ventura, R. J. O’Neill, S. Koren, K. D. Makova, A. M. Phillippy, and E. E. Eichler. 2024. Complete sequencing of ape genomes. bioRxiv.

Yule, G. U. 1925. A mathematical theory of evolution, based on the conclusions of Dr. J. C. Willis, F.R.S. Philosophical Transactions of the Royal Society of London. Series B, Containing Papers of a Biological Character 213:21–87.

Zhang, C. and S. Mirarab. 2022. Weighting by gene tree uncertainty improves accuracy of quartet-based species trees. 39:msac215.

Zhang, C., M. Rabiee, E. Sayyari, and S. Mirarab. 2018. ASTRAL-III: Polynomial time species tree reconstruction from partially resolved gene trees. 19:153.

