## Supplementary Materials for "Improved robustness to gene tree incompleteness, estimation errors, and systematic homology errors with weighted TREE-QMC"

September 27, 2024

#### Contents

**List of Tables** **2**

**List of Figures** **2**

**1 Supplemental Methods I: Quartet Weight Normalization** **3**

**2 Supplemental Methods II: Weighted Quartet Graph Construction** **3**

**References** **63**

S20 Testing for TREE-QMC-wh_n2 vs wASTRID on ASTRAL-II data with varying levels of ILS 58
S21 Testing for TREE-QMC-wh_n2 vs CAML on ASTRAL-II data with varying numbers of taxa 58
S22 Testing for TREE-QMC-wh_n2 vs ASTER-h on ASTRAL-II data with varying numbers of taxa 59
S23 Testing for TREE-QMC-wh_n2 vs wASTRID on ASTRAL-II data with varying numbers of taxa . . . 59

### 1 Supplemental Methods I: Quartet Weight Normalization

In this section, we provide the algorithm for computing taxon importance values for a given gene tree on the fly, picking up from Appendix A in the main text.

**Algorithm 1:** Computing Importance Values for Normalization on the Fly

**input :** Forest data structure $F$ that stores importance values for a subproblem and gene tree $g$, which may be complete or incomplete
**output:** Forest data structure with updated importance values based on the taxa in gene tree $g$

$Q \leftarrow$ all leaves $l$ in $F$ such that $l$ is also in $g$;
**while** $Q \neq \varnothing$ **do**
  $v \leftarrow Q$.pop()
  $F[v]$.importance $\leftarrow 1$
  **if** $v$.parent *in* $F$ *is not visited* **then**
    $Q$.push($F[v]$.parent)
    $F[v]$.parent.outdegree $\leftarrow 0$
  $F[v]$.parent.outdegree $\leftarrow F[v]$.parent.outdegree $+ 1$
$Q \leftarrow$ all roots $r$ in $F$ with $r$.importance $= 1$
**while** $Q \neq \varnothing$ **do**
  $v \leftarrow Q$.pop()
  **for** *each child* $u$ *of* $v$ **do**
    $Q$.push($u$)
    $F[u]$.importance $\leftarrow F[u]$.importance $\cdot F[v]$.outdegree$^{-1}$
**return** $F$

### 2 Supplemental Methods II: Weighted Quartet Graph Construction

In this section, we show how to construct the quartet graph (Fig. S1), picking up from section "Strategy for Computing Weighted Bad Edges and Good for Subproblem" in Appendix B of the main text. Recall the definitions for bad and good edges below vertex $t$ in gene tree $T$.

**Definition 1** (bad edges below $t$)**.** *The total weight of bad edges between taxa $X, Y$ below vertex $t$ in gene tree $T$ is defined as*

$$\Delta\mathbb{B}_t(X,Y) = \sum_{\substack{x,y,z,w \in \texttt{LeafSet}(T) \\ L(x)\neq L(y)\neq L(z)\neq L(w) \\ x\in \texttt{below}(t.l),\, y\in\texttt{below}(t.r) \\ L(x)=X,\, L(y)=Y \\ q=x,y|z,w}} \mathtt{I}(q)\cdot\Delta\mathtt{W}(q) \quad (1)$$

*where $x, y, z, w \in$ $\texttt{LeafSet}(T)$ enumerates all* unordered *tuples of four leaves from $T$ such that $L(x) \neq L(y) \neq L(z) \neq L(w)$, one leaf is below the left child of $t$ (call $x$), one leaf below the right child of $t$ (call $y$), one of $x, y$ is labeled by $X$ and the other is labeled by $Y$ (w.l.o.g. $X$ is the taxa labeling $x$ and $Y$ is the taxon labeling $y$), and $X$ and $Y$ are siblings in the resulting quartet.*

**Definition 2** (good edges below $t$)**.** *The total weight of good edges between taxa $X, Y$ below vertex $t$ in gene* *tree $T$ is defined as*

$$\Delta\mathbb{G}_t(X,Y) = \sum_{\substack{x,y,z,w\in\texttt{LeafSet}(T)\\ L(x)\neq L(y)\neq L(z)\neq L(w)\\ x\in\texttt{below}(t.l),y\in\texttt{below}(t.r)\\ L(x)=X,L(y)=Y\\ q=x,z|y,w}} \texttt{I}(q)\cdot\Delta\texttt{W}(q) \tag{2}$$

*where $x,y,z,w \in \texttt{LeafSet}(T)$ enumerates all* unordered *tuples of four unique leaves from $T$ such that $L(x)\neq L(y)\neq L(z)\neq L(w)$, one leaf is below the left child of $t$ (call $x$), one leaf below the right child of $t$ (call $y$), one of $x,y$ is labeled by $X$ and the other is labeled by $Y$ (w.l.o.g. $X$ is the taxa labeling $x$ and $Y$ is the taxon labeling $y$), and $X$ and $Y$ are NOT siblings in the resulting quartet.*

The taxa in the subproblem are broken down into two subsets: the set $\mathcal{S}$ of singleton taxa and the set $\mathcal{A}$ of artificial taxa. For time and space complexity analyses, $a=|\mathcal{S}|$, $b=|\mathcal{A}|$, $n$ is the number of leaves in gene tree $T$, and $h$ is the height of $T$.

#### 2.1 Auxiliary values w

Before giving equations to compute $\Delta\mathbb{B}_t(X,Y)$ and $\Delta\mathbb{G}_t(X,Y)$ from $T$, we define six auxiliary values, described below.

**Definition 3** (singlet auxiliary values below $t$)**.** *Let $\texttt{w}^1_X(t)$ be the sum of weights from leaves labeled by $X$ in the subtree* below *vertex $t$ in gene tree $T$. More precisely, the singlet auxiliary values of $X$* below *$t$ are defined as*

$$\texttt{w}^1_X(t) = \sum_{\substack{x\in\texttt{below}(t)\\ L(x)=X}} \texttt{I}(x)\cdot\texttt{exp}(\sum_{e\in x\to t} -\texttt{l}(e)) \tag{3}$$

*where $\texttt{s}(e)$ and $\texttt{l}(e)$ denote the support value and branch length of an edge $e$ in $T$.*

**Lemma 1** (singlet auxiliary values below $t$)**.** *$\texttt{w}^1_X(t)$ can be computed according to the following recurrence:*

$$\texttt{w}^1_X(t) = \begin{cases} \texttt{I}(t) & \textit{if } t \textit{ is a leaf in } T\\ \texttt{exp}(-\texttt{l}(t.l))\cdot\texttt{w}^1_X(t.l) + \texttt{exp}(-\texttt{l}(t.r))\cdot\texttt{w}^1_X(t.r) & \textit{otherwise} \end{cases} \tag{4}$$

*It takes $O((a+b)n)$ time to compute $\texttt{w}^1_X(t)$ for all vertices $t\in V(T)$ and for all taxa $X\in\mathcal{S}\cup\mathcal{A}$. Likewise, it takes $O((a+b)n)$ space to access these singlet auxiliary values later. The time and storage complexity drops to $O(bn)$ when limiting the computation to artificial taxa (i.e., $X\in\mathcal{A}$).*

*Proof.* In the base case, vertex $t$ is a leaf, so $\texttt{w}^1_X(t) = I(x)$ because there are no edges on the path from any leaf $x$ to itself so we only need to consider the importance value at the leaf. Otherwise $t$ is a non-leaf vertex, so we can split the weight into two parts: weight from leaves in the subtree below $t.l$ and weight from leaves in the subtree below $t.r$:

$$\begin{aligned}
\texttt{w}^1_X(t) &= \sum_{\substack{x\in\texttt{below}(t)\\ L(x)=X}} \texttt{I}(x)\cdot\texttt{exp}(\sum_{e\in x\to t} -\texttt{l}(e))\\
&= \sum_{\substack{x\in\texttt{below}(t.l)\\ L(x)=X}} \texttt{I}(x)\cdot\texttt{exp}(\sum_{e\in x\to t} -\texttt{l}(e)) + \sum_{\substack{x\in\texttt{below}(t.r)\\ L(x)=X}} \texttt{I}(x)\cdot\texttt{exp}(\sum_{e\in x\to t} -\texttt{l}(e))\\
&= \texttt{exp}(-\texttt{l}(t.l))\cdot \sum_{\substack{x\in\texttt{below}(t.l)\\ L(x)=X}} \texttt{I}(x)\cdot\texttt{exp}(\sum_{e\in x\to t.l} -\texttt{l}(e)) + \texttt{exp}(-\texttt{l}(t.r))\cdot \sum_{\substack{x\in\texttt{below}(t.r)\\ L(x)=X}} \texttt{I}(x)\cdot\texttt{exp}(\sum_{e\in x\to t.r} -\texttt{l}(e))\\
&= \texttt{exp}(-\texttt{l}(t.l))\cdot\texttt{w}^1_X(t.l) + \texttt{exp}(-\texttt{l}(t.r))\cdot\texttt{w}^1_X(t.r).
\end{aligned}$$

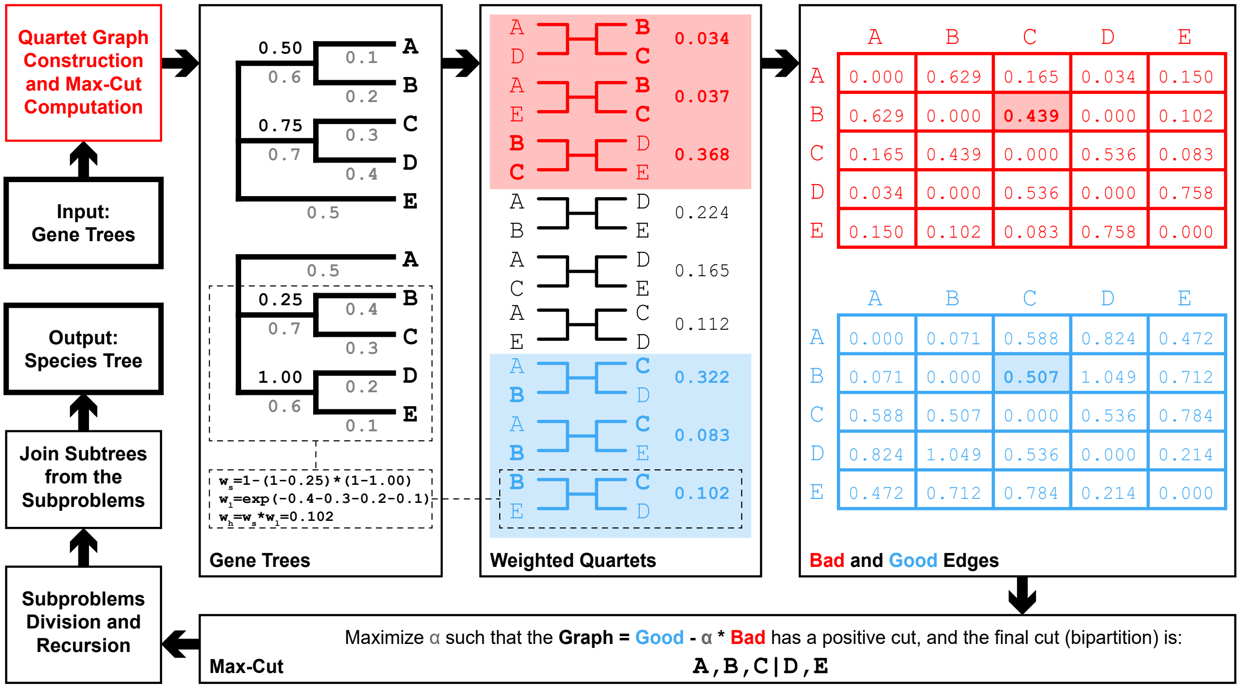

Figure S1: **Weighted Quartet Graph Construction.** The goal of weighted TREE-QMC is to take into account gene tree branch lengths and support values when constructing the quartet graph, whose (max) cut determines a bipartition of taxa in the output unrooted species tree. Consider the two input gene trees on 5 taxa with branches are labeled by branch lengths (below) and branch support values (above). The naive approach to construct quartet graph first extracts all weighted quartets from the gene trees. For example, the quartet $q = B, C|D, E$ only exists in one of the gene trees. Its two internal branches have support values of 0.25 and 1.00 and its four terminal branches have lengths of 0.4, 0.3, 0.2, and 0.1, respectively. As a result, its length weight is $\mathtt{W_l}(q) = \mathtt{exp}(-0.4-0.3-0.2-0.1) = 0.102$, its support weight is $\mathtt{W_s}(q) = 1-(1-0.25)\cdot(1-1) = 1$, and its hybrid weight is $\mathtt{W_h}(q) = \mathtt{W_s}(q)\cdot\mathtt{W_l}(q) = 0.102$, following Equations 1, 2, and 3 in the main text, respectively. To compute the good and bad edge hybrid weights between the taxa $B$ and $C$, we consider all quartets containing them. There are three quartets in which $B$ and $C$ are siblings—the total hybrid weight of the bad edges between $B$ and $C$, denoted $\mathbb{B}(B, C)$, is $0.034+0.039+0.368 = 0.439$. Similarly, we consider all quartets involving $B$ and $C$ in which $B$ and $C$ are not siblings—the total weight of the good edges between $B$ and $C$, denoted $\mathbb{G}(B, C)$, is $0.322 + 0.083 + 0.102 = 0.507$. Finally, we compute the optimal cut of the quartet graph and obtain the bipartition $A, B, C|D, E$, which is possible by brute in this example because the number of taxa is small. The TREE-QMC method continues by recursion on the subproblems defined by the two halves of the bipartition: $A, B, C$ and $D, E$, after the introduction of artificial taxa $\mathcal{A}_1$ and $\mathcal{A}_2$ to represent taxa on the other side of the bipartition $\{D, E\}$ and $\{A, B, C\}$, respectively. To solve the first subproblem, we would construct quartet graph relabeling the gene trees with taxa $\{A, B, C, \mathcal{A}_1\}$ and seek its max cut. The termination criterion is already satisfied for the second subproblem on $\{D, E, \mathcal{A}_2\}$ because only unrooted phylogenetic tree is possible on three leaves.

This recurrence can be computed by traversing the $O(n)$ vertices of gene tree $T$ in a postorder fashion. Evaluating the recurrence for space for vertex $t$ and taxon $X$ takes $O(1)$ time and storing the resulting auxiliary value requires $O(1)$. Repeating for all $O(a + b)$ taxa $X \in \mathcal{S} \cup \mathcal{A}$ gives the time and storage complexity. □

**Definition 4** (singlet auxiliary values above $t$)**.** *Let* $\overline{\mathtt{w}}^1_X(t)$ *be the sum of weights from leaves labeled by* $X$ *in the subtree* above *vertex* $t$ *in gene tree* $T$*. More precisely, the singlet auxiliary values of* $X$ above $t$ *are* *defined as*

$$\overline{\mathtt{w}}^1_X(t) = \sum_{\substack{x\in\mathtt{above}(t)\\ L(x)=X}} \mathtt{I}(x)\cdot \mathtt{exp}(\sum_{e\in x\to t} -\mathtt{l}(e)), \tag{5}$$

*where* $\mathtt{s}(e)$ *and* $\mathtt{l}(e)$ *denote the support value and branch length of an edge* $e$ *in* $T$*.*

**Lemma 2** (singlet auxiliary values above $t$)**.** $\overline{\mathtt{w}}^1_X(t)$ *can be computed according to the following recurrence*

$$\overline{\mathtt{w}}^1_X(t) = \begin{cases} 0 & \text{if } t \text{ is the root of } T \\ \mathtt{exp}(-\mathtt{l}(t))\cdot \overline{\mathtt{w}}^1_X(t.p) + \mathtt{exp}(-\mathtt{l}(t.s))\cdot \mathtt{w}^1_X(t.s) & \text{otherwise} \end{cases} \tag{6}$$

*After computing* $\mathtt{w}^1_X$ *(Lemma 1), the time and space complexity for* $\overline{\mathtt{w}}^1_X$ *is the same as for* $\mathtt{w}^1_X$ *(Lemma 1).*

*Proof.* In the base case, vertex $t$ is a root, so $\mathtt{w}^1_X(t) = 0$ because there are no edges or importance values associated with the root. Otherwise $t$ is a non-root vertex, so we can split the weight into two parts: weight from the leaves in the subtree above $t.p$ and weight from the leaves in the subtree below $t.s$:

$$\begin{aligned}
\overline{\mathtt{w}}^1_X(t) &= \sum_{\substack{x\in\mathtt{above}(t)\\ L(x)=X}} \mathtt{I}(x)\cdot \mathtt{exp}(\sum_{e\in x\to t} -\mathtt{l}(e)) \\
&= \sum_{\substack{x\in\mathtt{above}(t.p)\\ L(x)=X}} \mathtt{I}(x)\cdot \mathtt{exp}(\sum_{e\in x\to t} -\mathtt{l}(e)) + \sum_{\substack{x\in\mathtt{below}(t.s)\\ L(x)=X}} \mathtt{I}(x)\cdot \mathtt{exp}(\sum_{e\in x\to t} -\mathtt{l}(e)) \\
&= \mathtt{exp}(-\mathtt{l}(t))\cdot \sum_{\substack{x\in\mathtt{above}(t.p)\\ L(x)=X}} \mathtt{I}(x)\cdot \mathtt{exp}(\sum_{e\in x\to t.p} -\mathtt{l}(e)) + \mathtt{exp}(-\mathtt{l}(t.s))\cdot \sum_{\substack{x\in\mathtt{below}(t.s)\\ L(x)=X}} \mathtt{I}(x)\cdot \mathtt{exp}(\sum_{e\in x\to t.s} -\mathtt{l}(e)) \\
&= \mathtt{exp}(-\mathtt{l}(t))\cdot \overline{\mathtt{w}}^1_X(t.p) + \mathtt{exp}(-\mathtt{l}(t.s))\cdot \mathtt{w}^1_X(t.s).
\end{aligned}$$

After precomputation $\overline{\mathtt{w}}^1_X(t)$ can be computed by traversing the $O(n)$ vertices of gene tree $T$ in a preorder fashion. The remainder of the complexity analysis for $\overline{\mathtt{w}}^1_X(t)$ is the same as for $\mathtt{w}^1_X(t)$ (Lemma 1). □

**Definition 5** (doublet auxiliary values below $t$)**.** *Let* $\mathtt{w}^2_{X,Y}(t)$ *be the sum of the weights from leaf pairs* $x$ *and* $y$ *in the subtree* below *vertex* $t$ *in gene tree* $T$ *such that one leaf (w.l.o.g.* $x$*) is labeled* $X$*, the other leaf (w.l.o.g.* $y$*) is labeled* $Y$*, and* $X \neq Y$*. More precisely, the doublet auxiliary values of* $X$ *and* $Y$ below $t$ *are defined as*

$$\mathtt{w}^2_{X,Y}(t) = \mathtt{w}^2_{Y,X}(t) = \sum_{\substack{x,y\in\mathtt{below}(t)\\ L(x)=X, L(y)=Y}} \mathtt{I}(x,y)\cdot \mathtt{exp}(\sum_{e\in x,y\to u} -\mathtt{l}(e))\cdot \prod_{e\in u\to t}(1-\mathtt{s}(e)) \tag{7}$$

$$\tag{8}$$

*where* $u$ *is the lowest common ancestor of* $x, y$*, denoted* $lca(x, y)$*.*

**Lemma 3** (doublet auxiliary values below $t$)**.** $\mathtt{w}^2_{X,Y}(t)$ *can be computed according to the following recurrence:*

$$\mathtt{w}^2_{X,Y}(t) = \mathtt{w}^2_{Y,X}(t) = \begin{cases} 0 & \text{if } t \text{ is a leaf in } T \\ (1-\mathtt{s}(t.l))\cdot \mathtt{w}^2_{X,Y}(t.l) + (1-\mathtt{s}(t.r))\cdot \mathtt{w}^2_{X,Y}(t.r) & \\ +\mathtt{exp}(-\mathtt{l}(t.l))\cdot \mathtt{w}^1_X(t.l)\cdot \mathtt{exp}(-\mathtt{l}(t.r))\cdot \mathtt{w}^1_Y(t.r) & \\ +\mathtt{exp}(-\mathtt{l}(t.l))\cdot \mathtt{w}^1_Y(t.l)\cdot \mathtt{exp}(-\mathtt{l}(t.r))\cdot \mathtt{w}^1_X(t.r) & \text{otherwise} \end{cases} \tag{9}$$

*After computing* $\mathtt{w}^1_X$ *and* $\mathtt{w}^1_Y$ *(Lemma 1), it takes* $O((a+b)^2n)$ *time to compute* $\mathtt{w}^2_{X,Y}(t)$ *for all vertices* $t \in V(T)$ *and for all pairs of taxa* $X, Y \in \mathcal{S}\cup\mathcal{A}$*. Likewise, it takes* $O((a+b)^2n)$ *space to access these doublet auxiliary values later. The time and storage complexity drops to* $O(b^2n)$ *when only pairs of artificial taxa are considered (i.e.,* $X, Y \in \mathcal{A}$*).*

*Proof.* In the base case, we have $\mathtt{w}^2_{X,Y}(t) = 0$ because it is impossible to select the two leaves when $t$ is a leaf. Otherwise $t$ is a non-leaf vertex, so we split the sum into four cases:

$$\begin{aligned}
\mathtt{w}^2_{X,Y}(t) &= \sum_{\substack{x,y \in \mathtt{below}(t) \\ L(x)=X, L(y)=Y}} \mathtt{I}(x,y) \cdot \Delta\mathtt{W}(x,y|t) \\
&= \sum_{\substack{x,y \in \mathtt{below}(t.l) \\ L(x)=X, L(y)=Y}} \mathtt{I}(x,y) \cdot \Delta\mathtt{W}(x,y|t) + \sum_{\substack{x,y \in \mathtt{below}(t.r) \\ L(x)=X, L(y)=Y}} \mathtt{I}(x,y) \cdot \Delta\mathtt{W}(x,y|t) \\
&+ \sum_{\substack{x \in \mathtt{below}(t.l) \\ y \in \mathtt{below}(t.r) \\ L(x)=X, L(y)=Y}} \mathtt{I}(x,y) \cdot \Delta\mathtt{W}(x,y|t) + \sum_{\substack{x \in \mathtt{below}(t.r) \\ y \in \mathtt{below}(t.l) \\ L(x)=X, L(y)=Y}} \mathtt{I}(x,y) \cdot \Delta\mathtt{W}(x,y|t).
\end{aligned}$$

where

$$\Delta\mathtt{W}(x,y|t) = \sum_{\substack{x,y \in \mathtt{below}(t) \\ L(x)=X, L(y)=Y}} \mathtt{I}(x,y) \cdot \mathtt{exp}(\sum_{e \in x,y \to u} -\mathtt{l}(e)) \cdot \prod_{e \in u \to t} (1 - \mathtt{s}(e))$$

with $u = lca(x,y)$.

In the first and second terms, both $x$ and $y$ are from the same subtree so we recurse on the subtree. Assume $x, y \in v.l$ (the other case is similar) and we have

$$\begin{aligned}
&\sum_{\substack{x,y \in \mathtt{below}(t.l) \\ L(x)=X, L(y)=Y}} \mathtt{I}(x,y) \cdot \Delta\mathtt{W}(x,y|t) \\
=& \sum_{\substack{x,y \in \mathtt{below}(t.l) \\ L(x)=X, L(y)=Y}} \mathtt{I}(x,y) \cdot \mathtt{exp}(\sum_{e \in x,y \to u} -\mathtt{l}(e)) \cdot \prod_{e \in u \to t} (1 - \mathtt{s}(e)) \\
=& (1 - \mathtt{s}(t.l)) \cdot \left( \sum_{\substack{x,y \in \mathtt{below}(t.l) \\ L(x)=X, L(y)=Y}} \mathtt{I}(x,y) \cdot \mathtt{exp}(\sum_{e \in x,y \to u} -\mathtt{l}(e)) \cdot \prod_{e \in u \to t.l} (1 - \mathtt{s}(e)) \right) \\
=& (1 - \mathtt{s}(t.l)) \cdot \sum_{\substack{x,y \in \mathtt{below}(t.l) \\ L(x)=X, L(y)=Y}} \mathtt{I}(x,y) \cdot \Delta\mathtt{W}(x,y|t.l) \\
=& (1 - \mathtt{s}(t.l)) \cdot \mathtt{w}^2_{X,Y}(t.l).
\end{aligned}$$

In the third and fourth terms, $x$ and $y$ are from different subtrees so their lowest common ancestor is $t$. Assume $x \in \mathtt{below}(t.l)$ and $y \in \mathtt{below}(t.r)$. We may independently choose any $x$ from $below(t.l)$ and any $y$ from $below(t.r)$ to form a doublet. As a result, the total weight of doublets is a product of the total weights of singlets in the subtrees:

$$
\begin{aligned}
&\sum_{\substack{x\in\mathtt{below}(t.l)\\ y\in\mathtt{below}(t.r)\\ L(x)=X, L(y)=Y}} \mathtt{I}(x,y)\cdot\Delta\mathtt{W}(x,y|t)\\
=&\sum_{\substack{x\in\mathtt{below}(t.l)\\ y\in\mathtt{below}(t.r)\\ L(x)=X, L(y)=Y}} \mathtt{I}(x,y)\cdot\mathtt{exp}(\sum_{e\in x,y\to t}-\mathtt{l}(e))\\
=&\Big(\sum_{\substack{x\in\mathtt{below}(t.l)\\ L(x)=X}}\mathtt{I}(x)\cdot\mathtt{exp}(\sum_{e\in x\to t}-\mathtt{l}(e))\Big)\cdot\Big(\sum_{\substack{y\in\mathtt{below}(t.r)\\ L(y)=Y}}\mathtt{I}(y)\cdot\mathtt{exp}(\sum_{e\in y\to t}-\mathtt{l}(e))\Big)\\
=&\mathtt{exp}(-\mathtt{l}(t.l))\cdot\Big(\sum_{\substack{x\in\mathtt{below}(t.l)\\ L(x)=X}}\mathtt{I}(x)\cdot\mathtt{exp}(\sum_{e\in x\to t.l}-\mathtt{l}(e))\Big)\cdot\mathtt{exp}(-\mathtt{l}(t.l))\cdot\Big(\sum_{\substack{y\in\mathtt{below}(t.r)\\ L(y)=Y}}\mathtt{I}(y)\cdot\mathtt{exp}(\sum_{e\in y\to t.r}-\mathtt{l}(e))\Big)\\
=&\mathtt{exp}(-\mathtt{l}(t.l))\cdot\mathtt{w}^1_X(t.l)\cdot\mathtt{exp}(-\mathtt{l}(t.r))\cdot\mathtt{w}^1_Y(t.r)
\end{aligned}
$$

Adding the weights from four terms gives the final recurrence:

$$
\begin{aligned}
\mathtt{w}^2_{X,Y}(t) =&(1-\mathtt{s}(t.l))\cdot\mathtt{w}^2_{X,Y}(t.l)+(1-\mathtt{s}(t.r))\cdot\mathtt{w}^2_{X,Y}(t.r)\\
&+\mathtt{exp}(-\mathtt{l}(t.l))\cdot\mathtt{w}^1_X(t.l)\cdot\mathtt{exp}(-\mathtt{l}(t.r))\cdot\mathtt{w}^1_Y(t.r)\\
&+\mathtt{exp}(-\mathtt{l}(t.l))\cdot\mathtt{w}^1_Y(t.l)\cdot\mathtt{exp}(-\mathtt{l}(t.r))\cdot\mathtt{w}^1_X(t.r).
\end{aligned}
$$

After precomputation, $\mathtt{w}^2_{X,Y}(t)$ can be computed by traversing the $O(n)$ vertices of gene tree $T$ in a postorder fashion. Evaluating the recurrence for vertex $t$ and pair of taxa $X, Y$ takes $O(1)$ time and storing the resulting auxiliary value takes $O(1)$ space. Repeating for all $O((a+b)^2)$ pairs of taxa $X, Y \in \mathcal{S}\cup\mathcal{A}$ gives the time and storage complexity. □

**Definition 6** (doublet auxiliary values above $t$)**.** *Let $\overline{\mathtt{w}}^2_{X,Y}(t)$ be the sum of the weights from leaf pairs $x$ and $y$ in the subtree* above *vertex $t$ in gene tree $T$ such that one leaf (w.l.o.g. $x$) is labeled $X$, the other leaf (w.l.o.g. $y$) is labeled $Y$, and $X \neq Y$. More precisely, the doublet auxiliary values of $X$ and $Y$* above *$t$ are defined as*

$$
\overline{\mathtt{w}}^2_{X,Y}(t)=\overline{\mathtt{w}}^2_{Y,X}(t)=\sum_{\substack{x,y\in\mathtt{above}(t)\\ L(x)=X,L(y)=Y}}\mathtt{I}(x,y)\cdot\mathtt{exp}(\sum_{e\in u\to x,y}-\mathtt{l}(e))\cdot\prod_{e\in t\to u}(1-\mathtt{s}(e)) \tag{10}
$$

*where $u$ is either $lca(x,y)$, $lca(x,t)$ or $lca(y,t)$ depending on which of these three vertices is farthest from the root of $T$.*

**Lemma 4** (double auxiliary values above $t$)**.** *$\overline{\mathtt{w}}^2_{X,Y}(t)$ can be computed according to the following recurrence:*

$$
\overline{\mathtt{w}}^2_{X,Y}(t)=\overline{\mathtt{w}}^2_{Y,X}(t)=\begin{cases}0 & \text{if } t \text{ is the root of } T\\ (1-\mathtt{s}(t))\cdot\overline{\mathtt{w}}^2_{X,Y}(t.p)+(1-\mathtt{s}(t))\cdot(1-\mathtt{s}(t.s))\cdot\mathtt{w}^2_{X,Y}(t.s) & \\ +(1-\mathtt{s}(t))\cdot\mathtt{exp}(-\mathtt{l}(t.s))\cdot\overline{\mathtt{w}}^1_X(t.p)\cdot\mathtt{w}^1_Y(t.s) & \\ +(1-\mathtt{s}(t))\cdot\mathtt{exp}(-\mathtt{l}(t.s))\cdot\mathtt{w}^1_X(t.s)\cdot\overline{\mathtt{w}}^1_Y(t.p) & \text{otherwise}\end{cases} \tag{11}
$$

*After computing $\mathtt{w}^1_X$ and $\mathtt{w}^1_Y$ (Lemma 1), $\overline{\mathtt{w}}^1_X$ and $\overline{\mathtt{w}}^1_Y$ (Lemma 2), and $\mathtt{w}^2_{X,Y}$ (Lemma 3), the time and space complexity for $\overline{\mathtt{w}}^2_{X,Y}$ is the same as for $\mathtt{w}^2_{X,Y}$ (Lemma 3).*

*Proof.* In the base case, we have $\overline{\mathtt{w}}^2_{X,Y}(t)=0$ because the subtree above the root $t$ is empty. Otherwise $t$ is a non-root vertex, so we divide $\overline{\mathtt{w}}^2_{X,Y}(t)$ into four parts, which is similar to the proof for $\mathtt{w}^2_{X,Y}(t)$:

$$\begin{aligned}\overline{\mathtt{w}}^2_{X,Y}(t) &= \sum_{\substack{x,y\in\mathtt{above}(t)\\L(x)=X,L(y)=Y}} \mathtt{I}(x,y)\cdot\Delta\overline{\mathtt{W}}(x,y|t)\\ &= \sum_{\substack{x,y\in\mathtt{above}(t.p)\\L(x)=X,L(y)=Y}} \mathtt{I}(x,y)\cdot\Delta\overline{\mathtt{W}}(x,y|t) + \sum_{\substack{x,y\in\mathtt{below}(t.s)\\L(x)=X,L(y)=Y}} \mathtt{I}(x,y)\cdot\Delta\overline{\mathtt{W}}(x,y|t)\\ &+ \sum_{\substack{x\in\mathtt{above}(t.p)\\y\in\mathtt{below}(t.s)\\L(x)=X,L(y)=Y}} \mathtt{I}(x,y)\cdot\Delta\overline{\mathtt{W}}(x,y|t) + \sum_{\substack{x\in\mathtt{below}(t.s)\\y\in\mathtt{above}(t.p)\\L(x)=X,L(y)=Y}} \mathtt{I}(x,y)\cdot\Delta\overline{\mathtt{W}}(x,y|t)\end{aligned}$$

where

$$\Delta\overline{\mathtt{W}}(x,y|t) = \mathtt{I}(x,y)\cdot\mathtt{exp}(\sum_{e\in u\to x,y} -\mathtt{l}(e))\cdot\prod_{e\in t\to u}(1-\mathtt{s}(e))$$

and $u$ is $lca(x,y)$, $lca(x,t)$ or $lca(y,t)$ depending on which is farthest from the root of $T$. It is easy to see that $u$ is an ancestor of $t.p.p$ for the first term, $u$ is a descendant of $t.s$ for the second term, and $u$ is $t.p$ for third and fourth terms. Expanding the four terms in a similar fashion to Lemma 3 and adding the weights together gives the final recurrence:

$$\begin{aligned}\overline{\mathtt{w}}^2_{X,Y}(t) =&(1-\mathtt{s}(t))\cdot\overline{\mathtt{w}}^2_{X,Y}(t.p)\\ &+(1-\mathtt{s}(t))\cdot(1-\mathtt{s}(t.s))\cdot\mathtt{w}^2_{X,Y}(t.s)\\ &+(1-\mathtt{s}(t))\cdot\mathtt{exp}(-\mathtt{l}(t.s))\cdot\overline{\mathtt{w}}^1_X(t.p)\cdot\mathtt{w}^1_Y(t.s)\\ &+(1-\mathtt{s}(t))\cdot\mathtt{exp}(-\mathtt{l}(t.s))\cdot\mathtt{w}^1_X(t.s)\cdot\overline{\mathtt{w}}^1_Y(t.p)\end{aligned}$$

After precomputation, $\overline{\mathtt{w}}^2_{X,Y}(t)$ can be computed by traversing the $O(n)$ vertices of gene tree $T$ in a preorder fashion. The remainder of the complexity analysis for $\overline{\mathtt{w}}^2_{X,Y}(t)$ is the same as for $\mathtt{w}^2_{X,Y}(t)$ (Lemma 3). □

**Definition 7** (bad triplet auxiliary values)**.** *Let $\mathtt{w}^{x|z,w}_{X,Y}(t)$ be the sum of the weights from three leaves $x$, $z$, and $w$ in the subtree below $t$, where the topology of the triplet of $(x,z,w)$ is $x|z,w$, the outgroup of the triplet $(x)$ is labeled by $X$, and ingroups ($z$ and $w$) satisfy $L(z)\neq L(w)\neq X\neq Y$. Thus the triplet corresponds to a bad quartet for $X,Y$. The bad triplet auxiliary value below $t$ is defined as*

$$\mathtt{w}^{x|z,w}_{X,Y}(t) = \sum_{\substack{x,z,w\in\mathtt{below}(t)\\L(x)=X,L(z)\neq L(w)\neq X\neq Y\\t(x,z,w)=x|z,w}} \mathtt{I}(x,z,w)\cdot\mathtt{exp}(\sum_{\substack{e\in x,t\to u\\z,w\to v}} -\mathtt{l}(e))\cdot\prod_{e\in u\to v}(1-\mathtt{s}(e)) \tag{12}$$

*where $u$ and $v$ are anchor nodes of the quartet of $t,x|z,w$ and we assume $u$ is closer to $x$ than $v$. Unlike the case of doublets, $\mathtt{w}^{x|z,w}_{X,Y}(t)\neq\mathtt{w}^{x|z,w}_{Y,X}(t)$ because the former means that the outgroup $x$ is labeled $X$ and the latter means the outgroup $x$ is labeled $Y$ (in either case, the ingroup $z,w$ must satisfy $L(z)\neq L(w)\neq X\neq Y$). Lastly, observe that $\mathtt{w}^{x|z,w}_{X,Y}(t)$ is equal for all $Y\in\mathcal{S}$; we use $\mathtt{w}^{x|z,w}_{X,0}(t)$ to denote the bad triplet auxiliary value for any $Y\in\mathcal{S}$.*

**Lemma 5** (bad triplets auxiliary values)**.** *$\mathtt{w}^{x|z,w}_{X,Y}(t)$ can be computed according to the following recurrence:*

$$\mathtt{w}^{x|z,w}_{X,Y}(t) = \begin{cases} 0 & \text{if } t \text{ is a leaf in } T\\ \mathtt{exp}(-\mathtt{l}(t.l))\cdot\mathtt{w}^{x|z,w}_{X,Y}(t.l)+\mathtt{exp}(-\mathtt{l}(t.r))\cdot\mathtt{w}^{x|z,w}_{X,Y}(t.r) & \\ +\mathtt{exp}(-\mathtt{l}(t.l))\cdot\mathtt{w}^1_X(t.l)\cdot(1-\mathtt{s}(t.r))\cdot\sum_{\substack{Z,W\in\mathcal{S}\cup\mathcal{A}\\Z\neq W\neq X\neq Y}}\mathtt{w}^2_{Z,W}(t.r) & \\ +\mathtt{exp}(-\mathtt{l}(t.r))\cdot\mathtt{w}^1_X(t.r)\cdot(1-\mathtt{s}(t.l))\cdot\sum_{\substack{Z,W\in\mathcal{S}\cup\mathcal{A}\\Z\neq W\neq X\neq Y}}\mathtt{w}^2_{Z,W}(t.l) & \text{otherwise}\end{cases} \tag{13}$$

*where the summation enumerates all* unordered *pairs of taxa* $Z, W \in \mathcal{S} \cup \mathcal{A}$ *such that* $X \neq Y \neq Z \neq W$ *to avoid double counting. After computing* $\mathtt{w}^1_X$ *(Lemma 1) and* $\mathtt{w}^2_{Z,W}$ *for all pairs of taxa* $Z, W \in \mathcal{S} \cup \mathcal{A}$ *(Lemma 3), it takes* $O((a+b)^4 bn)$ *time to compute* $\mathtt{w}^{x|z,w}_{X,Y}(t)$ *for all vertices* $t \in V(T)$ *and for all unordered pairs of taxa* $X, Y \in \mathcal{S} \cup \mathcal{A}$*. Likewise, it takes* $O((a+b)^2 n)$ *space to access the bad triplet auxiliary values later. The time and storage complexity drops to* $O((a+b)^2 b^2 n)$ *and* $O(b^2 n)$*, respectively, when restricting* $X$ *to be an artificial taxa (i.e.,* $X \in \mathcal{A}$ *and* $Y \in \mathcal{A} \cup \{0\}$*).*

*Proof.* In the base case, $\mathtt{w}^{x|z,w}_{X,Y}(t) = 0$ because $t$ is a leaf. Otherwise, we must consider the positions of $x$, $z$, and $w$ in the subtree below $t$. Since the triplet is part of a quartet contributing a bad edge between $X$ and $Y$, $z$ and $w$ must be in the same subtree after deleting edges on the path $x \to t$ and their endpoints from $T$. Thus, it is impossible that $z \in \mathtt{below}(t.l)$ and $w \in \mathtt{below}(t.r)$ (or vise versa) and we only have four cases to consider:

(a) $x, z, w \in \mathtt{below}(t.l)$

(b) $x \in \mathtt{below}(t.l)$ and $z, w \in \mathtt{below}(t.r)$

(c) $x \in \mathtt{below}(t.r)$ and $z, w \in \mathtt{below}(t.l)$ (symmetric with case (b)

(d) $x, z, w \in \mathtt{below}(t.r)$ (symmetric with case (a))

We split $\mathtt{w}^{x|z,w}_{X,Y}(t)$ into four terms according to the cases above:

$$\begin{aligned}
\mathtt{w}^{x|z,w}_{X,Y}(t) &= \sum_{\substack{x,z,w \in \mathtt{below}(t) \\ L(z) \neq L(w) \neq X,Y \\ t(x,z,w) = x|z,w}} \mathtt{I}(x,z,w) \cdot \Delta\mathtt{W}(x,t|z,w) \\
&= \sum_{\substack{x,z,w \in \mathtt{below}(t.l) \\ L(z) \neq L(w) \neq X,Y \\ t(x,z,w) = x|z,w}} \mathtt{I}(x,z,w) \cdot \Delta\mathtt{W}(x,t|z,w) + \sum_{\substack{x,z,w \in \mathtt{below}(t.r) \\ L(z) \neq L(w) \neq X,Y \\ t(x,z,w) = x|z,w}} \mathtt{I}(x,z,w) \cdot \Delta\mathtt{W}(x,t|z,w) \\
&+ \sum_{\substack{x \in \mathtt{below}(t.l), \\ z,w \in \mathtt{below}(t.r) \\ L(z) \neq L(w) \neq X,Y \\ t(x,z,w) = x|z,w}} \mathtt{I}(x,z,w) \cdot \Delta\mathtt{W}(x,t|z,w) + \sum_{\substack{z,w \in \mathtt{below}(t.l), \\ x \in \mathtt{below}(t.r) \\ L(z) \neq L(w) \neq X,Y \\ t(x,z,w) = x|z,w}} \mathtt{I}(x,z,w) \cdot \Delta\mathtt{W}(x,t|z,w).
\end{aligned}$$

We only prove the recurrence of the cases (a) and (b) as the other two cases are similar due to symmetry. In the **case (a)**, we have

$$\begin{aligned}
&\sum_{\substack{x,z,w \in \mathtt{below}(t.l) \\ L(z) \neq L(w) \neq X,Y \\ t(x,z,w) = x|z,w}} \mathtt{I}(x,z,w) \cdot \Delta\mathtt{W}(x,t|z,w) \\
=&\sum_{\substack{x,z,w \in \mathtt{below}(t.l) \\ L(z) \neq L(w) \neq X,Y \\ t(x,z,w) = x|z,w}} \mathtt{I}(x,z,w) \cdot \mathtt{exp}(\sum_{\substack{e \in x,t \to u \\ z,w \to v}} -\mathtt{l}(e)) \cdot \prod_{e \in u \to v} (1 - \mathtt{s}(e)) \\
=&\mathtt{exp}(-\mathtt{l}(t.l)) \cdot \sum_{\substack{x,z,w \in \mathtt{below}(t.l) \\ L(z) \neq L(w) \neq X,Y \\ t(x,z,w) = x|z,w}} \mathtt{I}(x,z,w) \cdot \mathtt{exp}(\sum_{\substack{e \in x,t.l \to u \\ z,w \to v}} -\mathtt{l}(e)) \cdot \prod_{e \in u \to v} (1 - \mathtt{s}(e)) \\
=&\mathtt{exp}(-\mathtt{l}(t.l)) \cdot \mathtt{w}^{x|z,w}_{X,Y}(t.l).
\end{aligned}$$

In the **case (b)**, the topology must be $x|z,w$ so we have

$$\begin{aligned}
&\sum_{\substack{x\in\texttt{below}(t.l),\\ z,w\in\texttt{below}(t.r)\\ L(z)\neq L(w)\neq X,Y\\ t(x,z,w)=x|z,w}} \texttt{I}(x,z,w)\cdot\Delta\texttt{W}(x,t|z,w)\\
=&\sum_{\substack{x\in\texttt{below}(t.l),\\ z,w\in\texttt{below}(t.r)\\ L(z)\neq L(w)\neq X,Y}} \texttt{I}(x,z,w)\cdot\texttt{exp}(\sum_{\substack{e\in x\to t\\ z,w\to v}}-\texttt{l}(e))\cdot\prod_{e\in v\to t}(1-\texttt{s}(e))\\
=&\sum_{\substack{x\in\texttt{below}(t.l),\\ z,w\in\texttt{below}(t.r)\\ L(z)\neq L(w)\neq X,Y}} \texttt{I}(x)\cdot\texttt{exp}(-\sum_{e\in x\to t}\texttt{l}(e))\cdot\texttt{I}(z,w)\cdot\texttt{exp}(\sum_{e\in z,w\to v}-\texttt{l}(e))\cdot\prod_{e\in v\to t}(1-\texttt{s}(e))\\
=&\Big(\sum_{x\in\texttt{below}(t.l)}\texttt{I}(x)\cdot\texttt{exp}(-\sum_{e\in x\to t}\texttt{l}(e))\Big)\cdot\Big(\sum_{\substack{z,w\in\texttt{below}(t.r)\\ L(z)\neq L(w)\neq X,Y}}\texttt{I}(z,w)\cdot\texttt{exp}(\sum_{e\in z,w\to v}-\texttt{l}(e))\cdot\prod_{e\in v\to t}(1-\texttt{s}(e))\Big)\\
=&\texttt{exp}(-\texttt{l}(t.l))\cdot\texttt{w}^1_X(t.l)\cdot(1-\texttt{s}(t.r))\cdot\sum_{\substack{Z,W\in\mathcal{S}\cup\mathcal{A}\\ Z\neq W\neq X\neq Y}}\sum_{L(z)=Z,L(w)=W}\texttt{I}(z,w)\cdot\texttt{exp}(\sum_{e\in z,w\to v}-\texttt{l}(e))\cdot\prod_{e\in v\to t}(1-\texttt{s}(e))\\
=&\texttt{exp}(-\texttt{l}(t.l))\cdot\texttt{w}^1_X(t.l)\cdot(1-\texttt{s}(t.r))\cdot\sum_{\substack{Z,W\in\mathcal{S}\cup\mathcal{A}\\ Z\neq W\neq X\neq Y}}\texttt{w}^2_{Z,W}(t.r)
\end{aligned}$$

where the summation enumerates all unordered pairs of taxa $Z,W$ from the subproblem such that $X\neq Y\neq Z\neq W$ to avoid double counting. Adding the weights of the four cases together gives the final recurrence:

$$\begin{aligned}
\texttt{w}^{x|z,w}_{X,Y}(t) =&\texttt{exp}(-\texttt{l}(t.l))\cdot\texttt{w}^{x|z,w}_{X,Y}(t.l)+\texttt{exp}(-\texttt{l}(t.r))\cdot\texttt{w}^{x|z,w}_{X,Y}(t.r)\\
&+\texttt{exp}(-\texttt{l}(t.l))\cdot\texttt{w}^1_X(t.l)\cdot(1-\texttt{s}(t.r))\cdot\frac{1}{2}\cdot\sum_{\substack{Z,W\in\mathcal{S}\cup\mathcal{A}\\ Z\neq W\neq X\neq Y}}\texttt{w}^2_{Z,W}(t.r)\\
&+\texttt{exp}(-\texttt{l}(t.r))\cdot\texttt{w}^1_X(t.r)\cdot(1-\texttt{s}(t.l))\cdot\frac{1}{2}\cdot\sum_{\substack{Z,W\in\mathcal{S}\cup\mathcal{A}\\ Z\neq W\neq X\neq Y}}\texttt{w}^2_{Z,W}(t.l).
\end{aligned}$$

After precomputation, $\texttt{w}^{x|z,w}_{X,Y}(t)$ can be computed by traversing the $O(n)$ vertices of in a postorder traversal. Evaluating the recurrence for vertex $t$ and pair of taxa $X,Y$ takes $O((a+b)^2)$ time and storing the resulting auxiliary value requires $O(1)$ space. Repeating for all $O((a+b)^2)$ unordered pairs of taxa $X,Y\in\mathcal{S}\cup\mathcal{A}$ gives the time and storage complexity. □

**Definition 8** (good triplet auxiliary values)**.** *Let* $\texttt{w}^{x,z|w}_{X,Y}(t)$ *be the sum of the weights from three leaves* $x$, $z$, *and* $w$ *in the subtree below* $t$, *where the topology of the triplet of* $(x,z,w)$ *has one of the ingroups (w.l.o.g.* $x$*) labeled* $X$, *and the other two taxa satisfy* $L(z)\neq L(w)\neq X\neq Y$. *Thus, the triplet corresponds to a good quartet when selecting a leaf labeled* $Y$ *above* $t$. *The good triplet auxiliary value below* $t$ *is defined as*

$$\texttt{w}^{x,z|w}_{X,Y}(t)=\sum_{\substack{x,z,w\in\texttt{below}(t)\\ L(x)=X,L(z)\neq L(w)\neq X\neq Y\\ t(x,z,w)=x,z|w}}\texttt{I}(x,z,w)\cdot\texttt{exp}(\sum_{\substack{e\in x,z\to u\\ t,w\to v}}-\texttt{l}(e))\cdot\prod_{e\in u\to v}(1-\texttt{s}(e)) \tag{14}$$

*where* $u$ *and* $v$ *are anchor nodes of the quartet of* $x,z|w,t$ *and we assume* $u$ *is the closer to* $x$ *than* $v$. *Similar to bad triplet auxiliary values,* $\texttt{w}^{x,z|w}_{X,Y}(t)\neq\texttt{w}^{x,z|w}_{Y,X}(t)$ *and we use* $\texttt{w}^{x,z|w}_{X,0}(t)$ *to denote the quantity for any* $Y\in\mathcal{S}$.

**Lemma 6** (good triplet auxiliary values). $\mathtt{w}_{X,Y}^{x,z|w}(t)$ *can be computed according to the following recurrence:*

$$\mathtt{w}_{X,Y}^{x,z|w}(t) = \begin{cases} 0 & \text{if } t \text{ is a leaf in } T \\ \mathtt{exp}(-\mathtt{l}(t.l)) \cdot \mathtt{w}_{X,Y}^{x,z|w}(t.l) + \mathtt{exp}(-\mathtt{l}(t.r)) \cdot \mathtt{w}_{X,Y}^{x,z|w}(t.r) & \\ +(1-\mathtt{s}(t.l)) \cdot \mathtt{exp}(-\mathtt{l}(t.r)) \cdot \sum_{\substack{Z\in\mathcal{S}\cup\mathcal{A} \\ Z\neq X,Y}} \sum_{\substack{W\in\mathcal{S}\cup\mathcal{A} \\ W\neq X,Y,Z}} \mathtt{w}_{X,Z}^{2}(t.l) \cdot \mathtt{w}_{W}^{1}(t.r) & \\ +(1-\mathtt{s}(t.r)) \cdot \mathtt{exp}(-\mathtt{l}(t.l)) \cdot \sum_{\substack{Z\in\mathcal{S}\cup\mathcal{A} \\ Z\neq X,Y}} \sum_{\substack{W\in\mathcal{S}\cup\mathcal{A} \\ W\neq X,Y,Z}} \mathtt{w}_{X,Z}^{2}(t.r) \cdot \mathtt{w}_{W}^{1}(t.l) & \text{otherwise} \end{cases} \tag{15}$$

*where pairs of taxa $Z, W$ are ordered, as indicated by the nested summations, unlike when computing bad triplet auxiliary values. After computing $\mathtt{w}_{W}^{1}$ for all taxa $W \in \mathcal{S} \cup \mathcal{A}$ (Lemma 1) and $\mathtt{w}_{X,Z}^{2}$ for all taxa $Z \in \mathcal{S} \cup \mathcal{A}$ (Lemma 3), the time and space complexity for $\mathtt{w}_{X,Y}^{x,z|w}$ is the same as for $\mathtt{w}_{X,Y}^{x|z,w}$ (Lemma 5).*

*Proof.* In the base case, $\mathtt{w}_{X,Y}^{x,z|w}(t) = 0$ because $t$ is a leaf. Otherwise, we must consider the positions of $x$, $z$, and $w$ in the subtree below $t$. Since the triplet is part of a quartet contributing a good edge between $X$ and $Y$, $z$ and $w$ cannot be in the same subtree after deleting edges on the path $x \to t$ and their endpoints from $T$. Thus, we only have four cases to consider:

(a) $x, z, w \in \mathtt{below}(t.l)$

(b) $x, z \in \mathtt{below}(t.l)$ and $w \in \mathtt{below}(t.r)$

(c) $w \in \mathtt{below}(t.l)$ and $x, z \in \mathtt{below}(t.r)$ (symmetric with case (b))

(d) $x, z, w \in \mathtt{below}(t.r)$ (symmetric with case (a))

We split $\mathtt{w}_{X,Y}^{x,z|w}(t)$ into four parts according to the cases above:

$$\begin{aligned} \mathtt{w}_{X,Y}^{x,z|w}(t) &= \sum_{\substack{x,z,w\in\mathtt{below}(t) \\ L(z)\neq L(w)\neq X,Y \\ t(x,z,w)=x,z|w}} \mathtt{I}(x,z,w) \cdot \Delta\mathtt{W}(x,z|w,t) \\ &= \sum_{\substack{x,z,w\in\mathtt{below}(t.l) \\ L(z)\neq L(w)\neq X,Y \\ t(x,z,w)=x,z|w}} \mathtt{I}(x,z,w) \cdot \Delta\mathtt{W}(x,z|w,t) + \sum_{\substack{x,z,w\in\mathtt{below}(t.r) \\ L(z)\neq L(w)\neq X,Y \\ t(x,z,w)=x,z|w}} \mathtt{I}(x,z,w) \cdot \Delta\mathtt{W}(x,z|w,t) \\ &+ \sum_{\substack{x,z\in\mathtt{below}(t.l) \\ w\in\mathtt{below}(t.r) \\ L(z)\neq L(w)\neq X,Y \\ t(x,z,w)=x,z|w}} \mathtt{I}(x,z,w) \cdot \Delta\mathtt{W}(x,z|w,t) + \sum_{\substack{w\in\mathtt{below}(t.l) \\ x,z\in\mathtt{below}(t.r) \\ L(z)\neq L(w)\neq X,Y \\ t(x,z,w)=x,z|w}} \mathtt{I}(x,z,w) \cdot \Delta\mathtt{W}(x,z|w,t), \end{aligned}$$

We only prove the cases (a) and (b) since the other two cases are similar due to symmetry. In the **case (a)**, all three leaves comes from the left subtree, so we recurse on $t.l$ and append the edge from $t$ to $t.l$ to all triplets in $t.l$:

$$\begin{aligned}
&\sum_{\substack{x,z,w\in\texttt{below}(t.l)\\ L(z)\neq L(w)\neq X,Y\\ t(x,z,w)=x,z|w}} \texttt{I}(x,z,w)\cdot\Delta\texttt{W}(x,z|w,t)\\
=&\sum_{\substack{x,z,w\in\texttt{below}(t.l)\\ L(z)\neq L(w)\neq X,Y\\ t(x,z,w)=x,z|w}} \texttt{I}(x,z,w)\cdot\texttt{exp}(\sum_{\substack{e\in x,z\to u\\ t,w\to v}}-\texttt{l}(e))\cdot\prod_{e\in u\to v}(1-\texttt{s}(e))\\
=&\texttt{exp}(-\texttt{l}(t.l))\cdot\sum_{\substack{x,z,w\in\texttt{below}(t.l)\\ L(z)\neq L(w)\neq X,Y\\ t(x,z,w)=x,z|w}} \texttt{I}(x,z,w)\cdot\texttt{exp}(\sum_{\substack{e\in x,z\to u\\ t.l,w\to v}}-\texttt{l}(e))\cdot\prod_{e\in u\to v}(1-\texttt{s}(e))\\
=&\texttt{exp}(-\texttt{l}(t.l))\cdot\sum_{\substack{x,z,w\in\texttt{below}(t.l)\\ L(z)\neq L(w)\neq X,Y\\ t(x,z,w)=x,z|w}} \texttt{I}(x,z,w)\cdot\Delta\texttt{W}(x,z|w,t.l)\\
=&\texttt{exp}(-\texttt{l}(t.l))\cdot\texttt{w}^{x,z|w}_{X,Y}(t.l)
\end{aligned}$$

In the **case (b)** where $x,z\in\texttt{below}(t.l)$ and $w\in\texttt{below}(t.r)$ so the topology of the triplet must be $x,z|w$, the lowest common ancestor of $x,z$ and $w$ is $t$, i.e., $t$ is the anchor for $w,y$ in the quartet $x,z|w,y$. We then have

$$\begin{aligned}
&\sum_{\substack{x,z\in\texttt{below}(t.l)\\ w\in\texttt{below}(t.r)\\ L(z)\neq L(w)\neq X,Y\\ t(x,z,w)=x,z|w}} \texttt{I}(x,z,w)\cdot\Delta\texttt{W}(x,z|w,t)\\
=&\sum_{\substack{x,z\in\texttt{below}(t.l)\\ w\in\texttt{below}(t.r)\\ L(z)\neq L(w)\neq X,Y}} \texttt{I}(x,z,w)\cdot\texttt{exp}(\sum_{\substack{e\in x,z\to u\\ w\to t}}-\texttt{l}(e))\cdot\prod_{e\in u\to t}(1-\texttt{s}(e))\\
=&(1-\texttt{s}(t.l))\cdot\texttt{exp}(-\texttt{l}(t.r))\cdot\sum_{\substack{x,z\in\texttt{below}(t.l)\\ w\in\texttt{below}(t.r)\\ L(z)\neq L(w)\neq X,Y}} \texttt{I}(x,z,w)\cdot\texttt{exp}(\sum_{\substack{e\in x,z\to u\\ w\to t.r}}-\texttt{l}(e))\cdot\prod_{e\in u\to t.l}(1-\texttt{s}(e))
\end{aligned}$$

where

$$\begin{aligned}
&\sum_{\substack{x,z\in\texttt{below}(t.l)\\ w\in\texttt{below}(t.r)\\ L(z)\neq L(w)\neq X,Y}} \texttt{I}(x,z,w)\cdot\texttt{exp}(\sum_{\substack{e\in x,z\to u\\ w\to t.r}}-\texttt{l}(e))\cdot\prod_{e\in u\to t.l}(1-\texttt{s}(e))\\
=&\sum_{\substack{x,z\in\texttt{below}(t.l)\\ w\in\texttt{below}(t.r)\\ L(z)\neq L(w)\neq X,Y}} \Big(\texttt{I}(x,z)\cdot\texttt{exp}(\sum_{e\in x,z\to u}-\texttt{l}(e))\cdot\prod_{e\in u\to t.l}(1-\texttt{s}(e))\Big)\cdot\Big(\texttt{I}(w)\cdot\texttt{exp}(\sum_{e\in w\to t.r}-\texttt{l}(e))\Big)\\
=&\sum_{\substack{Z\in\mathcal{S}\cup\mathcal{A}\\ Z\neq X,Y}}\sum_{\substack{W\in\mathcal{S}\cup\mathcal{A}\\ W\neq X,Y,Z}}\sum_{\substack{x,z\in\texttt{below}(t.l)\\ L(x)=X,L(z)=Z}}\sum_{\substack{w\in\texttt{below}(t.r)\\ L(w)=W}} \Big(\texttt{I}(x,z)\cdot\texttt{exp}(\sum_{e\in x,z\to u}-\texttt{l}(e))\cdot\prod_{e\in u\to t.l}(1-\texttt{s}(e))\Big)\cdot\Big(\texttt{I}(w)\cdot\texttt{exp}(\sum_{e\in w\to t.r}-\texttt{l}(e))\Big)\\
=&\sum_{\substack{Z\in\mathcal{S}\cup\mathcal{A}\\ Z\neq X,Y}}\sum_{\substack{W\in\mathcal{S}\cup\mathcal{A}\\ W\neq X,Y,Z}}\Big(\sum_{\substack{x,z\in\texttt{below}(t.l)\\ L(x)=X,L(z)=Z}}\texttt{I}(x,z)\cdot\texttt{exp}(\sum_{e\in x,z\to u}-\texttt{l}(e))\cdot\prod_{e\in u\to t.l}(1-\texttt{s}(e))\Big)\cdot\Big(\sum_{\substack{w\in\texttt{below}(t.r)\\ L(w)=W}}\texttt{I}(w)\cdot\texttt{exp}(\sum_{e\in w\to t.r}-\texttt{l}(e))\Big)\\
=&\sum_{\substack{Z\in\mathcal{S}\cup\mathcal{A}\\ Z\neq X,Y}}\sum_{\substack{W\in\mathcal{S}\cup\mathcal{A}\\ W\neq X,Y,Z}}\texttt{w}^2_{X,Z}(t.l)\cdot\texttt{w}^1_W(t.r)
\end{aligned}$$

Note we can write the sum of products as the products of sums in the proof above because $Z \neq W$. Also, it is guaranteed that $z$ and $w$ are from different subtrees so there are no double counting problems. Adding the weights of the four cases together gives the final recurrence:

$$\begin{aligned}\mathtt{w}_{X,Y}^{x,z|w}(t) =& \mathtt{exp}(-\mathtt{l}(t.l)) \cdot \mathtt{w}_{X,Y}^{x,z|w}(t.l) + \mathtt{exp}(-\mathtt{l}(t.r)) \cdot \mathtt{w}_{X,Y}^{x,z|w}(t.r) \\ &+ (1-\mathtt{s}(t.l)) \cdot \mathtt{exp}(-\mathtt{l}(t.r)) \cdot \sum_{\substack{Z \in \mathcal{S}\cup\mathcal{A} \\ Z \neq X,Y}} \sum_{\substack{W \in \mathcal{S}\cup\mathcal{A} \\ W \neq X,Y,Z}} \mathtt{w}_{X,Z}^{2}(t.l) \cdot \mathtt{w}_{W}^{1}(t.r) \\ &+ (1-\mathtt{s}(t.r)) \cdot \mathtt{exp}(-\mathtt{l}(t.l)) \cdot \sum_{\substack{Z \in \mathcal{S}\cup\mathcal{A} \\ Z \neq X,Y}} \sum_{\substack{W \in \mathcal{S}\cup\mathcal{A} \\ W \neq X,Y,Z}} \mathtt{w}_{X,Z}^{2}(t.r) \cdot \mathtt{w}_{W}^{1}(t.l)\end{aligned}$$

After precomputation, $\mathtt{w}_{X,Y}^{x,z|w}(t)$ can be computed by traversing the $O(n)$ vertices of gene tree $T$ in a postorder traversal. The remainder of the complexity analysis for $\mathtt{w}_{X,Y}^{x,z|w}(t)$ is the same as for $\mathtt{w}_{X,Y}^{x|z,w}(t)$ (Lemma 5). □

#### 2.2 Quartet Graph Construction from Auxiliary Values w

We are now ready to define equations for the bad edges $\Delta\mathbb{B}_t(X,Y)$ and good edges $\Delta\mathbb{G}_t(X,Y)$ based on the six auxiliary values computed for gene tree $T$.

**Theorem 1** (bad edges below $t$)**.** *The weight of bad edges between taxa $X, Y$ below vertex $t$ in gene tree $T$ can be computed from the auxiliary values according to the following equation:*

$$\begin{aligned}\Delta\mathbb{B}_t(X,Y) =& \mathtt{w}_Y(t.r) \cdot \mathtt{exp}(-\mathtt{l}(t.r) \cdot \mathtt{w}_{X,Y}^{x|z,w}(t.l) \cdot \mathtt{exp}(-\mathtt{l}(t.l)) \\ &+ \mathtt{exp}(-\mathtt{l}(t.l)) \cdot \mathtt{w}_X(t.l) \cdot \mathtt{exp}(-\mathtt{l}(t.r)) \cdot \mathtt{w}_Y(t.r) \cdot \textstyle\sum_{\substack{Z,W \in \mathcal{S}\cup\mathcal{A} \\ Z \neq W \neq X \neq Y}} \overline{\mathtt{w}}_{Z,W}(t) \\ &+ \mathtt{w}_X(t.l) \cdot \mathtt{exp}(-\mathtt{l}(t.l) \cdot \mathtt{w}_{Y,X}^{x|z,w}(t.r) \cdot \mathtt{exp}(-\mathtt{l}(t.r))\end{aligned} \tag{16}$$

*where the summation enumerates all* unordered *pairs of taxa $Z, W \in \mathcal{S} \cup \mathcal{A}$ such that $X \neq Y \neq Z \neq W$.*

*Proof.* From Definition 1, we consider quartets $x, y|z, w$ such that $x \in \mathtt{below}(t.l)$, $y \in \mathtt{below}(t.r)$, $L(x) = X$, $L(y) = Y$, and $L(x) \neq L(y) \neq L(z) \neq L(w)$. Depending on the positions of $z$ and $w$, we divide the computation into several cases:

(a) $z, w \in \mathtt{below}(t.l)$

(b) $z \in \mathtt{below}(t.l)$ and $w \in \mathtt{above}(t)$

(c) $z, w \in \mathtt{above}(t)$

(d) $z \in \mathtt{above}(t)$ and $w \in \mathtt{below}(t.r)$ (symmetric with case (b))

(e) $z, w \in \mathtt{below}(t.r)$ (symmetric with case (a))

(f) $z \in \mathtt{below}(t.l)$ and $w \in \mathtt{below}(t.r)$

Only case (a), case (c), and case (e) result in bad edges, as both $z$ and $w$ must be from the same subtree after deleting edges on the path $x \to y$ and their endpoints from $T$. We introduce how to compute the weight of the cases (a) and (c) and the algorithm for the case (e) is similar to the case (a) according to symmetry. In the **case (a)** where $z, w \in \texttt{below}(t.l)$, we have

$$\begin{aligned}
\Delta\mathbb{B}_t^{(a)}(X,Y) =& \sum_{\substack{x\in\texttt{below}(t.l)\\ y\in\texttt{below}(t.r)\\ L(x)=X, L(y)=Y}} \sum_{\substack{z,w\in\texttt{below}(t.l)\\ L(z)\neq L(w)\neq X,Y\\ q(x,y,z,w)=x,y|z,w}} \Delta\texttt{W}(q)\cdot\texttt{I}(q)\\
=& \sum_{\substack{x\in\texttt{below}(t.l)\\ y\in\texttt{below}(t.r)\\ L(x)=X, L(y)=Y}} \sum_{\substack{z,w\in\texttt{below}(t.l)\\ L(z)\neq L(w)\neq X,Y\\ q(x,y,z,w)=x,y|z,w}} \texttt{exp}\Big(\sum_{\substack{e\in x,y\to u,\\ z,w\to v}} -\texttt{l}(e)\Big)\cdot\Big(\prod_{e\in u\to v}(1-\texttt{s}(e))\Big)\cdot\texttt{I}(x,y,z,w)\\
=&\Big(\sum_{\substack{y\in\texttt{below}(t.r)\\ L(y)=Y}} \texttt{I}(y)\cdot\texttt{exp}(\sum_{e\in y\to t} -\texttt{l}(e))\Big)\cdot\Big(\sum_{\substack{x,z,w\in\texttt{below}(t.l)\\ L(z)\neq L(w)\neq X,Y\\ t(x,z,w)=x|z,w}} \texttt{I}(x,z,w)\cdot\texttt{exp}(\sum_{\substack{e\in x,t\to u\\ z,w\to v}} -\texttt{l}(e))\cdot\prod_{e\in u\to v}(1-\texttt{s}(e))\Big)\\
=&\Big(\texttt{w}_Y^1(t.r)\cdot\texttt{exp}(-\texttt{l}(t.r)\Big)\cdot\Big(\texttt{w}_{X,Y}^{x|z,w}(t.l)\cdot\texttt{exp}(-\texttt{l}(t.l))\Big),
\end{aligned}$$

where $\texttt{w}_Y^1(t.r)$ counts the total weight of singlets below $t.r$ (which can be computed via the recurrence in Lemma 1) and $\texttt{w}_{X,Y}^{x|z,w}(t.l)$ counts the total weight of the triplets below $t.l$ whose topology is $x|z,w$ (which can be computed via the recurrence in Lemma 5). In the **case (c)**, $z$ and $w$ are in the subtree above $t$.

$$\begin{aligned}
\Delta\mathbb{B}_t^{(c)}(X,Y) =& \sum_{\substack{x\in\texttt{below}(t.l)\\ y\in\texttt{below}(t.r)\\ L(x)=X, L(y)=Y}} \sum_{\substack{z,w\in\texttt{above}(t)\\ L(z)\neq L(w)\neq X,Y\\ q(x,y,z,w)=x,y|z,w}} \Delta\texttt{W}(q)\cdot\texttt{I}(q)\\
=& \sum_{\substack{x\in\texttt{below}(t.l)\\ y\in\texttt{below}(t.r)\\ L(x)=X, L(y)=Y}} \sum_{\substack{z,w\in\texttt{above}(t)\\ L(z)\neq L(w)\neq X,Y\\ q(x,y,z,w)=x,y|z,w}} \texttt{exp}\Big(\sum_{\substack{e\in x,y\to t,\\ z,w\to u}} -\texttt{l}(e)\Big)\cdot\Big(\prod_{e\in u\to v}(1-\texttt{s}(e))\Big)\cdot\texttt{I}(x,y,z,w)\\
=&\Big(\sum_{\substack{x\in\texttt{below}(t.l)\\ L(x)=X}} \texttt{I}(x)\cdot\texttt{exp}(\sum_{e\in x\to t} -\texttt{l}(e))\Big)\cdot\Big(\sum_{\substack{y\in\texttt{below}(t.r)\\ L(y)=Y}} \texttt{I}(y)\cdot\texttt{exp}(\sum_{e\in y\to t} -\texttt{l}(e))\Big)\\
&\cdot\Big(\sum_{\substack{z,w\in\texttt{above}(t)\\ L(z)\neq L(w)\neq X,Y}} \texttt{I}(z,w)\cdot\texttt{exp}(\sum_{e\in z,w\to v} -\texttt{l}(e))\cdot\prod_{e\in v\to t}(1-\texttt{s}(e))\Big)\\
=&\texttt{exp}(-\texttt{l}(t.l))\cdot\texttt{w}_X(t.l)\cdot\texttt{exp}(-\texttt{l}(t.r))\cdot\texttt{w}_Y(t.r)\cdot\\
&\Big(\sum_{\substack{Z,W\in\mathcal{S}\cup\mathcal{A}\\ Z\neq W\neq X\neq Y}} \sum_{L(z)=Z,L(w)=W} \texttt{I}(z,w)\cdot\texttt{exp}(\sum_{e\in z,w\to v} -\texttt{l}(e))\cdot\prod_{e\in v\to t}(1-\texttt{s}(e))\Big)\\
=&\texttt{exp}(-\texttt{l}(t.l))\cdot\texttt{w}_X(t.l)\cdot\texttt{exp}(-\texttt{l}(t.r))\cdot\texttt{w}_Y(t.r)\cdot\sum_{\substack{Z,W\in\mathcal{S}\cup\mathcal{A}\\ Z\neq W\neq X\neq Y}}\overline{\texttt{w}}_{Z,W}(t).
\end{aligned}$$

where $\overline{\texttt{w}}_{X,Y}(t)$ counts the total weight of doublets in the subtree above $t$ (which can be computed via the recurrence in Lemma 3). Adding up weights from all three cases

$$\Delta\mathbb{B}_t(X,Y) = \sum_{\gamma\in\{a,c,e\}} \Delta\mathbb{B}_t^{(\gamma)}(X,Y).$$

gives us the final recurrence. □

**Theorem 2** (good edges below $t$)**.** *The weight of good edges between taxa $X, Y$ below vertex $t$ in gene tree*

*T can be derived from auxiliary values using the following equation:*

$$
\begin{aligned}
\Delta\mathbb{G}_t(X,Y) = {} & \mathtt{w}_Y^1(t.r)\cdot\mathtt{exp}(-\mathtt{l}(t.r))\cdot\mathtt{w}_{X,Y}^{x,z|w}(t.l)\cdot\mathtt{exp}(-\mathtt{l}(t.l)) \\
& + \mathtt{exp}(-\mathtt{l}(t.r))\cdot\mathtt{w}_Y^1(t.r)\cdot(1-\mathtt{s}(t.l))\cdot\textstyle\sum_{\substack{Z\in\mathcal{S}\cup\mathcal{A}\\ Z\neq X,Y}}\sum_{\substack{W\in\mathcal{S}\cup\mathcal{A}\\ W\neq X,Y,Z}}\mathtt{w}_{X,Z}^2(t.l)\cdot\overline{\mathtt{w}}_W^1(t) \\
& + (1-\mathtt{s}(t.l))\cdot(1-\mathtt{s}(t.r))\cdot\textstyle\sum_{\substack{Z\in\mathcal{S}\cup\mathcal{A}\\ Z\neq X,Y}}\sum_{\substack{W\in\mathcal{S}\cup\mathcal{A}\\ W\neq X,Y,Z}}\mathtt{w}_{X,Z}^2(t.l)\cdot\mathtt{w}_{Y,W}^2(t.r) \\
& + \mathtt{exp}(-\mathtt{l}(t.l))\cdot\mathtt{w}_X^1(t.l)\cdot(1-\mathtt{s}(t.r))\cdot\textstyle\sum_{\substack{Z\in\mathcal{S}\cup\mathcal{A}\\ Z\neq X,Y}}\sum_{\substack{W\in\mathcal{S}\cup\mathcal{A}\\ W\neq X,Y,Z}}\mathtt{w}_{Y,Z}^2(t.r)\cdot\overline{\mathtt{w}}_W^1(t) \\
& + \mathtt{w}_X^1(t.l)\cdot\mathtt{exp}(-\mathtt{l}(t.l))\cdot\mathtt{w}_{Y,X}^{x,z|w}(t.r)\cdot\mathtt{exp}(-\mathtt{l}(t.r))
\end{aligned}
\tag{17}
$$

*where $Z, W$ are ordered, as indicated by the nested summations, unlike when computing bad edges $\Delta\mathbb{B}_t(X,Y)$.*

*Proof.* From Definition 1, we consider quartets $x,z|y,w$ such that $x\in\mathtt{below}(t.l)$, $y\in\mathtt{below}(t.r)$, $L(x)=X$, $L(y)=Y$, and $L(x)\neq L(y)\neq L(z)\neq L(w)$. Depending on the positions of $z$ and $w$, we divide the computation into several cases:

(a) $z,w\in\mathtt{below}(t.l)$

(b) $z\in\mathtt{below}(t.l)$ and $w\in\mathtt{above}(t)$

(c) $z,w\in\mathtt{above}(t)$

(d) $z\in\mathtt{above}(t)$ and $w\in\mathtt{below}(t.r)$ (symmetric with case (b))

(e) $z,w\in\mathtt{below}(t.r)$ (symmetric with case (a))

(f) $z\in\mathtt{below}(t.l)$ and $w\in\mathtt{below}(t.r)$

Case (c) can be eliminated because when $z,w$ are from above $t.p$ (and $x,y$ are from below $t$), the quartet must have topology $x,y|z,w$, which corresponds to a bad edge between $X,Y$. Cases (a) and (e) are similar to each other and so are cases (b) and (d) according to the symmetry. We therefore only elaborate on the cases (a), (b), and (f). In the **case (a)**, both $z$ and $w$ are from *below*$(t.l)$ so $x$, $z$, and $w$ form a triplet in *below*$(t.l)$, whose topology has to be $x,z|w$. All triplets of $(x,z,w)$ in *below*$(t.l)$ with a topology of $x,z|w$ is able to form a valid quartet of topology $x,z|y,w$ with $y\in\mathtt{below}(t.r)$ as long as $L(z),L(w)\neq Y$. In this case, the value of $\Delta\mathbb{G}_t^{(a)}(X,Y)$ is

$$
\begin{aligned}
\Delta\mathbb{G}_t^{(a)}(X,Y) &= \sum_{\substack{x\in\mathtt{below}(t.l)\\ y\in\mathtt{below}(t.r)\\ L(x)=X,L(y)=Y}}\ \sum_{\substack{z,w\in\mathtt{below}(t.l)\\ L(z)\neq L(w)\neq X,Y\\ q(x,y,z,w)=x,z|y,w}}\Delta\mathtt{W}(q)\cdot\mathtt{I}(q) \\
&= \sum_{\substack{x\in\mathtt{below}(t.l)\\ y\in\mathtt{below}(t.r)\\ L(x)=X,L(y)=Y}}\ \sum_{\substack{z,w\in\mathtt{below}(t.l)\\ L(z)\neq L(w)\neq X,Y\\ q(x,y,z,w)=x,z|y,w}}\mathtt{exp}\Big(\sum_{\substack{e\in x,z\to u,\\ y,w\to v}}-\mathtt{l}(e)\Big)\cdot\Big(\prod_{e\in u\to v}(1-\mathtt{s}(e))\Big)\cdot\mathtt{I}(x,y,z,w) \\
&= \Big(\sum_{\substack{y\in\mathtt{below}(t.r)\\ L(y)=Y}}\mathtt{I}(y)\cdot\mathtt{exp}(\sum_{e\in y\to t}-\mathtt{l}(e))\Big) \\
&\quad\cdot\Big(\sum_{\substack{x,z,w\in\mathtt{below}(t.l)\\ L(z)\neq L(w)\neq X,Y\\ t(x,z,w)=x,z|w}}\mathtt{I}(x,z,w)\cdot\mathtt{exp}(\sum_{\substack{e\in x,z\to u\\ t,w\to v}}-\mathtt{l}(e))\cdot\prod_{e\in u\to v}(1-\mathtt{s}(e))\Big) \\
&= \Big(\mathtt{w}_Y^1(t.r)\cdot\mathtt{exp}(-\mathtt{l}(t.r))\Big)\cdot\Big(\mathtt{w}_{X,Y}^{x,z|w}(t.l)\cdot\mathtt{exp}(-\mathtt{l}(t.l))\Big)
\end{aligned}
$$

where $\mathtt{w}_Y^1(t.r)$ counts the total weight of singlets below $t_r$ (which can be computed via the recurrence in Lemma 1) and $\mathtt{w}_{X,Y}^{x,z|w}(t.l)$ counts the total weight of triplets below $t.l$ whose topology is $x,z|w$ (which can be computed via the recurrence in Lemma 6). In the **case (b)**, one of the leaves in the quartet is from the subtree above the node $t$ so the anchor node close to $y$ and $w$ is $t$. In this case, the total weight of the quartets is:

$$\begin{aligned}
\Delta\mathbb{G}_t^{(b)}(X,Y) &= \sum_{\substack{x\in\texttt{below}(t.l)\\ y\in\texttt{below}(t.r)\\ L(x)=X,L(y)=Y}} \sum_{\substack{z\in\texttt{below}(t.l)\\ w\in\texttt{above}(t)\\ L(z)\neq L(w)\neq X,Y\\ q(x,y,z,w)=x,z|y,w}} \Delta\texttt{W}(q)\cdot \texttt{I}(q) \\
&= \sum_{\substack{x\in\texttt{below}(t.l)\\ y\in\texttt{below}(t.r)\\ L(x)=X,L(y)=Y}} \sum_{\substack{z\in\texttt{below}(t.l)\\ w\in\texttt{above}(t)\\ L(z)\neq L(w)\neq X,Y\\ q(x,y,z,w)=x,z|y,w}} \texttt{exp}\Big(\sum_{\substack{e\in x,z\to u,\\ y,w\to t}} -\texttt{l}(e)\Big)\cdot\Big(\prod_{e\in u\to t}(1-\texttt{s}(e))\Big)\cdot \texttt{I}(x,y,z,w) \\
&=\Big(\sum_{\substack{y\in\texttt{below}(t.r)\\ L(y)=Y}} \texttt{I}(y)\cdot\texttt{exp}(\sum_{e\in y\to t} -\texttt{l}(e))\Big)\cdot\Big(\sum_{\substack{x,z\in\texttt{below}(t.l),w\in\texttt{above}(t)\\ L(x)=X,L(z)\neq L(w)\neq X,Y}} \texttt{I}(x,z,w)\cdot\Delta\texttt{W}(x,z|t)\cdot\texttt{exp}(\sum_{e\in w\to t}-\texttt{l}(e))\Big) \\
&=\texttt{exp}(-\texttt{l}(t.r))\cdot\texttt{w}^1_Y(t.r)\cdot(1-\texttt{s}(t.l))\cdot\Big(\sum_{\substack{x,z\in\texttt{below}(t.l),w\in\texttt{above}(t)\\ L(x)=X,L(z)\neq L(w)\neq X,Y}} \texttt{I}(x,z,w)\cdot\Delta\texttt{W}(x,z|t.l)\cdot\texttt{exp}(\sum_{e\in w\to t}-\texttt{l}(e))\Big)
\end{aligned}$$

where

$$\begin{aligned}
&\sum_{\substack{x,z\in\texttt{below}(t.l),w\in\texttt{above}(t)\\ L(x)=X,L(z)\neq L(w)\neq X,Y}} \texttt{I}(x,z,w)\cdot\Delta\texttt{W}(x,z|t.l)\cdot\texttt{exp}(\sum_{e\in w\to t}-\texttt{l}(e)) \\
&=\sum_{\substack{Z\in\mathcal{S}\cup\mathcal{A}\\ Z\neq X,Y}}\sum_{\substack{W\in\mathcal{S}\cup\mathcal{A}\\ W\neq X,Y,Z}}\sum_{\substack{x,z\in\texttt{below}(t.l)\\ L(x)=X,L(z)=Z}}\sum_{\substack{w\in\texttt{above}(t)\\ L(w)=W}} \texttt{I}(x,z,w)\cdot\Delta\texttt{W}(x,z|t.l)\cdot\texttt{exp}(\sum_{e\in w\to t}-\texttt{l}(e)) \\
&=\sum_{\substack{Z\in\mathcal{S}\cup\mathcal{A}\\ Z\neq X,Y}}\sum_{\substack{W\in\mathcal{S}\cup\mathcal{A}\\ W\neq X,Y,Z}}\sum_{\substack{x,z\in\texttt{below}(t.l)\\ L(x)=X,L(z)=Z}}\sum_{\substack{w\in\texttt{above}(t)\\ L(w)=W}} \Big(\texttt{I}(x,z)\cdot\texttt{exp}(\sum_{e\in x,z\to u}-\texttt{l}(e))\cdot(\prod_{e\in u\to t.l}(1-\texttt{s}(e)))\Big)\cdot\Big(\texttt{I}(w)\cdot\texttt{exp}(\sum_{e\in w\to t}-\texttt{l}(e))\Big) \\
&=\sum_{\substack{Z\in\mathcal{S}\cup\mathcal{A}\\ Z\neq X,Y}}\sum_{\substack{W\in\mathcal{S}\cup\mathcal{A}\\ W\neq X,Y,Z}}\Big(\sum_{\substack{x,z\in\texttt{below}(t.l)\\ L(x)=X,L(z)=Z}} \texttt{I}(x,z)\cdot\texttt{exp}(\sum_{e\in x,z\to u}-\texttt{l}(e))\cdot(\prod_{e\in u\to t.l}(1-\texttt{s}(e)))\Big)\cdot\Big(\sum_{\substack{w\in\texttt{above}(t)\\ L(w)=W}}\texttt{I}(w)\cdot\texttt{exp}(\sum_{e\in w\to t}-\texttt{l}(e))\Big) \\
&=\sum_{\substack{Z\in\mathcal{S}\cup\mathcal{A}\\ Z\neq X,Y}}\sum_{\substack{W\in\mathcal{S}\cup\mathcal{A}\\ W\neq X,Y,Z}} \texttt{w}^2_{X,Z}(t.l)\cdot\overline{\texttt{w}}^1_W(t)
\end{aligned}$$

where $\overline{\texttt{w}}^1_W(t)$ is the total weight of singlets above $t$ (which can be computed via the recurrence in Lemma 2) and $\texttt{w}^2_{X,Z}(t.l)$ is the total weight of doublets below $t.l$ (which can be computed via the recurrence in Lemma 3). As a result,

$$\Delta\mathbb{G}_t^{(c)}(X,Y)=\texttt{exp}(-\texttt{l}(t.r))\cdot\texttt{w}^1_Y(t.r)\cdot(1-\texttt{s}(t.l))\cdot\textstyle\sum_{\substack{Z\in\mathcal{S}\cup\mathcal{A}\\ Z\neq X,Y}}\sum_{\substack{W\in\mathcal{S}\cup\mathcal{A}\\ W\neq X,Y,Z}}\texttt{w}^2_{X,Z}(t.l)\cdot\overline{\texttt{w}}^1_W(t)$$

Lastly, in **case (f)**, $x$ and $z$ are from the left subtree while $y$ and $w$ are from the right subtree. In other words, we select a doublet $(x,z)$ from $v.l$ and a doublet $(y,w)$ from $v.r$ to form the quartet. As a result, we compute the total weight by multiplying the doublet weight sums in the subtrees while excluding the weights when $L(z)=L(w)$. Case (f) is special because the topology of the quartet must be $x,z|y,w$ so we have

$$\begin{aligned}
\Delta\mathbb{G}_t^{(f)}(X,Y) =& \sum_{\substack{x,z\in\texttt{below}(t.l),y,w\in\texttt{below}(t.r)\\ L(x)=X,L(y)=Y,L(z)\neq L(w)\neq X,Y\\ q(x,y,z,w)=x,z|y,w}} \Delta\texttt{W}(q)\cdot\texttt{I}(q)\\
=& \sum_{\substack{x,z\in\texttt{below}(t.l),y,w\in\texttt{below}(t.r)\\ L(x)=X,L(y)=Y,L(z)\neq L(w)\neq X,Y}} \texttt{exp}\Bigg(\sum_{\substack{e\in x,z\to u\\ y,w\to v}} -\texttt{l}(e)\Bigg)\cdot\Bigg(\prod_{e\in u\to v}(1-\texttt{s}(e))\Bigg)\cdot\texttt{I}(x,y,z,w)\\
=& \sum_{\substack{x,z\in\texttt{below}(t.l),y,w\in\texttt{below}(t.r)\\ L(x)=X,L(y)=Y,L(z)\neq L(w)\neq X,Y}} \Bigg((1-\texttt{s}(t.l))\cdot\texttt{I}(x,z)\cdot\texttt{exp}(\sum_{e\in x,z\to u}-\texttt{l}(e))\cdot\prod_{e\in u\to t.l}(1-\texttt{s}(e))\Bigg)\\
&\cdot\Bigg((1-\texttt{s}(t.r))\cdot\texttt{I}(y,w)\cdot\texttt{exp}(\sum_{e\in y,w\to v}-\texttt{l}(e))\cdot\prod_{e\in v\to t.r}(1-\texttt{s}(e))\Bigg)\\
=&(1-\texttt{s}(t.l))\cdot(1-\texttt{s}(t.r))\cdot\Bigg(\sum_{\substack{Z\in\mathcal{S}\cup\mathcal{A}\\ Z\neq X,Y}}\sum_{\substack{W\in\mathcal{S}\cup\mathcal{A}\\ W\neq X,Y,Z}} \texttt{w}^2_{X,Z}(t.l)\cdot\texttt{w}^2_{Y,Z}(t.r)\Bigg).
\end{aligned}$$

where $\texttt{w}^2_{X,Z}(t.l)$ counts the total weight of doublets below $t.l$ (which can be computed via the recurrence in Lemma 3). Adding up weights from all five cases

$$\Delta\mathbb{G}_t(X,Y) = \sum_{\gamma\in\{a,b,d,e,f\}} \Delta\mathbb{G}_t^{(\gamma)}(X,Y)$$

gives us the final recurrence. □

**Theorem 3.** *After precomputation of the six auxiliary values, $\Delta\mathbb{B}(X,Y)$ and $\Delta\mathbb{G}(X,Y)$ can be computed for all pairs of taxa $X,Y\in\mathcal{S}\cup\mathcal{A}$ in $O((a+b)^4n)$ time and $O((a+b)^2$ space.*

*Proof.* Bad and good edges between taxa $X,Y$ can be computed as

$$\Delta\mathbb{B}(X,Y) = \sum_{t\in V(T)}\Delta\mathbb{B}_t(X,Y) \text{ and } \Delta\mathbb{G}(X,Y) = \sum_{t\in V(T)}\Delta\mathbb{G}_t(X,Y) \tag{18}$$

where $\Delta\mathbb{B}_t(X,Y)$ and $\Delta\mathbb{G}_t(X,Y)$ are each solved in $O((a+b)^2)$ time with Equation 16 (Theorem 1) and Equation 17 (Theorem 2), respectively, after computing and saving the six auxiliary values. Repeating this computation for all pairs of taxa $X,Y\in\mathcal{S}\cup\mathcal{A}$ gives the time complexity result. We do not need to store $\Delta\mathbb{B}_t(X,Y)$, as it can be added to $\Delta\mathbb{B}(X,Y)$, which gives us the storage complexity result. □

#### 2.3 Time and Space Efficient Algorithms

We just introduced algorithms to compute $\Delta\mathbb{B}(X,Y)$ and $\Delta\mathbb{G}(X,Y)$. The algorithms relies heavily on precomputation of auxiliary values, which treats all pairs of taxa $X,Y$ in the same way regardless of whether they are singletons or artificial taxa. This is undesirable because we conjecture that the number $a$ of singletons will much greater than the number $b$ of artificial taxa in practice, and as we will show, singleton taxa can be treated more efficiently than artificial taxa. In the remainder of this section, we introduce time and space efficient algorithms, picking up from Section "Efficient Algorithm for Computing Weighted Bad Edges and Good for Subproblem" in Appendix B in the main text.

##### 2.3.1 Aggregation of singlets and doublets across all singleton taxa

In this section, we pick up from paragraph, "Reducing the time complexity of triplet auxiliary values by aggregating singletons," in the main text. We begin by aggregating the singlet and doublet auxiliary values across all singleton taxa, introducing special taxon label 1 to represent all singletons. We first consider extending the definition of singlet auxiliary values $\texttt{w}^1_X(t)$ (Definition 3) and $\overline{\texttt{w}}^1_X(t)$ (Definition 4) in the natural way.

**Definition 9** (singlet auxiliary values for aggregated singletons). $\mathtt{w}_1^1(t)$ *is the sum of weight from* singletons *in the subtree* below *vertex* $t$ *in gene tree* $T$. *Similarly,* $\overline{\mathtt{w}}_1^1(t)$ *is the sum of weight from* singletons *in the subtree above* $t$. *More precisely, the singlet weights of singletons* below *and* above $t$ *are defined as*

$$\mathtt{w}_1^1(t) = \sum_{\substack{x \in \mathtt{below}(t) \\ L(x) \in \mathcal{S}}} \mathtt{I}(x) \cdot \mathtt{exp}(\sum_{e \in x \to t} -\mathtt{l}(e)) \text{ and} \tag{19}$$

$$\overline{\mathtt{w}}_1^1(t) = \sum_{\substack{x \in \mathtt{above}(t) \\ L(x) \in \mathcal{S}}} \mathtt{I}(x) \cdot \mathtt{exp}(\sum_{e \in x \to t} -\mathtt{l}(e)), \text{ respectively.} \tag{20}$$

**Corollary 1** (singlet auxiliary values for aggregated singletons). $\mathtt{w}_1^1(t)$ *and* $\overline{\mathtt{w}}_1^1(t)$ *can be computed according to the following recurrences:*

$$\mathtt{w}_1^1(t) = \begin{cases} \mathtt{I}(t) & \text{if } t \text{ is a leaf in } T \\ \mathtt{exp}(-\mathtt{l}(t.l)) \cdot \mathtt{w}_1^1(t.l) + \mathtt{exp}(-\mathtt{l}(t.r)) \cdot \mathtt{w}_1^1(t.r) & \text{otherwise} \end{cases} \tag{21}$$

$$\overline{\mathtt{w}}_1^1(t) = \begin{cases} 0 & \text{if } t \text{ is the root of } T \\ \mathtt{exp}(-\mathtt{l}(t)) \cdot \overline{\mathtt{w}}_1^1(t.p) + \mathtt{exp}(-\mathtt{l}(t.s)) \cdot \mathtt{w}_1^1(t.s) & \text{otherwise} \end{cases} \tag{22}$$

*It takes* $O(n)$ *time to compute* $\mathtt{w}_1^1(t)$ *for all vertices* $t \in V(T)$. *Likewise, it takes* $O(n)$ *space to access the aggregated singlet auxiliary values later. The time and storage complexity are the same for* $\overline{\mathtt{w}}_X^1(t)$, *after computing* $\mathtt{w}_1^1(t)$.

*Proof.* Equation 21 and Equation 22 follow from Equation 4 (Lemma 1) and Equation 6 (Lemma 2). The "below" singlet is computed by traversing the $O(n)$ vertices of $T$ in a postorder fashion; after which the "above" singlet is computed by traversing the $O(n)$ vertices of $T$ in a preorder fashion. Evaluating the two recurrences for vertex $t$ takes $O(1)$ time and then storing the two auxiliary values requires $O(1)$ space. Putting this together gives the time and storage complexity. □

We continue in this fashion for doublet auxiliary values $\mathtt{w}_{X,Y}^2(t)$ and $\overline{\mathtt{w}}_{X,Y}^2(t)$, although now we could replace one or both of $X$ and $Y$ with the special taxon label 1. First, we consider the aggregating doublets when both $X$ and $Y$ are singletons.

**Definition 10** (doublet auxiliary values for two aggregated singletons). $\mathtt{w}_{1,1}^2(t)$ *is the sum of the weight from leaf pairs* $x$ *and* $y$ *in the subtree* below $t$ *in gene tree* $T$, *where* $x$ *and* $y$ *are labeled by singletons. Similarly,* $\overline{\mathtt{w}}_{1,1}^2(t)$ *is the sum of weight from leaf pairs in the subtree* above $t$. *The doublet weights of pairs of singletons* below *and* above $t$ *are defined as*

$$\mathtt{w}_{1,1}^2(t) = \sum_{\substack{x,y \in \mathtt{below}(t), x \neq y \\ L(x), L(y) \in \mathcal{S}}} \mathtt{I}(x,y) \cdot \mathtt{exp}(\sum_{e \in x,y \to u} -\mathtt{l}(e)) \cdot \prod_{e \in u \to t} (1 - \mathtt{s}(e)) \text{ and} \tag{23}$$

$$\overline{\mathtt{w}}_{1,1}^2(t) = \sum_{\substack{x,y \in \mathtt{above}(t), x \neq y \\ L(x), L(y) \in \mathcal{S}}} \mathtt{I}(x,y) \cdot \mathtt{exp}(\sum_{e \in x,y \to u} -\mathtt{l}(e)) \cdot \prod_{e \in u \to t} (1 - \mathtt{s}(e)), \tag{24}$$

*respectively, where* $u$ *is the lowest common ancestor of* $x$ *and* $y$ *for* $\mathtt{w}_{1,1}^2$ *and* $u$ *is* $lca(x,y)$, $lca(x,t)$, *or* $lca(y,t)$ *for* $\overline{\mathtt{w}}_{1,1}^2$ *depending on which is farthest from the root of* $T$.

**Corollary 2** (doublet auxiliary values for two aggregated singletons). $\mathtt{w}_{1,1}^2(t)$ *and* $\overline{\mathtt{w}}_{1,1}^2(t)$ *can be computed according to the following recurrences:*

$$\mathtt{w}_{1,1}^2(t) = \begin{cases} 0 & \text{if } t \text{ is a leaf of } T \\ (1 - \mathtt{s}(t.l)) \cdot \mathtt{w}_{1,1}^2(t.l) + (1 - \mathtt{s}(t.r)) \cdot \mathtt{w}_{1,1}^2(t.r) & \\ + \mathtt{exp}(-\mathtt{l}(t.l)) \cdot \mathtt{w}_1^1(t.l) \cdot \mathtt{exp}(-\mathtt{l}(t.r)) \cdot \mathtt{w}_1^1(t.r) & \text{otherwise} \end{cases} \tag{25}$$

$$\overline{\mathtt{w}}_{1,1}^2(t) = \begin{cases} 0 & \text{if } t \text{ is the root of } T \\ (1 - \mathtt{s}(t)) \cdot \overline{\mathtt{w}}_{1,1}^2(t.p) + (1 - \mathtt{s}(t)) \cdot (1 - \mathtt{s}(t.s)) \cdot \mathtt{w}_{1,1}^2(t.s) & \\ + (1 - \mathtt{s}(t)) \cdot \mathtt{exp}(-\mathtt{l}(t.s)) \cdot \overline{\mathtt{w}}_1^1(t.p) \cdot \mathtt{w}_1^1(t.s) & \text{otherwise} \end{cases} \tag{26}$$

*It takes $O(n)$ time to compute $\mathtt{w}^2_{1,1}(t)$ for all vertices $t \in V(T)$. Likewise, it takes $O(n)$ space to access the aggregated doublet auxiliary values later. The time and storage complexity is the same for $\overline{\mathtt{w}}^2_{1,1}(t)$, after computing $\mathtt{w}^1_1$ (Corollary 1).*

*Proof.* Equation 25 and Equation 26 are the similar as Equation 9 (Lemma 3) and Equation 11 (Lemma 11), respectively, but now instead of summing over all ways of picking two different leaves $x, y$ labeled by $X, Y$, we now sum over all ways of picking two different leaves labeled by singletons. This requires some small modifications to avoid double counting singleton taxa. The "below" doublet is computed by traversing the $O(n)$ vertices of $T$ in a postorder fashion; after which the "above" doublet is computed by traversing the $O(n)$ vertices of $T$ in a preorder fashion. Evaluating the two recurrences for vertex $t$ takes $O(1)$ time and then storing the two auxiliary values requires $O(1)$ space. Putting this together gives the time and storage complexity. □

Second, we consider the aggregating doublets when $Y$ is a singleton but not $X$.

**Definition 11** (doublet auxiliary values for one aggregated singletons)**.** *$\mathtt{w}^2_{X,1}(t) = \mathtt{w}^2_{1,X}(t)$ is the sum of the weight from leaf pairs in the subtree* below *$t$ in gene tree $T$, where one leaf is labeled by a singleton and the other is labeled by an artificial taxon $X$. Similarly, $\overline{\mathtt{w}}^2_{X,1}(t) = \overline{\mathtt{w}}^2_{1,X}(t)$ is the sum of weight from leaf pairs in the subtree* above *$t$. The doublet weights of pairs of singletons* below *and* above *$t$ are defined as*

$$\mathtt{w}^2_{X,1}(t) = \mathtt{w}^2_{1,X}(t) = \sum_{\substack{x,y \in \mathtt{below}(t) \\ L(x)=X, L(y)\in\mathcal{S}}} \mathtt{I}(x,y) \cdot \mathtt{exp}\Big(\sum_{e \in x,y \to u} -\mathtt{l}(e)\Big) \cdot \prod_{e \in u \to t} (1 - \mathtt{s}(e)) \text{ and} \tag{27}$$

$$\overline{\mathtt{w}}^2_{X,1}(t) = \overline{\mathtt{w}}^2_{1,X}(t) = \sum_{\substack{x,y \in \mathtt{above}(t) \\ L(x)=X, L(y)\in\mathcal{S}}} \mathtt{I}(x,y) \cdot \mathtt{exp}\Big(\sum_{e \in x,y \to u} -\mathtt{l}(e)\Big) \cdot \prod_{e \in u \to t} (1 - \mathtt{s}(e)), \tag{28}$$

*respectively, where $u$ is the lowest common ancestor of $x$ and $y$ for $\mathtt{w}^2_{X,1}$ and $u$ is $lca(x,y)$, $lca(x,t)$, or $lca(y,t)$ for $\overline{\mathtt{w}}^2_{X,1}$ depending on which is farthest from the root of $T$.*

**Corollary 3** (doublet auxiliary values for one aggregated singletons)**.** *$\mathtt{w}^2_{X,1}(t)$ and $\overline{\mathtt{w}}^2_{X,1}(t)$ can be computed according to the following recurrences:*

$$\mathtt{w}^2_{X,1}(t) = \mathtt{w}^2_{1,X}(t) = \begin{cases} 0 & \text{if } t \text{ is a leaf} \\ (1-\mathtt{s}(t.l)) \cdot \mathtt{w}^2_{X,1}(t.l) + (1-\mathtt{s}(t.r)) \cdot \mathtt{w}^2_{X,1}(t.r) & \\ +\mathtt{exp}(-\mathtt{l}(t.l)) \cdot \mathtt{w}^1_X(t.l) \cdot \mathtt{exp}(-\mathtt{l}(t.r)) \cdot \mathtt{w}^1_1(t.r) & \\ +\mathtt{exp}(-\mathtt{l}(t.l)) \cdot \mathtt{w}^1_1(t.l) \cdot \mathtt{exp}(-\mathtt{l}(t.r)) \cdot \mathtt{w}^1_X(t.r) & \text{otherwise} \end{cases} \tag{29}$$

$$\overline{\mathtt{w}}^2_{X,1}(t) = \overline{\mathtt{w}}^2_{1,X}(t) = \begin{cases} 0 & \text{if } t \text{ is the root} \\ (1-\mathtt{s}(t)) \cdot \overline{\mathtt{w}}^2_{X,1}(t.p) + (1-\mathtt{s}(t)) \cdot (1-\mathtt{s}(t.s)) \cdot \mathtt{w}^2_{X,1}(t.s) & \\ +(1-\mathtt{s}(t)) \cdot \mathtt{exp}(-\mathtt{l}(t.s)) \cdot \overline{\mathtt{w}}^1_X(t.p) \cdot \mathtt{w}^1_1(t.s) & \\ +(1-\mathtt{s}(t)) \cdot \mathtt{exp}(-\mathtt{l}(t.s)) \cdot \overline{\mathtt{w}}^1_X(t.s) \cdot \mathtt{w}^1_1(t.p) & \text{otherwise} \end{cases} \tag{30}$$

*After precomputation, it takes $O(bn)$ time to compute $\mathtt{w}^2_{X,1}(t)$ and $\overline{\mathtt{w}}^2_{X,1}(t)$ for all vertices $t \in V(T)$ and for all taxa $X \in \mathcal{A}$. Likewise, it takes $O(bn)$ space to access the aggregated doublet auxiliary values later.*

*Proof.* Equation 29 and Equation 30 follow from Equation 9 (Lemma 3) and Equation 11 (Lemma 11). The "below" doublet is computed by traversing the $O(n)$ vertices of $T$ in a postorder fashion; after which the "above" doublet is computed by traversing the $O(n)$ vertices of $T$ in a preorder fashion. Evaluating the two recurrences for vertex $t$ and artificial taxon $X$ takes $O(1)$ time and then storing the two auxiliary values requires $O(1)$ space. Putting this together gives the time and storage complexity. □

##### 2.3.2 Efficient algorithms for bad triplets based on aggregated singletons

Computing the auxiliary values for bad triplets is more complicated than singlets or doublets because the triplet recurrences enumerate all ways of selecting two other taxa $Z, W$ such that $Z \neq W \neq X \neq Y$. This leads to a quadratic term in the evaluation of the recurrences. To count the weights of bad triplets $\mathtt{w}_{X,Y}^{x|z,w}(t)$ via the recurrence in Equation 13 (Lemma 5), we need to evaluate the term

$$\sum_{\substack{Z,W\in\mathcal{S}\cup\mathcal{A}\\ Z\neq W\neq X\neq Y}} \mathtt{w}^2_{Z,W}(\cdot)$$

at both the left and right child of $t$. We speed up the computation for bad triplets by leveraging the doublet auxiliary quantities aggregated across singletons.

**Lemma 7.** *The quantity below can be computed in $O(b^2)$ time with the following equation:*

$$\sum_{\substack{Z,W\in\mathcal{S}\cup\mathcal{A}\\ Z\neq W\neq X,Y}} \mathtt{w}^2_{Z,W}(t) = \mathtt{w}^2_{1,1}(t) + \sum_{\substack{W\in\mathcal{A}\\ W\neq X,Y}} \mathtt{w}^2_{1,W}(t) + \sum_{\substack{Z,W\in\mathcal{A}\\ Z\neq W\neq X,Y}} \mathtt{w}^2_{Z,W}(t)$$

*after computing the doublet auxiliary values.*

*Proof.* We expand the computation of $\sum_{\substack{Z,W\in\mathcal{S}\cup\mathcal{A}\\ Z\neq W\neq X,Y}} \mathtt{w}^2_{Z,W}(t)$ into three cases: (1) both $Z$ and $W$ are both singleton taxa (i.e., $Z, W \in \mathcal{S}$), (2) one of $Z$ and $W$ is an artificial taxon and the other is a singleton taxon (w.l.o.g. $Z \in \mathcal{S}$ and $W \in \mathcal{A}$), and (3) $Z$ and $W$ are both artificial taxa (i.e., $Z, W \in \mathcal{A}$), where leaves labeled $Z$ and $W$ come from the same subtree by definition of $\mathtt{w}^2_{Z,W}(t)$. In case (1), both $Z$ and $W$ are singletons so the total weight of the doublets is $\mathtt{w}_{1,1}(t)$. In case (2), $Z$ is a singleton but $W$ is an artificial taxon, so the total weight is the sum of the $\mathtt{w}^2_{1,W}(t)$ for all $W$ in $\mathcal{A}$. In case (3), both $Z$ and $W$ are artificial so we still enumerate all unordered pairs of artificial taxa. Adding the weights of three cases together, we have

$$\begin{aligned}\sum_{\substack{Z,W\in\mathcal{S}\cup\mathcal{A}\\ Z\neq W\neq X,Y}} \mathtt{w}^2_{Z,W}(t) &= \sum_{\substack{Z,W\in\mathcal{S}\\ Z\neq W\neq X,Y}} \mathtt{w}^2_{Z,W}(t) + \sum_{\substack{Z\in\mathcal{S},W\in\mathcal{A}\\ Z\neq W\neq X,Y}} \mathtt{w}^2_{Z,W}(t) + \sum_{\substack{Z,W\in\mathcal{A}\\ Z\neq W\neq X,Y}} \mathtt{w}^2_{Z,W}(t)\\ &= \sum_{\substack{Z,W\in\mathcal{S}\\ Z\neq W\neq X,Y}} \mathtt{w}^2_{Z,W}(t) + \sum_{\substack{W\in\mathcal{A}\\ W\neq X,Y}} \sum_{\substack{Z\in\mathcal{S}\\ Z\neq X,Y}} \mathtt{w}^2_{Z,W}(t) + \sum_{\substack{Z,W\in\mathcal{A}\\ Z\neq W\neq X,Y}} \mathtt{w}^2_{Z,W}(t)\\ &= \mathtt{w}^2_{1,1}(t) + \sum_{\substack{W\in\mathcal{A}\\ W\neq X,Y}} \mathtt{w}^2_{1,W}(t) + \sum_{\substack{Z,W\in\mathcal{A}\\ Z\neq W\neq X,Y}} \mathtt{w}^2_{Z,W}(t)\end{aligned}$$

After precomputation, the time complexity of the first, second, and third terms is $O(1)$, $O(b)$, and $O(b^2)$, giving us the time complexity result. □

The proof above for $\mathtt{w}^2_{Z,W}(t)$ works for other function evaluations related to bad edges. We therefore define a function $\mathbb{P}^{\mathbb{B}}_{X,Y}[\mathtt{F}_{Z,Y}](t)$ that computes these nested summations efficiently, for example setting $\mathtt{F}_{Z,W} = \mathtt{w}^2_{Z,W}$ (which is evaluated at vertex $t$). This gives us the corollary below.

**Corollary 4.** *Let $\mathbf{P}^{\mathbb{B}}_{X,Y}[\mathtt{F}_{Z,W}](t)$ be $\sum_{\substack{Z,W\in\mathcal{S}\cup\mathcal{A}\\ Z\neq W\neq X,Y}} \mathtt{F}_{Z,W}(t)$, where $\mathtt{F}_{Z,W}$ is a mapping from a 3-tuple of two artificial taxa and a tree vertex to a weight value, i.e.,*

$$\mathtt{F}_{Z,W} : (\mathcal{S}\cup\mathcal{A}) \times (\mathcal{S}\cup\mathcal{A}) \times V(T) \to \mathbb{R},$$

*which could be $\mathtt{w}^2_{Z,W}(t)$ or $\overline{\mathtt{w}}^2_{Z,W}(t)$ in our algorithms. Then, $\mathbf{P}^{\mathbb{B}}_{X,Y}[\mathtt{F}_{Z,W}](t)$ can be computed in $O(b^2)$ time using the following equation:*

$$\mathbf{P}^{\mathbb{B}}_{X,Y}[\mathtt{F}_{Z,W}](t) = \sum_{\substack{Z,W\in\mathcal{S}\cup\mathcal{A}\\ Z\neq W\neq X,Y}} \mathtt{F}_{Z,W}(t) = \mathtt{F}_{1,1}(t) + \sum_{\substack{W\in\mathcal{A}\\ W\neq X,Y}} \mathtt{F}_{1,W}(t) + \sum_{\substack{Z,W\in\mathcal{A}\\ Z\neq W\neq X,Y}} \mathtt{F}_{Z,W}(t) \tag{31}$$

*after computing the relevant auxiliary values.*

Using Corollary 4, we can reduce the cost of computing a bad triplet for $X, Y$ below $t$ from $O((a+b)^2)$ to $O(b^2)$ by computing auxiliary values and aggregating the computation from singletons. The bottleneck now is evaluating the term enumerating the pairs of artificial taxa $Z$ and $W$ in the subtree $t$, i.e., $\mathbf{P}^{\mathbb{B}}_{X,Y}[\mathtt{F}_{Z,W}](t)$, which gives us the $O(b^2)$ term. In the following, we show how to reduce the time cost to $O(b)$.

**Lemma 8.** $\mathbf{P}^{\mathbb{B}}_{X,Y}[\mathtt{w}^2_{Z,W}](t)$ *can be computed in* $O(b)$ *time, after computing the doublet auxiliary values.*

*Proof.* According to Lemma 7, we have

$$\mathbf{P}^{\mathbb{B}}_{X,Y}[\mathtt{w}^2_{Z,W}](t) = \sum_{\substack{Z,W\in\mathcal{S}\cup\mathcal{A}\\ Z\neq W\neq X,Y}} \mathtt{w}^2_Z(t) = \mathtt{w}^2_{1,1}(t) + \sum_{\substack{W\in\mathcal{A}\\ W\neq X,Y}} \mathtt{w}^2_{1,W}(t) + \sum_{\substack{Z,W\in\mathcal{A}\\ Z\neq W\neq X,Y}} \mathtt{w}^2_{Z,W}(t)$$

where the first two terms $\mathtt{w}^2_{1,1}(t) + \sum_{\substack{W\in\mathcal{A}\\ W\neq X,Y}} \mathtt{w}^2_{1,W}(t)$ can be computed in $O(b)$ time. We rewrite the third term as

$$\sum_{\substack{Z,W\in\mathcal{A}\\ Z\neq W\neq X,Y}} \mathtt{w}^2_{Z,W}(t) = \sum_{Z,W\in\mathcal{A},Z\neq W} \mathtt{w}^2_{Z,W}(t) - \sum_{Z\in\mathcal{A}} \mathtt{w}^2_{Z,X}(t) - \sum_{Z\in\mathcal{A}} \mathtt{w}^2_{Z,Y}(t) + \mathtt{w}^2_{X,Y}(t), \tag{32}$$

The time complexity of the first, second, third, and fourth terms is $O(b^2)$, $O(b)$, $O(b)$, and $O(1)$, after precomputation. However, the first term does not depend on the selection of $X, Y$; therefore, we can compute this quantity during the precomputation phase after doublet auxiliary values but before bad triplet auxiliary values. The precomputation of the first term takes $O(b^2 n)$ time and space because there are $O(b^2)$ pairs of $Z, W$ of artificial taxa and $O(n)$ vertices $t$. Thus, it does not exceed the time and space complexity from precomputation of doublet auxiliary values even when only computing them for pairs of artificial taxa (Lemma 3).

Lastly, we do not have to exclude $X$ and/or $Y$ if they singletons. This simplifies the computation of the third term in $\mathbf{P}^{\mathbb{B}}_{X,Y}[\mathtt{w}^2_{Z,W}](t)$ to

$$\sum_{\substack{Z,W\in\mathcal{A}\\ Z\neq W\neq X,Y}} \mathtt{w}^2_{Z,W}(t) = \sum_{Z,W\in\mathcal{A},Z\neq W} \mathtt{w}^2_{Z,W}(t) - \sum_{Z\in\mathcal{A}} \mathtt{w}^2_{Z,Y}(t) \tag{33}$$

if $X$ but not $Y$ is a singleton,

$$\sum_{\substack{Z,W\in\mathcal{A}\\ Z\neq W\neq X,Y}} \mathtt{w}^2_{Z,W}(t) = \sum_{Z,W\in\mathcal{A},Z\neq W} \mathtt{w}^2_{Z,W}(t) - \sum_{Z\in\mathcal{A}} \mathtt{w}^2_{Z,X}(t) \tag{34}$$

if $Y$ but not $X$ is a singleton, and

$$\sum_{\substack{Z,W\in\mathcal{A}\\ Z\neq W\neq X,Y}} \mathtt{w}^2_{Z,W}(t) = \sum_{Z,W\in\mathcal{A},Z\neq W} \mathtt{w}^2_{Z,W}(t) \tag{35}$$

if both $X$ and $Y$ are singletons. It does not change the other computations because if $X$ and/or $Y$ are singletons we do not need to include them when drawing taxa from the set $\mathcal{A}$. □

Once again, the proof above for $\mathtt{w}^2_{Z,W}(t)$ works for other function evaluations related to bad edges. This gives us the following corollary.

**Corollary 5.** $\mathbf{P}^{\mathbb{B}}_{X,Y}[\mathtt{F}_{Z,W}](t)$ *can be computed in* $O(b)$ *time, after computing the relevant auxiliary values.*

Putting this all together enables us to reduce the time complexity of computing the bad triplet auxiliary values $\mathtt{w}^{x|z,w}_{X,Y}(t)$.

**Corollary 6** (efficient bad triplet auxiliary values)**.** $\mathtt{w}^{x|z,w}_{X,Y}(t)$ *can be computed according to the following recurrence*

$$\mathtt{w}^{x|z,w}_{X,Y}(t) = \begin{cases} 0 & \text{if } t \text{ is a leaf in } T \\ \mathtt{exp}(-\mathtt{l}(t.l)) \cdot \mathtt{w}^{x|z,w}_{X,Y}(t.l) + \mathtt{exp}(-\mathtt{l}(t.r)) \cdot \mathtt{w}^{x|z,w}_{X,Y}(t.r) & \\ +\mathtt{exp}(-\mathtt{l}(t.l)) \cdot \mathtt{w}^1_X(t.l) \cdot (1-\mathtt{s}(t.r)) \cdot \mathbf{P}^{\mathbb{B}}_{X,Y}[\mathtt{w}^2_{Z,W}](t.r) & \\ +\mathtt{exp}(-\mathtt{l}(t.r)) \cdot \mathtt{w}^1_X(t.r) \cdot (1-\mathtt{s}(t.l)) \cdot \mathbf{P}^{\mathbb{B}}_{X,Y}[\mathtt{w}^2_{Z,W}](t.l) & \text{otherwise} \end{cases} \tag{36}$$

*After computing singlet and doublet auxiliary values, it takes $O((a+b)^2bn)$ time to compute $\mathtt{w}^{x|z,w}_{X,Y}(t)$ for all vertices $t \in V(T)$ and for all unordered pairs of taxa $X, Y \in \mathcal{S} \cup \mathcal{A}$. Likewise, it takes $O((a+b)^2n)$ space to access the bad triplet auxiliary values later. The time and storage complexity drops to $O(b^3n)$ and $O(b^2n)$, respectively when restricting $X$ to be an artificial taxa (i.e., $X \in \mathcal{A}$ and $Y \in \mathcal{A} \cup \{0\}$).*

*Proof.* This correctness of the recurrence follows from Lemma 5 and Lemma 8. After precomputation, $\mathtt{w}^{x|z,w}_{X,Y}(t)$ can be computed by traversing the $O(n)$ vertices of gene tree $T$ in a postorder traversal. Evaluating the recurrence for vertex $t$ and pair of taxa $X, Y$ takes $O(b)$ time by Lemma 8, and storing the resulting auxiliary value requires $O(1)$ space. Repeating for all $O((a+b)^2)$ unordered pairs of taxa $X, Y \in \mathcal{S} \cup \mathcal{A}$ gives the time and storage complexity. □

###### 2.3.3 Efficient algorithms for good triplets based on aggregated singletons

The recurrence for good triplets also enumerates pairs of taxa, i.e., $\sum_{\substack{Z,W\in\mathcal{S}\cup\mathcal{A}\\ Z\neq W\neq X\neq Y}}\cdots$ and $\sum_{\substack{Z\in\mathcal{S}\cup\mathcal{A}\\ Z\neq X,Y}}\sum_{\substack{W\in\mathcal{S}\cup\mathcal{A}\\ W\neq X,Y,Z}}\cdots$ but with nested summations. We speed up the computation for good triplets by leveraging the doublet auxiliary quantities aggregated across singletons.

**Lemma 9.** *The quantity below can be computed in $O(b^2)$ time with the following equation:*

$$\sum_{\substack{Z\in\mathcal{S}\cup\mathcal{A}\\ Z\neq X,Y}}\sum_{\substack{W\in\mathcal{S}\cup\mathcal{A}\\ W\neq X,Y,Z}} \mathtt{w}^2_{X,Z}(t.l)\cdot \mathtt{w}^1_W(t.r) = \mathtt{w}^2_{X,1}(t.l)\cdot \mathtt{w}^1_1(t.r) + \sum_{\substack{W\in\mathcal{A}\\ W\neq X,Y}} \mathtt{w}^2_{X,1}(t.l)\cdot \mathtt{w}^1_W(t.r)$$
$$+ \sum_{\substack{Z\in\mathcal{A}\\ Z\neq X,Y}} \mathtt{w}^2_{X,Z}(t.l)\cdot \mathtt{w}^1_1(t.r) + \sum_{\substack{Z\in\mathcal{A}\\ Z\neq X,Y}}\sum_{\substack{W\in\mathcal{A}\\ W\neq X,Y,Z}} \mathtt{w}^2_{X,Z}(t.l)\cdot \mathtt{w}^1_W(t.r)$$

*after computing the singlet and doublet auxiliary values.*

*Proof.* We expand the computation of $\sum_{\substack{Z\in\mathcal{S}\cup\mathcal{A}\\ Z\neq X,Y}}\sum_{\substack{W\in\mathcal{S}\cup\mathcal{A}\\ W\neq X,Y,Z}} \mathtt{w}^2_{X,Z}(t.l)\cdot \mathtt{w}^1_W(t.r)$, where $Z$ into four cases: (1) $Z, W \in \mathcal{A}$, (2) $Z \in \mathcal{S}$ and $W \in \mathcal{A}$, (3) $Z \in \mathcal{A}$ and $W \in \mathcal{S}$, and (4) $Z, W \in \mathcal{S}$, where leaves labeled $Z$ and $W$ are from different subtrees by definition of $\mathtt{w}^1_W(t.r)$ and $\mathtt{w}^2_{X,Z}(t.l)$. Thus, cases (2) and (3) are not equivalent unlike the case of bad triplets (but they are similar due to symmetry). Breaking the computation into these cases gives us the following

$$\begin{aligned}
&\sum_{\substack{Z\in\mathcal{S}\cup\mathcal{A}\\ Z\neq X,Y}}\sum_{\substack{W\in\mathcal{S}\cup\mathcal{A}\\ W\neq X,Y,Z}} \mathtt{w}^2_{X,Z}(t.l)\cdot \mathtt{w}^1_W(t.r)\\
=&\sum_{\substack{Z\in\mathcal{S}\\ Z\neq X,Y}}\sum_{\substack{W\in\mathcal{S}\\ W\neq X,Y,Z}} \mathtt{w}^2_{X,Z}(t.l)\cdot \mathtt{w}^1_W(t.r) + \sum_{\substack{Z\in\mathcal{S}\\ Z\neq X,Y}}\sum_{\substack{W\in\mathcal{A}\\ W\neq X,Y,Z}} \mathtt{w}^2_{X,Z}(t.l)\cdot \mathtt{w}^1_W(t.r)\\
&+\sum_{\substack{Z\in\mathcal{A}\\ Z\neq X,Y}}\sum_{\substack{W\in\mathcal{S}\\ W\neq X,Y,Z}} \mathtt{w}^2_{X,Z}(t.l)\cdot \mathtt{w}^1_W(t.r) + \sum_{\substack{Z\in\mathcal{A}\\ Z\neq X,Y}}\sum_{\substack{W\in\mathcal{A}\\ W\neq X,Y,Z}} \mathtt{w}^2_{X,Z}(t.l)\cdot \mathtt{w}^1_W(t.r)\\
=&\left(\sum_{Z\in\mathcal{S}} \mathtt{w}^2_{X,Z}(t.l)\right)\cdot\left(\sum_{W\in\mathcal{S}} \mathtt{w}^1_W(t.r)\right) + \sum_{\substack{W\in\mathcal{A}\\ W\neq X,Y}}\left(\sum_{Z\in\mathcal{S}} \mathtt{w}^2_{X,Z}(t.l)\right)\cdot \mathtt{w}^1_W(t.r)\\
&+\sum_{\substack{Z\in\mathcal{A}\\ Z\neq X,Y}} \mathtt{w}^2_{X,Z}(t.l))\cdot\left(\sum_{W\in\mathcal{S}} \mathtt{w}^1_W(t.r)\right) + \sum_{\substack{Z\in\mathcal{A}\\ Z\neq X,Y}}\sum_{\substack{W\in\mathcal{A}\\ W\neq X,Y,Z}} \mathtt{w}^2_{X,Z}(t.l)\cdot \mathtt{w}^1_W(t.r)\\
=&\mathtt{w}^2_{X,1}(t.l)\cdot \mathtt{w}^1_1(t.r) + \sum_{\substack{W\in\mathcal{A}\\ W\neq X,Y}} \mathtt{w}^2_{X,1}(t.l)\cdot \mathtt{w}^1_W(t.r)\\
&+\sum_{\substack{Z\in\mathcal{A}\\ Z\neq X,Y}} \mathtt{w}^2_{X,Z}(t.l)\cdot \mathtt{w}^1_1(t.r) + \sum_{\substack{Z\in\mathcal{A}\\ Z\neq X,Y}}\sum_{\substack{W\in\mathcal{A}\\ W\neq X,Y,Z}} \mathtt{w}^2_{X,Z}(t.l)\cdot \mathtt{w}^1_W(t.r)
\end{aligned}$$

After computing the singlet and doublet auxiliary values, the time complexity of the first, second, third, and fourth term is $O(1)$, $O(b)$, $O(b)$, and $O(b^2)$, giving us the time complexity result. □

The proof for $\mathtt{w}^2_{X,Z}(t.l)$ and $\mathtt{w}^1_W(t.r)$ works for other pairs of function evaluations related to good edges. We therefore define a function $\mathbf{P}^{\mathbb{G}}_{X,Y}[\mathtt{G}_Z, \mathtt{H}_W](t_1, t_2)$ that computes these nested summations efficiently, for example setting $\mathtt{G}_Z = \mathtt{w}^2_{X,Z}$ (which will be evaluated at vertex $t1$) and $\mathtt{H}_W = \mathtt{w}^1_W$ (which will be evaluated at vertex $t2$). This gives us the corollary below.

**Corollary 7.** *Let* $\mathbf{P}^{\mathbb{G}}_{X,Y}[\mathtt{G}_Z, \mathtt{H}_W](t_1, t_2)$ *denote* $\sum_{\substack{Z\in\mathcal{S}\cup\mathcal{A}\\ Z\neq X,Y}} \sum_{\substack{W\in\mathcal{S}\cup\mathcal{A}\\ W\neq X,Y,Z}} \mathtt{G}_Z(t_1)\cdot\mathtt{H}_W(t_2)$*, where* $\mathtt{G}_Z(t_1)$ *and* $\mathtt{H}_W(t_2)$ *are both mappings from a 2-tuple of an artificial taxon and a tree vertex to a weight value, i.e.,*

$$\mathtt{G}_Z, \mathtt{H}_W : (\mathcal{S}\cup\mathcal{A}) \times V(T) \rightarrow \mathbb{R},$$

*which could be* $\mathtt{w}^1_Z(\cdot)$*,* $\overline{\mathtt{w}}^1_Z(\cdot)$*,* $\mathtt{w}^2_{C,Z}(\cdot)$*,* $\overline{\mathtt{w}}^2_{C,Z}(\cdot)$*, or* $\mathtt{p}^2_{C,Z}(\cdot)$ *in our algorithms (we treat taxon $C$ as a constant). Then,* $\mathbf{P}^{\mathbb{G}}_{X,Y}[\mathtt{G}_Z, \mathtt{H}_W](t_1, t_2)$ *can be computed in $O(b^2)$ time using the following equation:*

$$\begin{aligned}\mathbf{P}^{\mathbb{G}}_{X,Y}[\mathtt{G}_Z, \mathtt{H}_W](t_1, t_2) &= \sum_{\substack{Z\in\mathcal{S}\cup\mathcal{A}\\ Z\neq X,Y}} \sum_{\substack{W\in\mathcal{S}\cup\mathcal{A}\\ W\neq X,Y,Z}} \mathtt{G}_1(t_1)\cdot\mathtt{H}_1(t_2) = \mathtt{G}_1(t_1)\cdot\mathtt{H}_1(t_2) + \sum_{\substack{W\in\mathcal{A}\\ W\neq X,Y}} \mathtt{G}_1(t_1)\cdot\mathtt{H}_W(t_2)\\ &+ \sum_{\substack{Z\in\mathcal{A}\\ Z\neq X,Y}} \mathtt{G}_Z(t_1)\cdot\mathtt{H}_1(t_2) + \sum_{\substack{Z\in\mathcal{A}\\ Z\neq X,Y}} \sum_{\substack{W\in\mathcal{A}\\ W\neq X,Y,Z}} \mathtt{G}_Z(t_1)\cdot\mathtt{H}_W(t_2) \qquad (37)\end{aligned}$$

*after computing the relevant auxiliary values.*

Using Corollary 7, we can reduce the cost of computing a good triplet for $X, Y$ below $t$ from $O((a+b)^2)$ to $O(b^2)$ by computing auxiliary values and aggregating the computation from singletons. The bottleneck now is evaluating the term enumerating all pairs of artificial taxa $Z$ and $W$ in the subtree below $t$, which gives us the $O(b^2)$ terms. In the following, we show how to reduce the time cost to $O(b)$.

**Lemma 10.** $\mathbf{P}^{\mathbb{G}}_{X,Y}[\mathtt{w}^2_{X,Z}, \mathtt{w}^1_W](t_1, t_2)$ *can be computed in $O(b)$ time, after computing the singlet and doublet auxiliary values.*

*Proof.* According to Lemma 9, we have

$$\begin{aligned}\mathbf{P}^{\mathbb{G}}_{X,Y}[\mathtt{w}^2_{X,Z}, \mathtt{w}^1_W](t_1, t_2) &= \sum_{\substack{Z\in\mathcal{S}\cup\mathcal{A}\\ Z\neq X,Y}} \sum_{\substack{W\in\mathcal{S}\cup\mathcal{A}\\ W\neq X,Y,Z}} \mathtt{w}^2_{X,Z}(t_1)\cdot\mathtt{w}^1_1(t_2) = \mathtt{w}^2_{1,1}(t_1)\cdot\mathtt{w}^1_1(t_2) + \sum_{\substack{W\in\mathcal{A}\\ W\neq X,Y}} \mathtt{w}^2_{X,1}(t_1)\cdot\mathtt{w}^1_W(t_2)\\ &+ \sum_{\substack{Z\in\mathcal{A}\\ Z\neq X,Y}} \mathtt{w}^2_{X,Z}(t_1)\cdot\mathtt{w}^1_1(t_2) + \sum_{\substack{Z\in\mathcal{A}\\ Z\neq X,Y}} \sum_{\substack{W\in\mathcal{A}\\ W\neq X,Y,Z}} \mathtt{w}^2_{X,Z}(t_1)\cdot\mathtt{w}^1_W(t_2) \qquad (38)\end{aligned}$$

where the first three terms can be computed in $O(b)$ time. For the last term, we rewrite it as

$$\sum_{\substack{Z\in\mathcal{A}\\ Z\neq X,Y}} \sum_{\substack{W\in\mathcal{A}\\ W\neq X,Y,Z}} \mathtt{w}^2_{X,Z}(t_1)\cdot\mathtt{w}^1_W(t_2) = \sum_{Z\in\mathcal{A}, Z\neq X,Y} \mathtt{w}^2_{X,Z}(t_1)\cdot \sum_{Z\in\mathcal{A}, Z\neq X,Y} \mathtt{w}^1_Z(t_2) - \sum_{Z\in\mathcal{A}, Z\neq X,Y} \mathtt{w}^2_{X,Z}(t_1)\cdot\mathtt{w}^1_Z(t_2), \qquad (39)$$

where all three sums in the equation can be computed in $O(b)$ time. If $X$ or $Y$ is a singleton, then $Z \neq X$ or $Z \neq Y$ is always true but this does not affect the computation. □

Once again, the proof for $\mathbf{P}^{\mathbb{G}}_{X,Y}[\mathtt{w}^2_{X,Z}, \mathtt{w}^1_W](t_1, t_2)$ works for other pairs of function evaluations related to good edges besides $\mathtt{w}^2_{X,Z}(t1)$ and $\mathtt{w}^1_W(t2)$. This gives us the following corollary.

**Corollary 8.** $\mathbf{P}^{\mathbb{G}}_{X,Y}[\mathtt{G}_Z, \mathtt{H}_W](t_1, t_2)$ *can be computed in $O(b)$ time, after computing the relevant auxiliary values.*

Putting this all together enables us to reduce the time complexity of computing the good triplet weights $\mathtt{w}^{x,z|w}_{X,Y}(t)$.

**Corollary 9** (efficient good triplet auxiliary values)**.** $\mathtt{w}^{x,z|w}_{X,Y}(t)$ *can be computed according to the following recurrence:*

$$\mathtt{w}^{x,z|w}_{X,Y}(t) = \begin{cases} 0 & \text{if } t \text{ is a leaf in } T \\ \mathtt{exp}(-\mathtt{l}(t.l)) \cdot \mathtt{w}^{x,z|w}_{X,Y}(t.l) + \mathtt{exp}(-\mathtt{l}(t.r)) \cdot \mathtt{w}^{x,z|w}_{X,Y}(t.r) & \\ +(1-\mathtt{s}(t.l)) \cdot \mathtt{exp}(-\mathtt{l}(t.r)) \cdot \mathbf{P}^{\mathbb{G}}_{X,Y}[\mathtt{w}^2_{X,Z}, \mathtt{w}^1_W](t.r, t.l) & \\ +(1-\mathtt{s}(t.r)) \cdot \mathtt{exp}(-\mathtt{l}(t.l)) \cdot \mathbf{P}^{\mathbb{G}}_{X,Y}[\mathtt{w}^2_{X,Z}, \mathtt{w}^1_W](t.l, t.r) & \text{otherwise} \end{cases} \tag{40}$$

*After computing singlet and doublet auxiliary values, the time and space complexity for good triplets* $\mathtt{w}^{x,z|w}_{X,Y}(t)$ *is the same as for bad triplets* $\mathtt{w}^{x|z,w}_{X,Y}(t)$ *(Corollary 5).*

*Proof.* This correctness of the recurrence follows Lemma 6 and Lemma 10. After computing the singlet and doublet auxiliary values, $\mathtt{w}^{x,z|w}_{X,Y}(t)$ can be computed by traversing the $O(n)$ leaves of $T$ in a postorder fashion. The remainder of the time and space complexity analysis for $\mathtt{w}^{x,z|w}_{X,Y}(t)$ is the same as for bad triplets $\mathtt{w}^{x|z,w}_{X,Y}(t)$ (Corollary 5) □

###### 2.3.4 Reducing the storage and time of auxiliary values for singleton taxa

In this section, we pick up from paragraph, "Reducing the storage and time complexity of auxiliary values for singleton taxa," in the main text. We reduce the storage of auxiliary values for singleton taxa, breaking the computation into three cases:

(i) both $X$ and $Y$ are artificial taxa,

(ii) one of $X$ and $Y$ is an artificial taxa (w.l.o.g. we take $X$ to be the singleton and $Y$ to be the artificial taxon), and

(ii) both $X$ and $Y$ are singleton taxa.

As described in the main text, these cases allow us to reduce the storage and time complexity of auxiliary values for singleton taxa. We now give recurrences for computing the simplified recurrences for auxiliary values for singleton taxa, which are denoted by $\mathtt{p}$ instead of $\mathtt{w}$.

**Corollary 10** (simplified singlet auxiliary value for singleton taxa)**.** *Let $X$ be a singleton. Then,* $\mathtt{w}^1_X(t) = \mathtt{p}^1_X(t)$ *can be computed according to the following recurrence:*

$$\mathtt{p}^1_X(t) = \begin{cases} \mathtt{I}(t) & \text{if } t \text{ is a leaf in } T \text{ with } L(t) = X \in \mathcal{S} \\ \mathtt{exp}(-\mathtt{l}(t.l)) \cdot \mathtt{p}^1_X(t.l) & \text{if } X \in \mathtt{below}(t.l) \\ \mathtt{exp}(-\mathtt{l}(t.r)) \cdot \mathtt{p}^1_X(t.r) & \text{if } X \in \mathtt{below}(t.r) \\ \text{undefined} & \text{otherwise} \end{cases}. \tag{41}$$

*It takes $O(ah)$ time and $O(a)$ space to compute $\mathtt{p}^1_X(t)$ within the computation of $\Delta\mathbb{G}(X,Y)$ and $\Delta\mathbb{B}(X,Y)$ (Algorithm 4).*

*Proof.* The recurrence of $\mathtt{p}^1_X(t)$ follows from the recurrence for $\mathtt{w}^1_X(t)$ (Equation 4 in Lemma 1) but (1) replacing $\mathtt{w}^1_X(t)$ with $\mathtt{p}^1_X(t)$ and (2) simplifying based on whether $X \in \mathtt{below}(t.l)$ and $X \in \mathtt{below}(t.r)$. The recurrence for $\mathtt{p}^1_X(t)$ can be computed by traversing the $O(h)$ vertices on the path from the unique leaf labeled $X$ to the root of gene tree $T$. Evaluating the recurrence for vertex $t$ and taxon $X$ takes $O(1)$ time and $O(1)$ space. Repeating for all $O(a)$ taxa $X \in \mathcal{S}$ gives time and storage complexity of $O(ah)$. We reduce the space complexity to $O(a)$ because we only need to maintain $\mathtt{p}^1_X(t)$ at one vertex $t$ at a time during the postorder traversal of $T$ when also computing $\Delta\mathbb{G}(X,Y)$ and $\Delta\mathbb{B}(X,Y)$. □

**Corollary 11** (doublet auxiliary value for at least one singleton)**.** *Let $X$ be a singleton and $Y$ be an artificial taxon. Then, $\mathtt{w}^2_{X,Y}(t) = \mathtt{p}^2_{X,Y}(t) = \mathtt{p}^2_{Y,X}(t)$ can be computed according to the following recurrence:*

$$\mathtt{p}^2_{X,Y}(t) = \begin{cases} 0 & \text{if } L(t) = X \in \mathcal{S} \\ (1 - \mathtt{s}(t.l)) \cdot \mathtt{p}^2_{X,Y}(t.l) + \mathtt{exp}(-\mathtt{l}(t.l)) \cdot \mathtt{p}^1_X(t.l) \cdot \mathtt{exp}(-\mathtt{l}(t.r)) \cdot \mathtt{w}^1_Y(t.r) & \text{if } X \in \mathtt{below}(t.l) \\ (1 - \mathtt{s}(t.r)) \cdot \mathtt{p}^2_{X,Y}(t.r) + \mathtt{exp}(-\mathtt{l}(t.r)) \cdot \mathtt{p}^1_X(t.r) \cdot \mathtt{exp}(-\mathtt{l}(t.l)) \cdot \mathtt{w}^1_Y(t.l) & \text{if } X \in \mathtt{below}(t.r) \\ \text{undefined} & \text{otherwise} \end{cases}, \tag{42}$$

*After precomputation, it takes $O(abh)$ time and $O(ab)$ space to compute $\mathtt{p}^2_{X,Y}(t)$ within the computation of $\Delta\mathbb{G}(X,Y)$ and $\Delta\mathbb{B}(X,Y)$ (Algorithm 4). If $Y$ is also a singleton, the time and space complexity are $O(a^2h)$ time and $O(a^2)$, respectively.*

*Proof.* The recurrence for $\mathtt{p}^2_{X,Y}(t)$ follows from the recurrence for $\mathtt{w}^2_{X,Y}(t)$ (Equation 9 in Lemma 3) but (1) replacing $\mathtt{w}^1_X(t)$ with $\mathtt{p}^1_X(t)$, (2) replacing $\mathtt{w}^2_{X,Y}(t)$ with $\mathtt{p}^2_{X,Y}(t)$, and (3) simplifying based on whether $X \in \mathtt{below}(t.l)$ and $X \in \mathtt{below}(t.r)$ (etc). After precomputation, the recurrence can be computed by traversing the $O(h)$ vertices on the path from the unique labeled $X$ to the root of gene tree $T$. Evaluating the recurrence for vertex $t$ and a pair of taxa $X, Y$ takes $O(1)$ time and $O(1)$ space. Repeating for all $O(a)$ taxa $X \in \mathcal{S}$ and for all taxa $Y \in \mathcal{A}$ gives time and space complexity of $O(abh)$. We reduce the space complexity to $O(ab)$ because we only need to maintain $\mathtt{p}^2_{X,Y}(t)$ at one vertex of $t$ at a time during the postorder traversal of $T$ when also computing $\Delta\mathbb{G}(X,Y)$ and $\Delta\mathbb{B}(X,Y)$. If $Y$ is a singleton, then we simply change "for all taxa $Y \in \mathcal{A}$" to "for all taxa $Y \in \mathcal{S}$" in the analysis above. □

**Corollary 12** (efficient bad triplet auxiliary value for one singleton and one artificial taxa)**.** *Let $X$ be a singleton taxa and $Y$ be an artificial taxa. Then, $\mathtt{w}^{x|z,w}_{X,Y}(t) = \mathtt{p}^{x|z,w}_{X,Y}(t)$ can be computed according the following recurrence:*

$$\mathtt{p}^{x|z,w}_{X,Y}(t) = \begin{cases} 0 & \text{if } L(t) = X \\ \mathtt{exp}(-\mathtt{l}(t.l)) \cdot \mathtt{p}^{x|z,w}_{X,Y}(t.l) \\ \quad + \mathtt{exp}(-\mathtt{l}(t.l)) \cdot \mathtt{p}^1_X(t.l) \cdot (1 - \mathtt{s}(t.r)) \cdot \mathbf{P}^{\mathbb{B}}_{X,Y}[\mathtt{w}^2_{Z,W}](t.r) & \text{if } X \in \mathtt{below}(t.l) \\ \mathtt{exp}(-\mathtt{l}(t.r)) \cdot \mathtt{p}^{x|z,w}_{X,Y}(t.r) \\ \quad + \mathtt{exp}(-\mathtt{l}(t.r)) \cdot \mathtt{p}^1_X(t.r) \cdot (1 - \mathtt{s}(t.l)) \cdot \mathbf{P}^{\mathbb{B}}_{X,Y}[\mathtt{w}^2_{Z,W}](t.l) & \text{if } X \in \mathtt{below}(t.r) \\ \text{undefined} & \text{otherwise} \end{cases} \tag{43}$$

*After precomputation, it takes $O(ab^2h)$ time and $O(ab)$ space to compute $\mathtt{p}^{x|z,w}_{X,Y}(t)$ within the computation of $\Delta\mathbb{B}(X,Y)$ (Algorithm 4).*

*Proof.* The recurrence for $\mathtt{p}^{x|z,w}_{X,Y}$ follows Equation 36 (Corollary 6) but (1) replacing $\mathtt{w}^1_X(t)$ with $\mathtt{p}^1_X(t)$, (2) replacing $\mathtt{w}^{x|z,w}_{X,Y}(t)$ with $\mathtt{p}^{x|z,w}_{X,Y}(t)$, and (3) simplifying based on whether $X \in \mathtt{below}(t.l)$ and $X \in \mathtt{below}(t.r)$ (etc). After precomputation, the recurrence can be computed by traversing the $O(h)$ vertices on the path from the unique labeled $X$ to the root of gene tree $T$. Evaluating the recurrence for vertex $t$ and taxon $X$ takes $O(b)$ time and $O(1)$ space. Repeating for all $O(a)$ taxa $X \in \mathcal{S}$ and for all taxa $Y \in \mathcal{A}$ gives time and space complexity of $O(ab^2h)$ and $O(abh)$, respectively. We reduce the space complexity to $O(ab)$ because we only need to maintain $\mathtt{p}^{x|z,w}_{X,Y}$ at one vertex of $t$ at a time during the postorder traversal when also computing $\Delta\mathbb{G}(X,Y)$ and $\Delta\mathbb{B}(X,Y)$. □

**Corollary 13** (efficient good triplet auxiliary values for one singleton and one artificial taxa)**.** *Let $X$ be a singleton taxa and $Y$ be an artificial taxa. Then, $\mathtt{w}^{x,z|w}_{X,Y}(t) = \mathtt{p}^{x,z|w}_{X,Y}(t)$ can be computed using the following recurrence:*

$$\mathtt{p}^{x,z|w}_{X,Y}(t) = \begin{cases} 0 & \text{if } L(t) = X \\ \mathtt{exp}(-\mathtt{l}(t.l)) \cdot \mathtt{p}^{x,z|w}_{X,Y}(t.l) + (1 - \mathtt{s}(t.l)) \cdot \mathbf{P}^{\mathbb{G}}_{X,Y}[\mathtt{p}^2_{X,Z}, \mathtt{w}^1_W](t.l, t.r) & \text{if } X \in \mathtt{below}(t.l) \\ \mathtt{exp}(-\mathtt{l}(t.r)) \cdot \mathtt{p}^{x,z|w}_{X,Y}(t.r) + (1 - \mathtt{s}(t.r)) \cdot \mathbf{P}^{\mathbb{G}}_{X,Y}[\mathtt{p}^2_{X,Z}, \mathtt{w}^1_W](t.r, t.l) & \text{if } X \in \mathtt{below}(t.r) \\ \text{undefined} & \text{otherwise} \end{cases} \tag{44}$$

*After precomputation, the time and storage complexity for* $\mathtt{p}_{X,Y}^{x,z|w}(t)$ *is the same as for* $\mathtt{p}_{X,Y}^{x|z,w}(t)$ *(Corollary 12).*

*Proof.* The recurrence for $\mathtt{p}_{X,Y}^{x,z|w}$ follows Equation 40 (Corollary 9) but (1) replacing $\mathtt{w}_X^1(t)$ with $\mathtt{p}_X^1(t)$, (2) replacing $\mathtt{w}_{X,Y}^{x,z|w}(t)$ with $\mathtt{p}_{X,Y}^{x,z|w}(t)$, and (3) simplifying based on whether $X \in \mathtt{below}(t.l)$ and $X \in \mathtt{below}(t.r)$ (etc). The time and space cost analysis is the same as the auxiliary values for bad triplets (Corollary 12). □

**Corollary 14** (efficient bad and good triplet auxiliary values for two singletons)**.** *Let $X$ and $Y$ be singletons. Observe that good triplets auxiliary values $\mathtt{p}_{X,Y}^{x,z|w}(t)$ are equal for all $Y \in \mathcal{S}$ because $Y$ does not need to be excluded from the computation (Definition 15). We use $\mathtt{p}_{X,0}^{x,z|w}(t)$ to denote the bad triplet auxiliary value for any $Y \in \mathcal{S}$; it can be computed in $O(abh)$ time and $O(a)$ space. The same holds for bad triplet auxiliary values $\mathtt{p}_{X,Y}^{x|z,w}(t)$.*

*Proof.* The corollary follows from Corollary 13 but reduces the time and storage complexity by a factor of $(b)$ since we no longer need to evaluate the good triplet $\mathtt{p}_{X,Y}^{x,z|w}(t)$ for all $Y \in \mathcal{A}$ (we can select one singleton arbitrarily, denoted 0). The same holds for the bad triplet $\mathtt{p}_{X,Y}^{x,z|w}(t)$ but following Corollary 12. □

Lastly, recall that for bad triplets $\mathtt{p}_{X,Y}^{x|z,w}(t)$ does not equal $\mathtt{p}_{Y,X}^{x|z,w}(t)$ because they differ by whether the outgroup $x$ is labeled $X$ or $Y$; however, we can use the same recurrence due to symmetry swapping the ordering of the labels. The same is holds for good triplets.

##### 2.3.5 Efficient algorithms for computing $\Delta\mathbb{B}_t$ and $\Delta\mathbb{G}_t$

Finally, we incorporate our algorithmic improvements into the computation of $\Delta\mathbb{G}_t(X,Y)$ and $\Delta\mathbb{B}_t(X,Y)$, splitting it into three cases: (i) both $X$ and $Y$ are artificial taxa, (ii) one of $X$ and $Y$ is a singleton and the other is artificial (w.l.o.g. we take $X$ to be the singleton and $Y$ to be the artificial taxa), and (ii) both $X$ and $Y$ are singletons. Then we have the following corollaries:

**Corollary 15.** $\Delta\mathbb{B}_t(X,Y)$ *can be computed in $O(b)$ time according to the following equation:*

$$\Delta\mathbb{B}_t(X,Y) = \begin{cases} \mathtt{w}_Y^1(t.r)\cdot\mathtt{exp}(-\mathtt{l}(t.r)\cdot\mathtt{w}_{X,Y}^{x|z,w}(t.l)\cdot\mathtt{exp}(-\mathtt{l}(t.l)) \\ +\mathtt{exp}(-\mathtt{l}(t.l))\cdot\mathtt{w}_X^1(t.l)\cdot\mathtt{exp}(-\mathtt{l}(t.r))\cdot\mathtt{w}_Y^1(t.r)\cdot\mathbf{P}_{X,Y}^{\mathbb{B}}[\overline{\mathtt{w}}_{Z,W}^2](t) \\ +\mathtt{w}_X^1(t.l)\cdot\mathtt{exp}(-\mathtt{l}(t.l)\cdot\mathtt{w}_{Y,X}^{x|z,w}(t.r)\cdot\mathtt{exp}(-\mathtt{l}(t.r)) \\ \qquad\qquad \textit{if both } X \textit{ and } Y \textit{ are artificial} \\ \mathtt{w}_Y^1(t.r)\cdot\mathtt{exp}(-\mathtt{l}(t.r)\cdot\mathtt{p}_{X,Y}^{x|z,w}(t.l)\cdot\mathtt{exp}(-\mathtt{l}(t.l)) \\ +\mathtt{exp}(-\mathtt{l}(t.l))\cdot\mathtt{p}_X^1(t.l)\cdot\mathtt{exp}(-\mathtt{l}(t.r))\cdot\mathtt{w}_Y^1(t.r)\cdot\mathbf{P}_{X,Y}^{\mathbb{B}}[\overline{\mathtt{w}}_{Z,W}^2](t) \\ +\mathtt{p}_X^1(t.l)\cdot\mathtt{exp}(-\mathtt{l}(t.l)\cdot\mathtt{w}_{Y,0}^{x|z,w}(t.r)\cdot\mathtt{exp}(-\mathtt{l}(t.r)) \\ \qquad\qquad \textit{if } X \textit{ is a singleton and } Y \textit{ is artificial} \\ \mathtt{p}_Y^1(t.r)\cdot\mathtt{exp}(-\mathtt{l}(t.r)\cdot\mathtt{p}_{X,0}^{x|z,w}(t.l)\cdot\mathtt{exp}(-\mathtt{l}(t.l)) \\ +\mathtt{exp}(-\mathtt{l}(t.l))\cdot\mathtt{p}_X^1(t.l)\cdot\mathtt{exp}(-\mathtt{l}(t.r))\cdot\mathtt{p}_Y^1(t.r)\cdot\mathbf{P}_{X,Y}^{\mathbb{B}}[\overline{\mathtt{w}}_{Z,W}^2](t) \\ +\mathtt{p}_X^1(t.l)\cdot\mathtt{exp}(-\mathtt{l}(t.l)\cdot\mathtt{p}_{Y,0}^{x|z,w}(t.r)\cdot\mathtt{exp}(-\mathtt{l}(t.r)) \\ \qquad\qquad \textit{if both } X \textit{ and } Y \textit{ are singletons} \end{cases} \tag{45}$$

*Proof.* The corollary follows from Theorem 1 but (1) replacing $\mathtt{w}_A^1$ with $\mathtt{p}_A^1$ if $A$ is a singleton (Corollary 10), (2) replacing $\mathtt{w}_A^1$ with $\mathtt{p}_A^1$ if $A$ is a singleton (Corollary 11), (3) replacing $\mathtt{w}_{A,B}^{x|z,w}$ with $\mathtt{w}_{B,0}^{x|z,w}$ if $A$ is artificial and $B$ is singleton (Definition 7), (4) replacing $\mathtt{w}_{A,B}^{x|z,w}$ with $\mathtt{p}_{A,B}^{x|z,w}$ if $A$ is singleton and $B$ is artificial (Corollary 12), and (5) replacing with $\mathtt{w}_{A,B}^{x|z,w}$ with $\mathtt{p}_{A,0}^{x|z,w}$ if $A$ and $B$ are both singletons (Corollary 14). After precomputation, the previous quadratic term, now given by the function $\mathbf{P}_{X,Y}^{\mathbb{B}}$ can be computed according to Corollary 5 so the time cost to evaluate $\Delta\mathbb{B}_t(X,Y)$ is $O(b)$. □

**Corollary 16.** *$\Delta\mathbb{G}_t(X, Y)$ can be computed in $O(b)$ time according to the following equation:*

$$\Delta\mathbb{G}_t(X,Y) = \begin{cases} \mathtt{w}^1_Y(t.r)\cdot\mathtt{exp}(-\mathtt{l}(t.r))\cdot\mathtt{w}^{x,z|w}_{X,Y}(t.l)\cdot\mathtt{exp}(-\mathtt{l}(t.l)) \\ +\mathtt{exp}(-\mathtt{l}(t.r))\cdot\mathtt{w}^1_Y(t.r)\cdot(1-\mathtt{s}(t.l))\cdot\mathbf{P}^{\mathbb{G}}_{X,Y}[\mathtt{w}^2_{X,Z},\overline{\mathtt{w}}^1_W](t.l,t) \\ +(1-\mathtt{s}(t.l))\cdot(1-\mathtt{s}(t.r))\cdot\mathbf{P}^{\mathbb{G}}_{X,Y}[\mathtt{w}^2_{X,Z},\mathtt{w}^2_{Y,W}](t.l,t.r) \\ +\mathtt{exp}(-\mathtt{l}(t.l))\cdot\mathtt{w}^1_X(t.l)\cdot(1-\mathtt{s}(t.r))\cdot\mathbf{P}^{\mathbb{G}}_{X,Y}[\mathtt{w}^2_{Y,Z},\overline{\mathtt{w}}^1_W](t.r,t) \\ +\mathtt{w}^1_X(t.l)\cdot\mathtt{exp}(-\mathtt{l}(t.l))\cdot\mathtt{w}^{x,z|w}_{Y,X}(t.r)\cdot\mathtt{exp}(-\mathtt{l}(t.r)) \\ \qquad\qquad\qquad\qquad \textit{if both } X \textit{ and } Y \textit{ are artificial} \\ \mathtt{w}^1_Y(t.r)\cdot\mathtt{exp}(-\mathtt{l}(t.r))\cdot\mathtt{p}^{x,z|w}_{X,Y}(t.l)\cdot\mathtt{exp}(-\mathtt{l}(t.l)) \\ +\mathtt{exp}(-\mathtt{l}(t.r))\cdot\mathtt{w}^1_Y(t.r)\cdot(1-\mathtt{s}(t.l))\cdot\mathbf{P}^{\mathbb{G}}_{X,Y}[\mathtt{p}^2_{X,Z},\overline{\mathtt{w}}^1_W](t.l,t) \\ +(1-\mathtt{s}(t.l))\cdot(1-\mathtt{s}(t.r))\cdot\mathbf{P}^{\mathbb{G}}_{X,Y}[\mathtt{p}^2_{X,Z},\mathtt{w}^2_{Y,W}](t.l,t.r) \\ +\mathtt{exp}(-\mathtt{l}(t.l))\cdot\mathtt{p}^1_X(t.l)\cdot(1-\mathtt{s}(t.r))\cdot\mathbf{P}^{\mathbb{G}}_{X,Y}[\mathtt{w}^2_{Y,Z},\overline{\mathtt{w}}^1_W](t.r,t) \\ +\mathtt{p}^1_X(t.l)\cdot\mathtt{exp}(-\mathtt{l}(t.l))\cdot\mathtt{w}^{x,z|w}_{Y,0}(t.r)\cdot\mathtt{exp}(-\mathtt{l}(t.r)) \\ \qquad\qquad\qquad\qquad \textit{if } X \textit{ is a singleton and } Y \textit{ is artificial} \\ \mathtt{p}^1_Y(t.r)\cdot\mathtt{exp}(-\mathtt{l}(t.r))\cdot\mathtt{p}^{x,z|w}_{X,0}(t.l)\cdot\mathtt{exp}(-\mathtt{l}(t.l)) \\ +\mathtt{exp}(-\mathtt{l}(t.r))\cdot\mathtt{p}^1_Y(t.r)\cdot(1-\mathtt{s}(t.l))\cdot\mathbf{P}^{\mathbb{G}}_{X,Y}[\mathtt{p}^2_{X,Z},\overline{\mathtt{w}}^1_W](t.l,t) \\ +(1-\mathtt{s}(t.l))\cdot(1-\mathtt{s}(t.r))\cdot\mathbf{P}^{\mathbb{G}}_{X,Y}[\mathtt{p}^2_{X,Z},\mathtt{p}^2_{Y,W}](t.l,t.r) \\ +\mathtt{exp}(-\mathtt{l}(t.l))\cdot\mathtt{p}^1_X(t.l)\cdot(1-\mathtt{s}(t.r))\cdot\mathbf{P}^{\mathbb{G}}_{X,Y}[\mathtt{p}^2_{Y,Z},\overline{\mathtt{w}}^1_W](t.r,t) \\ +\mathtt{p}^1_X(t.l)\cdot\mathtt{exp}(-\mathtt{l}(t.l))\cdot\mathtt{p}^{x,z|w}_{Y,0}(t.r)\cdot\mathtt{exp}(-\mathtt{l}(t.r)) \\ \qquad\qquad\qquad\qquad \textit{if both } X \textit{ and } Y \textit{ are singletons} \end{cases} \tag{46}$$

*Proof.* The corollary follows from Theorem 2 but (1) replacing $\mathtt{w}^1_A$ with $\mathtt{p}^1_A$ if $A$ is a singleton (Corollary 10), (2) replacing $\mathtt{w}^1_A$ with $\mathtt{p}^1_A$ if $A$ is a singleton (Corollary 11), (3) replacing $\mathtt{w}^{x,z|w}_{A,B}$ with $\mathtt{w}^{x,z|w}_{B,0}$ if $A$ is artificial and $B$ is singleton (Definition 8), (4) replacing $\mathtt{w}^{x,z|w}_{A,B}$ with $\mathtt{p}^{x,z|w}_{A,B}$ if $A$ is singleton and $B$ is artificial (Corollary 13), and (5) replacing with $\mathtt{w}^{x,z|w}_{A,B}$ with $\mathtt{p}^{x,z|w}_{A,0}$ if $A$ and $B$ are both singletons (Corollary 14). After precomputation, the previous quadratic term, now given by the function $\mathbf{P}^{\mathbb{G}}_{X,Y}$ can be computed according to Corollary 8 so the time cost to evaluate $\Delta\mathbb{G}_t(X, Y)$ is $O(b)$. □

#### 2.4 Pseudocode

---

**Algorithm 2:** Weighted Quartet Graph Construction

---

**Input:** an input gene tree $T$
**Output:** the quartet graph represented by matrices $\mathbb{G}$ and $\mathbb{B}$

$T_1 \leftarrow$ a copy of $T$ with all branch support values being 0
arbitrarily resolve $T$ and $T_1$ (and set support and length values of new edges to 0)
$\mathbb{G}_1$, $\mathbb{B}_1 \leftarrow$ `edge-weight` $(T_1)$
$\Delta\mathbb{G}$, $\Delta\mathbb{B} \leftarrow$ `edge-weight` $(T)$
**return** $\mathbb{G}_1 - \Delta\mathbb{G}$, $\mathbb{B}_1 - \Delta\mathbb{B}$

---

**Algorithm 3:** Precomputation of auxiliary values for artificial taxa

**Input:** an input gene tree $T$

**Output:** all auxiliary values: $\mathtt{w}^1, \mathtt{w}^2, \overline{\mathtt{w}}^1, \overline{\mathtt{w}}^2, \mathtt{w}^{x|z,w}, \mathtt{w}^{x,z|w}$ and the precomputed *term* for evaluating $\mathbf{P}^{\mathbb{B}}$ efficiently according to Corollary 5

**for** *each node $t$ in a postordering of $T$* **do**
    $\mathtt{w}^1_1(t) \leftarrow$ Equation 21
    **for** *each $X$ in $\mathcal{A}$* **do**
        $\mathtt{w}^1_X(t) \leftarrow$ Equation 4
    $\mathtt{w}^2_{1,1}(t) \leftarrow$ Equation 25
    **for** *each $X$ in $\mathcal{A}$* **do**
        $\mathtt{w}^2_{1,X} = \mathtt{w}^2_{X,1} \leftarrow$ Equation 29
    **for** *each $(X, Y)$ in $\mathcal{A} \times \mathcal{A}$* **do**
        **if** $X \neq Y$ **then** $\mathtt{w}^2_{X,Y}(t) \leftarrow$ Equation 9

**for** *each vertex $t$ in a preorder traversal of $T$* **do**
    $\overline{\mathtt{w}}^1_1(t) \leftarrow$ Equation 22
    **for** *each $X$ in $\mathcal{A}$* **do**
        $\overline{\mathtt{w}}^1_X(t) \leftarrow$ Equation 6
    $\overline{\mathtt{w}}^2_{1,1}(t) \leftarrow$ Equation 26
    **for** *each $X$ in $\mathcal{A}$* **do**
        $\overline{\mathtt{w}}^2_{1,X} = \overline{\mathtt{w}}^2_{X,1} \leftarrow$ Equation 30
    **for** *each $(X, Y)$ in $\mathcal{A} \times \mathcal{A}$* **do**
        **if** $X \neq Y$ **then** $\overline{\mathtt{w}}^2_{X,Y}(t) \leftarrow$ Equation 11

**for** *each vertex $t$ in a postorder traversal of $T$* **do**
    *term*$(t) \leftarrow 0$ **for** *each $(X, Y)$ in $\mathcal{A} \times \mathcal{A} \cup \{0\}$* **do**
        **if** $X \neq Y$ **then**
            $\mathtt{w}^{x|z,w}_{X,Y}(t) \leftarrow$ Equation 36
            $\mathtt{w}^{x,z|w}_{X,Y}(t) \leftarrow$ Equation 40
            **if** $Y \neq 0$ **then** *term*$(t) \leftarrow$ *term*$(t) + w^2_{X,Y}(t)$

**return** $\mathtt{w}^1, \mathtt{w}^2, \overline{\mathtt{w}}^1, \overline{\mathtt{w}}^2, \mathtt{w}^{x|z,w}, \mathtt{w}^{x,z|w}$, *term*

**Algorithm 4: Good and Bad Edge Weight Computation.** Recall that the auxiliary quantities simplified for singleton taxa, denoted, $\mathtt{p}$ and are computed via a postorder traversal, along with $\Delta\mathbb{B}$ and $\Delta\mathbb{G}$. The goal is to reduce storage complexity for auxiliary values. At the leaves, there is one $\mathtt{p}$ auxiliary value for every singleton, and they are moved up tree during the postorder traversal. As an example, when we compute $\mathtt{p}^1_X(t)$, we can either overwrite $\mathtt{p}^1_X(t.l)$ or $\mathtt{p}^1_X(t.r)$, depending on whether $X \in \{L(x) : x \in \mathtt{below}(t.l)\}$ or $X \in \{L(x) : x \in \mathtt{below}(t.r)\}$ (only one is possible).

**Input:** an input gene tree $T$

**Output:** weights of good and bad edges, stored in the matrices $\Delta\mathbb{G}$ and $\Delta\mathbb{B}$

$\mathtt{w}^1, \mathtt{w}^2, \overline{\mathtt{w}}^1, \overline{\mathtt{w}}^2, \mathtt{w}^{x|z,w}, \mathtt{w}^{x,z|w}, term \leftarrow \mathtt{pre\text{-}processing}(T)$

**for** *each vertex $t$ in a postorder traversal of $T$* **do**

  **for** *each* $(X, Y)$ *in* $\{L(x) : x \in \mathtt{below}(t.l)\} \times \{L(x) : x \in \mathtt{below}(t.r)\}$ **do**

    **if** $X \neq Y$ **then**

      $\Delta\mathbb{G}_t(X, Y) \leftarrow$ Equation 46

      $\Delta\mathbb{B}_t(X, Y) \leftarrow$ Equation 45

      $\Delta\mathbb{G}(X, Y) \leftarrow \Delta\mathbb{G}(X, Y) + \Delta\mathbb{G}_t(X, Y)$

      $\Delta\mathbb{B}(X, Y) \leftarrow \Delta\mathbb{B}(X, Y) + \Delta\mathbb{B}_t(X, Y)$

  **for** *each* $X$ *in* $\{L(x) : x \in \mathtt{below}(t.l) \cup \mathtt{below}(t.r)\}$ **do**

    **if** $X \in \mathcal{S}$ **then**

      $\mathtt{p}^1_X(t) \leftarrow$ Equation 41, overwrite $\mathtt{p}^1_X(t.l)$ or $\mathtt{p}^1_X(t.r)$

      **for** *each* $Y \in \mathcal{A} \cup \{1\}$ **do**

        $\mathtt{p}^2_{X,Y}(t) \leftarrow$ Equation 42, overwrite $\mathtt{p}^2_{X,Y}(t.l)$ or $\mathtt{p}^2_{X,Y}(t.r)$

      $\mathtt{p}^{x|z,w}_{X,0}(t) \leftarrow$ Equation 43, overwrite $\mathtt{p}^{x|z,w}_{X,0}(t.l)$ or $\mathtt{p}^{x|z,w}_{X,0}(t.r)$

      $\mathtt{p}^{x,z|w}_{X,0}(t) \leftarrow$ Equation 44, overwrite $\mathtt{p}^{x,z|w}_{X,0}(t.l)$ or $\mathtt{p}^{x,z|w}_{X,0}(t.r)$

      **for** *each* $Y \in \mathcal{A}$ **do**

        $\mathtt{p}^{x|z,w}_{X,Y}(t) \leftarrow$ Equation 43, overwrite $\mathtt{p}^{x|z,w}_{X,Y}(t.l)$ or $\mathtt{p}^{x|z,w}_{X,Y}(t.r)$

        $\mathtt{p}^{x,z|w}_{X,Y}(t) \leftarrow$ Equation 44, overwrite $\mathtt{p}^{x,z|w}_{X,Y}(t.l)$ or $\mathtt{p}^{x,z|w}_{X,Y}(t.r)$

**return** $\Delta\mathbb{G}$, $\Delta\mathbb{B}$

#### 2.5 Final time and space complexity results

**Theorem 4.** *Let $n$ be the number leaves in the gene tree $T$, which is rooted arbitrarily with height $h$. Let $s = a + b$ be the number of leaf labels for the current subproblem, where $a$ is the number of singletons and $b$ is the number of non-singletons (or artificial taxa). Then, we can compute $\mathbb{G}$ and $\mathbb{B}$ correctly in $O(a^2b + ab^2n + b^3n)$ time and $O(a^2 + ab + b^2n)$ space using Algorithm 2.*

*Proof.* Correctness follows from Corollaries 15 and 16 because the modifications to the equations only change the computational efficiency not the results.

To construct the quartet graph from gene tree $T$ (Algorithm 2), we apply Algorithm 4 to the gene tree two times and the combines the results into the final graph . Therefore, we analyze the space and time complexity of Algorithm 4.

Algorithm 4 first computes the auxiliary values during a precomputation phase (Algorithm 3). Specifically, the auxiliary values $\mathtt{w}^1_X(t)$, $\mathtt{w}^2_{X,Y}(t)$, $\overline{\mathtt{w}}^1_X(t)$, $\overline{\mathtt{w}}^2_{X,Y}(t)$, $\mathtt{w}^{x|z,w}_{X,Y}(t)$, and $\mathtt{w}^{x,z|w}_{X,Y}(t)$ are computed for artificial taxa (plus the special taxa 0 and 1) and for all vertex $t \in V(T)$. We split the cost of the procomputation into three cases. In each case, the cost is determined by the number of values computed and the time cost to compute each value, i.e., the time complexity is $O(A) \times O(B) = O(AB)$ if there are $O(A)$ values and each value takes $O(B)$ time to compute. The cost of each auxiliary value is

- $\mathtt{w}^1_X(t)$ and $\overline{\mathtt{w}}^1_X(t)$: $O(bn) \times O(1) = O(bn)$, according to Lemmas 1–2 and Corollaries 1,
- $\mathtt{w}^2_{X,Y}(t)$ and $\overline{\mathtt{w}}^2_{X,Y}(t)$: $O(b^2n) \times O(1) = O(b^2n)$, according to Lemmas 3–4 and Corollaries 2–3
- $\mathtt{w}^{x|z,w}_{X,Y}(t)$ and $\mathtt{w}^{x,z|w}_{X,Y}(t)$: $O(b^2n) \times O(b) = O(b^3n)$, according to Corollaries 6 and 9.

Putting this together, the time and space complexity of precomputation for auxiliary values $\mathtt{w}$ is $O(b^3n)$ and $O(b^2n)$, respectively.

After the precomputation phase, Algorithm 4 computes $\Delta\mathbb{G}_t(X, Y)$ and $\Delta\mathbb{B}_t(X, Y)$, which are defined such that $X$ labels leaves below $t.l$ and $Y$ labels leaves below $t.r$. The time to compute $\Delta\mathbb{G}_t(X, Y)$ and $\Delta\mathbb{B}_t(X, Y)$ is determined by the number of $(X, Y, t)$ 3-tuples and the cost to process each 3-tuple. The number of $(X, Y, t)$ 3-tuples is $O(a^2 + abh + b^2n)$, where $h$ is the height of $T$. To show this, we split the problem into three cases and consider the number of $t$:

- $t$ is unique if both $X$ and $Y$ are singletons so the number of 3-tuples is $O(a^2) \times O(1) = O(a^2)$,
- the number of $t$ is $O(h)$ if one of $X$ and $Y$ is artificial so the number of 3-tuples is $O(ab) \times O(h) = O(abh)$,
- the number of $t$ is $O(n)$ if both $X$ and $Y$ are artificial so the number of 3-tuple is $O(b^2) \times O(n) = O(b^2n)$.

Adding the three cases up, we show the total number of 3-tuples is $O(a^2 + abh + b^2n)$. Given a 3-tuple of $(X, Y, t)$, both $\Delta\mathbb{G}_t(X, Y)$ and $\Delta\mathbb{B}_t(X, Y)$ takes $O(b)$ time to compute according to Corollaries 16-15, so the total time complexity of the procedure is $O(a^2 + abh + b^2n) \times O(b) = O(a^2b + ab^2h + b^3n)$. No extra space is needed because the quartet graph itself $O((a + b)^2) = O(a^2 + ab + b^2)$ because we update the weights of the good and bad edges in place.

Lastly, while computing $\Delta\mathbb{G}$ and $\Delta\mathbb{B}$ in a postorder traversal of $T$, we also compute $\mathtt{p}^1_X(t)$, $\mathtt{p}^2_{X,Y}(t)$, $\mathtt{p}^{x|z,w}_{X,Y}(t)$, and $\mathtt{p}^{x,z|w}_X(t)$. The cost of each quantity is

- $\mathtt{p}^1_X(t)$: $O(a) \times O(h) = O(a)$, according to Corollary 10,
- $\mathtt{p}^2_{X,Y}(t)$: $O(ab) \times O(h) = O(abh)$, according to Corollary 11,
- $\mathtt{p}^{x|z,w}_{X,Y}(t)$ and $\mathtt{p}^{x,z|w}_{X,Y}(t)$: $O(ab) \times O(bh) = O(ab^2h)$, according to Corollaries 12 and 13.

The storage cost of the $\mathtt{p}$ values is no greater than $O(ab)$ because they are updated on the fly. Consequently, computing the auxiliary values for singletons $\mathtt{p}(\cdot)$ does not affect the time or space complexity of Algorithm 4.

The final time complexity of Algorithm 2 is $O(a^2b+ab^2h+b^3n)$ by putting the cost of precomputation and edge weight computation together. The final space time complexity is $O(a^2 + ab + b^2n)$ because we combine the storage for the quartet graph $O((a+b)^2) = O(a^2+ab+b^2)$ with the storage from precomputation $O(b^2n)$

Although they do not impact the storage complexity, for completeness we mention storage for the subproblem forest data structure $O(n)$, which must be maintained across subproblems on the stack (for total of $O(n^2)$ which is the same as the quartet graph when $a = n$) and storage for the input gene trees $O(nk)$. □

**Theorem 5.** *Let $n$ be the number of species, and let $k$ be the number of gene trees. weighted TREE-QMC has time complexity of $O(n^4k)$ if subproblems are produced in a perfectly balanced fashion, and moreover a time complexity of $O(n^2 \log n \cdot k)$ if we assume the number of artificial taxa in each subproblem is a constant.*

*Proof.* At each step in the divide phase of the algorithm, we compute a bipartition and then recurse on two subproblems. Let $s$ be the number of taxa in the subproblem, i.e., $s = a + b$. The work per subproblem has time complexity $O(s^3nk + s^2) = O(s^3nk)$ where the first term comes from constructing the quartet graph by applying our algorithm to all gene trees ($O(a^2b + ab^2h + b^3n) = O(s^3n)$ by Theorem 4) and the second term comes from applying our approach for seeking a max-cut (Theorem 2 in [3]).

To proceed with the analysis, we assume the subproblem decomposition is perfectly balanced, as in Theorem 3 in [3]. Under this assumption, $T(s) = 2T(s/2)+O(s^3nk)$, and by the master theorem we get by we have $T(s) = O(s^3nk)$ for divide-and-conquer recurrence. This evaluates to $O(n^4k)$ if $s = n$. If we also assume the number of artificial taxa $b$ is a constant, the work per subproblem becomes $O((a^2+ah+n)k+s^2) = O(s^2k+snk)$. Then we have $T(s) = 2T(s/2) + O(s^2k + snk)$. By splitting the $T(s)$ into $T_1(s) = 2T_1(s/2) + O(s^2k)$ and $T_2(s) = 2T_2(s/2)+O(snk)$, we obtain $T_1(s) = O(s^2k)$ and $T_2(s) = O(s \log s \cdot nk)$ by the master theorem so the total time is $T_1(s) + T_2(s) = O(s^2k + s \log s \cdot nk)$. This evaluates to $O(n^2 \log n \cdot k)$ when $s = n$.

□

### 3 Supplemental Experimental Study

#### 3.1 Properties of Simulated Data

Table S1: **Properties of Asteroid simulated data from** [7]**.** Data were downloaded from https://cme.h-its.org/exelixis/material/asteroid_data.tar.gz in December 2022. For each replicate, ILS is the fraction of branches in true species tree missing from the true gene tree, averaged across all gene trees. GTEE is the fraction of branches in true gene tree that are missing from the estimated gene tree, averaged across all gene trees. AD is the fraction of branches in the true species tree that are missing from the estimated gene tree, averaged across all gene trees. The values in the table are the average (± standard deviation) across all replicates. Dup indicate the row is duplicated.

| | | ILS | GTEE | AD | Proportion missing taxa |
|---|---|---|---|---|---|
| Varying population size | 10 | $0.0000 \pm 0.0000$ | $0.3419 \pm 0.0453$ | $0.3419 \pm 0.0453$ | $0.8115 \pm 0.0107$ |
| | 50000000 | $0.0610 \pm 0.0297$ | $0.3530 \pm 0.0483$ | $0.3616 \pm 0.0496$ | $0.8106 \pm 0.0118$ |
| | 100000000 | $0.1119 \pm 0.0359$ | $0.3598 \pm 0.0440$ | $0.3836 \pm 0.0478$ | $0.8109 \pm 0.0115$ |
| | 500000000 | $0.3854 \pm 0.0722$ | $0.3904 \pm 0.0413$ | $0.5428 \pm 0.0582$ | $0.8131 \pm 0.0099$ |
| | 1000000000 | $0.5576 \pm 0.0626$ | $0.4071 \pm 0.0395$ | $0.6579 \pm 0.0501$ | $0.8118 \pm 0.0089$ |
| Varying number of taxa | 25 | $0.0492 \pm 0.0314$ | $0.3008 \pm 0.0456$ | $0.3085 \pm 0.0485$ | $0.7607 \pm 0.0104$ |
| | 75 | $0.0626 \pm 0.0225$ | $0.3872 \pm 0.0382$ | $0.3960 \pm 0.0388$ | $0.8255 \pm 0.0088$ |
| | 50 | $0.0610 \pm 0.0297$ | $0.3530 \pm 0.0483$ | $0.3616 \pm 0.0496$ | $0.8106 \pm 0.0118$ (dup) |
| | 100 | $0.0733 \pm 0.0270$ | $0.4125 \pm 0.0442$ | $0.4220 \pm 0.0454$ | $0.8320 \pm 0.0076$ |
| | 125 | $0.0835 \pm 0.0274$ | $0.4384 \pm 0.0397$ | $0.4487 \pm 0.0417$ | $0.8340 \pm 0.0073$ |
| | 150 | $0.0841 \pm 0.0284$ | $0.4524 \pm 0.0362$ | $0.4625 \pm 0.0371$ | $0.8370 \pm 0.0073$ |
| Varying number of genes | 250 | $0.0602 \pm 0.0256$ | $0.3504 \pm 0.0487$ | $0.3584 \pm 0.0500$ | $0.8127 \pm 0.0100$ |
| | 500 | $0.0592 \pm 0.0264$ | $0.3527 \pm 0.0435$ | $0.3603 \pm 0.0448$ | $0.8110 \pm 0.0094$ |
| | 1000 | $0.0610 \pm 0.0297$ | $0.3530 \pm 0.0483$ | $0.3616 \pm 0.0496$ | $0.8106 \pm 0.0118$ (dup) |
| | 2000 | $0.0618 \pm 0.0256$ | $0.3496 \pm 0.0464$ | $0.3582 \pm 0.0478$ | $0.8116 \pm 0.0101$ |
| Varying sequence length | 50 | $0.0618 \pm 0.0264$ | $0.4578 \pm 0.0452$ | $0.4628 \pm 0.0455$ | $0.8113 \pm 0.0093$ |
| | 100 | $0.0610 \pm 0.0297$ | $0.3530 \pm 0.0483$ | $0.3616 \pm 0.0496$ | $0.8106 \pm 0.0118$ (dup) |
| | 200 | $0.0586 \pm 0.0282$ | $0.2514 \pm 0.0454$ | $0.2634 \pm 0.0480$ | $0.8098 \pm 0.0100$ |
| | 500 | $0.0614 \pm 0.0273$ | $0.1571 \pm 0.0374$ | $0.1779 \pm 0.0423$ | $0.8135 \pm 0.0100$ |
| Varying branch length scaler | 0.05 | $0.0600 \pm 0.0274$ | $0.5530 \pm 0.0610$ | $0.5559 \pm 0.0612$ | $0.8135 \pm 0.0093$ |
| | 0.10 | $0.0608 \pm 0.0282$ | $0.4686 \pm 0.0555$ | $0.4722 \pm 0.0550$ | $0.8118 \pm 0.0090$ |
| | 1.00 | $0.0610 \pm 0.0297$ | $0.3530 \pm 0.0483$ | $0.3616 \pm 0.0496$ | $0.8106 \pm 0.0118$ (dup) |
| | 10.00 | $0.0600 \pm 0.0262$ | $0.4280 \pm 0.0412$ | $0.4347 \pm 0.0414$ | $0.8125 \pm 0.0097$ |
| | 100.00 | $0.0599 \pm 0.0270$ | $0.7035 \pm 0.0382$ | $0.7050 \pm 0.0378$ | $0.8126 \pm 0.0102$ |
| | 200.00 | $0.0604 \pm 0.0258$ | $0.7664 \pm 0.0322$ | $0.7674 \pm 0.0319$ | $0.8109 \pm 0.0098$ |
| Varying missingness parameter | 0.50 | $0.0767 \pm 0.0281$ | $0.3818 \pm 0.0430$ | $0.3936 \pm 0.0441$ | $0.7373 \pm 0.0111$ |
| | 0.55 | $0.0659 \pm 0.0265$ | $0.3644 \pm 0.0467$ | $0.3744 \pm 0.0481$ | $0.7792 \pm 0.0124$ |
| | 0.60 | $0.0610 \pm 0.0297$ | $0.3530 \pm 0.0483$ | $0.3616 \pm 0.0496$ | $0.8106 \pm 0.0118$ (dup) |
| | 0.65 | $0.0548 \pm 0.0250$ | $0.3340 \pm 0.0449$ | $0.3418 \pm 0.0459$ | $0.8378 \pm 0.0088$ |
| | 0.70 | $0.0471 \pm 0.0250$ | $0.3200 \pm 0.0462$ | $0.3263 \pm 0.0468$ | $0.8617 \pm 0.0093$ |
| | 0.75 | $0.0447 \pm 0.0267$ | $0.3023 \pm 0.0538$ | $0.3066 \pm 0.0539$ | $0.8789 \pm 0.0048$ |

Table S2: **Properties of S100 simulated data from** [10] **and also** [9]**.** Number of refinements is the number of refinements needed to make FastTrees binary (they can be non-binary due to identical sequences).

| Sequence length | ILS | GTEE | AD | # of refinements |
|---|---|---|---|---|
| 200 | $0.4575 \pm 0.0601$ | $0.5546 \pm 0.0792$ | $0.6634 \pm 0.0652$ | $2.6337 \pm 3.9622$ |
| 400 | $0.4575 \pm 0.0601$ | $0.4229 \pm 0.0823$ | $0.5894 \pm 0.0653$ | $0.8408 \pm 1.5477$ |
| 800 | $0.4575 \pm 0.0601$ | $0.3115 \pm 0.0789$ | $0.5385 \pm 0.0628$ | $0.2706 \pm 0.6069$ |
| 1600 | $0.4575 \pm 0.0601$ | $0.2258 \pm 0.0735$ | $0.5070 \pm 0.0611$ | $0.0972 \pm 0.2746$ |

Table S3: **Properties of 200-taxon ASTRAL-II simulated data from [6] and also [10].**

| ILS level (species tree height) | speciation | # of taxa | ILS | GTEE | AD |
|---|---|---|---|---|---|
| high (0.5X) | deep | 200 | $0.68 \pm 0.02$ | $0.44 \pm 0.14$ | $0.74 \pm 0.03$ |
| high (0.5X) | shallow | 200 | $0.69 \pm 0.02$ | $0.44 \pm 0.12$ | $0.74 \pm 0.03$ |
| medium (1X) | shallow | 10 | $0.17 \pm 0.06$ | $0.19 \pm 0.09$ | $0.28 \pm 0.08$ |
| medium (1X) | shallow | 50 | $0.31 \pm 0.04$ | $0.26 \pm 0.11$ | $0.42 \pm 0.08$ |
| medium (1X) | shallow | 100 | $0.33 \pm 0.02$ | $0.26 \pm 0.09$ | $0.44 \pm 0.05$ |
| medium (1X) | deep | 200 | $0.34 \pm 0.02$ | $0.34 \pm 0.12$ | $0.47 \pm 0.08$ |
| medium (1X) | shallow | 200 | $0.34 \pm 0.02$ | $0.27 \pm 0.12$ | $0.44 \pm 0.07$ |
| medium (1X) | shallow | 500 | $0.34 \pm 0.01$ | $0.28 \pm 0.11$ | $0.45 \pm 0.07$ |
| medium (1X) | shallow | 1000 | $0.35 \pm 0.01$ | $0.30 \pm 0.11$ | $0.47 \pm 0.08$ |
| low (5X) | deep | 200 | $0.09 \pm 0.01$ | $0.28 \pm 0.11$ | $0.31 \pm 0.10$ |
| low (5X) | shallow | 200 | $0.21 \pm 0.02$ | $0.21 \pm 0.13$ | $0.33 \pm 0.09$ |

#### 3.2 Software and Data availability

The scripts used to run methods and analyze the results are also available on Github: https://github.com/molloy-lab/tree-qmc-study/tree/main/han2024wtreeqmc. We analyzed a large number of analyses varying method parameters, so we encourage you to look at the scripts. We give a summary of commands below.

#### 3.3 Gene Tree Branch Support Estimation Commands

Some data sets included gene trees with branch supported estimated with the abayes technique [1], as implemented in ith IQ-TREE [5]. For the trees without abayes support (i.e., the ASTRAL-II data sets with 10, 50, 100, 500, and 1000 taxan), we estimated abayes support with 10, 50, 100, 500, and 1000 taxa using the following command:

```
iqtree2 \
    -s <input alignment> \
    -te <input gene tree> \
    -m GTR+G \
    -abayes \
    -pre <prefix for output file> \
    -T 1
```

Note that IQtree refines polytomies and gives them no support value. The default in weighted ASTRAL/ASTER [9], weighted ASTRID [4], and wTREE-QMC is to have these refined branches have no branch support when running the weighted versions of these methods. However, these (arbitrary?) refinements would be part of the input for the unweighted version of TREE-QMC (option: `--fast` or `-w f`).

#### 3.4 Species Tree Estimation Commands

**TREE-QMC.** TREE-QMC version 3.0.4 (commit b5bfe82) was run using the following command:

```
tree-qmc \
    -w <weighting scheme for quartets> \
    <branch support scheme> \
    <normalization scheme> \
    -i [input gene trees] \
    -o [output species tree] \
    &> [output log file]
```

The best version of TREE-QMC (denoted **TREE-QMC-wh**) uses the hybrid weighting scheme for quartets (option `-w h` or –hybrid) and the n2 normalization scheme for artificial taxa (option `--norm-atax 2`). If using branch support weighting, the branch support scheme set based on the method used to estimate support in the input gene trees, specifically `--lrt` for sh branch support (equivalent to `-n 0 -x 1 -d 0`), `--bootstrap` for bootstrap support (equivalent to `-n 0 -x 100 -d 0`), and `--bayes` for abayes branch support (equivalent to `-n 0.333 -x 1 -d 0.333`). We ran TREE-QMC with a variety of normalization and weighting schemes. We also ran TREE-QMC using the original algorithm method (no weighting) from [3] (option `--fast` or `-w f`) with the n2 normalization scheme for artificial taxa (option: `--norm-atax 2`); this method (denoted **TREE-QMC-n2**) does not support polytomies in the input gene trees and thus it internally refines any polytomies in the gene trees at random prior to species tree estimation. Note that we built TREE-QMC from source on the EPYC-7313 compute nodes.

**ASTRAL/ASTER.** ASTRAL/ASTER (no weighting) and ASTRAL/ASTER [9] was downloaded from https://github.com/chaoszhang/ASTER (commit 0c61214) and built from source on an EPYC-7313 compute node. For Asteroid data sets, ASTRAL/ASTER v1.16.3.4 was run using the following command:

```
astral \
    -u 0 \
    -i [input gene trees] \
    -o [output species tree] \
    &> [output log file]
```

The `-u 0` option turns of branch support calculations to enable a fair runtime comparison. For the other data sets, weighted ASTRAL/ASTER (hybrid) v1.16.3.4 was run using the following command:

```
astral-hybrid \
    --bayes \
    -u 0  \
    -i [input gene trees] \
    -o [output species tree] \
    &> [output log file]
```

The `--bayes` option was replaced with `--bootstrap` when branch support was estimated with bootstrapping. For the scalability study, we added flags `-t 1` or `-t 16` to indicate 1 or 16 threads, respectively.

**ASTRID.** We were unable to build wASTRID from source on an EPYC-7313 compute node, so we downloaded wASTRID version 0.0.7 [9] from https://github.com/RuneBlaze/internode/releases/tag/v0.0.7-snapshot. wASTRID was run with the following command:

```
wastrid \
    -b <support min>-<support max>  \
    -i [input gene trees] \
    -o [output species tree] \
    &> [output log file]
```

with minimum and maximum support set based on the input gene trees as previously noted. The default mode is support weighting (i.e., `-m support`). To use length weighting, we added `-m n-length` and removed the branch support information. To use no weighting, we added `-m internode` or `--preset vanilla`) and removed the branch support information.

**Asteroid.** Asteroid [7] was downloaded from https://github.com/BenoitMorel/Asteroid (commit f69b761) and built from source (without mpi) on an EPYC-7313 compute node. Astroid was run with the following command:

```
asteroid \
    -i [input gene trees] \
    -p [prefix for output species tree] \
    &> [output log file]
```

#### 3.5 Species Tree Branch Support Estimation Commands

Quartet score and branch support with hybrid quartet weights were computed using the following command:

```
astral-hybrid \
    --scoring -u 2 -t 16 \
    -i [input gene trees] \
    -c [input species tree] \
    -o [output species tree annotated with support] \
    &> [output log file]
```

For analyses of simulated data, we replaced astral-hybrid with astral to perform the computation with unweighted quartets. For analyses of avian biological data sets, we used astral-hybrid with the `--mode` flag to change the weighting scheme. For the plant biological data sets, we used an older version of ASTRAL (v5.7.7) with the following command:

```
java -Xmx36G -D"java.library.path=<path to ASTRAL>/lib" -jar <path to ASTRAL>/astral.5.7.7.jar \
     -t2 \
     -q [input species tree] \
     -i [input gene trees] \
     -o [output scored species tree] &> [output log file]
```

because this version handles explicitly outputs EN and gives warnings for low EN, which is useful information when gene trees have high rates of missing taxa.

#### 3.6 Scalability Study

We evaluated scalability for increasing numbers of taxa using the ASTRAL-II data sets [6] with 1000 genes, reporting the wall-clock time (i.e., the amount of time that the user waits from the beginning to the end of the computation). These computational experiments were performed on compute nodes outfitted with 32 AMD EPYC-7313 cores and 2TB of RAM. There are 28 nodes with these specifications on the CBCB cluster, we requested exclusive access of these compute nodes via slurm scheduler when performing timing experiments. We ran all methods were run with 1 thread and limited to 64 GB of memory. ASTER-h was also evaluated with 16 threads.

#### 3.7 Replicates Excluded from ASTRAL-II Data

All comparisons are made on the same set of replicate data sets. As in [6], we excluded replicate data set from analyses whenever more than half of the 1000 gene trees had the majority of their branches unresolved (due to identical sequences). Specifically, we excluded

- 1 replicate (# 41) from the 10-taxon model condition (50, 200, and 1000 genes)
- 2 replicates (# 21, 41) from the 50-taxon model condition (50, 200, and 1000 genes)
- 2 replicates (# 8, 47) from the 100-taxon model condition (50, 200, and 1000 genes)
- 3 replicates (# 8, 15, 49) from the 200 taxon model condition with very high ILS and shallow speciation (50, 200, and 1000 genes)

We also excluded replicates because the concatenation trees (downloaded from [6]) were missing on the following data sets:

- 1 replicate (# 12) from the 10-taxon, 1000-gene model condition
- 1 replicate (# 27) from the 50-taxon model condition (50, 200, and 1000 genes)

**Scalability study.** For the scalability study, 3 additional replicates were excluded because ASTER-h (single-threaded) failed to complete within 20 hours for the following data sets:

- 1 replicate (# 21) from the 200 taxon, 1000-gene model condition with very high ILS and deep speciation
- 2 replicates (# 6, 38) from the 1000-taxon, 1000-gene model condition

Lastly, the concatenation tree was also missing from the 1000-taxon, 1000-gene model condition (all 50 replicates) so we don't show results for concatenation on this model condition.

#### 3.8 Statistical Tests

We tested for significant differences between methods using two-sided, paired Wilcoxon signed-rank tests, as implemented in the R coin library. We tried two different ways of correcting for ties. The Wilcoxon method of correcting for ties was run with the following command:

```
pval <-  pvalue(wilcoxsign_test(mthd1 ~ mthd2,
                                zero.method="Wilcoxon",
                                distribution = "exact",
                                paired = TRUE,
                                alternative = "two.sided",
                                conf.int = TRUE))
```

The Pratt method of correcting for ties was run with the following command:

```
pval <- pvalue(wilcoxsign_test(mthd1 ~ mthd2,
                                zero.method="Pratt",
                                distribution = "exact",
                                paired = TRUE,
                                alternative = "two.sided",
                                conf.int = TRUE))
```

We took the **higher** of the two p-values to be conservative. We say the resulting p-value was significant after Bonferroni multiple comparisons correction if it was less than 0.05 divided by the number of tests for an experiment (we consider analyses of the Asteroid, S100, and ASTRAL-II data sets as separate experiments).

### 4 Supplemental Results

#### 4.1 Results on Asteroid data

Table S4: **Species tree (FP) error rate for Asteroid data.** Mean error rate is given across 50 replicates for each method.

| | | ASTRID | ASTER | Asteroid | TREE-QMC-n2 | n1 | n0 | n2-shared |
|---|---|---|---|---|---|---|---|---|
| Varying population size | 10 | 0.21200 | 0.17660 | 0.14430 | **0.14370** | 0.14630 | 0.17560 | 0.21490 |
| | 50000000 | 0.24600 | 0.20380 | 0.18130 | **0.18120** | 0.18190 | 0.20710 | 0.24580 |
| | 100000000 | 0.29660 | 0.26170 | 0.23150 | **0.22390** | 0.22740 | 0.25870 | 0.32100 |
| | 500000000 | 0.47530 | 0.46210 | 0.43450 | **0.41560** | 0.42080 | 0.45200 | 0.51840 |
| | 1000000000 | 0.58470 | 0.58810 | **0.55280** | 0.55330 | 0.55430 | 0.60310 | 0.66180 |
| Varying number of taxa | 25 | 0.26910 | 0.27540 | 0.22540 | 0.22280 | **0.22160** | 0.26480 | 0.27030 |
| | 75 | 0.24390 | 0.20890 | 0.18310 | **0.18070** | 0.18420 | 0.20710 | 0.26900 |
| | 50 | 0.24600 | 0.20380 | 0.18130 | **0.18120** | 0.18190 | 0.20710 | 0.24580 (dup) |
| | 100 | 0.25070 | 0.21710 | 0.19650 | **0.18560** | 0.18980 | 0.22000 | 0.28340 |
| | 125 | 0.24670 | 0.21120 | 0.18480 | **0.17650** | 0.18010 | 0.20990 | 0.27040 |
| | 150 | 0.25580 | 0.21690 | 0.18680 | **0.18160** | 0.18600 | 0.21720 | 0.28880 |
| Varying number of genes | 250 | 0.47080 | 0.43140 | **0.35980** | 0.36450 | 0.36500 | 0.42440 | 0.46060 |
| | 500 | 0.34820 | 0.31340 | 0.26010 | **0.25260** | 0.25960 | 0.29610 | 0.33560 |
| | 1000 | 0.24600 | 0.20380 | 0.18130 | **0.18120** | 0.18190 | 0.20710 | 0.24580 (dup) |
| | 2000 | 0.18170 | 0.14940 | 0.13110 | **0.12050** | 0.12630 | 0.14860 | 0.18800 |
| Varying sequence length | 50 | 0.29180 | 0.29470 | **0.24580** | 0.25020 | 0.25440 | 0.29120 | 0.34300 |
| | 100 | 0.24600 | 0.20380 | 0.18130 | **0.18120** | 0.18190 | 0.20710 | 0.24580 (dup) |
| | 200 | 0.23060 | 0.17110 | 0.16430 | **0.15320** | 0.15450 | 0.17300 | 0.20780 |
| | 500 | 0.19740 | 0.14640 | 0.13230 | 0.11520 | **0.11170** | 0.13870 | 0.15590 |
| Varying branch length scaler | 0.05 | 0.51020 | 0.48550 | 0.45060 | 0.45540 | **0.44880** | 0.48060 | 0.58390 |
| | 0.10 | 0.37830 | 0.34000 | **0.29280** | 0.30920 | 0.30210 | 0.35010 | 0.42360 |
| | 1.00 | 0.24600 | 0.20380 | 0.18130 | **0.18120** | 0.18190 | 0.20710 | 0.24580 (dup) |
| | 10.00 | 0.26130 | 0.24510 | 0.20720 | **0.19000** | 0.20270 | 0.23790 | 0.26610 |
| | 100.00 | 0.42680 | 0.42550 | **0.35790** | 0.36260 | 0.37720 | 0.42630 | 0.51750 |
| | 200.00 | 0.50080 | 0.49530 | **0.45620** | 0.48220 | 0.48530 | 0.52540 | 0.61540 |
| Varying missingness parameter | 0.50 | 0.13360 | 0.11790 | **0.09870** | 0.10280 | 0.10490 | 0.12080 | 0.14070 |
| | 0.55 | 0.17320 | 0.17190 | 0.14510 | **0.14020** | 0.14300 | 0.17010 | 0.20660 |
| | 0.60 | 0.24600 | 0.20380 | 0.18130 | **0.18120** | 0.18190 | 0.20710 | 0.24580 (dup) |
| | 0.65 | 0.38490 | 0.32350 | 0.28050 | **0.24730** | 0.25690 | 0.30250 | 0.35640 |
| | 0.70 | 0.60670 | 0.49840 | 0.43100 | **0.41460** | 0.41930 | 0.45520 | 0.53320 |
| | 0.75 | 0.82050 | 0.73420 | 0.62240 | 0.61020 | **0.60340** | 0.65720 | 0.72410 |

Table S5: **Species tree (FN) error rate for Asteroid data.** Mean error rate is given across 50 replicates for each method.

| | | ASTRID | ASTER | Asteroid | TREE-QMC-n2 | n1 | n0 | n2-shared |
|---|---|---|---|---|---|---|---|---|
| Varying population size | 10 | 0.21200 | 0.17660 | **0.14430** | 0.15750 | 0.16090 | 0.18860 | 0.22640 |
| | 50000000 | 0.24600 | 0.20380 | **0.18130** | 0.19620 | 0.19620 | 0.22040 | 0.25830 |
| | 100000000 | 0.29660 | 0.26170 | **0.23150** | 0.23960 | 0.24340 | 0.27400 | 0.33400 |
| | 500000000 | 0.47530 | 0.46210 | 0.43450 | **0.42430** | 0.42980 | 0.46080 | 0.52430 |
| | 1000000000 | 0.58470 | 0.58810 | **0.55280** | 0.56130 | 0.56170 | 0.61020 | 0.66890 |
| Varying number of taxa | 25 | 0.26910 | 0.27540 | **0.22540** | 0.24180 | 0.24000 | 0.28090 | 0.28820 |
| | 75 | 0.24390 | 0.20890 | **0.18310** | 0.19560 | 0.19970 | 0.22190 | 0.28420 |
| | 50 | 0.24600 | 0.20380 | **0.18130** | 0.19620 | 0.19620 | 0.22040 | 0.25830 (dup) |
| | 100 | 0.25070 | 0.21710 | **0.19650** | 0.20310 | 0.20640 | 0.23570 | 0.29770 |
| | 125 | 0.24670 | 0.21120 | **0.18480** | 0.19430 | 0.19900 | 0.22590 | 0.28560 |
| | 150 | 0.25580 | 0.21690 | **0.18680** | 0.19820 | 0.20250 | 0.23100 | 0.30290 |
| Varying number of genes | 250 | 0.47080 | 0.43140 | **0.35980** | 0.41230 | 0.41310 | 0.46560 | 0.50060 |
| | 500 | 0.34820 | 0.31340 | **0.26010** | 0.28310 | 0.29040 | 0.32360 | 0.36020 |
| | 1000 | 0.24600 | 0.20380 | **0.18130** | 0.19620 | 0.19620 | 0.22040 | 0.25830 (dup) |
| | 2000 | 0.18170 | 0.14940 | 0.13110 | **0.13060** | 0.13570 | 0.15830 | 0.19700 |
| Varying sequence length | 50 | 0.29180 | 0.29470 | **0.24580** | 0.26320 | 0.26790 | 0.30100 | 0.35390 |
| | 100 | 0.24600 | 0.20380 | **0.18130** | 0.19620 | 0.19620 | 0.22040 | 0.25830 (dup) |
| | 200 | 0.23060 | 0.17110 | **0.16430** | 0.17530 | 0.17570 | 0.19280 | 0.22720 |
| | 500 | 0.19740 | 0.14640 | **0.13230** | 0.13740 | 0.13400 | 0.15960 | 0.17570 |
| Varying branch length scaler | 0.05 | 0.51020 | 0.48550 | **0.45060** | 0.46850 | 0.46300 | 0.48980 | 0.59190 |
| | 0.10 | 0.37830 | 0.34000 | **0.29280** | 0.32260 | 0.31570 | 0.36080 | 0.43360 |
| | 1.00 | 0.24600 | 0.20380 | **0.18130** | 0.19620 | 0.19620 | 0.22040 | 0.25830 (dup) |
| | 10.00 | 0.26130 | 0.24510 | **0.20720** | 0.21150 | 0.22260 | 0.25570 | 0.28380 |
| | 100.00 | 0.42680 | 0.42550 | **0.35790** | 0.37230 | 0.38770 | 0.43450 | 0.52770 |
| | 200.00 | 0.50080 | 0.49530 | **0.45620** | 0.49320 | 0.49490 | 0.53280 | 0.62340 |
| Varying missingness parameter | 0.50 | 0.13360 | 0.11790 | **0.09870** | 0.10470 | 0.10680 | 0.12300 | 0.14260 |
| | 0.55 | 0.17320 | 0.17190 | **0.14510** | 0.14720 | 0.14940 | 0.17450 | 0.21280 |
| | 0.60 | 0.24600 | 0.20380 | **0.18130** | 0.19620 | 0.19620 | 0.22040 | 0.25830 (dup) |
| | 0.65 | 0.38490 | 0.32350 | **0.28050** | 0.29030 | 0.29970 | 0.33970 | 0.38820 |
| | 0.70 | 0.60670 | 0.49840 | **0.43100** | 0.48830 | 0.49130 | 0.51890 | 0.58580 |
| | 0.75 | 0.82050 | 0.73420 | **0.62240** | 0.69620 | 0.69020 | 0.72750 | 0.77880 |

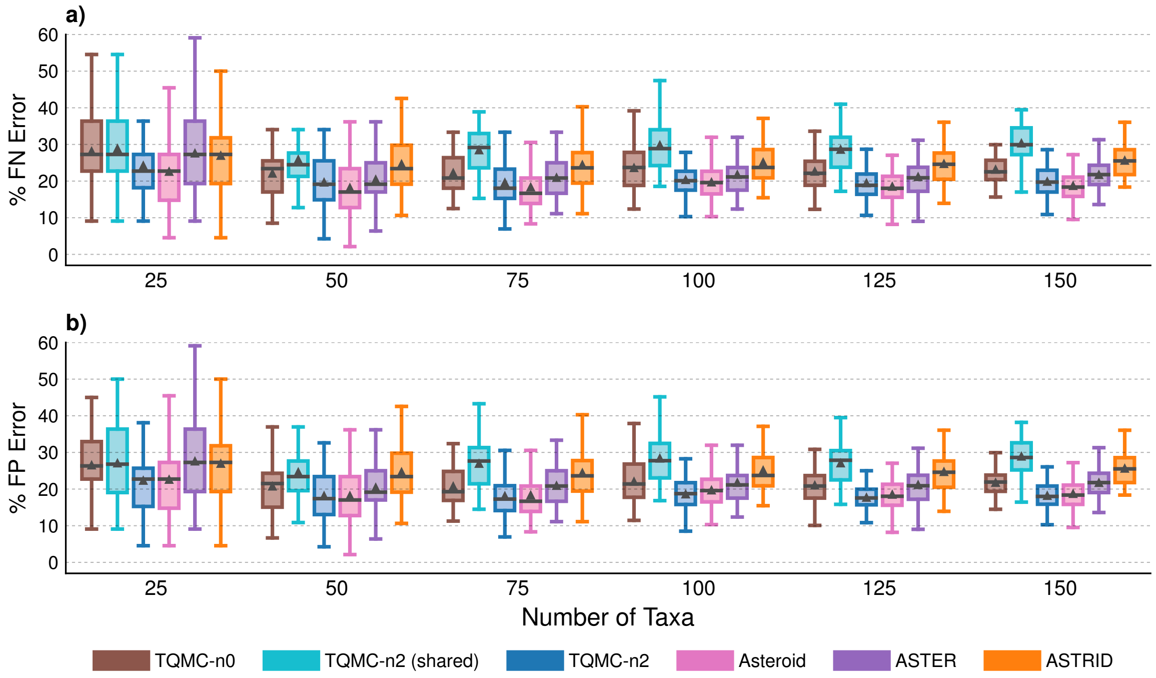

Figure S2: **Species tree error for Asteroid data with varying numbers of taxa.** Percent species tree (FN or FP) error is shown on $y$-axis for 50 replicates in each model condition. Bars represent medians, triangles represent means, outliers are not shown.

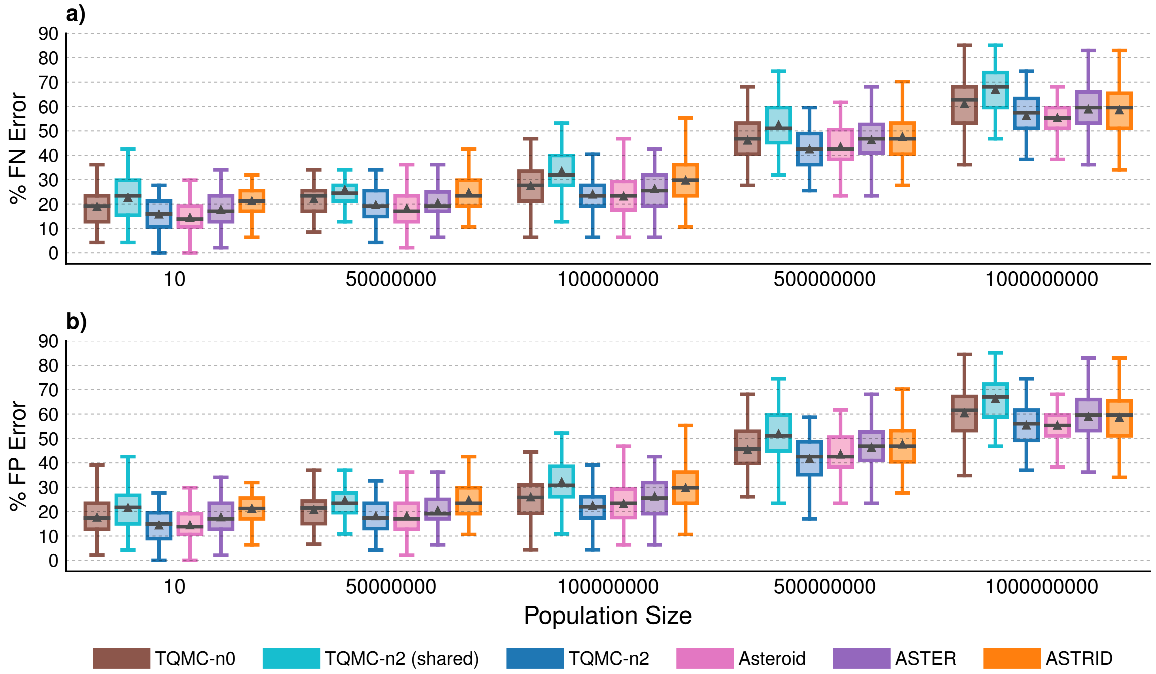

Figure S3: **Species tree error for Asteroid data with varying population size.** Percent species tree (FN or FP) error is shown on $y$-axis for 50 replicates in each model condition. Bars represent medians, triangles represent means, outliers are not shown.

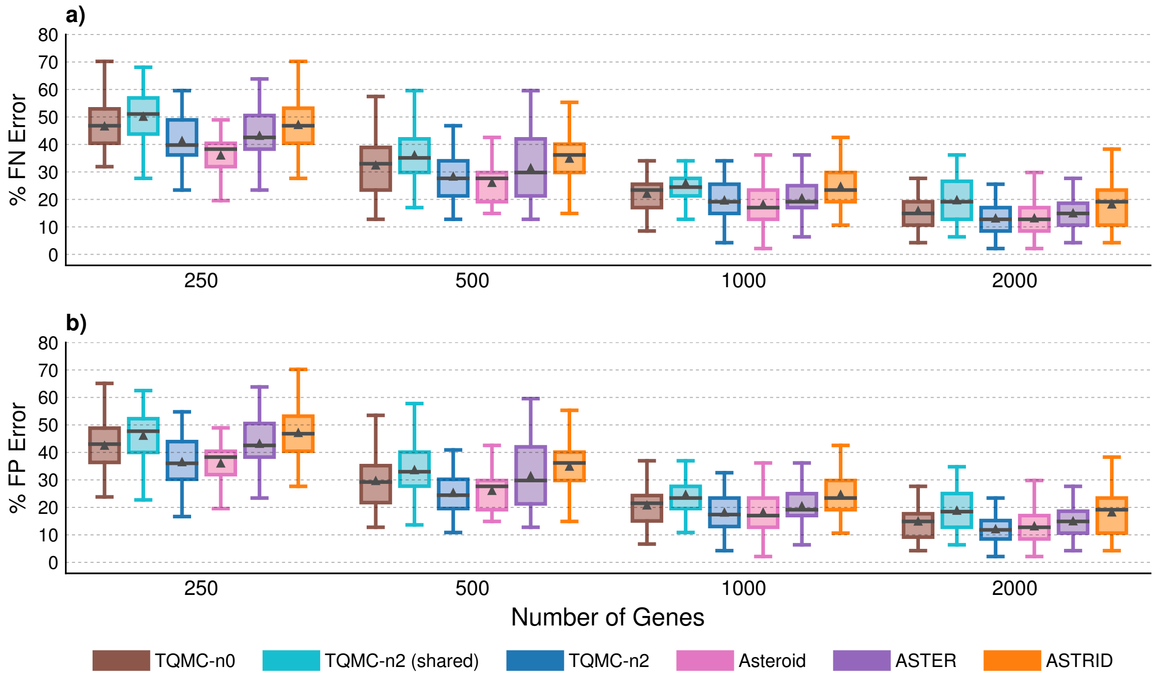

Figure S4: **Species tree error for Asteroid data with varying numbers of genes.** Percent species tree (FN or FP) error is shown on $y$-axis for 50 replicates in each model condition. Bars represent medians, triangles represent means, outliers are not shown.

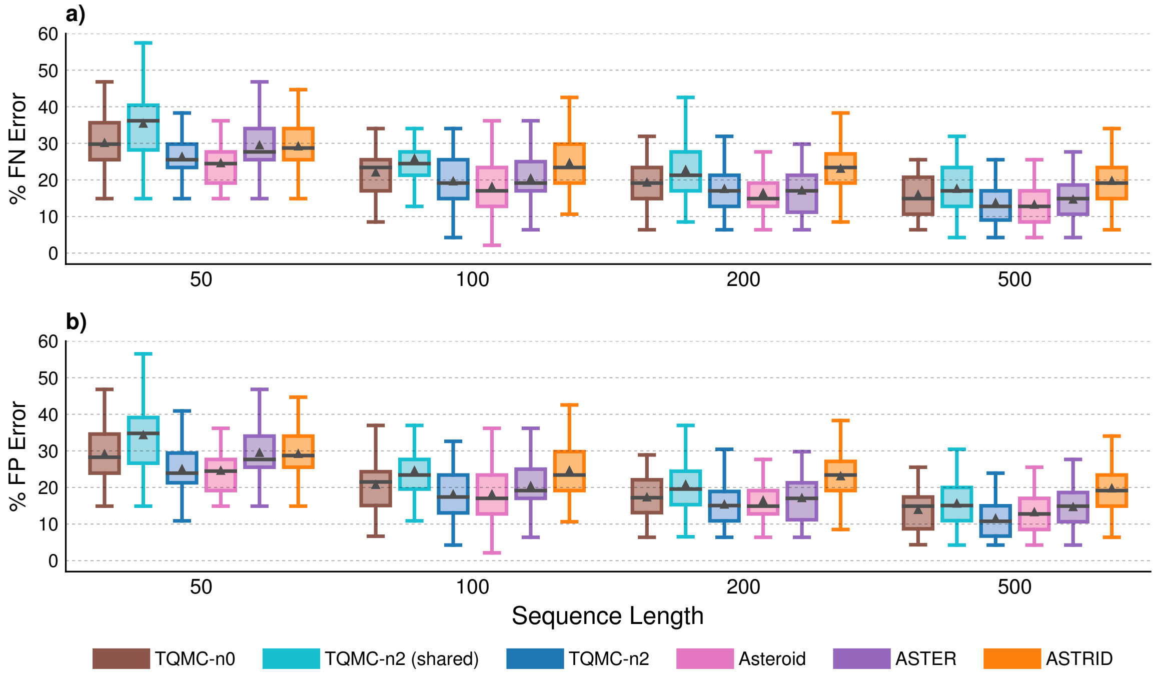

Figure S5: **Species tree error for Asteroid data with varying sequence length.** Percent species tree (FN or FP) error is shown on $y$-axis for 50 replicates in each model condition. Bars represent medians, triangles represent means, outliers are not shown.

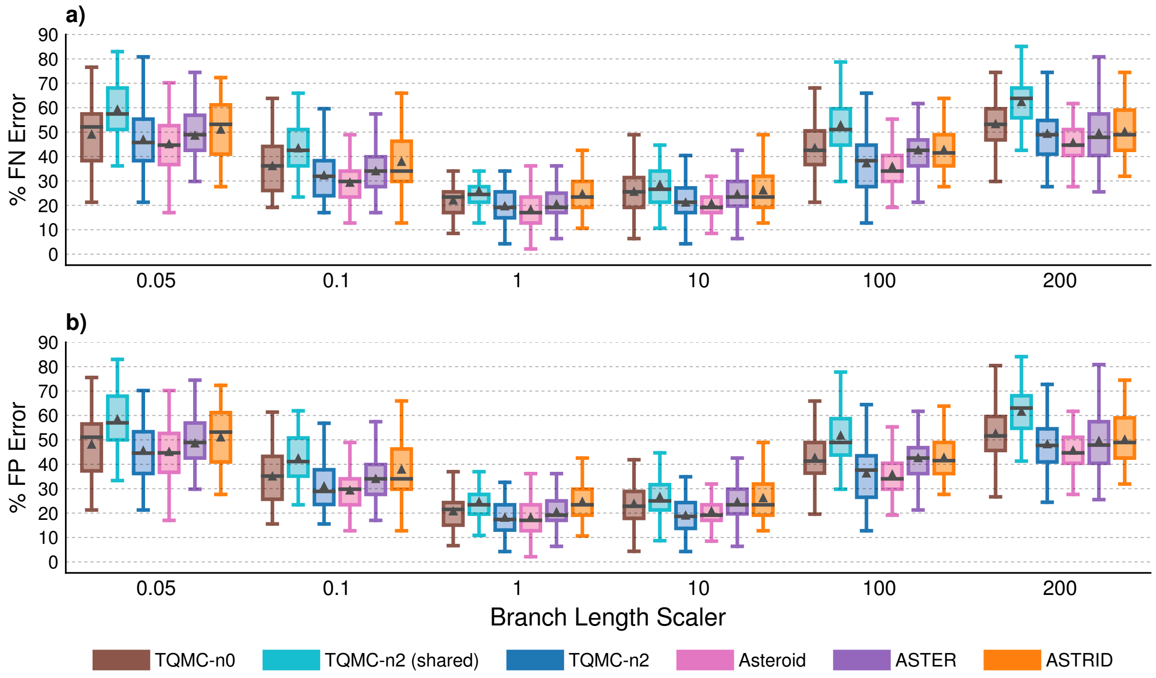

Figure S6: **Species tree error for Asteroid data with varying branch length scalar.** Percent species tree (FN or FP) error is shown on $y$-axis for 50 replicates in each model condition. Bars represent medians, triangles represent means, outliers are not shown.

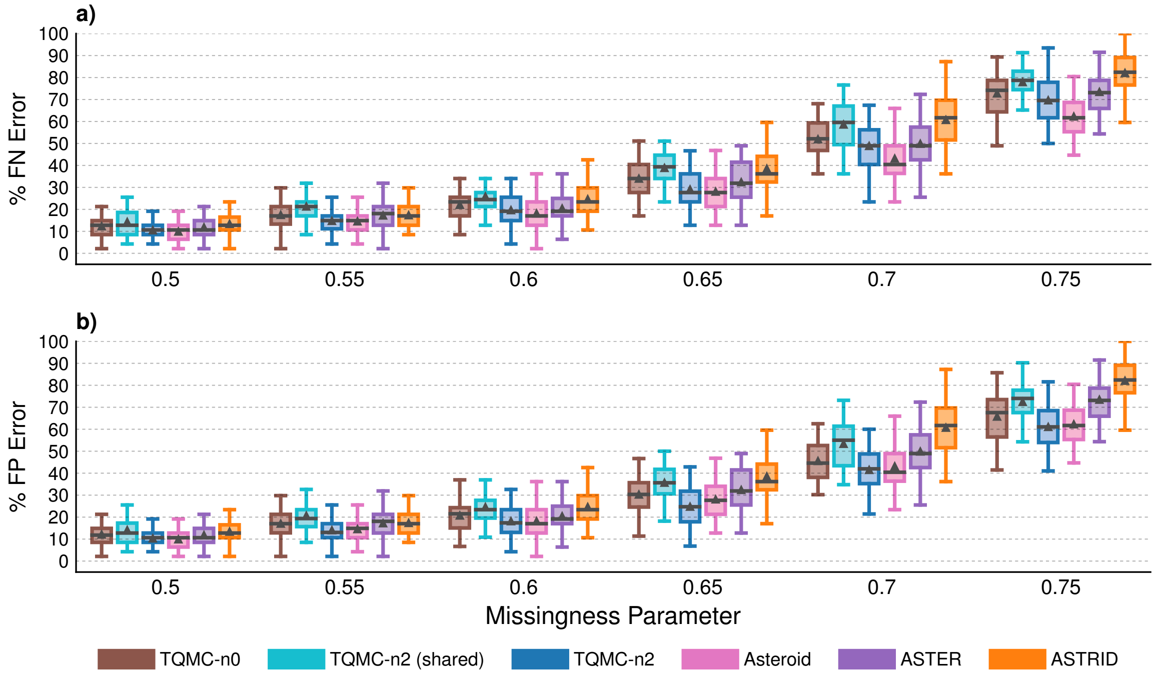

Figure S7: **Species tree error for Asteroid data with varying missingness parameter.** Percent species tree (FN or FP) error is shown on $y$-axis for 50 replicates in each model condition. Bars represent medians, triangles represent means, outliers are not shown.

Table S6: **Testing for differences between TREE-QMC-n2 vs ASTEROID on the Asteroid data.** BET is the number of replicates for which TREE-QMC-n2 has lower species tree error and thus is better than ASTEROID, and WOR is the opposite. Significance is evaluated using Wilcoxon signed-rank tests on the error rates. The symbols *, **, ***, ****, ***** indicate significance at $p < 0.5$, 0.005, and so on. MC indicates significance after Bonferroni correction, i.e., $p < 0.05 / 312 = 2e\text{-}04$ for the 312 tests made on the Asteroid data.

| TREE-QMC-n2 vs ASTEROID | | | | | | | | | |
|---|---|---|---|---|---|---|---|---|---|
| | | BET | WOR | TIE | p-val | sig | MC | note | |
| **False negative error rate** | | | | | | | | | |
| Varying population size | 10 | 15 | 24 | 11 | 0.04 | * | | ASTEROID better | |
| | 50000000 | 12 | 27 | 11 | 0.01 | * | | ASTEROID better | |
| | 100000000 | 20 | 20 | 10 | 0.4 | | | | |
| | 500000000 | 26 | 20 | 4 | 0.3 | | | | |
| | 1000000000 | 16 | 25 | 9 | 0.4 | | | | |
| Varying number of taxa | 25 | 13 | 18 | 19 | 0.3 | | | | |
| | 75 | 14 | 27 | 9 | 0.008 | * | | ASTEROID better | |
| | 50 | 12 | 27 | 11 | 0.01 | * | | ASTEROID better | (dup) |
| | 100 | 13 | 28 | 9 | 0.1 | | | | |
| | 125 | 13 | 29 | 8 | 0.003 | ** | | ASTEROID better | |
| | 150 | 15 | 32 | 3 | 0.002 | ** | | ASTEROID better | |
| Varying number of genes | 250 | 12 | 36 | 2 | 3e-06 | ***** | MC | ASTEROID better | |
| | 500 | 15 | 28 | 7 | 0.004 | ** | | ASTEROID better | |
| | 1000 | 12 | 27 | 11 | 0.01 | * | | ASTEROID better | (dup) |
| | 2000 | 13 | 15 | 22 | 0.8 | | | | |
| Varying sequence length | 50 | 13 | 28 | 9 | 0.02 | * | | ASTEROID better | |
| | 100 | 12 | 27 | 11 | 0.01 | * | | ASTEROID better | (dup) |
| | 200 | 13 | 24 | 13 | 0.03 | * | | ASTEROID better | |
| | 500 | 15 | 20 | 15 | 0.3 | | | | |
| Varying branch length scaler | 0.05 | 14 | 26 | 10 | 0.07 | | | | |
| | 0.1 | 9 | 34 | 7 | 7e-04 | ** | | ASTEROID better | |
| | 1 | 12 | 27 | 11 | 0.01 | * | | ASTEROID better | (dup) |
| | 10 | 18 | 18 | 14 | 0.7 | | | | |
| | 100 | 21 | 25 | 4 | 0.3 | | | | |
| | 200 | 12 | 29 | 9 | 0.002 | ** | | ASTEROID better | |
| Varying missingness parameter | 0.5 | 14 | 23 | 13 | 0.3 | | | | |
| | 0.55 | 15 | 17 | 18 | 0.8 | | | | |
| | 0.6 | 12 | 27 | 11 | 0.01 | * | | ASTEROID better | (dup) |
| | 0.65 | 15 | 24 | 11 | 0.08 | | | | |
| | 0.7 | 10 | 32 | 8 | 5e-06 | ***** | MC | ASTEROID better | |
| | 0.75 | 7 | 40 | 3 | 5e-09 | ***** | MC | ASTEROID better | |
| **False positive error rate** | | | | | | | | | |
| Varying population size | 10 | 20 | 23 | 7 | 1 | | | | |
| | 50000000 | 24 | 23 | 3 | 0.7 | | | | |
| | 100000000 | 30 | 19 | 1 | 0.2 | | | | |
| | 500000000 | 30 | 18 | 2 | 0.04 | * | | | |
| | 1000000000 | 22 | 24 | 4 | 0.8 | | | | |
| Varying number of taxa | 25 | 20 | 16 | 14 | 0.7 | | | | |
| | 75 | 23 | 25 | 2 | 0.8 | | | | |
| | 50 | 24 | 23 | 3 | 0.7 | | | | (dup) |
| | 100 | 29 | 20 | 1 | 0.03 | * | | | |
| | 125 | 30 | 19 | 1 | 0.01 | * | | | |
| | 150 | 31 | 19 | 0 | 0.1 | | | | |
| Varying number of genes | 250 | 28 | 22 | 0 | 0.9 | | | | |
| | 500 | 26 | 23 | 1 | 0.3 | | | | |
| | 1000 | 24 | 23 | 3 | 0.7 | | | | (dup) |
| | 2000 | 21 | 13 | 16 | 0.09 | | | | |
| Varying sequence length | 50 | 19 | 27 | 4 | 0.3 | | | | |
| | 100 | 24 | 23 | 3 | 0.7 | | | | (dup) |
| | 200 | 27 | 19 | 4 | 0.2 | | | | |
| | 500 | 32 | 11 | 7 | 8e-04 | ** | | | |
| Varying branch length scaler | 0.05 | 26 | 23 | 1 | 0.9 | | | | |
| | 0.1 | 16 | 29 | 5 | 0.04 | * | | ASTEROID better | |
| | 1 | 24 | 23 | 3 | 0.7 | | | | (dup) |
| | 10 | 31 | 16 | 3 | 0.02 | * | | | |
| | 100 | 25 | 23 | 2 | 0.7 | | | | |
| | 200 | 17 | 29 | 4 | 0.02 | * | | ASTEROID better | |
| Varying missingness parameter | 0.5 | 17 | 23 | 10 | 0.3 | | | | |
| | 0.55 | 22 | 16 | 12 | 0.3 | | | | |
| | 0.6 | 24 | 23 | 3 | 0.7 | | | | (dup) |
| | 0.65 | 39 | 11 | 0 | 1e-05 | **** | MC | | |
| | 0.7 | 33 | 17 | 0 | 0.07 | | | | |
| | 0.75 | 28 | 22 | 0 | 0.2 | | | | |

Table S7: **Testing for differences between TREE-QMC-n2 vs ASTER on the Asteroid data.** BET is the number of replicates for which TREE-QMC-n2 has lower species tree error and thus is better than ASTER, and WOR is the opposite. Significance is evaluated using Wilcoxon signed-rank tests on the error rates. The symbols *, **, ***, ****, ***** indicate significance at $p < 0.5$, 0.005, and so on. MC indicates significance after Bonferroni correction, i.e., $p < 0.05 / 312 = 2e\text{-}04$ for the 312 tests made on the Asteroid data.

| TREE-QMC-n2 vs ASTER | | | | | | | | |
|---|---|---|---|---|---|---|---|---|
| | | BET | WOR | TIE | p-val | sig | MC | note |
| **False negative error rate** | | | | | | | | |
| Varying population size | 10 | 30 | 13 | 7 | 0.005 | * | | |
| | 50000000 | 24 | 19 | 7 | 0.3 | | | |
| | 100000000 | 26 | 16 | 8 | 0.02 | * | | |
| | 500000000 | 32 | 12 | 6 | 5e-04 | *** | | |
| | 1000000000 | 32 | 16 | 2 | 0.005 | * | | |
| Varying number of taxa | 25 | 29 | 9 | 12 | 8e-04 | ** | | |
| | 75 | 26 | 18 | 6 | 0.1 | | | |
| | 50 | 24 | 19 | 7 | 0.3 | | | (dup) |
| | 100 | 30 | 15 | 5 | 0.01 | * | | |
| | 125 | 31 | 16 | 3 | 0.003 | ** | | |
| | 150 | 38 | 6 | 6 | 3e-06 | ***** | MC | |
| Varying number of genes | 250 | 28 | 16 | 6 | 0.1 | | | |
| | 500 | 30 | 14 | 6 | 0.006 | * | | |
| | 1000 | 24 | 19 | 7 | 0.3 | | | (dup) |
| | 2000 | 34 | 9 | 7 | 0.001 | ** | | |
| Varying sequence length | 50 | 29 | 11 | 10 | 0.001 | ** | | |
| | 100 | 24 | 19 | 7 | 0.3 | | | (dup) |
| | 200 | 17 | 26 | 7 | 0.4 | | | |
| | 500 | 21 | 17 | 12 | 0.3 | | | |
| Varying branch length scaler | 0.05 | 27 | 17 | 6 | 0.09 | | | |
| | 0.1 | 27 | 15 | 8 | 0.05 | | | |
| | 1 | 24 | 19 | 7 | 0.3 | | | (dup) |
| | 10 | 29 | 13 | 8 | 6e-04 | ** | | |
| | 100 | 37 | 10 | 3 | 2e-06 | ***** | MC | |
| | 200 | 25 | 21 | 4 | 0.6 | | | |
| Varying missingness parameter | 0.5 | 27 | 11 | 12 | 0.02 | * | | |
| | 0.55 | 31 | 9 | 10 | 5e-04 | *** | | |
| | 0.6 | 24 | 19 | 7 | 0.3 | | | (dup) |
| | 0.65 | 29 | 7 | 14 | 2e-04 | *** | MC | |
| | 0.7 | 22 | 20 | 8 | 0.4 | | | |
| | 0.75 | 28 | 15 | 7 | 0.01 | * | | |
| **False positive error rate** | | | | | | | | |
| Varying population size | 10 | 36 | 12 | 2 | 3e-05 | **** | MC | |
| | 50000000 | 28 | 18 | 4 | 0.009 | * | | |
| | 100000000 | 36 | 14 | 0 | 5e-05 | *** | MC | |
| | 500000000 | 40 | 10 | 0 | 1e-05 | **** | MC | |
| | 1000000000 | 32 | 16 | 2 | 2e-04 | *** | | |
| Varying number of taxa | 25 | 36 | 7 | 7 | 8e-06 | **** | MC | |
| | 75 | 33 | 16 | 1 | 3e-04 | *** | | |
| | 50 | 28 | 18 | 4 | 0.009 | * | | (dup) |
| | 100 | 39 | 10 | 1 | 4e-08 | ***** | MC | |
| | 125 | 41 | 9 | 0 | 2e-08 | ***** | MC | |
| | 150 | 46 | 4 | 0 | 4e-13 | ***** | MC | |
| Varying number of genes | 250 | 40 | 10 | 0 | 2e-08 | ***** | MC | |
| | 500 | 39 | 11 | 0 | 3e-07 | ***** | MC | |
| | 1000 | 28 | 18 | 4 | 0.009 | * | | (dup) |
| | 2000 | 39 | 8 | 3 | 2e-06 | ***** | MC | |
| Varying sequence length | 50 | 36 | 10 | 4 | 2e-05 | **** | MC | |
| | 100 | 28 | 18 | 4 | 0.009 | * | | (dup) |
| | 200 | 29 | 18 | 3 | 0.009 | * | | |
| | 500 | 33 | 14 | 3 | 2e-04 | *** | | |
| Varying branch length scaler | 0.05 | 32 | 16 | 2 | 0.002 | ** | | |
| | 0.1 | 29 | 15 | 6 | 0.002 | ** | | |
| | 1 | 28 | 18 | 4 | 0.009 | * | | (dup) |
| | 10 | 37 | 12 | 1 | 5e-07 | ***** | MC | |
| | 100 | 41 | 9 | 0 | 8e-08 | ***** | MC | |
| | 200 | 28 | 20 | 2 | 0.2 | | | |
| Varying missingness parameter | 0.5 | 28 | 11 | 11 | 0.009 | * | | |
| | 0.55 | 36 | 9 | 5 | 3e-05 | **** | MC | |
| | 0.6 | 28 | 18 | 4 | 0.009 | * | | (dup) |
| | 0.65 | 45 | 4 | 1 | 1e-12 | ***** | MC | |
| | 0.7 | 43 | 7 | 0 | 8e-09 | ***** | MC | |
| | 0.75 | 45 | 5 | 0 | 3e-10 | ***** | MC | |

Table S8: **Testing for differences between TREE-QMC-n2 vs ASTRID on the Asteroid data.** BET is the number of replicates for which TREE-QMC-n2 has lower species tree error and thus is better than ASTRID, and WOR is the opposite. Significance is evaluated using Wilcoxon signed-rank tests on the error rates. The symbols *, **, ***, ****, ***** indicate significance at $p < 0.5$, 0.005, and so on. MC indicates significance after Bonferroni correction, i.e., $p < 0.05 / 312 = 2e\text{-}04$ for the 312 tests made on the Asteroid data.

| **TREE-QMC-n2 vs ASTRID** | | | | | | | | |
|---|---|---|---|---|---|---|---|---|
| | | BET | WOR | TIE | p-val | sig | MC | note |
| **False negative error rate** | | | | | | | | |
| Varying population size | 10 | 40 | 5 | 5 | 3e-09 | ***** | MC | |
| | 50000000 | 35 | 10 | 5 | 1e-06 | ***** | MC | |
| | 100000000 | 38 | 8 | 4 | 1e-06 | ***** | MC | |
| | 500000000 | 34 | 11 | 5 | 5e-05 | **** | MC | |
| | 1000000000 | 30 | 15 | 5 | 0.05 | | | |
| Varying number of taxa | 25 | 25 | 14 | 11 | 0.04 | * | | |
| | 75 | 36 | 9 | 5 | 5e-07 | ***** | MC | |
| | 50 | 35 | 10 | 5 | 1e-06 | ***** | MC | (dup) |
| | 100 | 40 | 8 | 2 | 6e-09 | ***** | MC | |
| | 125 | 45 | 4 | 1 | 3e-11 | ***** | MC | |
| | 150 | 48 | 2 | 0 | 5e-13 | ***** | MC | |
| Varying number of genes | 250 | 38 | 9 | 3 | 1e-05 | **** | MC | |
| | 500 | 40 | 7 | 3 | 6e-08 | ***** | MC | |
| | 1000 | 35 | 10 | 5 | 1e-06 | ***** | MC | (dup) |
| | 2000 | 36 | 8 | 6 | 3e-07 | ***** | MC | |
| Varying sequence length | 50 | 30 | 17 | 3 | 0.007 | * | | |
| | 100 | 35 | 10 | 5 | 1e-06 | ***** | MC | (dup) |
| | 200 | 41 | 6 | 3 | 9e-08 | ***** | MC | |
| | 500 | 42 | 4 | 4 | 6e-10 | ***** | MC | |
| Varying branch length scaler | 0.05 | 30 | 14 | 6 | 0.005 | * | | |
| | 0.1 | 35 | 10 | 5 | 1e-05 | **** | MC | |
| | 1 | 35 | 10 | 5 | 1e-06 | ***** | MC | (dup) |
| | 10 | 34 | 13 | 3 | 3e-05 | **** | MC | |
| | 100 | 34 | 14 | 2 | 7e-04 | ** | | |
| | 200 | 26 | 23 | 1 | 0.6 | | | |
| Varying missingness parameter | 0.5 | 29 | 10 | 11 | 4e-04 | *** | | |
| | 0.55 | 32 | 15 | 3 | 0.003 | ** | | |
| | 0.6 | 35 | 10 | 5 | 1e-06 | ***** | MC | (dup) |
| | 0.65 | 35 | 9 | 6 | 4e-08 | ***** | MC | |
| | 0.7 | 43 | 3 | 4 | 2e-11 | ***** | MC | |
| | 0.75 | 46 | 1 | 3 | 7e-14 | ***** | MC | |
| **False positive error rate** | | | | | | | | |
| Varying population size | 10 | 44 | 4 | 2 | 7e-12 | ***** | MC | |
| | 50000000 | 39 | 9 | 2 | 1e-08 | ***** | MC | |
| | 100000000 | 40 | 8 | 2 | 3e-08 | ***** | MC | |
| | 500000000 | 38 | 10 | 2 | 5e-06 | **** | MC | |
| | 1000000000 | 34 | 15 | 1 | 0.006 | * | | |
| Varying number of taxa | 25 | 31 | 12 | 7 | 0.002 | ** | | |
| | 75 | 42 | 7 | 1 | 9e-10 | ***** | MC | |
| | 50 | 39 | 9 | 2 | 1e-08 | ***** | MC | (dup) |
| | 100 | 43 | 6 | 1 | 1e-11 | ***** | MC | |
| | 125 | 47 | 3 | 0 | 2e-13 | ***** | MC | |
| | 150 | 49 | 1 | 0 | 9e-15 | ***** | MC | |
| Varying number of genes | 250 | 45 | 5 | 0 | 1e-10 | ***** | MC | |
| | 500 | 43 | 7 | 0 | 1e-10 | ***** | MC | |
| | 1000 | 39 | 9 | 2 | 1e-08 | ***** | MC | (dup) |
| | 2000 | 40 | 6 | 4 | 1e-09 | ***** | MC | |
| Varying sequence length | 50 | 31 | 17 | 2 | 3e-04 | *** | | |
| | 100 | 39 | 9 | 2 | 1e-08 | ***** | MC | (dup) |
| | 200 | 45 | 5 | 0 | 1e-10 | ***** | MC | |
| | 500 | 45 | 2 | 3 | 4e-12 | ***** | MC | |
| Varying branch length scaler | 0.05 | 35 | 14 | 1 | 3e-04 | *** | | |
| | 0.1 | 38 | 10 | 2 | 2e-07 | ***** | MC | |
| | 1 | 39 | 9 | 2 | 1e-08 | ***** | MC | (dup) |
| | 10 | 39 | 10 | 1 | 1e-07 | ***** | MC | |
| | 100 | 35 | 14 | 1 | 2e-04 | *** | | |
| | 200 | 27 | 23 | 0 | 0.3 | | | |
| Varying missingness parameter | 0.5 | 31 | 10 | 9 | 2e-04 | *** | | |
| | 0.55 | 33 | 15 | 2 | 9e-04 | ** | | |
| | 0.6 | 39 | 9 | 2 | 1e-08 | ***** | MC | (dup) |
| | 0.65 | 45 | 5 | 0 | 4e-13 | ***** | MC | |
| | 0.7 | 49 | 1 | 0 | 4e-15 | ***** | MC | |
| | 0.75 | 49 | 1 | 0 | 4e-15 | ***** | MC | |

Table S9: **Testing for differences between TREE-QMC-n2 vs TREE-QMC-n2-shared on the Asteroid data.** BET is the number of replicates for which TREE-QMC-n2 has lower species tree error and thus is better than TREE-QMC-n2-shared, and WOR is the opposite. Significance is evaluated using Wilcoxon signed-rank tests on the error rates. The symbols *, **, ***, ****, ***** indicate significance at $p < 0.5$, 0.005, and so on. MC indicates significance after Bonferroni correction, i.e., $p < 0.05 / 312 =$ 2e-04 for the 312 tests made on the Asteroid data.

| **TREE-QMC-n2 vs TREE-QMC-n2-shared** | | | | | | | | |
|---|---|---|---|---|---|---|---|---|
| | | BET | WOR | TIE | p-val | sig | MC | note |
| **False negative error rate** | | | | | | | | |
| Varying population size | 10 | 41 | 5 | 4 | 1e-09 | ***** | MC | |
| | 50000000 | 37 | 7 | 6 | 5e-08 | ***** | MC | |
| | 100000000 | 45 | 1 | 4 | 2e-13 | ***** | MC | |
| | 500000000 | 44 | 3 | 3 | 2e-10 | ***** | MC | |
| | 1000000000 | 46 | 4 | 0 | 4e-11 | ***** | MC | |
| Varying number of taxa | 25 | 28 | 9 | 13 | 1e-04 | *** | MC | |
| | 75 | 46 | 2 | 2 | 1e-12 | ***** | MC | |
| | 50 | 37 | 7 | 6 | 5e-08 | ***** | MC | (dup) |
| | 100 | 50 | 0 | 0 | 2e-15 | ***** | MC | |
| | 125 | 50 | 0 | 0 | 2e-15 | ***** | MC | |
| | 150 | 50 | 0 | 0 | 2e-15 | ***** | MC | |
| Varying number of genes | 250 | 41 | 5 | 4 | 4e-10 | ***** | MC | |
| | 500 | 41 | 6 | 3 | 3e-10 | ***** | MC | |
| | 1000 | 37 | 7 | 6 | 5e-08 | ***** | MC | (dup) |
| | 2000 | 41 | 4 | 5 | 8e-11 | ***** | MC | |
| Varying sequence length | 50 | 41 | 3 | 6 | 2e-11 | ***** | MC | |
| | 100 | 37 | 7 | 6 | 5e-08 | ***** | MC | (dup) |
| | 200 | 37 | 7 | 6 | 9e-08 | ***** | MC | |
| | 500 | 40 | 4 | 6 | 4e-08 | ***** | MC | |
| Varying branch length scaler | 0.05 | 49 | 1 | 0 | 5e-15 | ***** | MC | |
| | 0.1 | 48 | 2 | 0 | 1e-13 | ***** | MC | |
| | 1 | 37 | 7 | 6 | 5e-08 | ***** | MC | (dup) |
| | 10 | 43 | 6 | 1 | 3e-10 | ***** | MC | |
| | 100 | 48 | 2 | 0 | 2e-14 | ***** | MC | |
| | 200 | 46 | 4 | 0 | 1e-11 | ***** | MC | |
| Varying missingness parameter | 0.5 | 34 | 7 | 9 | 2e-06 | ***** | MC | |
| | 0.55 | 39 | 5 | 6 | 1e-09 | ***** | MC | |
| | 0.6 | 37 | 7 | 6 | 5e-08 | ***** | MC | (dup) |
| | 0.65 | 45 | 5 | 0 | 1e-10 | ***** | MC | |
| | 0.7 | 44 | 1 | 5 | 2e-13 | ***** | MC | |
| | 0.75 | 41 | 6 | 3 | 4e-09 | ***** | MC | |
| **False positive error rate** | | | | | | | | |
| Varying population size | 10 | 42 | 5 | 3 | 7e-10 | ***** | MC | |
| | 50000000 | 39 | 7 | 4 | 1e-08 | ***** | MC | |
| | 100000000 | 46 | 1 | 3 | 4e-14 | ***** | MC | |
| | 500000000 | 45 | 4 | 1 | 2e-10 | ***** | MC | |
| | 1000000000 | 46 | 4 | 0 | 3e-11 | ***** | MC | |
| Varying number of taxa | 25 | 28 | 11 | 11 | 2e-04 | *** | | |
| | 75 | 46 | 3 | 1 | 7e-13 | ***** | MC | |
| | 50 | 39 | 7 | 4 | 1e-08 | ***** | MC | (dup) |
| | 100 | 50 | 0 | 0 | 2e-15 | ***** | MC | |
| | 125 | 50 | 0 | 0 | 2e-15 | ***** | MC | |
| | 150 | 50 | 0 | 0 | 2e-15 | ***** | MC | |
| Varying number of genes | 250 | 41 | 8 | 1 | 8e-10 | ***** | MC | |
| | 500 | 45 | 5 | 0 | 9e-11 | ***** | MC | |
| | 1000 | 39 | 7 | 4 | 1e-08 | ***** | MC | (dup) |
| | 2000 | 42 | 4 | 4 | 3e-10 | ***** | MC | |
| Varying sequence length | 50 | 42 | 6 | 2 | 1e-11 | ***** | MC | |
| | 100 | 39 | 7 | 4 | 1e-08 | ***** | MC | (dup) |
| | 200 | 38 | 7 | 5 | 1e-07 | ***** | MC | |
| | 500 | 40 | 6 | 4 | 1e-08 | ***** | MC | |
| Varying branch length scaler | 0.05 | 49 | 1 | 0 | 4e-15 | ***** | MC | |
| | 0.1 | 48 | 2 | 0 | 6e-14 | ***** | MC | |
| | 1 | 39 | 7 | 4 | 1e-08 | ***** | MC | (dup) |
| | 10 | 44 | 6 | 0 | 6e-11 | ***** | MC | |
| | 100 | 49 | 1 | 0 | 1e-14 | ***** | MC | |
| | 200 | 47 | 3 | 0 | 9e-12 | ***** | MC | |
| Varying missingness parameter | 0.5 | 34 | 7 | 9 | 3e-06 | ***** | MC | |
| | 0.55 | 38 | 6 | 6 | 2e-09 | ***** | MC | |
| | 0.6 | 39 | 7 | 4 | 1e-08 | ***** | MC | (dup) |
| | 0.65 | 46 | 4 | 0 | 1e-11 | ***** | MC | |
| | 0.7 | 46 | 3 | 1 | 5e-13 | ***** | MC | |
| | 0.75 | 44 | 6 | 0 | 2e-10 | ***** | MC | |

Table S10: **Testing for differences between TREE-QMC-n2 vs TREE-QMC-n1 on the Asteroid data.** BET is the number of replicates for which TREE-QMC-n2 has lower species tree error and thus is better than TREE-QMC-n1, and WOR is the opposite. Significance is evaluated using Wilcoxon signed-rank tests on the error rates. The symbols *, **, ***, ****, ***** indicate significance at $p < 0.5$, 0.005, and so on. MC indicates significance after Bonferroni correction, i.e., $p < 0.05 / 312 = 2e\text{-}04$ for the 312 tests made on the Asteroid data.

| **TREE-QMC-n2 vs TREE-QMC-n1** | | | | | | | | |
|---|---|---|---|---|---|---|---|---|
| | | BET | WOR | TIE | p-val | sig | MC | note |
| **False negative error rate** | | | | | | | | |
| Varying population size | 10 | 17 | 12 | 21 | 0.6 | | | |
| | 50000000 | 11 | 13 | 26 | 0.9 | | | |
| | 100000000 | 18 | 16 | 16 | 0.6 | | | |
| | 500000000 | 19 | 17 | 14 | 0.5 | | | |
| | 1000000000 | 14 | 15 | 21 | 0.8 | | | |
| Varying number of taxa | 25 | 7 | 8 | 35 | 1 | | | |
| | 75 | 22 | 15 | 13 | 0.2 | | | |
| | 50 | 11 | 13 | 26 | 0.9 | | | (dup) |
| | 100 | 21 | 17 | 12 | 0.2 | | | |
| | 125 | 26 | 16 | 8 | 0.04 | * | | |
| | 150 | 27 | 16 | 7 | 0.03 | * | | |
| Varying number of genes | 250 | 18 | 14 | 18 | 0.5 | | | |
| | 500 | 20 | 13 | 17 | 0.2 | | | |
| | 1000 | 11 | 13 | 26 | 0.9 | | | (dup) |
| | 2000 | 14 | 8 | 28 | 0.2 | | | |
| Varying sequence length | 50 | 16 | 17 | 17 | 0.5 | | | |
| | 100 | 11 | 13 | 26 | 0.9 | | | (dup) |
| | 200 | 14 | 14 | 22 | 1 | | | |
| | 500 | 6 | 13 | 31 | 0.09 | | | |
| Varying branch length scaler | 0.05 | 15 | 24 | 11 | 0.3 | | | |
| | 0.1 | 14 | 22 | 14 | 0.2 | | | |
| | 1 | 11 | 13 | 26 | 0.9 | | | (dup) |
| | 10 | 21 | 10 | 19 | 0.01 | * | | |
| | 100 | 30 | 12 | 8 | 0.002 | ** | | |
| | 200 | 22 | 19 | 9 | 0.6 | | | |
| Varying missingness parameter | 0.5 | 16 | 12 | 22 | 0.4 | | | |
| | 0.55 | 16 | 12 | 22 | 0.7 | | | |
| | 0.6 | 11 | 13 | 26 | 0.9 | | | (dup) |
| | 0.65 | 21 | 12 | 17 | 0.09 | | | |
| | 0.7 | 21 | 15 | 14 | 0.3 | | | |
| | 0.75 | 14 | 22 | 14 | 0.2 | | | |
| **False positive error rate** | | | | | | | | |
| Varying population size | 10 | 17 | 13 | 20 | 0.5 | | | |
| | 50000000 | 13 | 13 | 24 | 1 | | | |
| | 100000000 | 18 | 17 | 15 | 0.7 | | | |
| | 500000000 | 19 | 17 | 14 | 0.5 | | | |
| | 1000000000 | 17 | 18 | 15 | 0.9 | | | |
| Varying number of taxa | 25 | 7 | 8 | 35 | 1 | | | |
| | 75 | 22 | 16 | 12 | 0.3 | | | |
| | 50 | 13 | 13 | 24 | 1 | | | (dup) |
| | 100 | 21 | 17 | 12 | 0.2 | | | |
| | 125 | 27 | 16 | 7 | 0.2 | | | |
| | 150 | 28 | 16 | 6 | 0.03 | * | | |
| Varying number of genes | 250 | 16 | 19 | 15 | 1 | | | |
| | 500 | 22 | 16 | 12 | 0.2 | | | |
| | 1000 | 13 | 13 | 24 | 1 | | | (dup) |
| | 2000 | 14 | 8 | 28 | 0.2 | | | |
| Varying sequence length | 50 | 16 | 17 | 17 | 0.7 | | | |
| | 100 | 13 | 13 | 24 | 1 | | | (dup) |
| | 200 | 14 | 14 | 22 | 0.8 | | | |
| | 500 | 6 | 13 | 31 | 0.06 | | | |
| Varying branch length scaler | 0.05 | 16 | 25 | 9 | 0.4 | | | |
| | 0.1 | 14 | 23 | 13 | 0.2 | | | |
| | 1 | 13 | 13 | 24 | 1 | | | (dup) |
| | 10 | 24 | 10 | 16 | 0.02 | * | | |
| | 100 | 31 | 12 | 7 | 0.01 | * | | |
| | 200 | 22 | 19 | 9 | 0.5 | | | |
| Varying missingness parameter | 0.5 | 16 | 12 | 22 | 0.5 | | | |
| | 0.55 | 17 | 12 | 21 | 0.8 | | | |
| | 0.6 | 13 | 13 | 24 | 1 | | | (dup) |
| | 0.65 | 22 | 12 | 16 | 0.05 | | | |
| | 0.7 | 24 | 13 | 11 | 0.3 | | | |
| | 0.75 | 18 | 24 | 8 | 0.3 | | | |

Table S11: **Testing for differences between TREE-QMC-n2 vs TREE-QMC-n0 on the Asteroid data.** BET is the number of replicates for which TREE-QMC-n2 has lower species tree error and thus is better than TREE-QMC-n0, and WOR is the opposite. Significance is evaluated using Wilcoxon signed-rank tests on the error rates. The symbols *, **, ***, ****, ***** indicate significance at $p < 0.5$, 0.005, and so on. MC indicates significance after Bonferroni correction, i.e., $p < 0.05$ / 312 = 2e-04 for the 312 tests made on the Asteroid data.

| **TREE-QMC-n2 vs TREE-QMC-n0** | | | | | | | | |
|---|---|---|---|---|---|---|---|---|
| | | BET | WOR | TIE | p-val | sig | MC | note |
| **False negative error rate** | | | | | | | | |
| Varying population size | 10 | 33 | 10 | 7 | 4e-05 | **** | MC | |
| | 50000000 | 30 | 12 | 8 | 6e-04 | ** | | |
| | 100000000 | 33 | 11 | 6 | 3e-05 | **** | MC | |
| | 500000000 | 33 | 8 | 9 | 1e-04 | *** | MC | |
| | 1000000000 | 35 | 11 | 4 | 3e-06 | ***** | MC | |
| Varying number of taxa | 25 | 29 | 8 | 13 | 6e-05 | *** | MC | |
| | 75 | 30 | 11 | 9 | 3e-04 | *** | | |
| | 50 | 30 | 12 | 8 | 6e-04 | ** | | (dup) |
| | 100 | 39 | 7 | 4 | 7e-09 | ***** | MC | |
| | 125 | 39 | 6 | 5 | 1e-07 | ***** | MC | |
| | 150 | 46 | 3 | 1 | 7e-13 | ***** | MC | |
| Varying number of genes | 250 | 34 | 8 | 8 | 2e-06 | ***** | MC | |
| | 500 | 33 | 10 | 7 | 1e-05 | **** | MC | |
| | 1000 | 30 | 12 | 8 | 6e-04 | ** | | (dup) |
| | 2000 | 36 | 8 | 6 | 2e-05 | **** | MC | |
| Varying sequence length | 50 | 31 | 9 | 10 | 1e-04 | *** | MC | |
| | 100 | 30 | 12 | 8 | 6e-04 | ** | | (dup) |
| | 200 | 26 | 11 | 13 | 0.004 | ** | | |
| | 500 | 27 | 8 | 15 | 0.002 | ** | | |
| Varying branch length scaler | 0.05 | 29 | 14 | 7 | 0.01 | * | | |
| | 0.1 | 34 | 10 | 6 | 7e-05 | *** | MC | |
| | 1 | 30 | 12 | 8 | 6e-04 | ** | | (dup) |
| | 10 | 34 | 12 | 4 | 1e-05 | **** | MC | |
| | 100 | 37 | 7 | 6 | 3e-08 | ***** | MC | |
| | 200 | 31 | 13 | 6 | 6e-04 | ** | | |
| Varying missingness parameter | 0.5 | 33 | 8 | 9 | 0.001 | ** | | |
| | 0.55 | 31 | 6 | 13 | 3e-05 | **** | MC | |
| | 0.6 | 30 | 12 | 8 | 6e-04 | ** | | (dup) |
| | 0.65 | 38 | 8 | 4 | 3e-07 | ***** | MC | |
| | 0.7 | 31 | 11 | 8 | 0.003 | ** | | |
| | 0.75 | 30 | 15 | 5 | 0.01 | * | | |
| **False positive error rate** | | | | | | | | |
| Varying population size | 10 | 35 | 9 | 6 | 3e-05 | **** | MC | |
| | 50000000 | 32 | 13 | 5 | 5e-04 | ** | | |
| | 100000000 | 35 | 11 | 4 | 1e-05 | **** | MC | |
| | 500000000 | 34 | 11 | 5 | 1e-04 | *** | MC | |
| | 1000000000 | 36 | 14 | 0 | 2e-06 | ***** | MC | |
| Varying number of taxa | 25 | 30 | 10 | 10 | 3e-04 | *** | | |
| | 75 | 30 | 13 | 7 | 4e-04 | *** | | |
| | 50 | 32 | 13 | 5 | 5e-04 | ** | | (dup) |
| | 100 | 40 | 8 | 2 | 3e-09 | ***** | MC | |
| | 125 | 42 | 5 | 3 | 5e-08 | ***** | MC | |
| | 150 | 47 | 3 | 0 | 8e-14 | ***** | MC | |
| Varying number of genes | 250 | 35 | 11 | 4 | 3e-06 | ***** | MC | |
| | 500 | 36 | 9 | 5 | 4e-06 | ***** | MC | |
| | 1000 | 32 | 13 | 5 | 5e-04 | ** | | (dup) |
| | 2000 | 36 | 8 | 6 | 1e-05 | **** | MC | |
| Varying sequence length | 50 | 32 | 9 | 9 | 5e-05 | **** | MC | |
| | 100 | 32 | 13 | 5 | 5e-04 | ** | | (dup) |
| | 200 | 28 | 13 | 9 | 0.004 | ** | | |
| | 500 | 28 | 9 | 13 | 4e-04 | *** | | |
| Varying branch length scaler | 0.05 | 33 | 15 | 2 | 0.004 | ** | | |
| | 0.1 | 35 | 12 | 3 | 3e-05 | **** | MC | |
| | 1 | 32 | 13 | 5 | 5e-04 | ** | | (dup) |
| | 10 | 37 | 12 | 1 | 5e-06 | ***** | MC | |
| | 100 | 40 | 8 | 2 | 3e-08 | ***** | MC | |
| | 200 | 35 | 13 | 2 | 3e-04 | *** | | |
| Varying missingness parameter | 0.5 | 33 | 9 | 8 | 0.002 | ** | | |
| | 0.55 | 35 | 6 | 9 | 1e-05 | **** | MC | |
| | 0.6 | 32 | 13 | 5 | 5e-04 | ** | | (dup) |
| | 0.65 | 40 | 7 | 3 | 2e-07 | ***** | MC | |
| | 0.7 | 32 | 13 | 5 | 9e-04 | ** | | |
| | 0.75 | 36 | 13 | 1 | 5e-04 | ** | | |

##### 4.2 Results on S100 data

Table S12: **Species tree (RF) error rate for S100 simulated data.** Mean error rate is given across 50 replicates for each method. Note: TREE-QMC-n2 refines polytomies in the input gene trees randomly (note that it is given trees with bootstrap support because IQTree refines polytomies when computing abayes support)

| # of genes | sequence length | ASTRID ws | ASTER wh | TREE-QMC wh-n2 | TREE-QMC wh-n1 | TREE-QMC wh-n0 | TREE-QMC ws-n2 | TREE-QMC n2 |
|---|---|---|---|---|---|---|---|---|
| **bootstrap support for weighted methods** | | | | | | | | |
| 50 | 200 | 0.16310 | 0.15530 | **0.15390** | 0.15630 | 0.16740 | 0.15610 | 0.17590 |
| 50 | 400 | 0.13120 | 0.12550 | **0.12220** | 0.12350 | 0.13200 | 0.12350 | 0.13100 |
| 50 | 800 | 0.11350 | 0.10390 | **0.10310** | 0.10430 | 0.11040 | 0.10530 | 0.11450 |
| 50 | 1600 | 0.10350 | 0.09590 | **0.09290** | 0.09530 | 0.09920 | 0.09690 | 0.09710 |
| 200 | 200 | 0.09710 | 0.09630 | **0.09490** | 0.09880 | 0.10020 | 0.09820 | 0.10960 |
| 200 | 400 | 0.07960 | 0.07670 | **0.07290** | 0.07690 | 0.08330 | 0.07470 | 0.08550 |
| 200 | 800 | 0.06900 | 0.06470 | **0.06290** | 0.06430 | 0.06800 | 0.06410 | 0.06980 |
| 200 | 1600 | 0.06140 | 0.05820 | **0.05550** | 0.05880 | 0.06160 | 0.05610 | 0.06410 |
| 500 | 200 | 0.07740 | **0.07530** | 0.07550 | 0.07920 | 0.08020 | 0.07940 | 0.09260 |
| 500 | 400 | 0.06160 | 0.05960 | **0.05550** | 0.05960 | 0.06450 | 0.06000 | 0.07220 |
| 500 | 800 | 0.05080 | 0.04900 | **0.04800** | 0.04960 | 0.05350 | 0.04920 | 0.05470 |
| 500 | 1600 | 0.04410 | 0.04290 | **0.04060** | 0.04260 | 0.04430 | 0.04240 | 0.04880 |
| 1000 | 200 | 0.06690 | **0.06530** | 0.06730 | 0.07160 | 0.07020 | 0.07040 | 0.08000 |
| 1000 | 400 | 0.05160 | 0.05140 | **0.04800** | 0.05020 | 0.05350 | 0.05060 | 0.06330 |
| 1000 | 800 | 0.04200 | 0.04180 | **0.03900** | 0.04260 | 0.04490 | 0.04120 | 0.05040 |
| 1000 | 1600 | 0.03590 | 0.03610 | **0.03510** | 0.03630 | 0.03820 | 0.03530 | 0.04100 |
| **abayes support for weighted methods** | | | | | | | | |
| 50 | 200 | 0.15040 | 0.14570 | **0.14140** | 0.14570 | 0.15860 | 0.14550 | 0.17590 |
| 50 | 400 | 0.12390 | 0.12180 | **0.11120** | 0.11690 | 0.12330 | 0.11530 | 0.13100 |
| 50 | 800 | 0.11140 | 0.10840 | **0.09670** | 0.10370 | 0.11260 | 0.10060 | 0.11450 |
| 50 | 1600 | 0.10450 | 0.09550 | **0.08630** | 0.09310 | 0.09940 | 0.08820 | 0.09710 |
| 200 | 200 | 0.09470 | 0.09220 | **0.08310** | 0.08710 | 0.09800 | 0.09430 | 0.10960 |
| 200 | 400 | 0.07430 | 0.07220 | **0.06710** | 0.07120 | 0.07780 | 0.07180 | 0.08550 |
| 200 | 800 | 0.06780 | 0.06410 | **0.06180** | 0.06410 | 0.06760 | 0.06240 | 0.06980 |
| 200 | 1600 | 0.05880 | 0.05650 | **0.05160** | 0.05490 | 0.05800 | 0.05370 | 0.06410 |
| 500 | 200 | 0.07530 | 0.07160 | **0.06920** | 0.07290 | 0.07780 | 0.07840 | 0.09260 |
| 500 | 400 | 0.06080 | **0.05760** | 0.05860 | 0.05840 | 0.06080 | 0.06160 | 0.07220 |
| 500 | 800 | 0.04880 | 0.04690 | **0.04370** | 0.04590 | 0.04960 | 0.04820 | 0.05470 |
| 500 | 1600 | 0.04080 | 0.03920 | **0.03530** | 0.03860 | 0.04220 | 0.03820 | 0.04880 |
| 1000 | 200 | 0.06220 | 0.06310 | **0.05350** | 0.06100 | 0.06840 | 0.05920 | 0.08000 |
| 1000 | 400 | 0.05120 | 0.05160 | **0.04650** | 0.05160 | 0.05610 | 0.05240 | 0.06330 |
| 1000 | 800 | 0.04220 | 0.04180 | **0.03630** | 0.04060 | 0.04650 | 0.04140 | 0.05040 |
| 1000 | 1600 | 0.03740 | 0.03530 | **0.02880** | 0.03490 | 0.04000 | 0.03410 | 0.04100 |

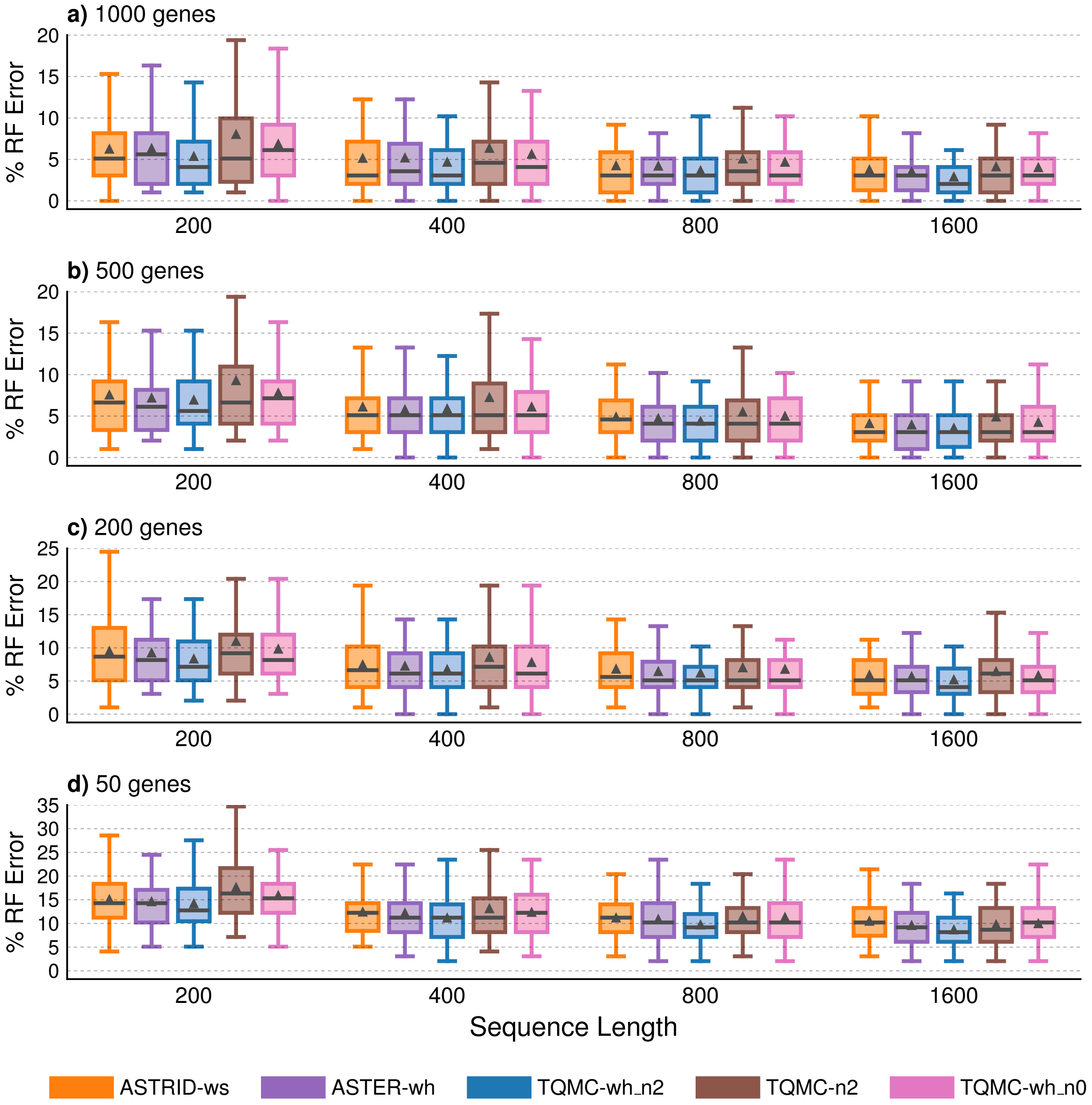

Figure S8: **Species tree error for S100 data with abayes support.** Percent species tree (RF) error is shown on $y$-axis for 50 replicates in each model condition. Bars represent medians, triangles represent means, outliers are not shown.

Table S13: **Testing for differences between TREE-QMC-wh_n2 vs ASTER-wh on the S100 simulated data.** BET is the number of replicates for which TREE-QMC-wh_n2 has lower species tree (RF) error and thus is better than ASTER-wh , WOR is the number of replicates for which wTREE-QMC-wh_n2 has higher RF error and thus is worse than ASTER-wh , and TIE is the number of replicates where the two methods tie. Significance is evaluated using paired, two-sided Wilcoxon signed-rank tests on the RF error rates. The symbols *, **, ***, ****, ***** indicate significance at $p < 0.5$, 0.005, and so on. MC indicates significance after Bonferroni correction, i.e., $p < 0.05 / 96 = 5e\text{-}04$ for the 96 tests made on the S100 data.

| **TREE-QMC-wh_n2 vs ASTER-wh** | | | | | | | | |
|---|---|---|---|---|---|---|---|---|
| # of genes | sequence length | BET | WOR | TIE | p-val | sig | MC | note |
| **Abayes Support for weighted methods** | | | | | | | | |
| 50 | 200 | 23 | 17 | 10 | 0.3 | | | |
| 50 | 400 | 33 | 7 | 10 | 4e-04 | *** | MC | |
| 50 | 800 | 27 | 8 | 15 | 3e-04 | *** | MC | |
| 50 | 1600 | 23 | 6 | 21 | 4e-04 | *** | MC | |
| 200 | 200 | 30 | 9 | 11 | 0.007 | * | | |
| 200 | 400 | 21 | 14 | 15 | 0.2 | | | |
| 200 | 800 | 18 | 11 | 21 | 0.2 | | | |
| 200 | 1600 | 22 | 10 | 18 | 0.02 | * | | |
| 500 | 200 | 22 | 15 | 13 | 0.8 | | | |
| 500 | 400 | 14 | 22 | 14 | 0.5 | | | |
| 500 | 800 | 18 | 11 | 21 | 0.5 | | | |
| 500 | 1600 | 21 | 13 | 16 | 0.2 | | | |
| 1000 | 200 | 20 | 9 | 21 | 0.006 | * | | |
| 1000 | 400 | 20 | 6 | 24 | 0.003 | ** | | |
| 1000 | 800 | 20 | 7 | 23 | 0.008 | * | | |
| 1000 | 1600 | 19 | 5 | 26 | 0.002 | ** | | |
| **Boostrap Support for weighted methods** | | | | | | | | |
| 50 | 200 | 22 | 17 | 11 | 0.4 | | | |
| 50 | 400 | 19 | 14 | 17 | 0.5 | | | |
| 50 | 800 | 23 | 18 | 9 | 0.8 | | | |
| 50 | 1600 | 21 | 16 | 13 | 0.2 | | | |
| 200 | 200 | 19 | 17 | 14 | 0.6 | | | |
| 200 | 400 | 21 | 11 | 18 | 0.06 | | | |
| 200 | 800 | 17 | 18 | 15 | 0.6 | | | |
| 200 | 1600 | 17 | 11 | 22 | 0.2 | | | |
| 500 | 200 | 23 | 18 | 9 | 0.8 | | | |
| 500 | 400 | 21 | 14 | 15 | 0.1 | | | |
| 500 | 800 | 17 | 10 | 23 | 0.3 | | | |
| 500 | 1600 | 13 | 8 | 29 | 0.2 | | | |
| 1000 | 200 | 16 | 19 | 15 | 0.6 | | | |
| 1000 | 400 | 16 | 10 | 24 | 0.1 | | | |
| 1000 | 800 | 17 | 6 | 27 | 0.09 | | | |
| 1000 | 1600 | 13 | 10 | 27 | 0.4 | | | |

Table S14: **Testing for differences between TREE-QMC-wh_n2 vs ASTRID-ws on the S100 simulated data.** BET is the number of replicates for which TREE-QMC-wh_n2 has lower species tree (RF) error and thus is better than ASTRID-ws , WOR is the number of replicates for which wTREE-QMC-wh_n2 has higher RF error and thus is worse than ASTRID-ws , and TIE is the number of replicates where the two methods tie. Significance is evaluated using paired, two-sided Wilcoxon signed-rank tests on the RF error rates. The symbols *, **, ***, ****, ***** indicate significance at $p < 0.5$, 0.005, and so on. MC indicates significance after Bonferroni correction, i.e., $p < 0.05 / 96 = 5e\text{-}04$ for the 96 tests made on the S100 data.

| **TREE-QMC-wh_n2 vs ASTRID-ws** | | | | | | | | |
|---|---|---|---|---|---|---|---|---|
| # of genes | sequence length | BET | WOR | TIE | p-val | sig | MC | note |
| **Abayes Support for weighted methods** | | | | | | | | |
| 50 | 200 | 30 | 16 | 4 | 0.03 | * | | |
| 50 | 400 | 33 | 12 | 5 | 0.001 | ** | | |
| 50 | 800 | 35 | 10 | 5 | 2e-04 | *** | MC | |
| 50 | 1600 | 34 | 10 | 6 | 2e-05 | **** | MC | |
| 200 | 200 | 27 | 7 | 16 | 3e-04 | *** | MC | |
| 200 | 400 | 25 | 13 | 12 | 0.03 | * | | |
| 200 | 800 | 25 | 13 | 12 | 0.03 | * | | |
| 200 | 1600 | 29 | 7 | 14 | 0.006 | * | | |
| 500 | 200 | 24 | 16 | 10 | 0.1 | | | |
| 500 | 400 | 23 | 13 | 14 | 0.1 | | | |
| 500 | 800 | 22 | 11 | 17 | 0.02 | * | | |
| 500 | 1600 | 24 | 9 | 17 | 0.04 | * | | |
| 1000 | 200 | 25 | 9 | 16 | 0.008 | * | | |
| 1000 | 400 | 21 | 13 | 16 | 0.07 | | | |
| 1000 | 800 | 21 | 8 | 21 | 0.004 | ** | | |
| 1000 | 1600 | 23 | 5 | 22 | 4e-04 | *** | MC | |
| **Boostrap Support for weighted methods** | | | | | | | | |
| 50 | 200 | 30 | 12 | 8 | 0.004 | ** | | |
| 50 | 400 | 31 | 12 | 7 | 0.002 | ** | | |
| 50 | 800 | 31 | 13 | 6 | 0.002 | ** | | |
| 50 | 1600 | 32 | 9 | 9 | 2e-04 | *** | MC | |
| 200 | 200 | 20 | 18 | 12 | 0.6 | | | |
| 200 | 400 | 22 | 14 | 14 | 0.03 | * | | |
| 200 | 800 | 23 | 11 | 16 | 0.02 | * | | |
| 200 | 1600 | 19 | 11 | 20 | 0.04 | * | | |
| 500 | 200 | 17 | 14 | 19 | 0.6 | | | |
| 500 | 400 | 25 | 9 | 16 | 0.002 | ** | | |
| 500 | 800 | 17 | 12 | 21 | 0.2 | | | |
| 500 | 1600 | 20 | 9 | 21 | 0.03 | * | | |
| 1000 | 200 | 23 | 18 | 9 | 1 | | | |
| 1000 | 400 | 21 | 11 | 18 | 0.09 | | | |
| 1000 | 800 | 15 | 9 | 26 | 0.1 | | | |
| 1000 | 1600 | 15 | 12 | 23 | 0.4 | | | |

Table S15: **Testing for differences between TREE-QMC-wh_n2 vs TREE-QMC-n2 on the S100 simulated data.** BET is the number of replicates for which TREE-QMC-wh_n2 has lower species tree (RF) error and thus is better than TREE-QMC-n2 , WOR is the number of replicates for which wTREE-QMC-wh_n2 has higher RF error and thus is worse than TREE-QMC-n2 , and TIE is the number of replicates where the two methods tie. Significance is evaluated using paired, two-sided Wilcoxon signed-rank tests on the RF error rates. The symbols *, **, ***, ****, ***** indicate significance at $p < 0.5$, 0.005, and so on. MC indicates significance after Bonferroni correction, i.e., $p < 0.05$ / 96 = 5e-04 for the 96 tests made on the S100 data. Note: TREE-QMC-n2 refines polytomies in the input gene trees randomly (note that it is given trees with bootstrap support because IQTree refines polytomies when computing abayes support)

| **TREE-QMC-wh_n2 vs TREE-QMC-n2** | | | | | | | | |
|---|---|---|---|---|---|---|---|---|
| # of genes | sequence length | BET | WOR | TIE | p-val | sig | MC | note |
| **Abayes Support for weighted methods** | | | | | | | | |
| 50 | 200 | 38 | 8 | 4 | 3e-08 | ***** | MC | |
| 50 | 400 | 31 | 11 | 8 | 9e-05 | *** | MC | |
| 50 | 800 | 31 | 8 | 11 | 9e-05 | *** | MC | |
| 50 | 1600 | 24 | 16 | 10 | 0.05 | * | | |
| 200 | 200 | 33 | 11 | 6 | 1e-05 | **** | MC | |
| 200 | 400 | 34 | 8 | 8 | 3e-06 | ***** | MC | |
| 200 | 800 | 22 | 12 | 16 | 0.04 | * | | |
| 200 | 1600 | 29 | 9 | 12 | 9e-04 | ** | | |
| 500 | 200 | 28 | 10 | 12 | 0.001 | ** | | |
| 500 | 400 | 28 | 8 | 14 | 9e-05 | *** | MC | |
| 500 | 800 | 22 | 9 | 19 | 0.008 | * | | |
| 500 | 1600 | 23 | 6 | 21 | 4e-04 | *** | MC | |
| 1000 | 200 | 27 | 7 | 16 | 4e-05 | **** | MC | |
| 1000 | 400 | 28 | 4 | 18 | 2e-06 | ***** | MC | |
| 1000 | 800 | 28 | 5 | 17 | 9e-06 | **** | MC | |
| 1000 | 1600 | 22 | 5 | 23 | 0.002 | ** | | |
| **Boostrap Support for weighted methods** | | | | | | | | |
| 50 | 200 | 29 | 13 | 8 | 3e-04 | *** | MC | |
| 50 | 400 | 25 | 20 | 5 | 0.4 | | | |
| 50 | 800 | 26 | 15 | 9 | 0.04 | * | | |
| 50 | 1600 | 22 | 18 | 10 | 0.3 | | | |
| 200 | 200 | 24 | 16 | 10 | 0.05 | * | | |
| 200 | 400 | 29 | 12 | 9 | 0.005 | * | | |
| 200 | 800 | 17 | 14 | 19 | 0.3 | | | |
| 200 | 1600 | 24 | 9 | 17 | 0.003 | ** | | |
| 500 | 200 | 24 | 19 | 7 | 0.05 | * | | |
| 500 | 400 | 26 | 9 | 15 | 2e-04 | *** | MC | |
| 500 | 800 | 18 | 16 | 16 | 0.3 | | | |
| 500 | 1600 | 17 | 12 | 21 | 0.2 | | | |
| 1000 | 200 | 26 | 17 | 7 | 0.04 | * | | |
| 1000 | 400 | 27 | 10 | 13 | 0.001 | ** | | |
| 1000 | 800 | 21 | 9 | 20 | 0.005 | ** | | |
| 1000 | 1600 | 12 | 11 | 27 | 0.6 | | | |

#### 4.3 Results on ASTRAL-II data

Table S16: **Runtime for ASTRAL-II simulated data.** Mean runtime (in seconds) is given across replicates for each method. All weighted methods were given gene trees with abayes support. CA-ML trees from original study were not available for 1000-taxon, 1000-gene data sets.

| # of taxa | ILS level | speciation location | # of genes | ASTRID ws | TREE-QMC (n2) | TREE-QMC wh (n2) | ASTER wh (16 threads) | ASTER wh (1 thread) |
|---|---|---|---|---|---|---|---|---|
| 10 | medium | shallow | 1000 | 0.0 | 0.2 | 0.3 | 0.1 | 0.7 |
| 50 | medium | shallow | 1000 | 0.1 | 3.3 | 7.5 | 2.4 | 34.5 |
| 100 | medium | shallow | 1000 | 0.1 | 13.8 | 40.5 | 10.7 | 131.6 |
| 200 | low | deep | 1000 | 0.3 | 51.8 | 170.8 | 41.5 | 700.0 |
| 200 | low | shallow | 1000 | 0.3 | 50.4 | 159.9 | 36.9 | 645.5 |
| 200 | medium | deep | 1000 | 0.3 | 55.7 | 181.3 | 44.0 | 734.3 |
| 200 | medium | shallow | 1000 | 0.3 | 51.7 | 167.1 | 41.1 | 661.1 |
| 200 | high | deep | 1000 | 0.4 | 61.1 | 201.6 | 54.0 | 955.0 |
| 200 | high | shallow | 1000 | 0.4 | 58.8 | 192.7 | 53.8 | 869.3 |
| 500 | medium | shallow | 1000 | 1.9 | 322.8 | 1084.0 | 273.1 | 5145.7 |
| 1000 | medium | shallow | 1000 | 7.3 | 1303.3 | 4469.8 | 1272.5 | 28758.8 |

Table S17: **Species tree (RF) error rate for ASTRAL-II simulated data.** Mean error rate is given across replicates for each method. All weighted methods were given gene trees with abayes support. CA-ML trees from original study were not available for 1000-taxon, 1000-gene data sets.

| # of taxa | ILS level | speciation location | # of genes | CA-ML | ASTER wh | TREE-QMC wh (n2) | TREE-QMC (n2) | ASTRID ws |
|---|---|---|---|---|---|---|---|---|
| 10 | medium | shallow | 50 | 0.03830 | 0.03060 | 0.03060 | 0.03570 | **0.02810** |
| 10 | medium | shallow | 200 | 0.01790 | **0.00760** | 0.01020 | 0.01790 | **0.00760** |
| 10 | medium | shallow | 1000 | 0.02080 | **0.01560** | **0.01560** | **0.01560** | **0.01560** |
| 50 | medium | shallow | 50 | 0.07800 | 0.06070 | 0.05850 | 0.07090 | **0.05760** |
| 50 | medium | shallow | 200 | 0.04520 | 0.03060 | **0.02840** | 0.04120 | 0.03550 |
| 50 | medium | shallow | 1000 | 0.02660 | 0.01860 | **0.01770** | 0.02530 | 0.01910 |
| 100 | medium | shallow | 50 | 0.09120 | 0.07160 | **0.06720** | 0.07230 | 0.07040 |
| 100 | medium | shallow | 200 | 0.04740 | 0.03830 | **0.03610** | 0.04680 | 0.03930 |
| 100 | medium | shallow | 1000 | 0.02490 | **0.01910** | 0.01980 | 0.03040 | 0.02400 |
| 200 | low | deep | 50 | **0.03980** | 0.05830 | 0.05160 | 0.06710 | 0.06490 |
| 200 | low | deep | 200 | **0.02230** | 0.03520 | 0.03200 | 0.05060 | 0.05130 |
| 200 | low | deep | 1000 | **0.01780** | 0.03010 | 0.02480 | 0.04520 | 0.04850 |
| 200 | low | shallow | 50 | 0.05360 | 0.04810 | **0.04240** | 0.05280 | 0.04430 |
| 200 | low | shallow | 200 | 0.03110 | 0.02350 | 0.02240 | 0.02910 | **0.02230** |
| 200 | low | shallow | 1000 | 0.01440 | 0.01260 | **0.01120** | 0.01880 | 0.01300 |
| 200 | medium | deep | 50 | 0.10300 | 0.08920 | **0.08570** | 0.09750 | 0.09100 |
| 200 | medium | deep | 200 | 0.05690 | 0.05230 | **0.04660** | 0.05700 | 0.05500 |
| 200 | medium | deep | 1000 | **0.02840** | 0.03520 | 0.03090 | 0.03980 | 0.03920 |
| 200 | medium | shallow | 50 | 0.09220 | 0.07060 | **0.06690** | 0.08090 | 0.07280 |
| 200 | medium | shallow | 200 | 0.05520 | 0.04030 | **0.03820** | 0.04750 | 0.04170 |
| 200 | medium | shallow | 1000 | 0.02780 | 0.02410 | **0.02270** | 0.03170 | 0.02600 |
| 200 | high | deep | 50 | 0.28160 | 0.18980 | **0.18580** | 0.20670 | 0.20630 |
| 200 | high | deep | 200 | 0.16100 | **0.09130** | 0.09190 | 0.10940 | 0.10560 |
| 200 | high | deep | 1000 | 0.08010 | **0.04710** | 0.04860 | 0.06170 | 0.04930 |
| 200 | high | shallow | 50 | 0.27950 | 0.18180 | **0.17430** | 0.18980 | 0.20220 |
| 200 | high | shallow | 200 | 0.16250 | 0.08940 | **0.08300** | 0.09600 | 0.10130 |
| 200 | high | shallow | 1000 | 0.07950 | 0.04000 | **0.03770** | 0.04930 | 0.04710 |
| 500 | medium | shallow | 50 | 0.09240 | 0.07400 | **0.06650** | 0.07630 | 0.07400 |
| 500 | medium | shallow | 200 | 0.04720 | 0.04000 | **0.03410** | 0.04260 | 0.04100 |
| 500 | medium | shallow | 1000 | 0.02330 | 0.02430 | **0.02030** | 0.02810 | 0.02610 |
| 1000 | medium | shallow | 50 | 0.09760 | 0.08670 | **0.07680** | 0.10110 | 0.08810 |
| 1000 | medium | shallow | 200 | 0.05150 | 0.04850 | **0.04180** | 0.05880 | 0.05180 |
| 1000 | medium | shallow | 1000 | nan | 0.03050 | **0.02460** | 0.03640 | 0.03320 |

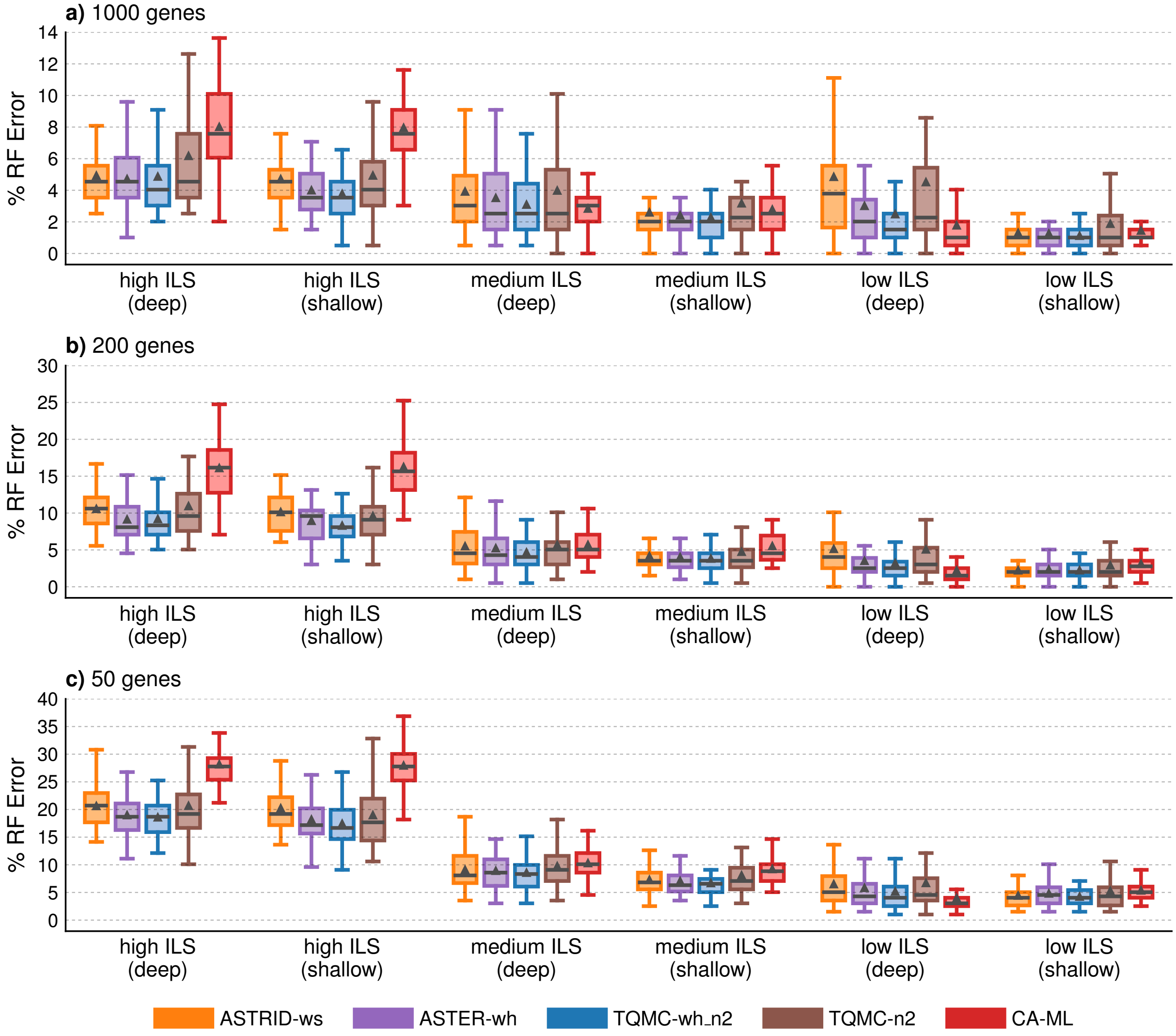

Figure S9: **Species tree error for ASTRAL-II data (abayes support) with varying levels of ILS.** Percent species tree (RF) error across replicates (bars represent medians; triangles represent means; outliers are not shown).

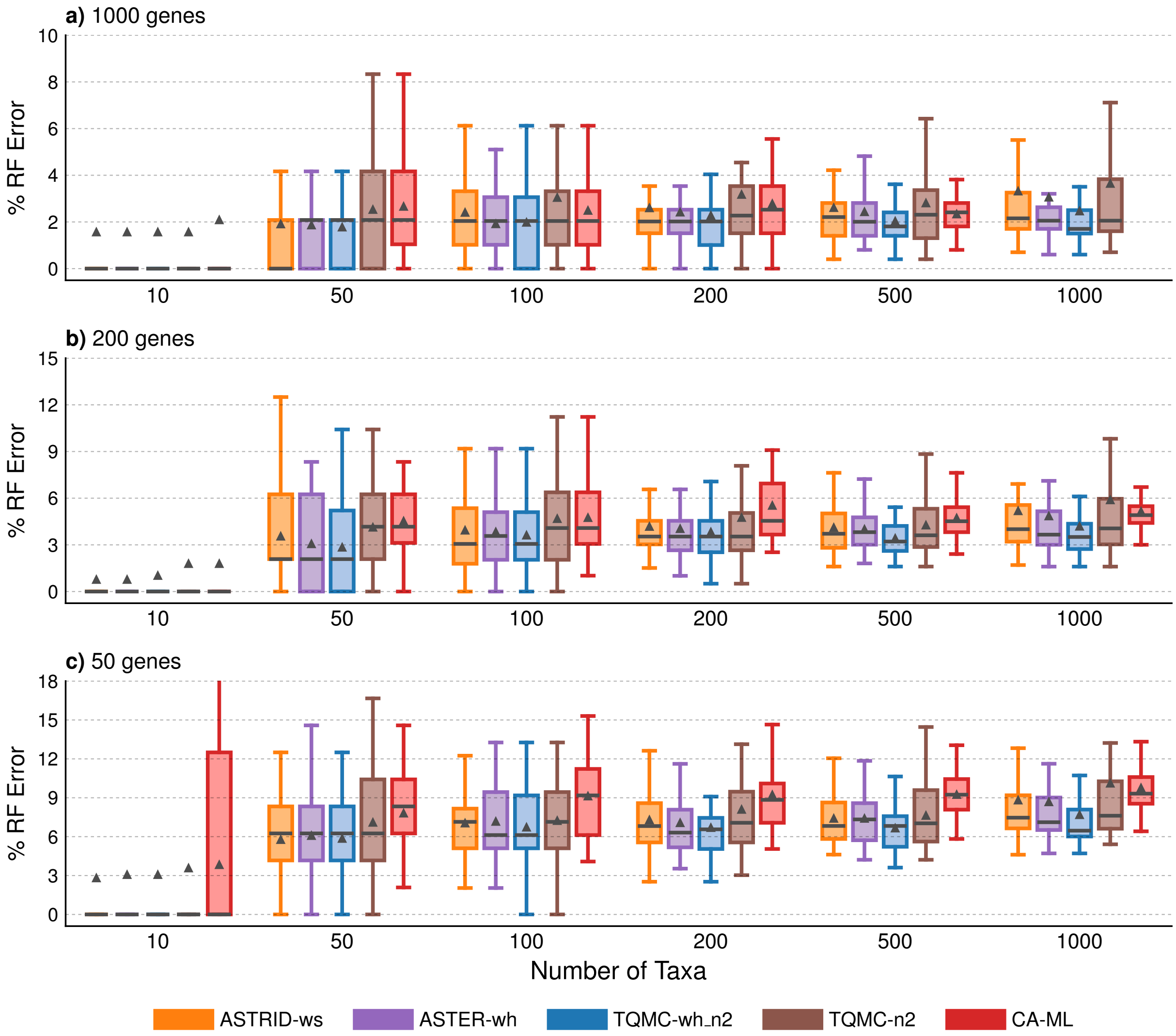

Figure S10: **Species tree error for ASTRAL-II data (abayes support) with varying numbers of taxa.** Percent species tree (RF) error across replicates (bars represent medians; triangles represent means; outliers are not shown). Note that CA-ML trees from original study were not available for 1000-taxon, 1000-gene data sets. Also, the results for 200-taxon data sets are duplicated from Figure S9

Table S18: **Testing for differences between TREE-QMC-wh_n2 vs CAML on the ASTRAL-II simulated data (abayes support) with varying levels of ILS.** BET is the number of replicates for which TREE-QMC-wh_n2 has lower species tree (RF) error and thus is better than CAML , WOR is the number of replicates for which wTREE-QMC-wh_n2 has higher RF error and thus is worse than CAML , and TIE is the number of replicates where the two methods tie. Significance is evaluated using paired, two-sided Wilcoxon signed-rank tests on the RF error rates. The symbols *, **, ***, ****, ***** indicate significance at $p < 0.5$, 0.005, and so on. MC indicates significance after Bonferroni correction, i.e., $p < 0.05$ / 99 = 5e-04 for the 99 tests made on the S100 data.

| **TREE-QMC-wh_n2 vs CAML** | | | | | | | | | |
|---|---|---|---|---|---|---|---|---|---|
| ILS Level | Speciation | # of genes | BET | WOR | TIE | p-val | sig | MC | note |
| low | deep | 50 | 12 | 31 | 7 | 0.001014695 | | ** | ( CAML better) |
| low | deep | 200 | 13 | 28 | 9 | 0.001628425 | | ** | ( CAML better) |
| low | deep | 1000 | 13 | 29 | 8 | 0.01314316 | | * | ( CAML better) |
| low | shallow | 50 | 36 | 9 | 5 | 5.632589e-06 | | **** | MC |
| low | shallow | 200 | 38 | 6 | 6 | 8.63588e-06 | | **** | MC |
| low | shallow | 1000 | 29 | 10 | 11 | 0.005499648 | | * | |
| medium | deep | 50 | 38 | 10 | 2 | 4.729318e-06 | | ***** | MC |
| medium | deep | 200 | 37 | 9 | 4 | 0.0004051359 | | *** | MC |
| medium | deep | 1000 | 24 | 19 | 7 | 0.9784243 | | | |
| medium | shallow | 50 | 44 | 3 | 3 | 3.538503e-12 | | ***** | MC |
| medium | shallow | 200 | 40 | 4 | 6 | 2.048751e-09 | | ***** | MC |
| medium | shallow | 1000 | 33 | 8 | 9 | 0.0003452434 | | *** | MC |
| high | deep | 50 | 50 | 0 | 0 | 1.776357e-15 | | ***** | MC |
| high | deep | 200 | 50 | 0 | 0 | 1.776357e-15 | | ***** | MC |
| high | deep | 1000 | 45 | 4 | 0 | 1.20962e-07 | | ***** | MC |
| high | shallow | 50 | 47 | 0 | 0 | 1.421085e-14 | | ***** | MC |
| high | shallow | 200 | 47 | 0 | 0 | 1.421085e-14 | | ***** | MC |
| high | shallow | 1000 | 46 | 0 | 1 | 2.842171e-14 | | ***** | MC |

Table S19: **Testing for differences between TREE-QMC-wh_n2 vs ASTER-h on the ASTRAL-II simulated data (abayes support) with varying levels of ILS.** BET is the number of replicates for which TREE-QMC-wh_n2 has lower species tree (RF) error and thus is better than ASTER-h , WOR is the number of replicates for which wTREE-QMC-wh_n2 has higher RF error and thus is worse than ASTER-h , and TIE is the number of replicates where the two methods tie. Significance is evaluated using paired, two-sided Wilcoxon signed-rank tests on the RF error rates. The symbols *, **, ***, ****, ***** indicate significance at $p < 0.5$, 0.005, and so on. MC indicates significance after Bonferroni correction, i.e., $p < 0.05$ / 99 = 5e-04 for the 99 tests made on the S100 data.

| **TREE-QMC-wh_n2 vs ASTER-h** | | | | | | | | | |
|---|---|---|---|---|---|---|---|---|---|
| ILS Level | Speciation | # of genes | BET | WOR | TIE | p-val | sig | MC | note |
| low | deep | 50 | 30 | 3 | 17 | 2.060551e-07 | | ***** | MC |
| low | deep | 200 | 21 | 11 | 18 | 0.02371259 | | * | |
| low | deep | 1000 | 26 | 2 | 22 | 1.696497e-05 | | **** | MC |
| low | shallow | 50 | 32 | 10 | 8 | 6.118376e-05 | | *** | MC |
| low | shallow | 200 | 21 | 13 | 16 | 0.1606247 | | | |
| low | shallow | 1000 | 17 | 5 | 28 | 0.03724957 | | * | |
| medium | deep | 50 | 24 | 14 | 12 | 0.05695275 | | | |
| medium | deep | 200 | 29 | 8 | 13 | 0.0001611809 | | *** | MC |
| medium | deep | 1000 | 22 | 11 | 17 | 0.0186442 | | * | |
| medium | shallow | 50 | 22 | 17 | 11 | 0.07322669 | | | |
| medium | shallow | 200 | 25 | 10 | 15 | 0.01056484 | | * | |
| medium | shallow | 1000 | 17 | 5 | 28 | 0.01173639 | | * | |
| high | deep | 50 | 25 | 21 | 4 | 0.2112505 | | | |
| high | deep | 200 | 24 | 21 | 5 | 0.9229223 | | | |
| high | deep | 1000 | 20 | 17 | 12 | 0.9970997 | | | |
| high | shallow | 50 | 29 | 13 | 5 | 0.01055602 | | * | |
| high | shallow | 200 | 29 | 12 | 6 | 0.01537982 | | * | |
| high | shallow | 1000 | 26 | 14 | 7 | 0.07923641 | | | |

Table S20: **Testing for differences between TREE-QMC-wh_n2 vs wASTRID on the ASTRAL-II simulated data (abayes support) with varying levels of ILS.** BET is the number of replicates for which TREE-QMC-wh_n2 has lower species tree (RF) error and thus is better than wASTRID , WOR is the number of replicates for which wTREE-QMC-wh_n2 has higher RF error and thus is worse than wASTRID , and TIE is the number of replicates where the two methods tie. Significance is evaluated using paired, two-sided Wilcoxon signed-rank tests on the RF error rates. The symbols *, **, ***, ****, ***** indicate significance at $p < 0.5$, 0.005, and so on. MC indicates significance after Bonferroni correction, i.e., $p < 0.05$ / 99 = 5e-04 for the 99 tests made on the S100 data.

| **TREE-QMC-wh_n2 vs wASTRID** | | | | | | | | | |
|---|---|---|---|---|---|---|---|---|---|
| ILS Level | Speciation | # of genes | BET | WOR | TIE | p-val | sig | MC | note |
| low | deep | 50 | 31 | 7 | 12 | 1.008374e-05 | | **** | MC |
| low | deep | 200 | 38 | 5 | 7 | 2.443812e-09 | | ***** | MC |
| low | deep | 1000 | 43 | 2 | 5 | 3.581135e-12 | | ***** | MC |
| low | shallow | 50 | 22 | 13 | 15 | 0.2696331 | | | |
| low | shallow | 200 | 12 | 20 | 18 | 0.2902047 | | | |
| low | shallow | 1000 | 14 | 7 | 29 | 0.2642612 | | | |
| medium | deep | 50 | 25 | 20 | 5 | 0.05764566 | | | |
| medium | deep | 200 | 33 | 10 | 7 | 6.837434e-05 | | *** | MC |
| medium | deep | 1000 | 30 | 6 | 14 | 4.754454e-05 | | **** | MC |
| medium | shallow | 50 | 30 | 13 | 7 | 0.001086662 | | ** | |
| medium | shallow | 200 | 26 | 13 | 11 | 0.02047486 | | * | |
| medium | shallow | 1000 | 26 | 12 | 12 | 0.01566447 | | * | |
| high | deep | 50 | 35 | 9 | 6 | 9.161746e-06 | | **** | MC |
| high | deep | 200 | 37 | 10 | 3 | 1.476171e-05 | | **** | MC |
| high | deep | 1000 | 29 | 13 | 7 | 0.1618871 | | | |
| high | shallow | 50 | 39 | 5 | 3 | 8.781171e-10 | | ***** | MC |
| high | shallow | 200 | 40 | 5 | 2 | 5.238405e-08 | | ***** | MC |
| high | shallow | 1000 | 33 | 8 | 6 | 1.492041e-06 | | ***** | MC |

Table S21: **Testing for differences between TREE-QMC-wh_n2 vs CAML on the ASTRAL-II simulated data (abayes support) with varying numbers of taxa.** BET is the number of replicates for which TREE-QMC-wh_n2 has lower species tree (RF) error and thus is better than CAML , WOR is the number of replicates for which wTREE-QMC-wh_n2 has higher RF error and thus is worse than CAML , and TIE is the number of replicates where the two methods tie. Significance is evaluated using paired, two-sided Wilcoxon signed-rank tests on the RF error rates. The symbols *, **, ***, ****, ***** indicate significance at $p < 0.5$, 0.005, and so on. MC indicates significance after Bonferroni correction, i.e., $p < 0.05$ / 99 = 5e-04 for the 99 tests made on the S100 data.

| **TREE-QMC-wh_n2 vs CAML** | | | | | | | | |
|---|---|---|---|---|---|---|---|---|
| # of taxa | # of genes | BET | WOR | TIE | p-val | sig | MC | note |
| 10 | 50 | 8 | 6 | 35 | 0.6334229 | | | |
| 10 | 200 | 4 | 2 | 43 | 0.53125 | | | |
| 10 | 1000 | 2 | 1 | 45 | 0.75 | | | |
| 50 | 50 | 28 | 9 | 10 | 0.0001925013 | *** | MC | |
| 50 | 200 | 29 | 12 | 6 | 0.0009076932 | ** | | |
| 50 | 1000 | 21 | 3 | 23 | 0.0007148981 | ** | | |
| 100 | 50 | 38 | 5 | 5 | 1.594049e-08 | ***** | MC | |
| 100 | 200 | 32 | 9 | 7 | 0.001508224 | ** | | |
| 100 | 1000 | 23 | 13 | 12 | 0.04959638 | * | | |
| 200 | 50 | 44 | 3 | 3 | 3.538503e-12 | ***** | MC | (dup) |
| 200 | 200 | 40 | 4 | 6 | 2.048751e-09 | ***** | MC | (dup) |
| 200 | 1000 | 33 | 8 | 9 | 0.0003452434 | *** | MC | (dup) |
| 500 | 50 | 46 | 3 | 1 | 8.881784e-14 | ***** | MC | |
| 500 | 200 | 42 | 7 | 1 | 3.823587e-09 | ***** | MC | |
| 500 | 1000 | 36 | 11 | 3 | 0.004434673 | ** | | |
| 1000 | 50 | 45 | 5 | 0 | 8.173611e-08 | ***** | MC | |
| 1000 | 200 | 42 | 8 | 0 | 1.173318e-06 | ***** | MC | |

Table S22: **Testing for differences between TREE-QMC-wh_n2 vs ASTER-h on the ASTRAL-II simulated data (abayes support) with varying numbers of taxa.** BET is the number of replicates for which TREE-QMC-wh_n2 has lower species tree (RF) error and thus is better than ASTER-h , WOR is the number of replicates for which wTREE-QMC-wh_n2 has higher RF error and thus is worse than ASTER-h , and TIE is the number of replicates where the two methods tie. Significance is evaluated using paired, two-sided Wilcoxon signed-rank tests on the RF error rates. The symbols *, **, ***, ****, ***** indicate significance at $p < 0.5$, 0.005, and so on. MC indicates significance after Bonferroni correction, i.e., $p < 0.05$ / 99 = 5e-04 for the 99 tests made on the S100 data.

| **TREE-QMC-wh_n2 vs ASTER-h** | | | | | | | | |
|---|---|---|---|---|---|---|---|---|
| # of taxa | # of genes | BET | WOR | TIE | p-val | sig | MC | note |
| 10 | 50 | 1 | 1 | 47 | 1 | | | |
| 10 | 200 | 0 | 1 | 48 | 1 | | | |
| 10 | 1000 | 0 | 0 | 48 | NA | NA | NA | |
| 50 | 50 | 11 | 8 | 28 | 0.4520187 | | | |
| 50 | 200 | 8 | 3 | 36 | 0.2265625 | | | |
| 50 | 1000 | 5 | 3 | 39 | 0.7265625 | | | |
| 100 | 50 | 17 | 8 | 23 | 0.0395084 | * | | |
| 100 | 200 | 15 | 7 | 26 | 0.1132421 | | | |
| 100 | 1000 | 8 | 10 | 30 | 0.67379 | | | |
| 200 | 50 | 22 | 17 | 11 | 0.07322669 | | | (dup) |
| 200 | 200 | 25 | 10 | 15 | 0.01056484 | * | | (dup) |
| 200 | 1000 | 17 | 5 | 28 | 0.01173639 | * | | (dup) |
| 500 | 50 | 38 | 10 | 2 | 1.885201e-07 | ***** | MC | |
| 500 | 200 | 40 | 6 | 4 | 5.277997e-09 | ***** | MC | |
| 500 | 1000 | 36 | 8 | 6 | 1.304655e-05 | **** | MC | |
| 1000 | 50 | 44 | 2 | 4 | 4.547474e-13 | ***** | MC | |
| 1000 | 200 | 38 | 4 | 8 | 4.101821e-10 | ***** | MC | |
| 1000 | 1000 | 42 | 2 | 4 | 2.273737e-11 | ***** | MC | |

Table S23: **Testing for differences between TREE-QMC-wh_n2 vs wASTRID on the ASTRAL-II simulated data (abayes support) with varying numbers of taxa.** BET is the number of replicates for which TREE-QMC-wh_n2 has lower species tree (RF) error and thus is better than wASTRID , WOR is the number of replicates for which wTREE-QMC-wh_n2 has higher RF error and thus is worse than wASTRID , and TIE is the number of replicates where the two methods tie. Significance is evaluated using paired, two-sided Wilcoxon signed-rank tests on the RF error rates. The symbols *, **, ***, ****, ***** indicate significance at $p < 0.5$, 0.005, and so on. MC indicates significance after Bonferroni correction, i.e., $p < 0.05$ / 99 = 5e-04 for the 99 tests made on the S100 data.

| **TREE-QMC-wh_n2 vs wASTRID** | | | | | | | | |
|---|---|---|---|---|---|---|---|---|
| # of taxa | # of genes | BET | WOR | TIE | p-val | sig | MC | note |
| 10 | 50 | 1 | 2 | 46 | 1 | | | |
| 10 | 200 | 0 | 1 | 48 | 1 | | | |
| 10 | 1000 | 0 | 0 | 48 | NA | NA | NA | |
| 50 | 50 | 12 | 13 | 22 | 0.8604026 | | | |
| 50 | 200 | 18 | 3 | 26 | 0.00435257 | ** | | |
| 50 | 1000 | 6 | 3 | 38 | 0.5078125 | | | |
| 100 | 50 | 21 | 14 | 13 | 0.1545381 | | | |
| 100 | 200 | 19 | 13 | 16 | 0.1799249 | | | |
| 100 | 1000 | 14 | 7 | 27 | 0.0765667 | | | |
| 200 | 50 | 30 | 13 | 7 | 0.001086662 | ** | | (dup) |
| 200 | 200 | 26 | 13 | 11 | 0.02047486 | * | | (dup) |
| 200 | 1000 | 26 | 12 | 12 | 0.01566447 | * | | (dup) |
| 500 | 50 | 35 | 13 | 2 | 0.0001402717 | *** | MC | |
| 500 | 200 | 41 | 7 | 2 | 4.245093e-08 | ***** | MC | |
| 500 | 1000 | 29 | 9 | 12 | 3.824975e-05 | **** | MC | |
| 1000 | 50 | 43 | 6 | 1 | 1.760259e-09 | ***** | MC | |
| 1000 | 200 | 46 | 3 | 1 | 7.94742e-12 | ***** | MC | |
| 1000 | 1000 | 39 | 6 | 3 | 6.072582e-10 | ***** | MC | |

#### 4.4 Results on biological data sets

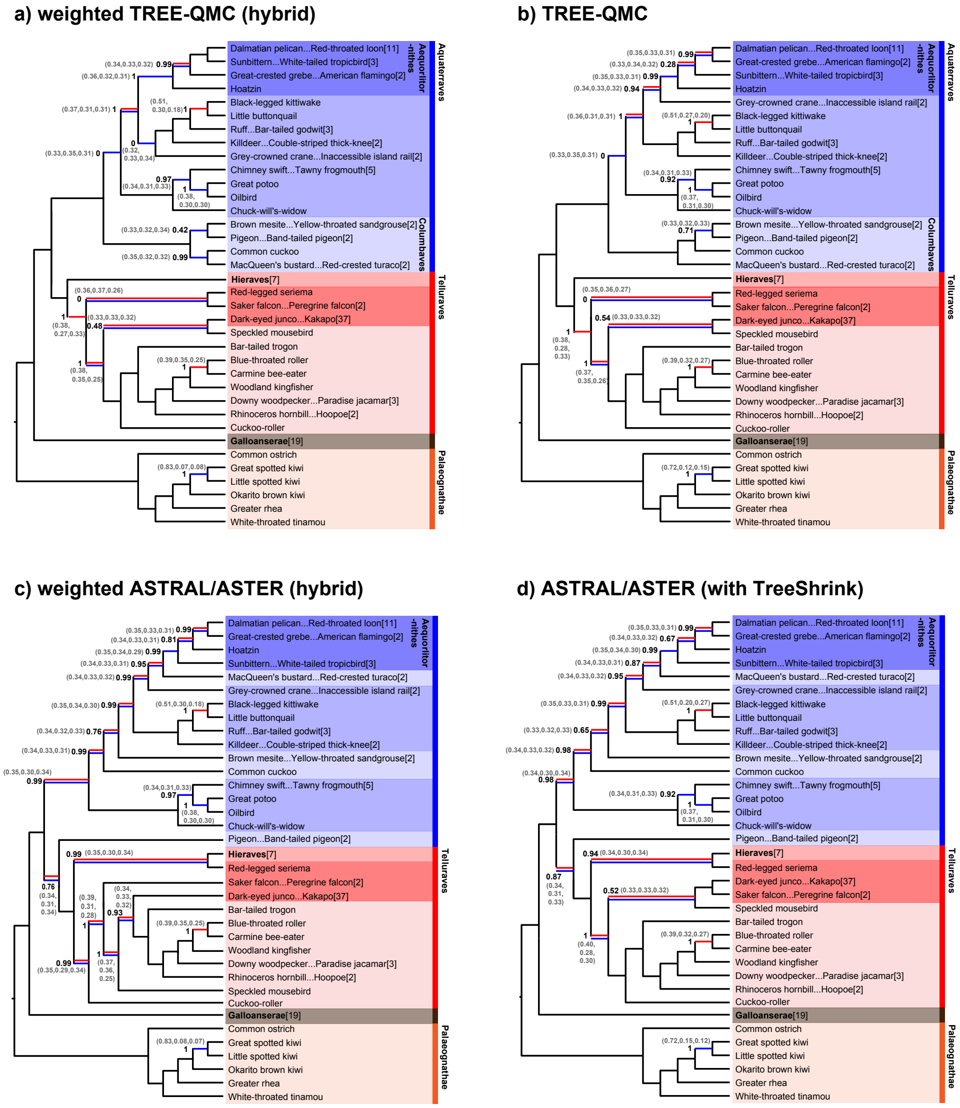

Figure S11: **Species trees estimated from Wu *et al.*, 2024 data** [8]**.** Subfigures (a)–(d) shows species tree we estimated on CDs and introns, continuing from the figure in the main text. For unweighted ASTRAL only (subfigure d), outlier taxa were removed via TreeShrink prior to species tree estimation. Red lines are differences with the NJst tree. Blue lines are differences with the RAxML tree. Branch support (i.e., ASTRAL's local posterior probability) is shown on branches where there are disagreements between the NJst and RAxML trees. QQS values for branch are in parentheses. QQS values are computed with hybrid weighted quartets for subfigures (a) and (c) and unweighted quartets (on input with taxon filtering with TreeShrink) except for subfigures (c) and (d).

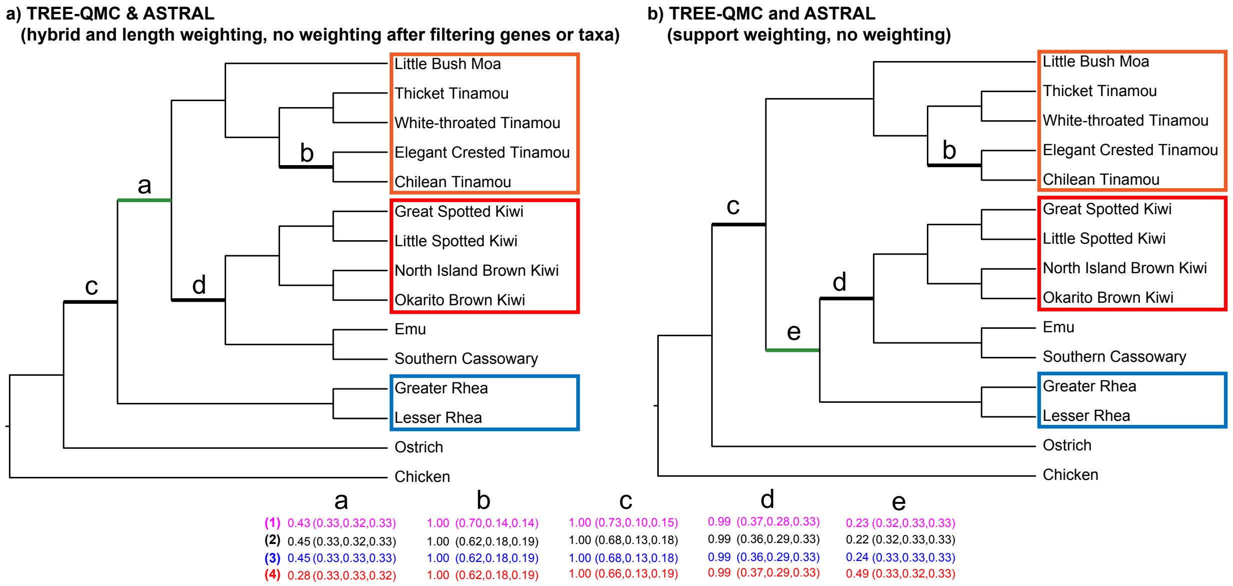

Figure S12: **Species trees estimated from Cloutier *et al.*, 2019 data** [8]**.** We used two types of data filtering: (1) removing the impacted taxa (Chicken and White-Throated Tinamou) from the 105 gene trees with homology errors and (2) these 105 gene trees entirely. Only two species trees were recovered by all methods. ASTRID and Asteroid also recovered the tree shown in subfigure (b) when using support, length, or no weighting (before or after either type of filtering). Branch support (ASTRAL's local posterior probability followed by the three normalized QQS values) is shown at bottom of the figure. Support was estimated in four ways: (1) hybrid quartet weighting (magenta), (2) no quartet weighting after filtering impacted taxa (black), (3) no quartet weighting after filtering gene trees (blue), (4) no quartet weighting (red).

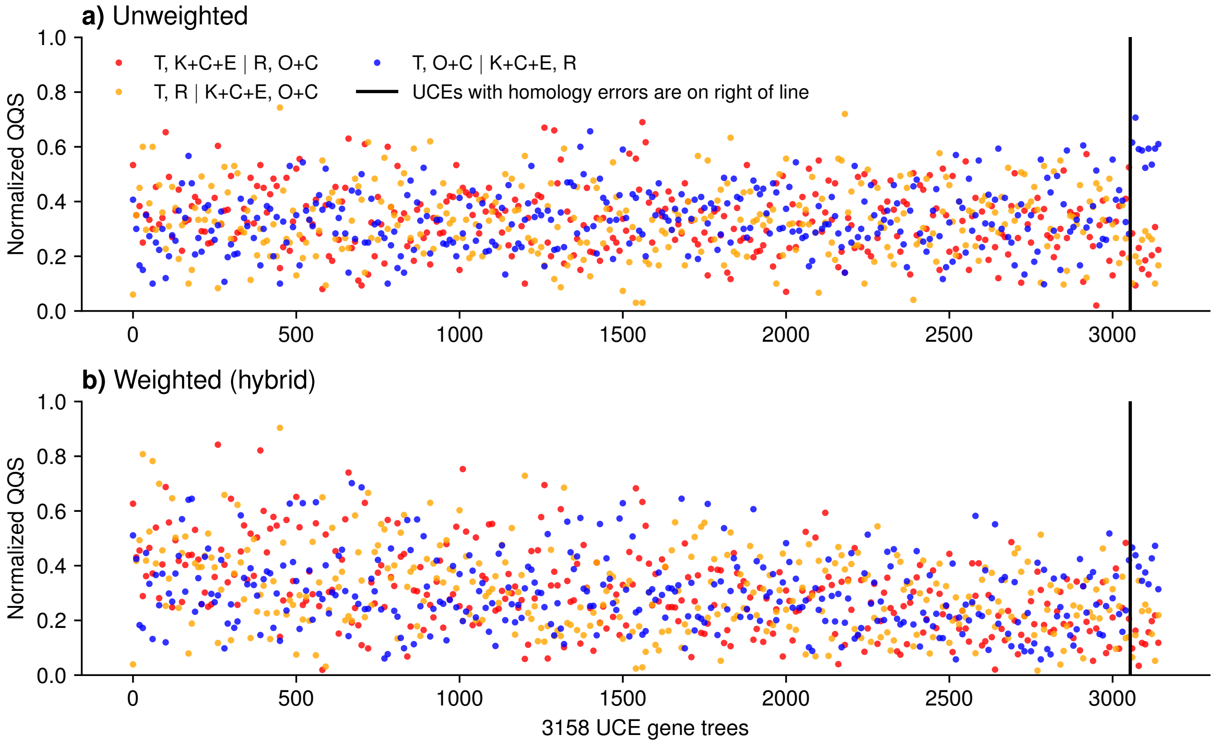

Figure S13: **Partitioned Coalescent Support (PCS) analysis of Cloutier *et al.*, 2019 data** [2]. The $x$-axis shows the 3,053 UCE gene trees sorted by mean abayes branch support, plus the 105 UCE gene trees with homology errors at the end. Subfigures (a) and (b) show normalized QQS values on the $y$-axis for unweighted and weighted (hybrid) quartets, respectively. Dots are averages across 10 gene trees (non-overlapping windows). The red, orange, and blue colors indicate the three possible resolutions of the focal branch. $T$ corresponds to four Tinamou species plus Little Bush Moa, $K+C+E$ corresponds to the four Kiwi species plus Southern Cassowary and Emu, and $R$ corresponds to the two Rhea species. $O+C$ indicates Ostrich plus Chicken. The homology errors cluster White-Throated Tinamou and Chicken, resulting in increased support for the related topology (blue).

### References

[1] Maria Anisimova, Manuel Gil, Jean-François Dufayard, Christophe Dessimoz, and Olivier Gascuel. Survey of Branch Support Methods Demonstrates Accuracy, Power, and Robustness of Fast Likelihood-based Approximation Schemes. *Systematic Biology*, 60(5):685–699, 05 2011.

[2] A. Cloutier, T. B. Sackton, P. Grayson, M. Clamp, A. J. Baker, and S. V. Edwards. Whole-genome analyses resolve the phylogeny of flightless birds (Palaeognathae) in the presence of an empirical anomaly zone. *Systematic Biology*, 68:937–955, 2019.

[3] Yunheng Han and Erin K Molloy. Improving quartet graph construction for scalable and accurate species tree estimation from gene trees. *Genome Research*, 33(7):1042–1052, July 2023.

[4] Baqiao Liu and Tandy Warnow. Weighted astrid: fast and accurate species trees from weighted internode distances. *Algorithms for Molecular Biology*, 18:6, 2023.

[5] Bui Quang Minh, Heiko A Schmidt, Olga Chernomor, Dominik Schrempf, Michael D Woodhams, Arndt von Haeseler, and Robert Lanfear. IQ-TREE 2: New Models and Efficient Methods for Phylogenetic Inference in the Genomic Era. *Molecular Biology and Evolution*, 37(5):1530–1534, 2020.

[6] Siavash Mirarab and Tandy Warnow. ASTRAL-II: coalescent-based species tree estimation with many hundreds of taxa and thousands of genes. *Bioinformatics*, 31(12):i44–i52, 2015.

[7] Benoit Morel, Tom A Williams, and Alexandros Stamatakis. Asteroid: a new algorithm to infer species trees from gene trees under high proportions of missing data. *Bioinformatics*, 2022. btac832.

[8] Shaoyuan Wu, Frank E. Rheindt, Jin Zhang, Jiajia Wang, Lei Zhang, Cheng Quan, Zhiheng Li, Min Wang, Feixiang Wu, Yanhua Qu, Scott V. Edwards, Zhonghe Zhou, and Liang Liu. Genomes, fossils, and the concurrent rise of modern birds and flowering plants in the late cretaceous. *Proceedings of the National Academy of Sciences*, 121(8):e2319696121, 2024.

[9] Chao Zhang and Siavash Mirarab. Weighting by gene tree uncertainty improves accuracy of quartet-based species trees. *Molecular Biology and Evolution*, 39(12):msac215, 2022.

[10] Chao Zhang, Maryam Rabiee, Erfan Sayyari, and Siavash Mirarab. ASTRAL-III: Polynomial time species tree reconstruction from partially resolved gene trees. *BMC Bioinformatics*, 19(6):153, 2018.
